# Robust latent-variable interpretation of *in vivo* regression models by nested resampling

**DOI:** 10.1101/703470

**Authors:** Alexander W. Caulk, Kevin A. Janes

## Abstract

Simple multilinear methods, such as partial least squares regression (PLSR), are effective at interrelating dynamic, multivariate datasets of cell–molecular biology through high-dimensional arrays. However, data collected *in vivo* are more difficult, because animal-to-animal variability is often high, and each time-point measured is usually a terminal endpoint for that animal. Observations are further complicated by the nesting of cells within tissues or tissue sections, which themselves are nested within animals. Here, we introduce principled resampling strategies that preserve the tissue-animal hierarchy of individual replicates and compute the uncertainty of multidimensional decompositions applied to global averages. Using molecular–phenotypic data from the mouse aorta and colon, we find that interpretation of decomposed latent variables (LVs) changes when PLSR models are resampled. Lagging LVs, which statistically improve global-average models, are unstable in resampled iterations that preserve nesting relationships, arguing that these LVs should not be mined for biological insight. Interestingly, resampling is less discriminatory for multidimensional regressions of *in vitro* data, suggesting it is unnecessary when replicate-to-replicate variance is low. Our work illustrates the challenges and opportunities in translating systems-biology approaches from cultured cells to living organisms. Nested resampling adds a straightforward quality-control step aiding the interpretability of *in vivo* regression models.

## INTRODUCTION

Modern biology and physiology demand rich, quantitative, time-resolved observations obtained by different methods^1^. To analyze such datasets, statistical “data-driven” modeling^2^ approaches have been productively deployed *in vitro* to examine network-level relationships between signal transduction and cell phenotype^3–9^. One class of models uses partial least squares regression (PLSR) to factorize data by the measured biological variables^10^. Linear combinations are iteratively extracted as latent variables (LVs) that optimize the covariation between independent and dependent datasets to enable input-output predictions. Highly multivariate data are efficiently modeled by a small number of LVs because of the mass-action kinetic processes underlying biological regulation^11^.

The success of PLSR at capturing biological function extends to nonlinear derivatives^12^ and structured multidimensional data arrays^13^ (tensors) from cell lines. By contrast, *in vivo* applications of PLSR have not gone beyond qualitative classification of inputs or outcomes^14–17^. The gap is unfortunate, because *in vivo* studies are the gold standard to compare phenotypes across species^18,19^, disease models^20,21^, and laboratories^22–26^. Animal surrogates can offer insight into the (patho)physiologic function of individual proteins, but interpreting the consequences of *in vivo* perturbations is complicated^27,28^. Applying PLSR quantitatively to *in vivo* data may better identify the underlying networks that, when perturbed, yield clinically relevant phenotypes.

For predictive modeling, there are many hurdles to using PLSR- and other LV-based approaches with *in vivo* data. Unlike spectroscopy (where PLSR originated^10^) or experiments in cultured cells, variation among *in vivo* replicates is often large even within inbred strains^29–31^, and this uncertainty does not get transmitted to standard models built from global averages. Including all replicates fixes the problem but creates others related to crossvalidation^32^ and the nesting of replicates in the study design^33^. *In vivo* data are typically grouped by replicate within a time point but are unpaired between time points, complicating model construction. An open question is whether the combinatorics of replicated, multivariate *in vivo* datasets can be tackled algorithmically within a multidimensional PLSR framework.

In this study, we apply computational statistics^34^ to the construction and interpretation of *in vivo* PLSR models built from multidimensional arrays (Fig. 1). Replicate-to-replicate uncertainty is propagated by resampling strategies that maintain the nesting relationships of the data acquisition. Nested resampling separates robust latent variables, which arise regardless of replicate configuration, from those that are statistically important in the global-average model but fragile upon resampling. Interpretations of robustness are more conservative when nested resampling is executed by bootstrapping (a leave-one-in approach) than by jackknifing (a leave-one-out approach). By contrast, neither is especially informative at discriminating latent variables when applied to a highly reproducible^35^ multidimensional dataset collected *in vitro*, bolstering the claims of earlier studies with cultured cells^3–9^. By leveraging the structure of multidimensional arrays, nested resampling provides a rapid numerical means to incorporate the uncertainty of *in vivo* observations into data-driven models without violating their mathematical assumptions.

**Figure 1.**
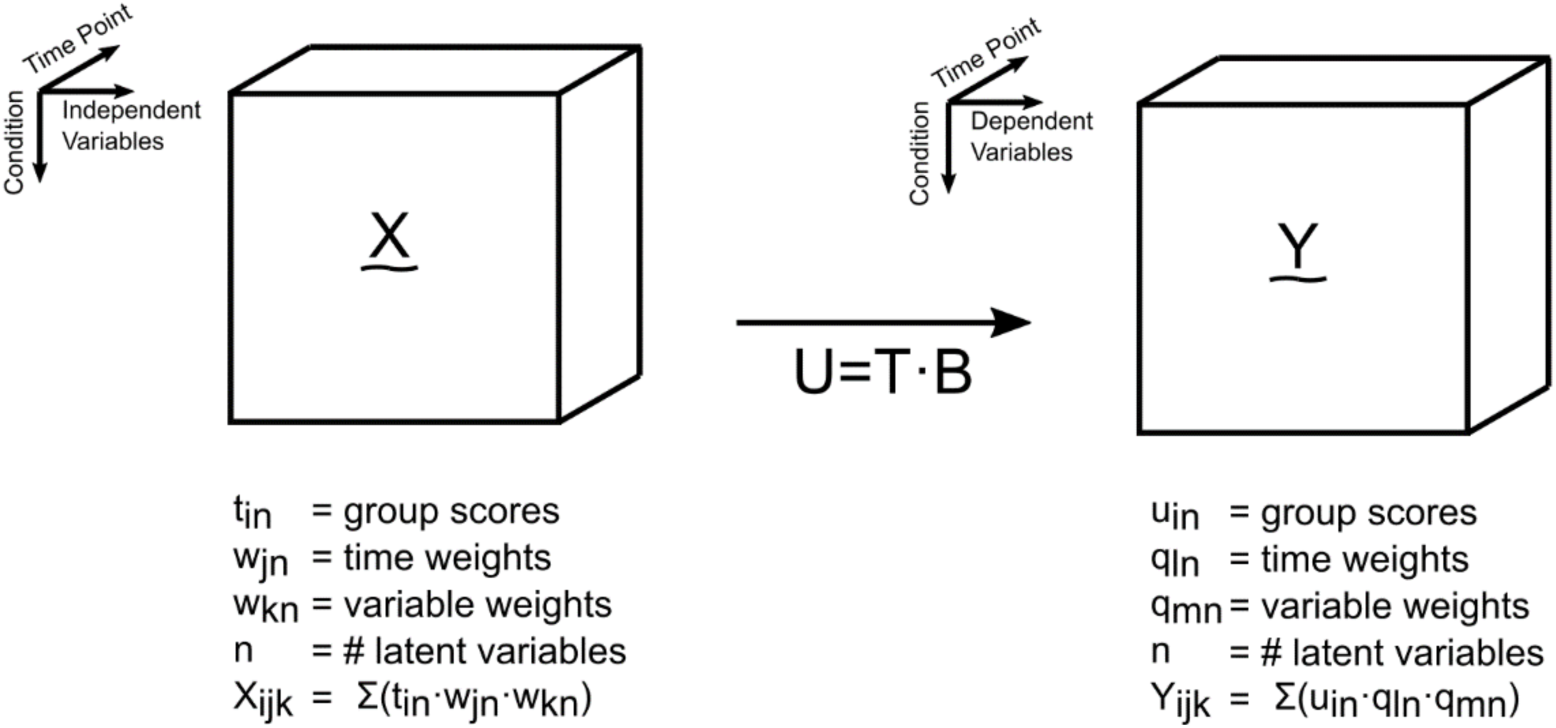
A three-mode structure efficiently models dynamic, multivariate data as multi-dimensional arrays. Data were organized as multidimensional arrays (**X** and **Y**) with mode 1 (indexed as i) delineating experimental conditions, mode 2 (indexed as j) delineating time course of experimental endpoints, and mode 3 (indexed as k) delineating variables measured for each experiment. Independent and dependent variables were selected according to the original datasets as detailed in Tables 1–3. PLSR derives condition scores (**U**) for the dependent data that are linearly related by regression coefficients (**B**) to the scores for the independent data (**T**).

## RESULTS

We sought an implementation of PLSR that robustly analyzes *in vivo* datasets comprised of temporal, multiparameter, and interrelated responses to perturbations. At the core of a PLSR model are its LVs (alternatively, principal components), which capture separable covariations among measured observations^2,36^. Interpreting LV features—for example, a “score” related to a condition or a “weight” (“loading”) related to a measured observation—is aided by computational randomization approaches that build hundreds of null models from the same data but without any true structure^13,37^. Scores and loadings that are similar between the null model and the actual model indicate data artifacts (biases, batch effects, etc.) that should not be used for hypothesis generation. Thus, by systematically building many alternative models, the randomization approach contextualizes the meaning of the true model.

We reasoned that a conceptually analogous approach might be useful for handling *in vivo* datasets that are inherently more variable than is typical for PLSR^31,32^. Iterative leave-one-out approaches such as jackknifing^38^ or crossvalidation^10^ are established approaches for omitting individual conditions during PLSR training and validation. Unexplored is whether there could be value in adapting such a strategy to replicates rather than conditions. To resample replicates by jackknifing, one biological replicate (*i.e.*, animal) is randomly omitted from each condition. All observations from that replicate are removed as a group to reflect the nesting relationships within the dataset. After one replicate is left out, averages are recalculated and a resampled PLSR model is built. The distribution of hundreds of jackknifed iterations indicates the extent to which the global-average model requires all of the data available.

Reciprocally, one could ask whether the global-average model is sufficiently reconstructed from any of the data by using bootstrapping instead of jackknifing. For bootstrap resampling, the nested observations from one biological replicate (animal) are randomly selected from each condition to build an n-of-one dataset that is modeled by PLSR. As with jackknife resampling, hundreds of iterations are compiled, yielding a bootstrap distribution of models and LVs based on a single instance of the data. Together, nested jackknife–bootstrap resampling should provide numerical estimates for the fragility and robustness of PLSR models constructed from global-average data with high inter-replicate variance.

The premise of nested resampling was tested in three contexts. First, we used a multidimensional dataset from Bersi *et al*.^39^ to build a new PLSR model, which warranted reinterpretation after nested resampling. We next tested general applicability of the approach by repurposing *in vivo* data from Lau *et al*.^14^ to construct a second multidimensional PLSR model for nested-resampling analysis. Last, we asked whether the same tools were similarly informative when applied to an existing multidimensional PLSR model from Chitforoushzadeh *et al*.^13^, which was calibrated with highly reproducible data from cultured cells. The results collectively support nested resampling as a useful complement to PLSR models applied to *in vivo* settings when biological variability is large.

### Nested resampling uncovers PLSR model fragilities missed by randomization

In the study by Bersi *et al*.^39^, *ApoE^−/−^* mice (used for their highly maladaptive hypertension-induced vascular remodeling^40^) were continuously administered Angiotensin II (AngII) and evaluated for enzymatic, cellular, and mechanical changes in four regions of aortic tissue (Table 1). Enzymatic–cellular (immuno)histology was collected at three time points and mechanical data at five time points over 28 days along with baseline controls (*N* = 2–7 animals; Fig. 2). For multidimensional PLSR modeling, data were separated by histological (input) and mechanical (output) data (Fig. 1) and standardized to predict mechanical metrics from histological and immunohistochemical data (see Methods). The working hypothesis of the model was that regionally disparate inflammatory and enzymatic changes in the aorta predictably drive differential changes in tissue mechanical properties.

**Table 1.**
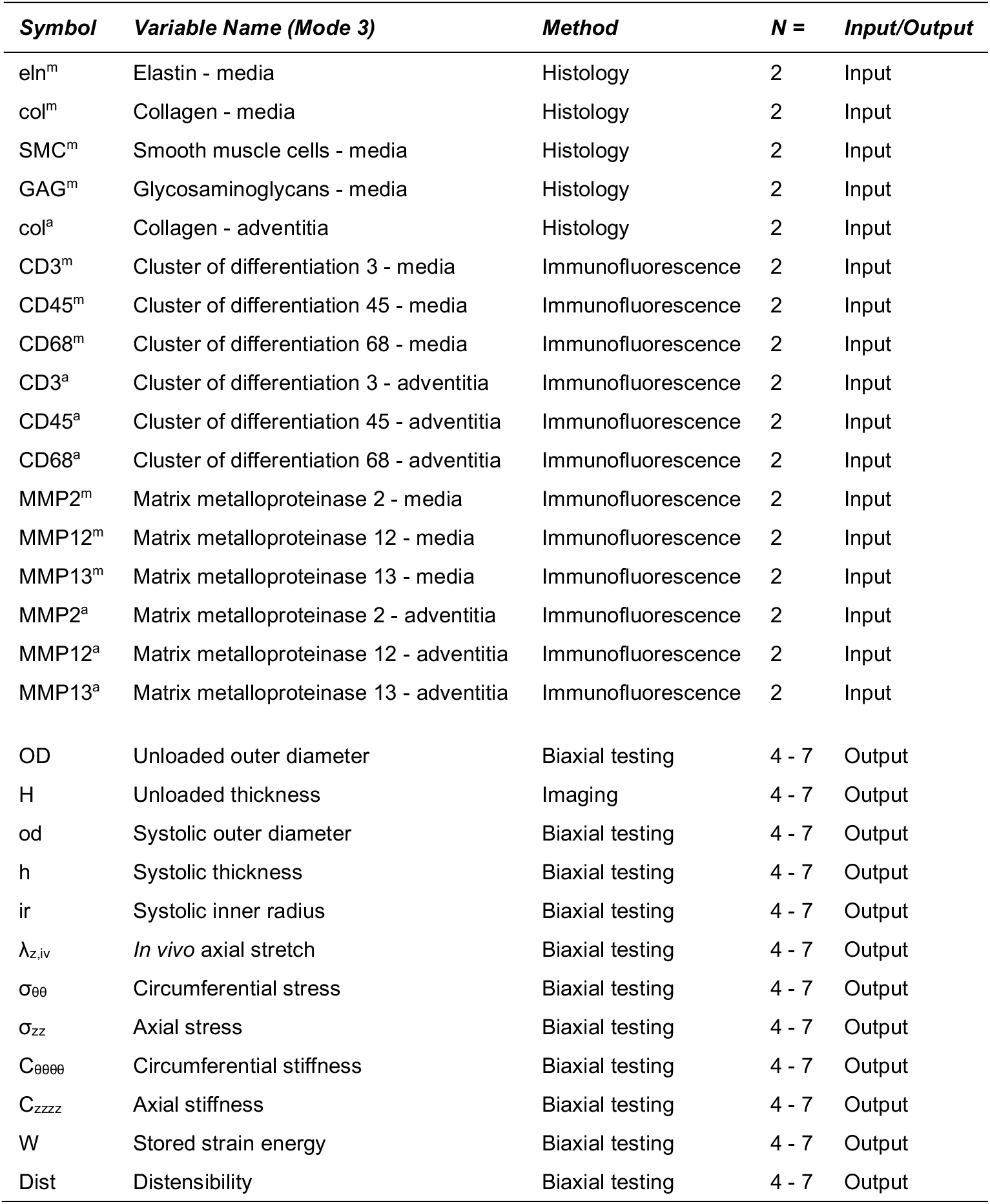
Symbols, metrics, methods of acquisition, and sample sizes per condition per time point (*N* =) for the PLSR model of Bersi *et al*.^39^. Histological stains used for matrix quantification include Elastica van Gieson (elastin – black stain), Movat’s Pentachrome (smooth muscle cells – red stain, GAGs – blue stain), and Picrosirius Red (collagen). Output samples were whole aortic sections from one mouse which were formalin-fixed after testing. Input samples were slides from output samples chosen for sectioning and staining based on their proximity to the mean thickness of their associated groups. Inputs were averages of three sections per slide.

| *Symbol* | *Variable Name (Mode 3)* | *Method* | *N =* | *Input/Output* |
| --- | --- | --- | --- | --- |
| eln<sup>m</sup> | Elastin - media | Histology | 2 | Input |
| col<sup>m</sup> | Collagen - media | Histology | 2 | Input |
| SMC<sup>m</sup> | Smooth muscle cells - media | Histology | 2 | Input |
| GAG<sup>m</sup> | Glycosaminoglycans - media | Histology | 2 | Input |
| col<sup>a</sup> | Collagen - adventitia | Histology | 2 | Input |
| CD3<sup>m</sup> | Cluster of differentiation 3 - media | Immunofluorescence | 2 | Input |
| CD45<sup>m</sup> | Cluster of differentiation 45 - media | Immunofluorescence | 2 | Input |
| CD68<sup>m</sup> | Cluster of differentiation 68 - media | Immunofluorescence | 2 | Input |
| CD3<sup>a</sup> | Cluster of differentiation 3 - adventitia | Immunofluorescence | 2 | Input |
| CD45<sup>a</sup> | Cluster of differentiation 45 - adventitia | Immunofluorescence | 2 | Input |
| CD68<sup>a</sup> | Cluster of differentiation 68 - adventitia | Immunofluorescence | 2 | Input |
| MMP2<sup>m</sup> | Matrix metalloproteinase 2 - media | Immunofluorescence | 2 | Input |
| MMP12<sup>m</sup> | Matrix metalloproteinase 12 - media | Immunofluorescence | 2 | Input |
| MMP13<sup>m</sup> | Matrix metalloproteinase 13 - media | Immunofluorescence | 2 | Input |
| MMP2<sup>a</sup> | Matrix metalloproteinase 2 - adventitia | Immunofluorescence | 2 | Input |
| MMP12<sup>a</sup> | Matrix metalloproteinase 12 - adventitia | Immunofluorescence | 2 | Input |
| MMP13<sup>a</sup> | Matrix metalloproteinase 13 - adventitia | Immunofluorescence | 2 | Input |
| OD | Unloaded outer diameter | Biaxial testing | 4 - 7 | Output |
| H | Unloaded thickness | Imaging | 4 - 7 | Output |
| od | Systolic outer diameter | Biaxial testing | 4 - 7 | Output |
| h | Systolic thickness | Biaxial testing | 4 - 7 | Output |
| ir | Systolic inner radius | Biaxial testing | 4 - 7 | Output |
| $\lambda_{z,iv}$ | *In vivo* axial stretch | Biaxial testing | 4 - 7 | Output |
| $\sigma_{\theta\theta}$ | Circumferential stress | Biaxial testing | 4 - 7 | Output |
| $\sigma_{zz}$ | Axial stress | Biaxial testing | 4 - 7 | Output |
| $C_{\theta\theta\theta\theta}$ | Circumferential stiffness | Biaxial testing | 4 - 7 | Output |
| $C_{zzzz}$ | Axial stiffness | Biaxial testing | 4 - 7 | Output |
| W | Stored strain energy | Biaxial testing | 4 - 7 | Output |
| Dist | Distensibility | Biaxial testing | 4 - 7 | Output |

**Figure 2.**
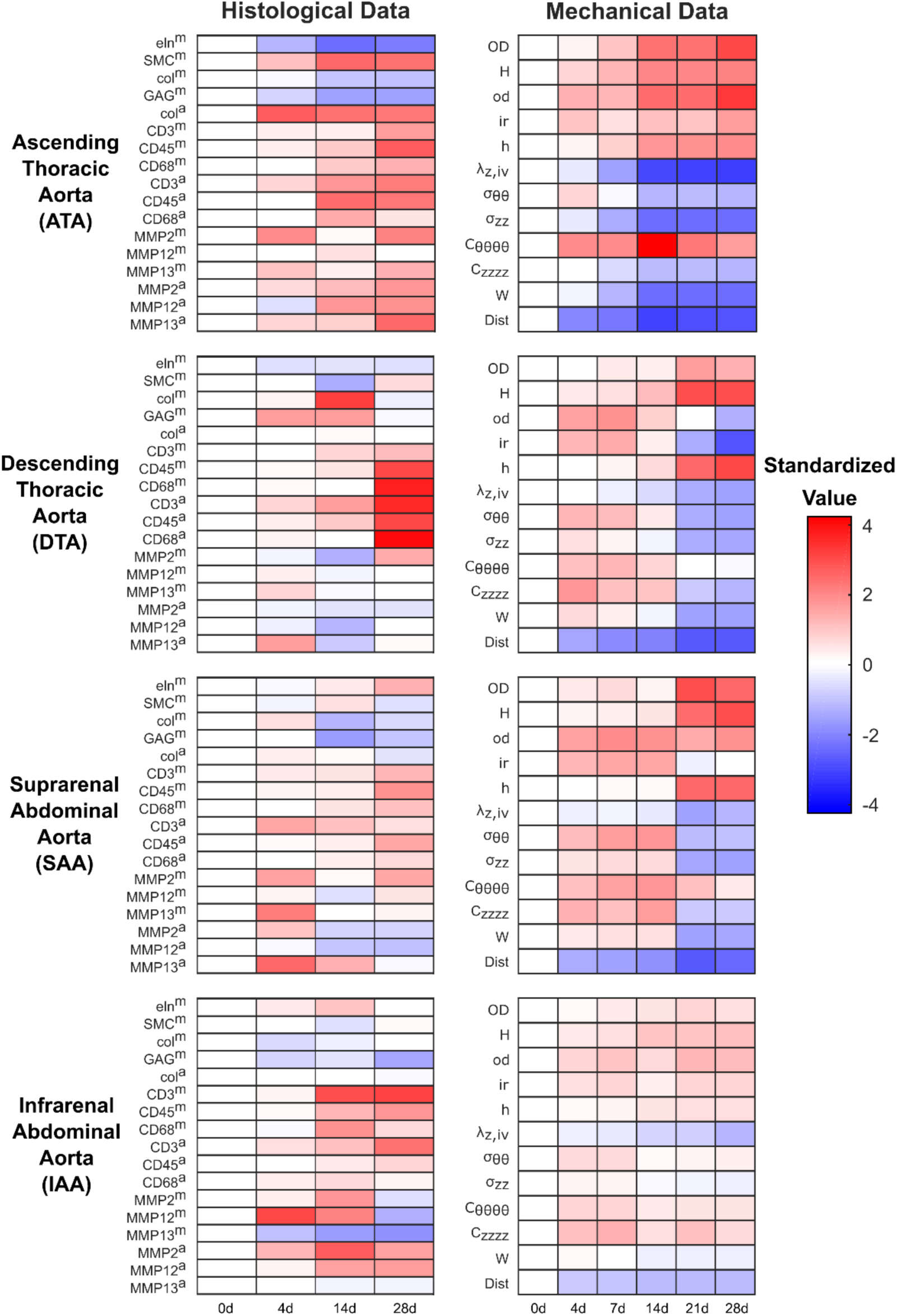
Time-resolved profiling of cellular infiltration, extracellular matrix production– turnover, and aortic geometry and mechanics during pharmacologically-induced hypertension. Mice were treated with AngII and tissue harvested at the indicated time points for subsequent histological and mechanical analysis (Table 1). Data are separated by independent (left) and dependent data (right) and aortic region (rows). Standardized differential changes (see Methods) from the 0-day baseline value are shaded red (increase) or blue (decrease).

LVs were iteratively defined for the multidimensional arrays by established approaches^13,41^, and the model root mean squared error (RMSE) of prediction was minimized with four LVs (Fig. 3a). By leave-one-out crossvalidation, we found that standardized predictions of the four-LV model were accurate to within ~75% of the measured result when averaged across all conditions (Fig. 3b), suggesting good predictive capacity. The four LVs of the multidimensional PLSR model thereby parse the regional, temporal, and molecular–cellular–mechanical covariations in the global-average dataset (Supplementary Fig. S1).

**Figure 3.**
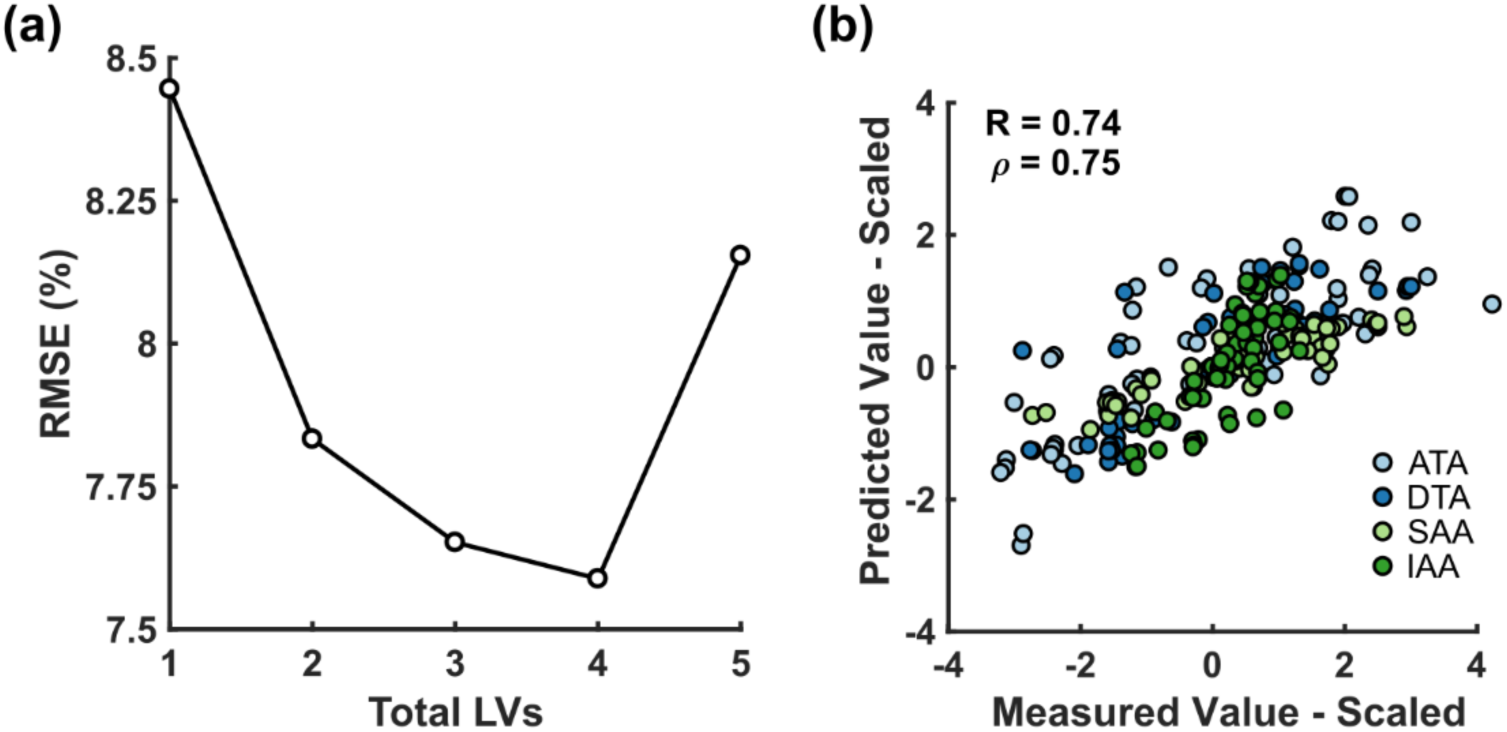
A four-component multidimensional PLSR model predicts AngII-induced evolution of aortic geometry and mechanics from matrix production and turnover, proteolytic enzyme expression, and inflammatory cell infiltrate. **(a)** Root mean squared error (RMSE) of cross-validated predictions is minimized with four LVs. **(b)** Pearson (R) and Spearman (*ρ*) correlation coefficients of the four-LV PLSR model for all aortic regions and time points. Cross-validated predictions were made by leaving out one entire aortic region at a time. ATA – ascending thoracic aorta, DTA – descending thoracic aorta, SAA – suprarenal abdominal aorta, IAA – infrarenal abdominal aorta.

For LV interpretation and hypothesis generation from the Bersi *et al*.^39^ dataset, we compared existing randomization methods^13,37^ to nested resampling. Across the four LVs, nearly all mechanical observations were weighted beyond the standard deviation of random null models (Fig. 4a,b), supporting interpretation of the weights. For example, inner radius was positively weighted on LV3 (ir; Fig. 4b) whereas thickness measures were negatively weighted on LV3 (H and h; Fig. 4b), suggesting that LV3 may discriminate aneurysmal dilatation, which predisposes to aortic dissection and rupture^42^, and fibrotic thickening, which predisposes to myocardial infarction and stroke via increased arterial stiffness^43^. However, interpretations changed when biological variability of the underlying *in vivo* data was considered through nested resampling (Fig. 4c–f). Both jackknifed and bootstrapped resampling suggested that LV3 and LV4 were too unstable to justify interpreting any parameters in these LVs (Fig. 4d,f). LV1 and LV2 yielded nonzero weights that were more robust, even retaining certain thickness and outer-diameter observations that were excluded by randomization (H, od, and OD; Fig. 4c). However, nested resampling revealed considerable uncertainty in the weights of LV3 and LV4 (Fig. 4d,f), arguing against any quantitative comparison of mechanical observations along these LVs. In contrast to standard performance metrics for PLSR (Fig. 3 and 4a,b), nested resampling provisioned the Bersi *et al*.^39^ model as fragile in its lagging LVs compared to the robustness of LV1 and LV2.

**Figure 4.**
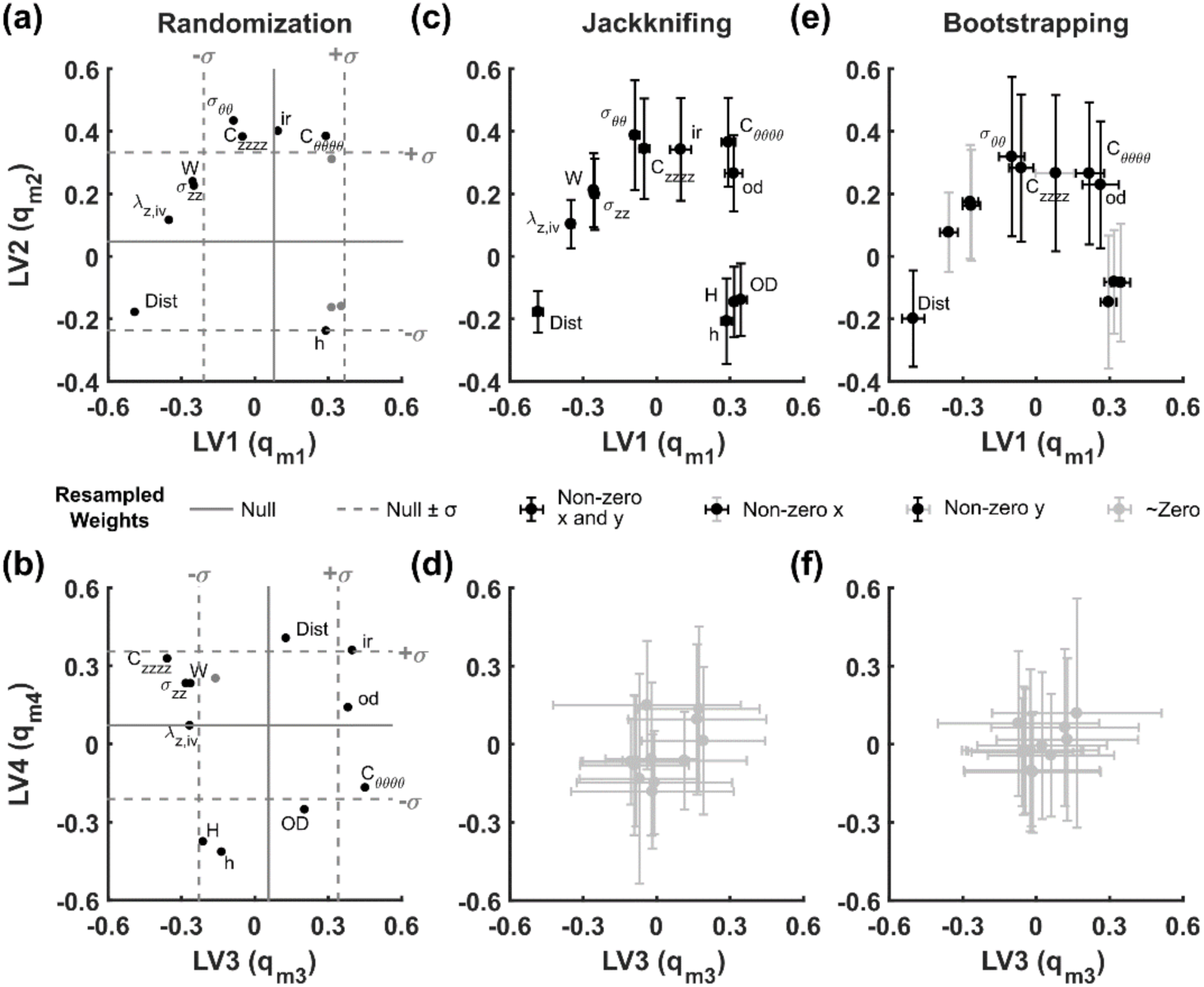
Resampling PLSR distinguishes robust dependent variable weights (q_mn_) in a four-LV model of AngII-induced hypertension. **(a, b)** Generation of a null PLSR model via data randomization of data to identify parameters of interest. Dependent variable weights (q_mn_) in the original PLSR model lying outside of a single standard deviation of the null PLSR model are labeled in black (see Table 1 for abbreviations). Solid gray lines denote the mean of *N* = 500 reshufflings within mode 1 (*i.e.*, time and measured variables were shuffled within each aortic region). Dotted-gray lines denote mean ± standard deviation of weights. **(c, d)** Replicate resampling (*N* = 500) by jackknifing (“leave one out”) changes confidence of predictions for parameters compared with randomization. Black dots denote variable weights with error bars that do not intersect with zero (*i.e.*, parameters weight consistently in a single region). Gray error bars denote errors that intersect with zero. **(e, f)** Replicate resampling (*N* = 500) by bootstrapping (“leave one in”) decreases confidence of parameters compared to jackknifing and yields no significant identifications in LV3 or LV4. Color delineations are identical to those in (c, d). The top row depicts results for LV1 and LV2, and the bottom row depicts results for LV3 and LV4.

One possible explanation for such high uncertainty is that some resampled models might switch the sign of an LV weight together with the associated LV score, which mutually offset as a degenerate solution (Fig. 1). We accounted for sign switching by looking for symmetric bimodal distributions about zero and flipping signs to the dominant mode when switching was evident. Some bimodal scores were asymmetrically distributed with a near-zero mode (*e.g.*, the distribution of LV1–LV2 scores for the DTA condition; Supplementary Fig. S2), indicating that their LV assignments were heavily dependent on the resampling iteration. For LV3 and LV4, however, the distribution of scores was broad among resampling replicates and mostly indistinguishable from zero (Supplementary Fig. S2). Uncertainty in the trailing LVs may stem from model iterations requiring less than 4 LVs to explain the variance in that iteration. The analysis further supports that the lagging LVs of this model do not contain prevailing trends in the data but instead capture a specific replicate configuration of the animals used.

### Data pairing does not significantly alter results of nested resampling

In the Bersi *et al*.^39^ study, inbred animals sacrificed at several time points were doubly used to collect enzymatic–cellular histology (X) and mechanical data (Y; Fig. 1). Possibly, the paired animal-by-animal covariation of histology and mechanics was greater than the condition-wide averages. We sought to evaluate the relative importance of within-animal pairing between independent and dependent datasets by applying nested resampling. To do so, we built a second PLSR model using only the time points with paired enzymatic–cellular and mechanical data: 0, 4, 14, and 28 days (Fig. 2). For the second model, resampling was coupled between X and Y to retain the paired information of each animal selected by bootstrapping. The interpretation of bootstrap-resampled time weights for the paired model was then compared with the original unpaired model to determine if conclusions were fundamentally different.

We found that the LV1–LV2 time weights obtained by paired sampling were indistinguishable from those obtained by unpaired sampling (Fig. 5, upper). Relative to their corresponding global-average model, both analyses indicated that the dynamics associated with LV1 and LV2 were robust, consistent with the prior assessment of mechanical weights for these LVs (Fig. 4). Histological time weights were similarly reliable for LV3 and LV4, but mechanical time weights were highly variable and largely overlapping with zero (Fig. 5, lower). No statistically significant differences were identified between paired and unpaired time weights in LV3 or LV4 (*p* > 0.25 following two-way ANOVA with Tukey’s post-hoc test for differences between paired–unpaired or independent–dependent time weights), indicating that pairing does not add statistical power to the trailing LVs for this dataset. More generally, the analysis suggests that unpaired *in vivo* designs may be sufficient for nested resampling to assess the stability of model components.

**Figure 5.**
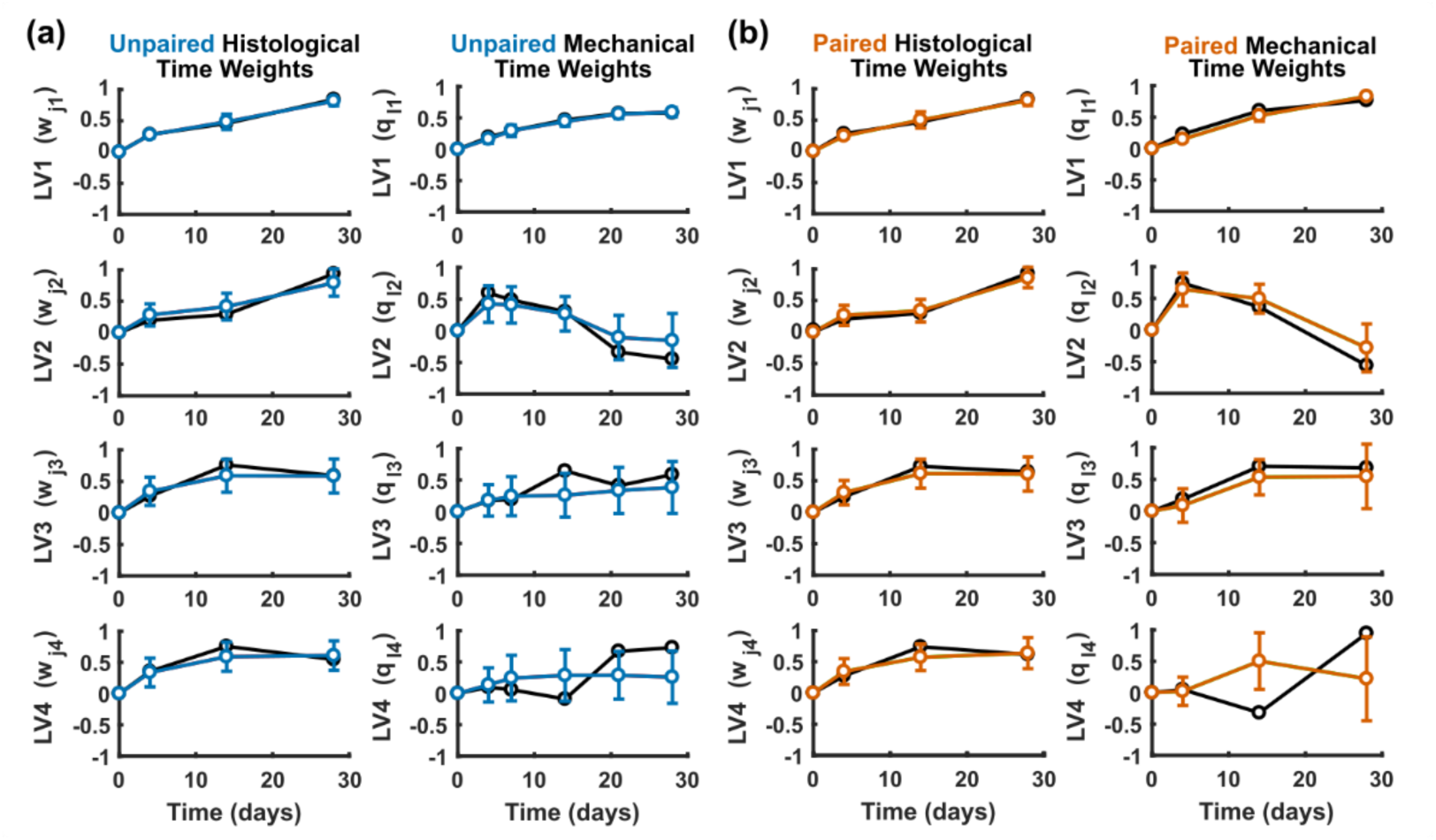
Bootstrapping PLSR with paired data shows similar performance to bootstrapping with unpaired data. Time weights (w_jn_, q_ln_) from a PLSR model using **(a)** unpaired (blue) and **(b)** paired (orange) bootstrapping of histological and biomechanical data were generated 500 times for unpaired and paired sampling each. Note that paired sampling required omission of the 7 and 21 day time points in the dependent variables because histological data were not collected for those time points. Paired data were available for only two samples per aortic region and time point, both of which were chosen based on the proximity of the thickness value to the mean thickness value for the corresponding region and time point.

### Generality of nested resampling to other multidimensional *in vivo* and *in vitro* datasets

The LV fragilities revealed by nested resampling could be specific to the Bersi *et al*.^39^ dataset. We thus sought another *in vivo* study comprised of multiple molecular–cellular measurements, time points, and animals where nested replicate information could be recovered confidently. Raw data was obtained from Lau *et al*.^14^, who examined the molecular and cellular inflammatory response of the small intestine to the cytokine tumor necrosis factor α (TNFα). Animals (*N* = 5) were administered one of two doses of TNFα and sacrificed at one of six time points after administration. From each animal, two intestinal regions were analyzed for signaling by Luminex phosphoproteomics, cell proliferation by immunohistochemistry, and overall cell death by western blotting (Table 2). The data were used previously to classify cell-fate responses^14^—we asked here whether cell proliferation and death were predicted quantitatively from the time-resolved phosphoproteomic observations. If so, then nested resampling could address how robust or fragile those predictions were to the animals included.

**Table 2.** Symbols, metrics, methods of acquisition, and sample sizes per condition per time point (*N* =) for the PLSR model of Lau *et al*.^14^ All input and output samples represent mice per time point and one intestinal segment each. qWB – Quantitative western blotting, IHC – Immunohistochemistry.

| *Symbol* | *Variable Name (Mode 3)* | *Marker* | *Method* | *N =* | *Input/Output* |
| --- | --- | --- | --- | --- | --- |
| pIκ $\beta$ $\alpha$ | Inhibitor of nuclear factor $\kappa$ β-α | Phospho Ser<sup>32/36</sup> | Bio-Plex | 5 | Input |
| pJnk | c-Jun N-terminal kinase | Phospho Thr<sup>183</sup>/Tyr<sup>185</sup> | Bio-Plex | 5 | Input |
| pMek1 | MAPK and ERK kinase 1 | Phospho Ser<sup>217/221</sup> | Bio-Plex | 5 | Input |
| pErk1/2 | Extracellular signal-related kinase 1/2 | Phospho Thr<sup>202</sup>/Tyr<sup>204</sup> (1), Thr<sup>185</sup>/Tyr<sup>187</sup> (2) | Bio-Plex | 5 | Input |
| pRsk | Ribosomal S6 kinase | Phospho Thr<sup>359</sup>/Ser<sup>363</sup> | Bio-Plex | 5 | Input |
| pp38 | p38 mitogen-activated protein kinase | Phospho Thr<sup>180</sup>/Tyr<sup>182</sup> | Bio-Plex | 5 | Input |
| pc-Jun | c-Jun | Phospho Ser<sup>63</sup> | Bio-Plex | 5 | Input |
| pAtf2 | Activating transcription factor 2 | Phospho Thr<sup>71</sup> | Bio-Plex | 5 | Input |
| pAkt | Akt/Protein kinase B | Phospho Ser<sup>473</sup> | Bio-Plex | 5 | Input |
| pS6 | Ribosomal protein S6 | Phospho Ser<sup>235/236</sup> | Bio-Plex | 5 | Input |
| pStat3<sup>S727</sup> | Signal transducer and activator of transcription 3 | Phospho Ser<sup>727</sup> | Bio-Plex | 5 | Input |
| pStat3<sup>Y705</sup> | Signal transducer and activator of transcription 3 | Phospho Tyr<sup>705</sup> | Bio-Plex | 5 | Input |
| cc3 | Cleaved caspase 3 | Cleaved levels | qWB | 5 | Output |
| ph3 | Phosphorylated histone 3 | Number positive cells | IHC | 5 | Output |

We organized and standardized the data (Supplementary Fig. S3), building a single PLSR model of the global averages along with 500 null models by randomization. For the Lau *et al*.^14^ dataset, a three-LV model was optimal and yielded good predictive accuracy (Fig. 6). LV1 of the global-average model did not discriminate between tissues or outcomes, but LV2 separated cell proliferation (ph3) vs. death (cc3) readouts and LV3 stratified duodenal vs. ileal segments of the intestine (Supplementary Fig. S4). Furthermore, randomization suggested that the ph3–cc3 distinction along LV2 was far outside chance expectation (Fig. 7a, left). Nested resampling, however, revealed a pronounced fragility of output weights when accounting for inter-replicate variability. Both jackknifing and bootstrapping eliminated any discrimination along LV2 (Fig. 7a, middle and right), undermining model interpretations based on it. Similarly, the time-dependent behavior associated with LV2 and LV3 (Fig. 7b) mostly reverted to near zero after bootstrap resampling (Fig. 7c). Therefore, as with the Bersi *et al*.^39^ study, the lagging components of this multidimensional PLSR model capture *in vivo* replicate instabilities instead of salient trends in the data.

**Figure 6.**
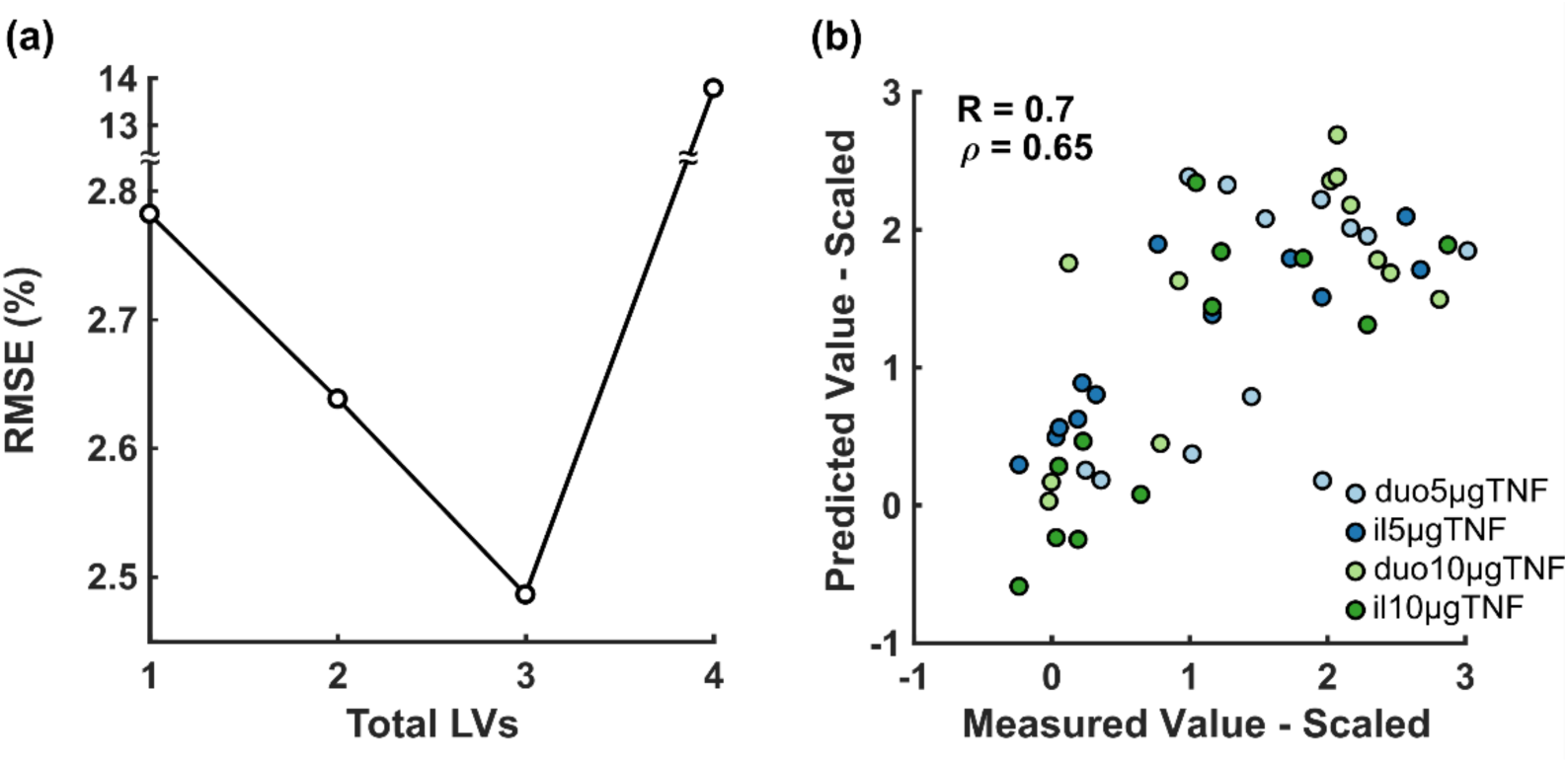
A three-component multidimensional PLSR model predicts TNFα-induced apoptosis and proliferation of intestinal cells from cell signaling in the duodenum and ileum. **(a)** Root mean squared error (RMSE) of cross-validated predictions is minimized with three LVs. **(b)** Pearson (R) and Spearman (*ρ*) correlation coefficients of the three-LV PLSR model for all intestinal regions and time points. Cross-validated predictions were made using the leave-one-out approach. duo – duodenum, il – ileum, 5µgTNF – 5 µg TNFα treatment, 10µgTNF – 10 µg TNFα treatment.

**Figure 7.**
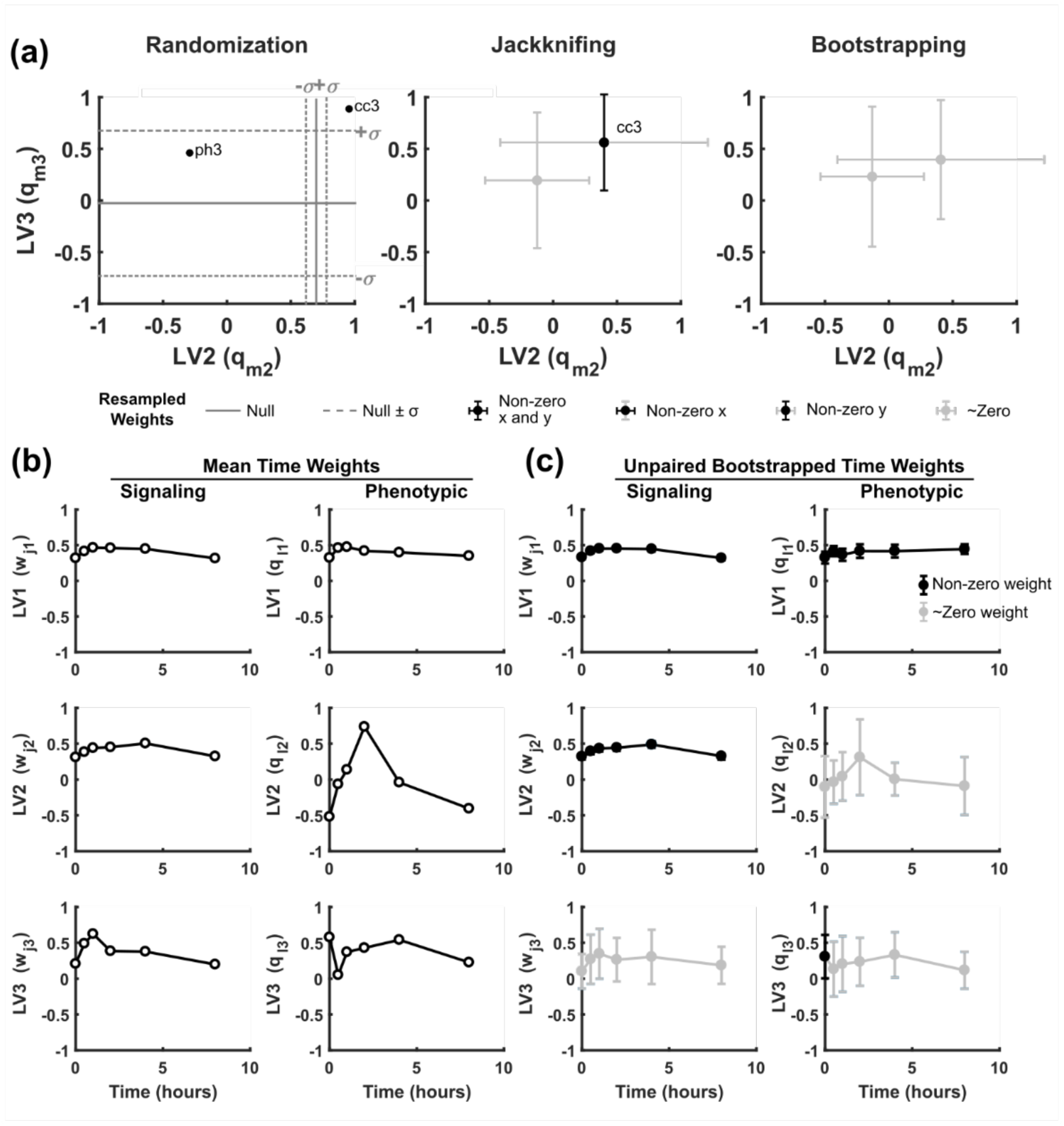
Bootstrapping PLSR of a second *in vivo* dataset reveals poor repeatability in trailing LVs. **(a)** Dependent variable weights (q_mn_) for LV2 vs. LV3 following randomization, jackknifing, and bootstrapping. LV1 is omitted for clarity. Graphs are labeled as in Fig. 4. **(b)** Time weights for the global-average model delineating temporal behaviors of each LV. **(c)** Bootstrapped time weights (*N* = 500) show good agreement with the mean dataset on LV1 and LV3 with less agreement on LV2. Data are presented as mean ± standard deviation, with black markers indicating error bars that do not intersect with zero and gray markers indicating error bars that intersect with zero.

It is possible that nested resampling excludes lagging LVs in any multidimensional dataset irrespective of its origin. To determine if fragility is tied to the higher biological variability of *in vivo* datasets, we reassessed an earlier multidimensional PLSR model built from global averages of *in vitro* measurements. The model of Chitforoushzadeh *et al*.^13^ predicts gene-expression cluster dynamics from intracellular signaling in a colon-cancer cell line stimulated with combinations of cytokines and growth factors^3,35,44^. Cell extracts (*N* = 2–6) were collected at three or 13 time points and measured transcriptomically by microarray or for signaling by various methods (Table 3). The prior hypothesis was that quantitative predictions of gene-expression dynamics would uncover novel upstream signaling regulators of transcriptional programs^13^.

**Table 3.** Symbols, metrics, methods of acquisition, and sample sizes per condition per time point (*N* =) for the PLSR model of Chitforoushzadeh *et al*.^13^. All input and output data represent cell extracts per time point. Ab – antibody, µ-array – microarray, qWB – Quantitative western blotting, CLICK – Cluster Identification via Connectivity Kernels.

| *Symbol* | *Variable Name (Mode 3)* | *Marker* | *Method* | *N =* | *Input/Output* |
| --- | --- | --- | --- | --- | --- |
| ERK | Extracellular signal-related kinase | Kinase activity | Kinase assay | 3 - 6 | Input |
| Akt | Akt/Protein kinase B | Kinase activity | Kinase assay | 3 - 6 | Input |
| pAkt<sub>Ab</sub> | Akt/Protein kinase B | Phospho Ser<sup>473</sup> | Ab μ-array | 3 - 6 | Input |
| pAkt<sub>WB</sub> | Akt/Protein kinase B | Phospho Ser<sup>473</sup> | qWB | 3 - 6 | Input |
| tAkt | Akt/Protein kinase B | Total amount | Ab μ-array | 3 - 6 | Input |
| ptAkt | Akt/Protein kinase B | Phospho/total ratio | Ab μ-array | 3 - 6 | Input |
| JNK1 | Jun N-terminal kinase 1 | Kinase activity | Kinase assay | 3 - 6 | Input |
| IKK | IκB kinase | Kinase activity | Kinase assay | 3 - 6 | Input |
| MK2 | MAP kinase-activated protein kinase 2 | Kinase activity | Kinase assay | 3 - 6 | Input |
| pMEK | MAPK and ERK kinase 1 | Phospho Ser<sup>217/221</sup> | qWB | 3 - 6 | Input |
| pFKHR | Forkhead in rhabdomyosarcoma | Phospho Ser<sup>256</sup> | qWB | 3 - 6 | Input |
| pIRS1<sub>636</sub> | Insulin receptor substrate 1 | Phospho Ser<sup>636</sup> | qWB | 3 - 6 | Input |
| pIRS1<sub>896</sub> | Insulin receptor substrate 1 | Phospho Tyr<sup>896</sup> | qWB | 3 - 6 | Input |
| proC8 | Caspase-8 | Zymogen amount | qWB | 3 - 6 | Input |
| cc8 | Caspase-8 | Cleaved amount | qWB | 3 - 6 | Input |
| proC3 | Caspase-3 | Zymogen amount | qWB | 3 - 6 | Input |
| pEGFR | Epidermal growth factor receptor | Phospho Tyr<sup>1068</sup> | Ab μ-array | 3 - 6 | Input |
| tEGFR | Epidermal growth factor receptor | Total amount | Ab μ-array | 3 - 6 | Input |
| ptEGFR | Epidermal growth factor receptor | Phospho/total ratio | Ab μ-array | 3 - 6 | Input |
| c1 - c9 | Gene clusters 1-9 | Transcription level | μ-array + CLICK | 2 | Output |

After obtaining the original dataset and confirming the nested replicate structure, we modeled the mean dataset (Supplementary Fig. S5) standardized as before^13^. The global-average model was optimally decomposed with four LVs, and randomizing 500 null models reproduced all the meaningfully weighted parameters (*e.g.*, gene-cluster weights) described in the original study (Fig. 8a,b). Remarkably, when nested resampling was applied to this PLSR model, the conclusions were largely unaltered. Cluster weights were retained in ~90% of LV2 and LV3 and even ~56% of LV4 (Fig. 8c–f), bolstering prior interpretations of this PLSR model along with others built upon highly reproducible *in vitro* data^3–9,13^.

**Figure 8.**
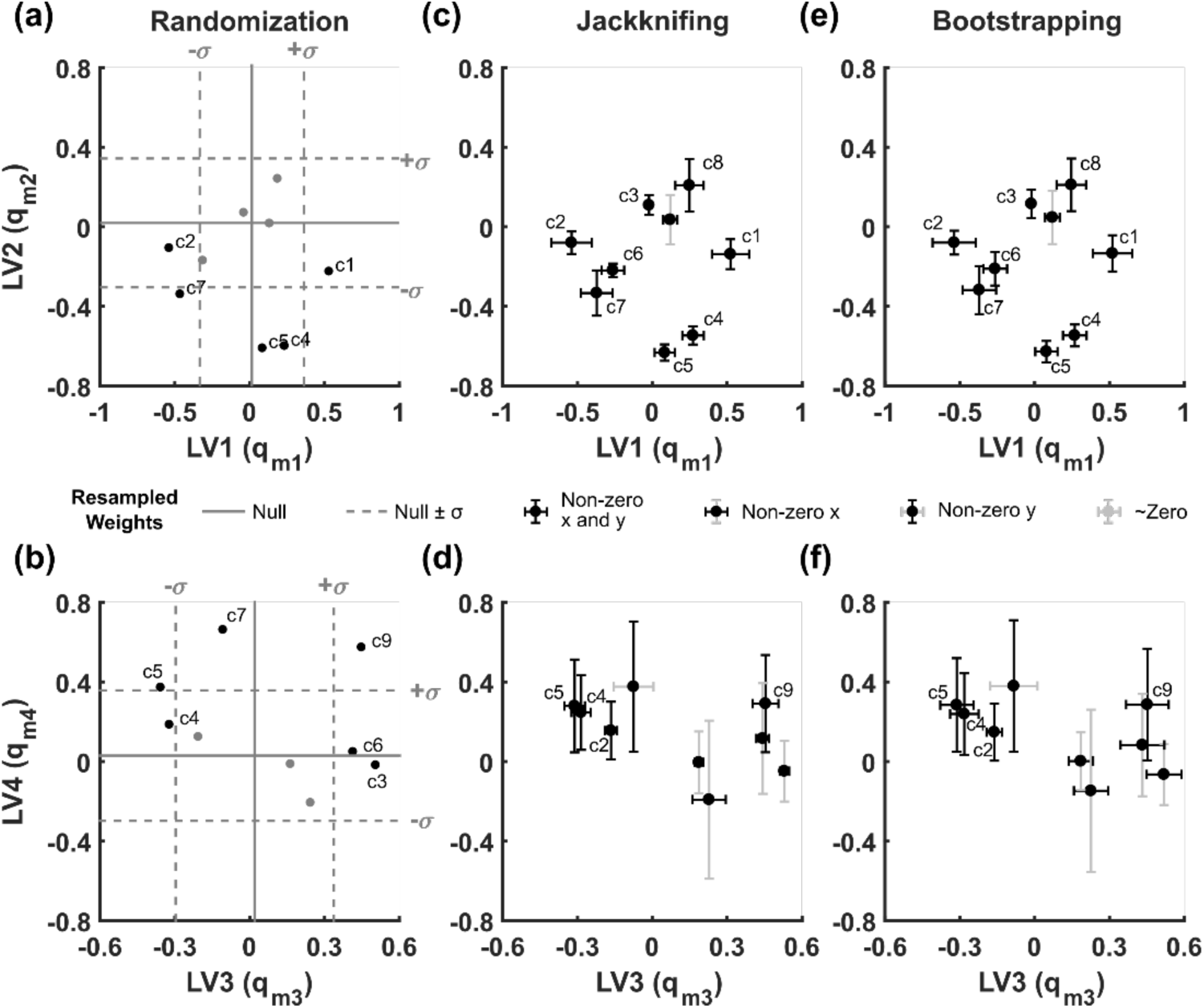
Resampling PLSR validates the robustness of higher-order LVs in multidimensional arrays. **(a, b)** Generation of a null PLSR model via randomization (*N* = 500 reshufflings within mode 1) identifies parameters of interest as variable weights in the original PLSR model (black dots) lying outside of a single standard deviation of the null PLSR model. **(c, d)** Replicate resampling (*N* = 500) by jackknifing (“leave one out”) increases confidence of most LV parameters. **(e, f)** Replicate resampling (*N* = 500) by bootstrapping (“leave one in”) yields very similar results to jackknifing, as expected given the *N* = 2 sample size for output data (Table 3). Graphs are labeled as in Fig. 4.

Using all three models resampled here, we plotted RMSE as a function of increasing LV for the global-average model compared to its mean jackknife or bootstrap replication. For the Chitforoushzadeh *et al*.^13^ model built from *in vitro* data, jackknife and bootstrap resamplings were superimposable with the global average (Fig. 9a). However, for the two *in vivo* studies, the resampled variants were consistently less accurate than the corresponding global average (Fig. 9b,c). Taken together, the results indicate that nested resampling is an effective strategy—distinct from prevailing methods—to benchmark meaningful LVs extracted from *in vivo* datasets.

**Figure 9.**
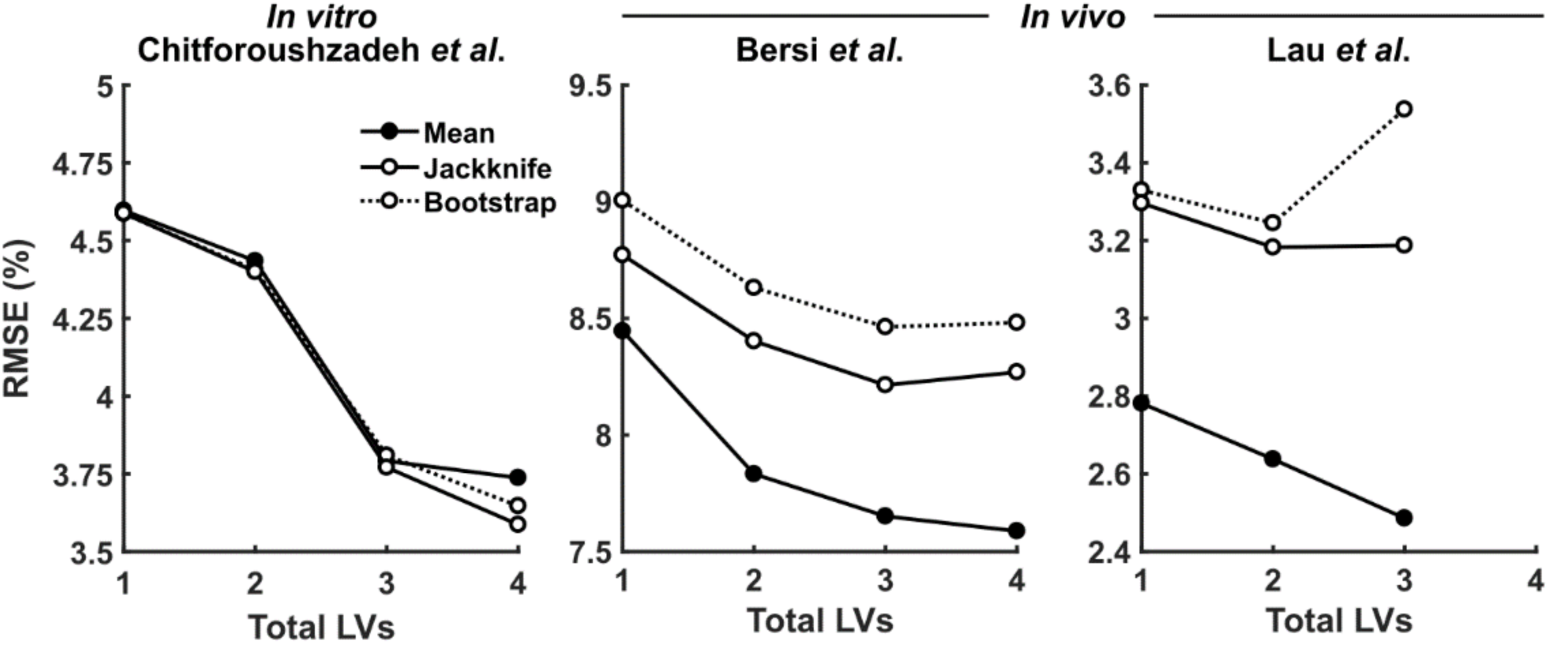
Nested resampling PLSR vets the robustness of *in vivo* multidimensional arrays. Root mean squared error (RMSE) as a function of total included LVs is reported for PLSR models of mean datasets (solid lines and filled circles; reprinted from Fig. 3a, 6a, and Chitforoushzadeh *et al*.^13^), mean predictions from jackknifed models (solid lines and open circles), and mean predictions from bootstrapped models (dotted lines and open circles).

## DISCUSSION

When applied to *in vivo* PLSR models, nested resampling is an effective way to hone in on latent variables that are robust to the replicate fluctuations of individual inbred animals. For high-variance observations, the method gives information complementary to that obtained by condition-specific jackknifing^38^ or crossvalidation^10^. In building hundreds of instances around the global-average model, nested resampling does not rely on any further assumptions to execute. However, it is important to recognize the nesting relationships within a study design and ensure that they are retained during resampling. The diversity of study designs^33^ precludes a universal software for nested resampling, but we provide code for the specific implementations here, which can readily be adapted for other *in vivo* datasets (Supplementary File S1).

Normally, direct use of replicated data in PLSR is discouraged, because replicates inflate the number of observations and reduce the stringency of crossvalidation^32^. Resampling avoids data inflation but is minimally effective for latent-variable assessment when replicates are highly reproducible. The *in vitro* model^13^ resampled here uses data with a median coefficient of variation of ~11% (Ref. ^44^), which is too small to impact the latent variables of the model. In mice, however, phenotypic variability within inbred strains is typically 3–5 times greater^31^, competing with the biological effect size of many studies. Replicates are essential for more reliable central estimates and statistical power^45^. This work shows how replicates can be repurposed to reflect better the internal variability of *in vivo* datasets and identify the robust vs. fragile components of regression models that are ordinarily limited to using replication indirectly.

The *in vivo* datasets modeled here used inbred strains of mice to minimize genotypic differences. Modeling outbred strains of animals^31^ or diverse human populations^46^ will involve very different approaches. Rather than averaging (followed by jackknife–bootstrap resampling), each individual will be better handled as a separate observation if the independent and dependent data can be reliably paired to that individual. Data pairing may be particularly difficult when X and Y observations are collected at multiple time points. The paired-vs.-unpaired resampling comparison involving the Bersi *et al*.^39^ dataset (Fig. 5) provides a useful guide for determining when less conservative experimental designs (*i.e.*, averaging without pairing) are acceptable.

The nested methods proposed here differ from prior resampling approaches that focus on defining observation sets for proper model selection^47^. Numerical Monte-Carlo simulations have a rich history in PLSR originating in chemometrics^48,49^. However, applications to replicated data have not been considered previously, likely because of the high reproducibility of measured chemical spectra. In nested resampling, the bootstrap and jackknife gauge different ends of latent-variable robustness. Bootstrapping is highly conservative, evaluating whether any random draw of replicates yields essentially the same model. Latent variables that survive bootstrapping capture large, reproducible effect sizes and thus are highly robust. Conversely, jackknifing is a much weaker test of model fragility. Global-average relationships that disappear with jackknifing are severely underpowered and should be ignored or followed up with more replicates. Together, these established tools from computational statistics^34^ enable formal examination of data qualities that would otherwise be inaccessible by PLSR alone.

The concepts put forth here generalize to other data-driven approaches besides PLSR. For example, when classifying observations by support vector machines^50^, the handling of replicated observations is often heuristic. Heinemann *et al*.^51^ investigated the effects of replicate downsampling on classification by metabolomics data with small or large variance, but nesting of replicates within observations was not considered as we did. Nested resampling of PLSR models shares conceptual analogies with the method of random forests^52^ for decision tree classifiers. Individual decision trees are unstable in their predictions, but robustness is improved when training data are randomly resampled to make ensemble classifications. Biological data *in vivo* are typically noisy and the number of observations is often limited, suggesting that some form of nested resampling would be beneficial for many data-driven methods seeking to identify molecular– cellular drivers of organismal phenotypes.

A primary motivation for applying PLSR in biological systems is to simplify complex observations and generate testable hypotheses^2,36^. The latter goal is impossible when chasing latent variables that are statistically significant overall but fragile upon replication. By using all of the *in vivo* data available, nested resampling identifies where PLSR stops modeling effect sizes and starts fitting biologically noisy averages. It contributes to the ongoing effort to improve the reproducibility of models^53^ and preclinical research^26,54^.

## MATERIALS AND METHODS

### Experimental models

Three studies were selected in which an inflammatory agent was administered *in vivo* or *in vitro* and subsequent temporal and/or spatial analyses were performed^13,14,39^. First, source data were obtained from Bersi *et al*.^39^ in which male *ApoE^−/−^* mice were infused with Angiotensin II (AngII, 1000 ng/kg/min) via an implantable osmotic mini-pump for 4, 7, 14, 21, or 28 days. Following treatment, the aorta was harvested and separated into four regions: 1) the ascending thoracic aorta (ATA) spanning from the aortic root to the brachiocephalic artery, 2) the descending thoracic aorta (DTA) spanning from the left subclavian artery to the 4th or 5th pair of intercostal arteries, 3) the suprarenal abdominal aorta (SAA) spanning from the diaphragm to the left renal artery, and 4) the infrarenal abdominal aorta (IAA) spanning from the left renal artery to the iliac trifurcation. Vessels were cleaned, sutured, and mounted on an opposing glass cannula and subjected to passive biomechanical testing without contribution from smooth muscle as previously described^55^. Briefly, vessels were preconditioned to minimize viscoelastic behavior of the material and then subjected to three fixed-length, pressure-diameter inflation tests and four fixed-pressure, force-length extension tests. Following testing, vessels were fixed in 10% neutral buffered formalin, embedded in paraffin, and sectioned and stained with Movat’s pentachrome, Picrosirius red, or Elastica van Gieson to quantify layer-specific matrix content. Additional slides were stained for CD3, CD45, CD68, MMP2, MMP12, or MMP13 expression. Details regarding region- and layer-specific matrix, inflammatory cell, and enzyme content can be found in the original publication^39^. Animal housing and experimental procedures were carried out in compliance with regulations and protocols approved by the Institutional Animal Care and Use Committee at Yale University.

Passive mechanical properties of the tissue were quantified using a microstructurally-motivated strain energy function assuming hyperelasticity. The analytical methods for determining mechanical metrics have been described in detail previously^55^. Briefly, biaxial Cauchy wall stresses were calculated as

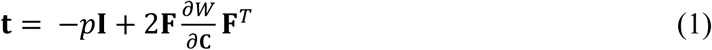

where **t** is the Cauchy stress tensor, *p* is the Lagrange multiplier enforcing incompressibility, **I** is the second-order identity matrix, **F** is the deformation gradient mapping spatial coordinates from a reference to deformed configuration, **C** is the right Cauchy-Green deformation tensor (**C** = **F***^T^***F**), and *W* is a microstructurally-motivated strain energy density function reflecting contributions of matrix constituents to material behavior. Linearized biaxial material stiffnesses were determined in terms of the second derivative of *W* with respect to deformations. These metrics, along with associated loaded geometry, were evaluated at group-specific blood pressures and at estimated *in vivo* axial stretch values.

For the second study, source data were obtained from Lau *et al*.^14^ in which male C57BL/6J mice were injected with 5 or 10 µg TNFα by retro-orbital injection for 0.5, 1, 2, 4, or 8 hours. Following treatment, mice were euthanized, and two regions of the small intestine were harvested: 1) the duodenum consisting of the 1 cm of area immediately distal to the stomach, and 2) the ileum consisting of the 3 cm of area immediately proximal to the cecum. Tissue samples were rinsed in PBS and lysed and homogenized in Bio-Plex lysis buffer or fixed in formalin for immunohistochemical analysis. Data characterizing apoptosis and proliferation were obtained by quantitative immunoblotting for cleaved caspase 3 (cc3) and by immunohistochemistry for phosphorylated histone 3 (ph3), respectively. Signaling data were obtained via Bio-Plex signaling analysis. The targets included pIκβα, pJnk, pMek1, pErk1/2, pRsk, pp38, pc-Jun, pAtf2, pAkt, pS6, pStat3, and Mek1, totaling 12 signaling targets. Details regarding the quantification of apoptosis, proliferation, and signaling are in the original publication^14^. Animal housing and experimental procedures were carried out in compliance with regulations and protocols approved by the Subcommittee on Research Animal Care at Massachusetts General Hospital.

For the third study, source data were obtained from Chitforoushzadeh *et al*.^13^ in which HT-29 cells were pretreated with interferon γ (IFNγ; 200 U/mL) for 24 hours and subsequently treated with various combinations and concentrations of TNFα, insulin, and epidermal growth factor (EGF) for 5 min, 15 min, 30 min, 1 hours, 1.5 hours, 2 hours, 4 hours, 8 hours, 12 hours, 16 hours, 20 hours, or 24 hours. Signaling metrics included 12 proteins that were evaluated via kinase activity, protein phosphorylation, total protein, phospho-total ratio, zymogen amount, or cleaved amount. Proteins included ERK, Akt, JNK1, IKK, MK2, pMEK, pFKHR, pIRS1, caspase 8, caspase 3, and EGFR. The combination of 12 proteins and multiple possible proteoforms (*e.g.*, phosphorylated protein and total protein) yielded a total of 19 signaling metrics. Additionally, microarray profiling of HT-29 cells was performed on Affymetrix U133A arrays and organized by Cluster Identification via Connectivity Kernels (CLICK). Briefly, cells were pretreated with IFNγ (200 U/mL) for 24 hours before stimulation with TNFα (0, 5, or 100 ng/mL), insulin (0, 5, or 500 ng/mL), and EGF (0, 1, or 100 ng/mL) for 4, 8, or 16 hours. CLICK clustering of microarray data yielded 9 clusters for each condition and time point^13^.

For all studies, global averages were calculated as the mean among replicates.

### Multidimensional partial least squares modeling

Multidimensional PLSR was performed in MATLAB using version 2.02 of the NPLS Toolbox^56^ after dividing each study into independent and dependent datasets according to the stated hypothesis. Model variables for the three studies are listed in Tables 1–3 with associated abbreviations, methods of acquisition, sample sizes, and input–output classifications. The algorithm for PLSR has been described in detail previously with specific application to multi-linear frameworks^13,57^. Briefly, PLSR is a simultaneous decomposition of two matrices where the scores of each decomposition are linearly related (Fig. 1). Various options exist for exact algorithms. The algorithm applied in this study is detailed below:

1. Organize independent data into an i x j x k array **X**, where i is the number of experimental conditions, j is the number of time points, and k is the number of variables in the independent dataset. In parallel, organize the dependent data into an i x l x m array **Y** where l is the number of time points, and m is the number of variables in the dependent dataset. Note that the algorithm requires the first dimension of each matrix to be equal but numbers of variables and time points need not be equal.
2. Standardize the data by mean centering and/or variance scaling the data. Different standardization techniques can yield markedly different results^58^. For Bersi *et al*.^39^, only variance scaling across mode 3 was performed, and time 0 values were subtracted for a given condition and variable from all other corresponding time points within the same condition and variable such that regional differences are not considered at baseline. For Lau *et al*.^14^ and Chitforoushzadeh *et al*.^13^, variance scaling across modes 2 and 3 was performed.
3. Initialize an i x 1 vector for the n^th^ latent variable for the dependent condition scores, **u**, and the independent condition scores, **t**. Here, **u** is initialized by performing principal components analysis on the standardized residual **Y** matrix (which equals the original scaled **Y** matrix for the first LV) and setting **u** = principal component 1. The vector **t** is randomly initialized.
4. Calculate variable and time weights for the independent data, **w**, by back projecting the independent data, **X**, onto **u**,

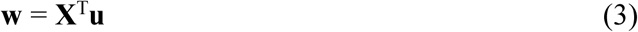

Back projection requires unfolding **X** into an i x (j*k) matrix, **X**.
5. Update independent condition scores, **t**, by projecting **X** onto **w**,

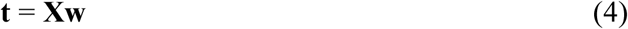
6. Calculate variable and time weights for the dependent data, **q**, by back projecting the residual of the **Y** matrix onto **t**,

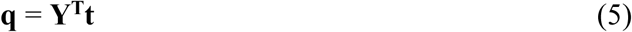

Back projection requires unfolding **Y** into an i x (l*m) matrix, **Y**.
7. Update dependent condition scores, **u**, by projecting the residual of **Y** onto **q**,

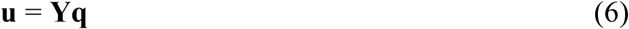
8. Calculate the difference in magnitude between the updated **t** from step 5 and the original **t** from step 3 (or the previously calculated **t** if on iteration 2 or more) and return to step 4 as long as the change in magnitude remains above a critical threshold (here, 10^−10^).
9. Calculate the regression coefficient between the independent and dependent condition scores,

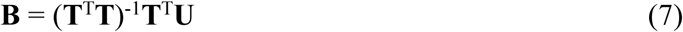

where **B** is an n x n matrix where n is the number of the current LV. If the calculation is for the first LV, then **B** becomes a scalar calculated as *b* = (**t**^T^**t**)^−1^**t**^T^**u**.
10. Calculate the residuals of **X** and **Y** by subtracting the decomposed matrices from the previous residual matrices.
11. Complete steps 4 – 10 for the desired number of LVs using **X** and **Y_res_**.

Statistical significance of variable weights was determined by calculating a null PLSR model in which raw data were shuffled within mode 1 (*i.e.*, time and variable data were shuffled within each condition) and re-standardized, and the scores and weights recalculated according to the previously mentioned algorithm. Average scores and weights were calculated for 500 iterations of reshuffling, and meaningful scores–weights were considered to be those exceeding one standard deviation from the mean. The PLSR model was cross-validated using a leave-one-out approach in which predictions for one condition are calculated from parameters derived from the remaining conditions. The root mean squared error (RMSE) for the cross-validated predictions was calculated with the addition of each LV, and the optimal number of LVs was determined by the number of LVs that minimized the RMSE in the global-average model.

### Nested resampling

Data subsets were generated by sampling individual replicates for each condition and time point by using a jackknifing (leave-one-out) approach or bootstrapping (leave-one-in) approach, and PLSR models were developed for each sampled dataset. Data were resampled 500 times with or without retention of data pairing by animal if pairing information was available. Replicate sizes per condition per time point are denoted in Tables 1–3. From Bersi *et al*.^39^, the majority of the histological samples were paired to one of the biomechanical datasets and were chosen based on the nearness of the unloaded thickness to the mean within each condition (aortic region) and time point. For ph3 data in Lau *et al*.^14^, source data for individual replicates was not available because of blinding in the original study. Therefore, sets of 5 individual samples for each condition (intestinal region and TNFα dose) and time point were simulated from published means and standard deviations by assuming the data were normally distributed.

For each randomly generated dataset, scores and weights were calculated using the number of LVs required for the corresponding mean dataset to facilitate comparison to the global-average model. Each model was cross-validated using the leave-one-out approach as previously described, and scores, weights, and cross-validated predictions were summarized and compared to the corresponding values derived from the model of the mean dataset.

## ACKNOWLEDGEMENTS

The authors would like to thank Dr. Matthew Bersi and Prof. Ken Lau for generously providing source data and the interpretations needed for its reuse in this study and Prof. Jay Humphrey for his valuable insight and feedback during the study design and manuscript preparation. This work was supported by the David & Lucile Packard Foundation #2009-34710 (K.A.J.) and the National Institutes of Health #U01-CA215794 (K.A.J.) and #R01-HL105297 (C.A. Figueroa and J.D. Humphrey).

## COMPETING INTERESTS STATEMENT

The authors declare no competing interests.

## AUTHOR CONTRIBUTIONS

A.W.C. participated in study design, code generation, model development and interpretation, and manuscript preparation. K.A.J. led the study design and participated in model interpretation and manuscript preparation. Both authors reviewed and approved the manuscript.

## DATA AVAILABILITY

All code and source data are available in Supplementary File S1. Parameter values for the the Bersi *et al*.^39^, Lau *et al*.^14^, and Chitforoushzadeh *et al*.^13^ PLSR models are available in Supplementary File S2.

