## Supplementary Figures for "Robust latent-variable interpretation of *in vivo* regression models by nested resampling"

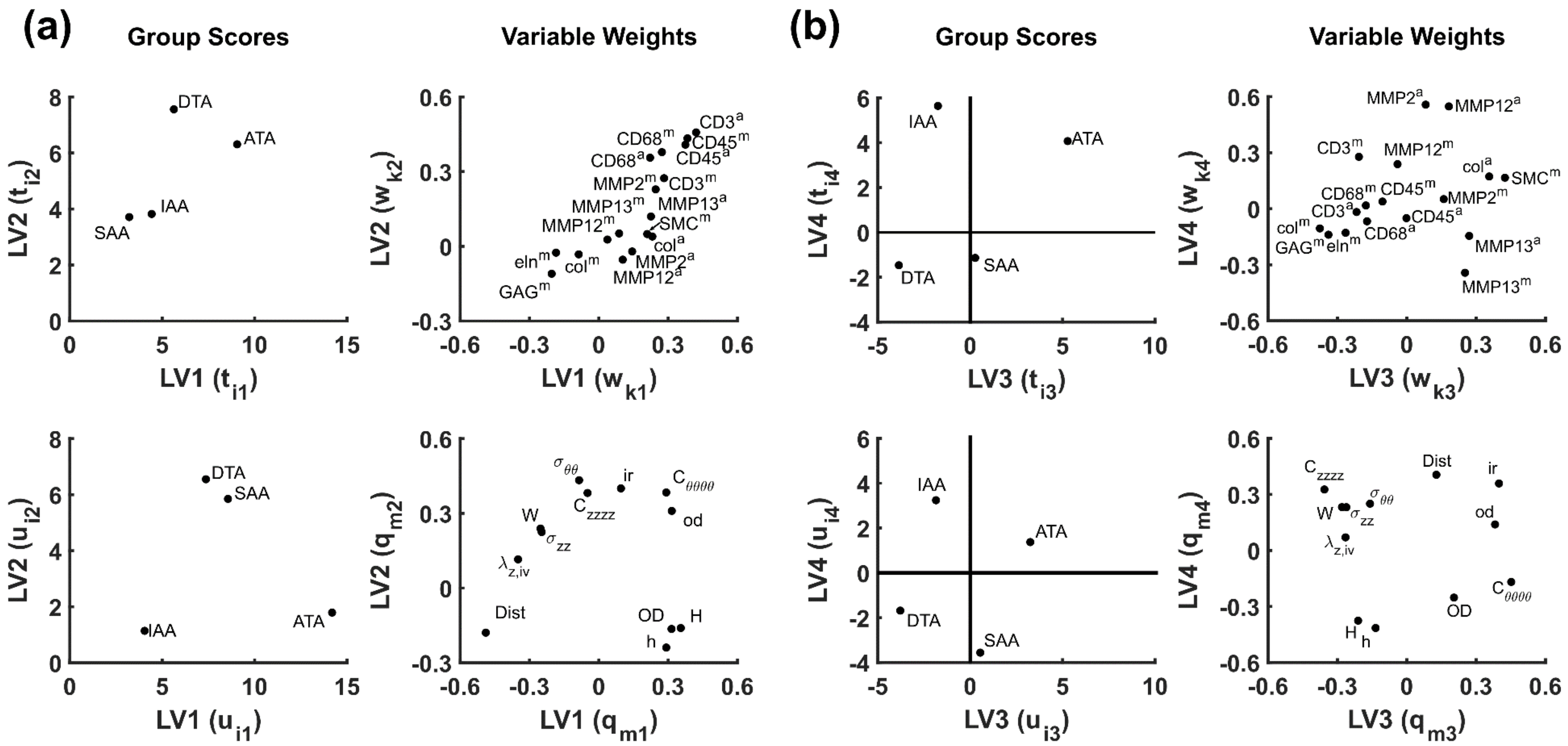


**Supplementary Figure S1.** Group scores (mode 1) and variable weights (mode 3) for all latent variables (LVs) from a model of the mean data from Bersi *et al.*[1] **(a)** Leading LVs (i.e., LV1–2) capture salient features of the data observed in original study. **(b)** Trailing LVs (i.e., LV3–4) differentiate spatial variations in aortic remodeling. Independent scores ($t_{in}$) and weights ($w_{kn}$) are depicted in the top row. Dependent scores ($u_{in}$) and weights ($q_{mn}$) are depicted in the bottom row. Sign conventions for independent and dependent group scores yielded positive inner relationships for all experimental conditions.

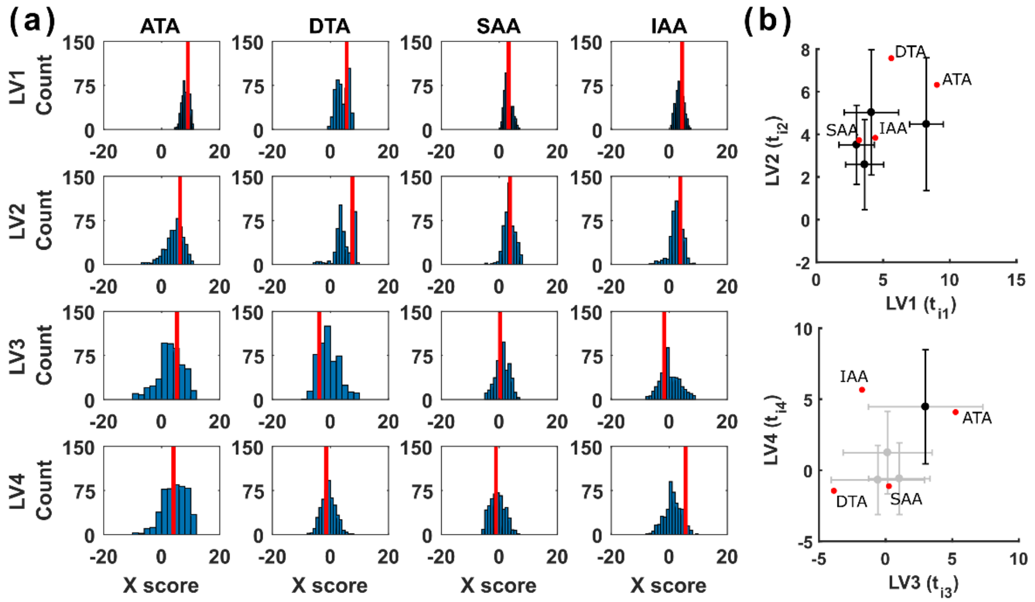


**Supplementary Figure S2.** Nested resampling uncertainty is not caused by sign flipping of LV axes. **(a)** Lack of bimodal, zero-centered X score distributions among bootstrapped replicates. Solid red lines denote scores obtained from the model of the mean data set. **(b)** Bootstrap resampling ($N = 500$) of independent scores are shown as the mean ± standard deviation. Values from the global-average model of the mean data are denoted in red and correspond to red vertical lines in (a). ATA – ascending thoracic aorta, DTA – descending thoracic aorta, SAA – suprarenal abdominal aorta, IAA – infrarenal abdominal aorta.

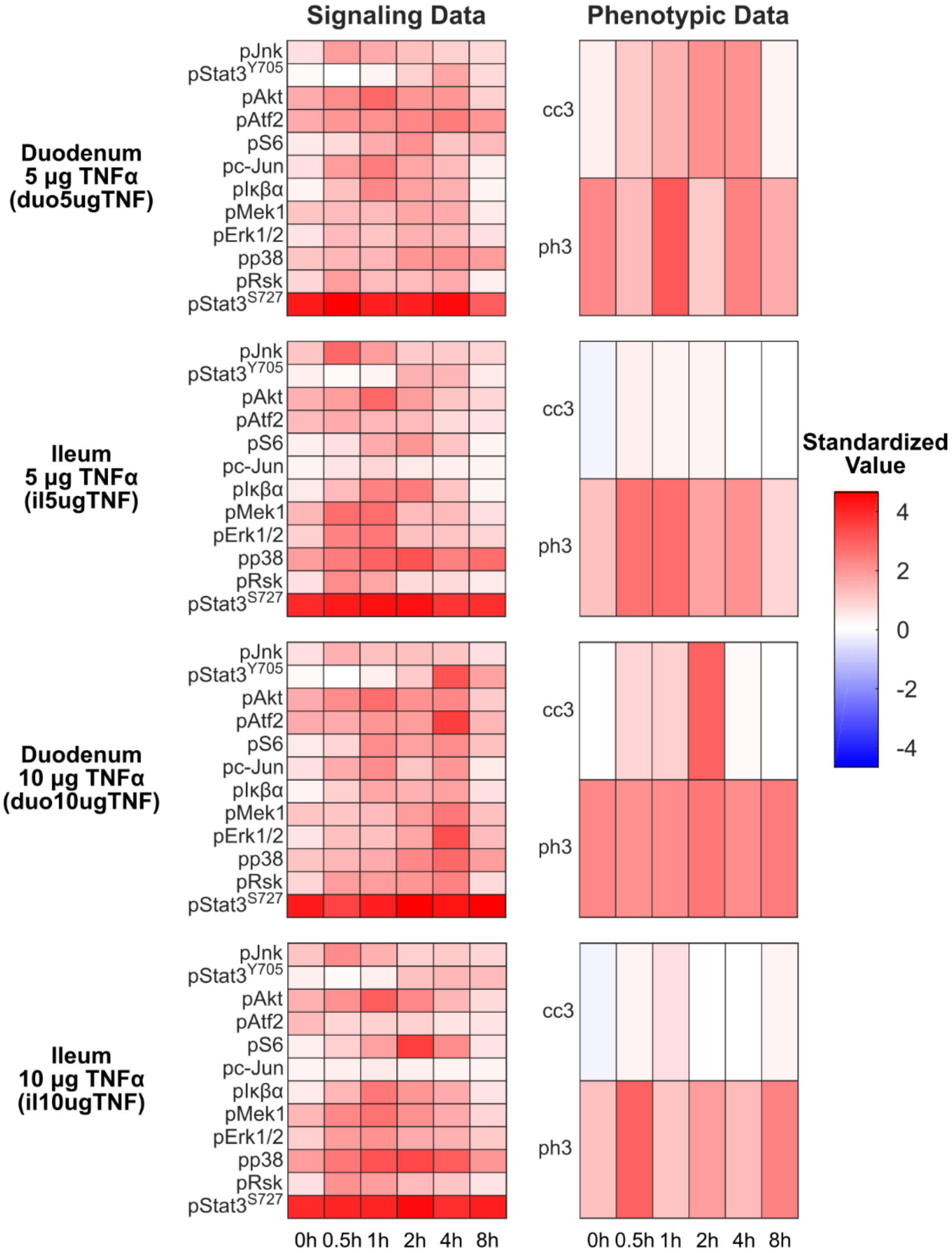


**Supplementary Figure S3.** Time-resolved profiling of cellular signaling, apoptosis, and proliferation during TNFα-induced intestinal inflammation. Mice were treated with TNFα at the indicated dose and intestinal tissue harvested at the indicated time points for subsequent molecular and histological analysis (Table 2). Data from Lau *et al.*[2] are separated by independent (left) and dependent data (right) and the combination of intestinal region and treatment (rows).

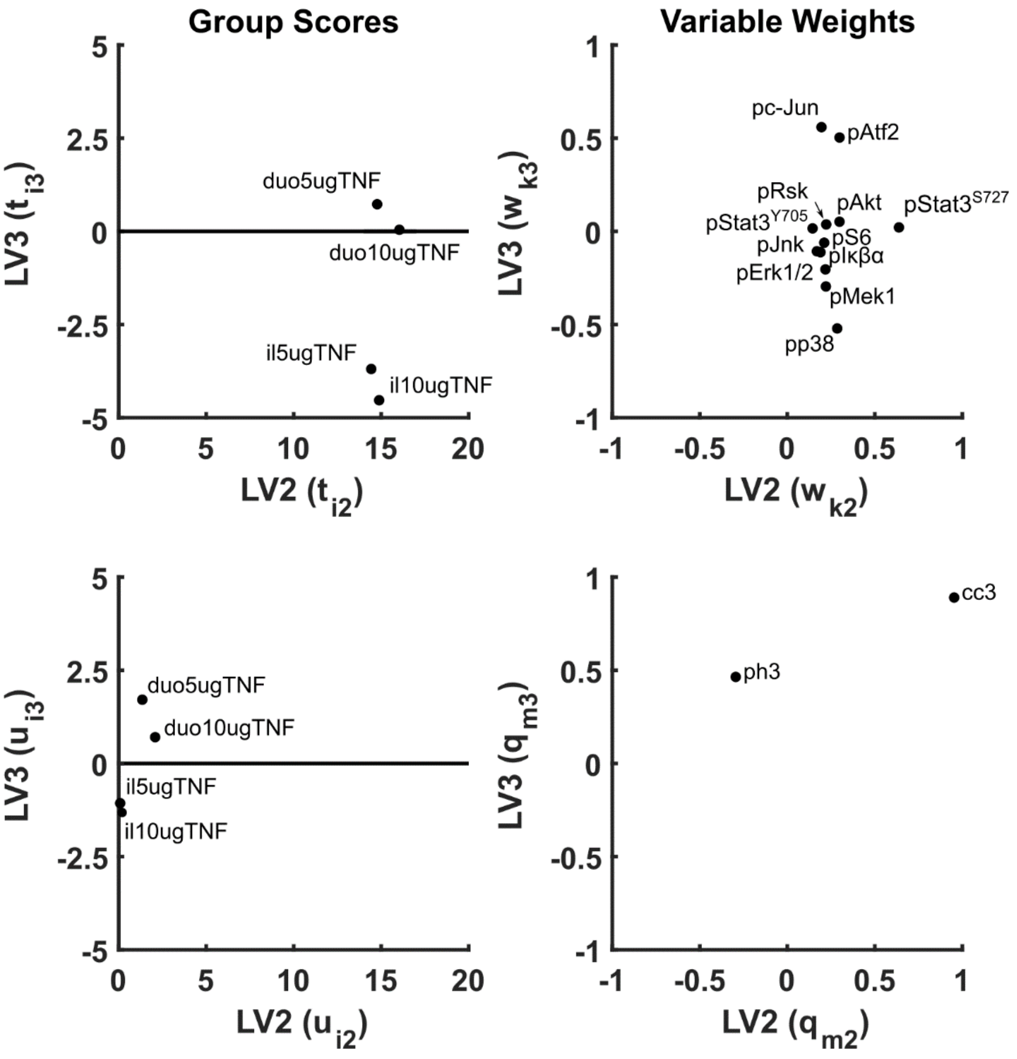


**Supplementary Figure S4.** Group scores (mode 1) and variable weights (mode 3) for a model of the mean data from Lau *et al*.[2] Independent scores ($t_{in}$) and weights ($w_{kn}$) are depicted in the top row. Dependent scores ($u_{in}$) and weights ($q_{mn}$) are depicted in the bottom row. Sign conventions for independent and dependent group scores yielded positive inner relationships for all experimental conditions. LV1 is omitted for clarity.

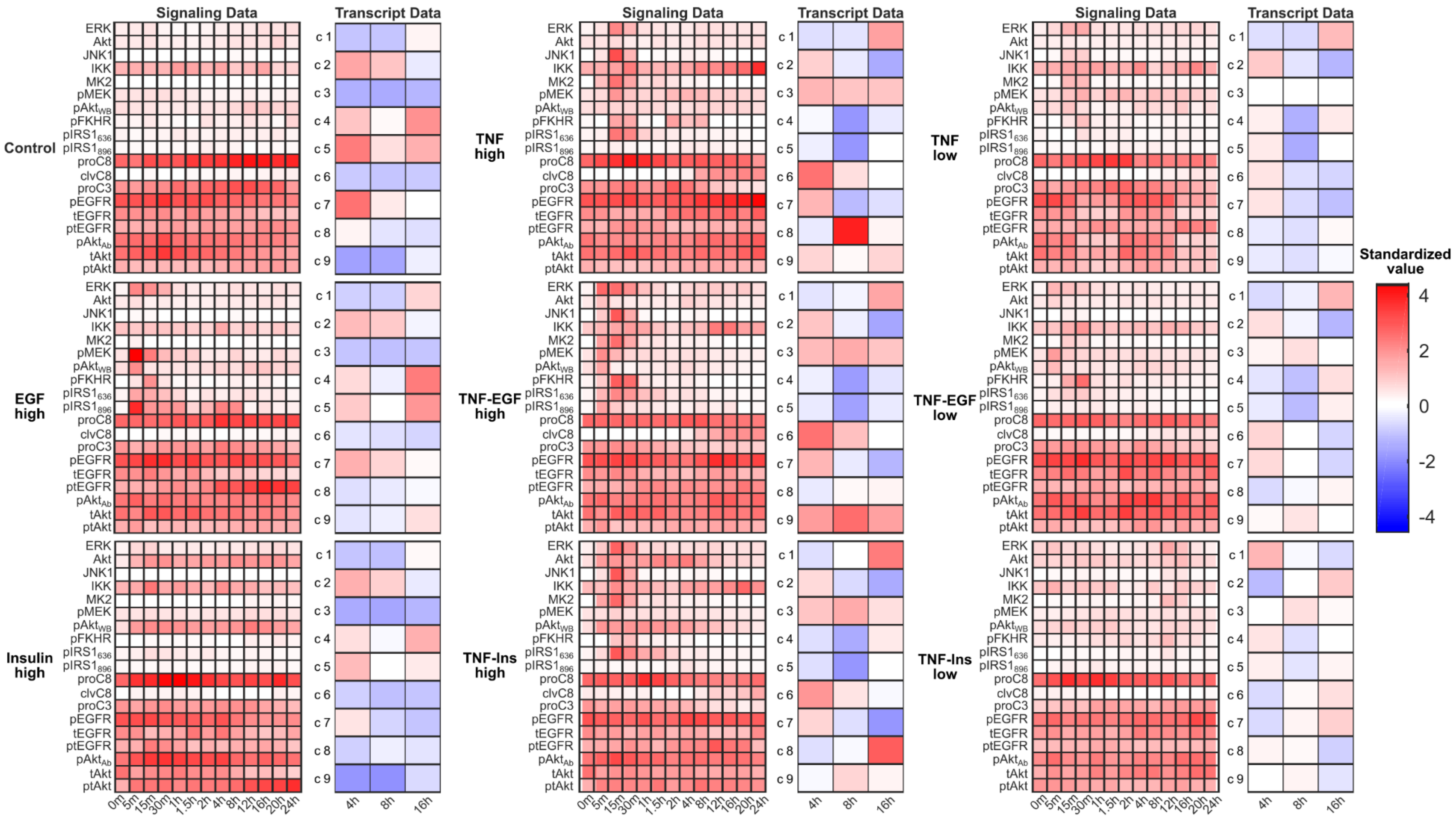


**Supplementary Figure S5.** Time-resolved profiling of cellular signaling and gene expression during *in vitro* stimulation with combinations of TNFα, EGF, and insulin. HT29 colon adenocarcinoma cells were sensitized and treated with various combinations of TNFα, EGF, and insulin at the indicated dose and cells harvested at the indicated time points for subsequent molecular analysis (Table 3). Data from Chitforoushzadeh *et al.*[3] are separated by independent (left) and dependent data (right) and treatment conditions.
