## Supplementary File S1 for "Robust latent-variable interpretation of *in vivo* regression models by nested resampling": herrorbar.m

```
function hh = herrorbar(x, y, l, u, symbol)
%HERRORBAR Horizontal Error bar plot.
%   HERRORBAR(X,Y,L,R) plots the graph of vector X vs. vector Y with
%   horizontal error bars specified by the vectors L and R. L and R contain the
%   left and right error ranges for each point in X. Each error bar
%   is L(i) + R(i) long and is drawn a distance of L(i) to the right and R(i)
%   to the right the points in (X,Y). The vectors X,Y,L and R must all be
%   the same length. If X,Y,L and R are matrices then each column
%   produces a separate line.
%
%   HERRORBAR(X,Y,E) or HERRORBAR(Y,E) plots X with error bars [X-E X+E].
%   HERRORBAR(...,'LineSpec') uses the color and linestyle specified by
%   the string 'LineSpec'. See PLOT for possibilities.
%
%   H = HERRORBAR(...) returns a vector of line handles.
%
%   Example:
%      x = 1:10;
%      y = sin(x);
%      e = std(y)*ones(size(x));
%      herrorbar(x,y,e)
%   draws symmetric horizontal error bars of unit standard deviation.
%
%   This code is based on ERRORBAR provided in MATLAB.   
%
%   See also ERRORBAR

%   Jos van der Geest
%  
%
%   File history:
%   August 2006 (Jos): I have taken back ownership. I like to thank Greg Aloe from
%   The MathWorks who originally introduced this piece of code to the
%   Matlab File Exchange. 
%   September 2003 (Greg Aloe): This code was originally provided by Jos
%   from the newsgroup comp.soft-sys.matlab:
%   http://newsreader.mathworks.com/WebX?50@118.fdnxaJz9btF^1@.eea3ff9
%   After unsuccessfully attempting to contact the orignal author, I
%   decided to take ownership so that others could benefit from finding it
%   on the MATLAB Central File Exchange.

if min(size(x))==1,
    npt = length(x);
    x = x(:);
    y = y(:);
    if nargin > 2,
        if ~isstr(l),
            l = l(:);
        end
        if nargin > 3
            if ~isstr(u)
                u = u(:);
            end
        end
    end
else
    [npt,n] = size(x);
end

if nargin == 3
    if ~isstr(l)
        u = l;
        symbol = '-';
    else
        symbol = l;
        l = y;
        u = y;
        y = x;
        [m,n] = size(y);
        x(:) = (1:npt)'*ones(1,n);;
    end
end

if nargin == 4
    if isstr(u),
        symbol = u;
        u = l;
    else
        symbol = '-';
    end
end

if nargin == 2
    l = y;
    u = y;
    y = x;
    [m,n] = size(y);
    x(:) = (1:npt)'*ones(1,n);;
    symbol = '-';
end

u = abs(u);
l = abs(l);

if isstr(x) | isstr(y) | isstr(u) | isstr(l)
    error('Arguments must be numeric.')
end

if ~isequal(size(x),size(y)) | ~isequal(size(x),size(l)) | ~isequal(size(x),size(u)),
    error('The sizes of X, Y, L and U must be the same.');
end

tee = (max(y(:))-min(y(:)))/100; % make tee .02 x-distance for error bars
% changed from errorbar.m
xl = x - l;
xr = x + u;
ytop = y + tee;
ybot = y - tee;
n = size(y,2);
% end change

% Plot graph and bars
hold_state = ishold;
cax = newplot;
next = lower(get(cax,'NextPlot'));

% build up nan-separated vector for bars
% changed from errorbar.m
xb = zeros(npt*9,n);
xb(1:9:end,:) = xl;
xb(2:9:end,:) = xl;
xb(3:9:end,:) = NaN;
xb(4:9:end,:) = xl;
xb(5:9:end,:) = xr;
xb(6:9:end,:) = NaN;
xb(7:9:end,:) = xr;
xb(8:9:end,:) = xr;
xb(9:9:end,:) = NaN;

yb = zeros(npt*9,n);
yb(1:9:end,:) = ytop;
yb(2:9:end,:) = ybot;
yb(3:9:end,:) = NaN;
yb(4:9:end,:) = y;
yb(5:9:end,:) = y;
yb(6:9:end,:) = NaN;
yb(7:9:end,:) = ytop;
yb(8:9:end,:) = ybot;
yb(9:9:end,:) = NaN;
% end change


[ls,col,mark,msg] = colstyle(symbol); if ~isempty(msg), error(msg); end
symbol = [ls mark col]; % Use marker only on data part
esymbol = ['-' col]; % Make sure bars are solid

h = plot(xb,yb,esymbol); hold on
h = [h;plot(x,y,symbol)];

if ~hold_state, hold off; end

if nargout>0, hh = h; end
```
