## Supplementary File S1 for "Robust latent-variable interpretation of *in vivo* regression models by nested resampling": calcore.m

```
function [G]=calcore(X,Factors,Options,O,MissingExist);

%CALCORE Calculate the Tucker core
%
%	
% [G]=calcore(X,Factors,Options);
% [G]=calcore(X,Factors);
%
% This algorithm applies to the general N-way case, so
% the unfolded X can have any number of dimensions. The principles of
% 'projections' and 'systematic unfolding methodology (SUM)' are used
% in this algorithm so orthogonality is required.
% This algorithm can handle missing values in X and
% also allows for TUCKER2 models using the an empty matrix in the
% corresponding cell of Factors.
% The variable 'Factors' must contain the stringed-out factors.

% Copyright (C) 1995-2006  Rasmus Bro & Claus Andersson
% Copenhagen University, DK-1958 Frederiksberg, Denmark,

format compact
format long

DimX = size(X);
X = reshape(X,DimX(1),prod(DimX(2:end)));
ff = [];
for f=1:length(Factors)
   ff=[ff;Factors{f}(:)];
   Fac(f)=size(Factors{f},2);
   if isempty(Factors{f}) % 'Tucker2' - i.e. no compression in that mode
      Fac(f) = -1;
   end
end
Factors = ff;

% Initialize system variables
if length(Fac)==1,
   Fac=Fac*ones(size(DimX));
end;

Fac_orig=Fac;
i=find(Fac==-1);
Fac(i)=zeros(1,length(i));
N=size(Fac,2);
FIdx0=zeros(1,N);
FIdx1=zeros(1,N);
if ~exist('MissingExist')
   if sum(isnan(X(:)))>0,
      MissingExist=1;
   else
      MissingExist=0;
   end;
end;
FIdx0=cumsum([1 DimX(1:N-1).*Fac(1:N-1)]);
FIdx1=cumsum([DimX.*Fac]);
if ~exist('O') | isempty(O),
   O=1;
end;


if O, %means orthogonality
   CurDimX=DimX;
   RedData=X;
   for c=1:N,
      
      if Fac_orig(c)==-1,
         kthFactor=eye(DimX(c));
         CurDimX(c)=DimX(c);
      else
         kthFactor=reshape(Factors(FIdx0(c):FIdx1(c)),DimX(c),Fac(c));
         CurDimX(c)=Fac(c);
      end;      
      if MissingExist
         RedData=missmult(kthFactor',RedData);
      else
         RedData=kthFactor'*RedData;
      end;
      
      if c~=N,
         newi=CurDimX(c+1);
         newj=prod(CurDimX)/CurDimX(c+1);
      else
         newi=CurDimX(1);
         newj=prod(CurDimX)/CurDimX(1);
      end;
      
      RedData=reshape(RedData',newi,newj);
   end;
   G=RedData;
else %oblique factors
   
   LMatTmp=1;
   if Fac_orig(1)==-1,
      LMatTmp=eye(DimX(c));
   else
      LMatTmp=reshape(Factors(FIdx0(1):FIdx1(1)),DimX(1),Fac(1));
   end;    
   
   RMatTmp=1;
   for c=2:N,
      if Fac_orig(c)==-1,
         kthFactor=eye(DimX(c));
      else
         kthFactor=reshape(Factors(FIdx0(c):FIdx1(c)),DimX(c),Fac(c));
      end;    
      RMatTmp=ckron(kthFactor',RMatTmp);
   end;
   
   if MissingExist
      RedData=missmult(pinv(LMatTmp),X);
      RedData=missmult(RedData,pinv(RMatTmp));
   else
      RedData=LMatTmp\X;
      RedData=RedData/RMatTmp;
   end;
   
   G=RedData;
   
end;    

for i = 1:length(Fac)
   if Fac(i)==0
      Fac(i) = DimX(i);
   end
end
G = reshape(G,Fac);

return
```
