## Supplementary File S1 for "Robust latent-variable interpretation of *in vivo* regression models by nested resampling": ckron.m

```
function C=ckron(A,B)
%CKRON
% C=ckron(A,B)
%
% Copyright (C) 1995-2006  Rasmus Bro & Claus Andersson
% Copenhagen University, DK-1958 Frederiksberg, Denmark,
%

% Claus Andersson, Jan. 1996

% Should not be compiled to overwrite ckron.mex

[mA,nA] = size(A);
[mB,nB] = size(B);

C = zeros(mA*mB,nA*nB);
if mA*nA <= mB*nB
  for i = 1:mA
  iC = 1+(i-1)*mB:i*mB;
    for j = 1:nA
      jC = 1+(j-1)*nB:j*nB;
      C(iC,jC) = A(i,j)*B;
    end
  end
else
  for i = 1:mB
    iC = i:mB:(mA-1)*mB+i;
    for j = 1:nB
      jC = j:nB:(nA-1)*nB+j;
      C(iC,jC) = B(i,j)*A;
    end
  end
end
```
