## Supplementary File S1 for "Robust latent-variable interpretation of *in vivo* regression models by nested resampling": cmatrep.m

```
function [A]=cmatrep(A,crit,v1,v2)

%CMATREP optimized matrep
% 
%[A]=cmatrep(A,crit,v1,v2)
%Criteria   all values   will be replaced with
% 'eq'         =v1               v2
% 'lt'         <v1               v2
% 'gt'         >v1               v2

[a1 a2]=size(A);

if (crit=='lt') | (crit=='LT'), 
for i=1:a1;
  for j=1:a2;
    if A(i,j)<v1,
    	A(i,j)=v2;
    end;
  end;
end;
end;

if (crit=='eq') | (crit=='EQ'), 
for i=1:a1;
  for j=1:a2;
    if A(i,j)==v1,
    	A(i,j)=v2;
    end;
  end;
end;
end;

if (crit=='gt') | (crit=='GT'), 
for i=1:a1;
  for j=1:a2;
    if A(i,j)>v1,
    	A(i,j)=v2;
    end;
  end;
end;
end;
```
