## Supplementary File S1 for "Robust latent-variable interpretation of *in vivo* regression models by nested resampling": contents.m

```
% The N-way Toolbox for MATLAB (ver. 7.3)
% Version 3.2 March-2012
% Download from http://www.models.life.ku.dk/source/nwaytoolbox/
% $Revision: 3.2.0$  $1-Mar-2012$  $CA/RB$
%
% Copyright (C) 1995-2012  Rasmus Bro & Claus Andersson
% University of Copenhagen , DK-1958 Frederiksberg, Denmark,
%
%
%-----------------------------------------------
%MODELS
%
%TUCKER        Multi-way tucker model
%NPLS          Multilinear partial least squares regression
%NPRED         Prediction with NPLS model
%PARAFAC       Multiway parafac model
%GRAM          Generalized rank annihilation method
%DTLD          Direct trilinear decomposition
%
%
%-----------------------------------------------
%CENTER AND SCALE
%
%NPROCESS      Pre- and postprocessing of multiway arrays
%
%-----------------------------------------------
%MODEL EVALUATION
%
%NCROSSDECOMP  Crossvalidation of PARAFAC/Tucker/PCA
%NCROSSREG     Cross-validation of regression model
%NMODEL        Make model of data from loadings
%NCOSINE       Multiple cosine/Tuckers congruence coefficient
%FAC2LET       Convert 'Factors' to component matrices
%CORCOND       Core consistency for PARAFAC model
%PFTEST        Find the number of PARAFAC components
%TUCKTEST      Find the number of Tucker components
%
%-----------------------------------------------
%TUCKER CORE 
%
%EXPLCORE      For interpretation of cores and arrays
%MAXDIA3       Maximize core diagonality
%MAXSWD3       Maximize core slice-wise diagonality
%MAXVAR3       Maximize core squared variance
%COREDIAN      Calculates core diagonality
%CORESWDN      Calculates the 'core-slice-wise-diagonality'.
%COREVARN      Calculates the 'core-variance' 
%T3CORE        Calculate Tucker core
%CALCORE       Calculate the Tucker core
%
%-----------------------------------------------
%AUXILARY MULTI-WAY THINGS
%
%NEYE          Produces a super-diagonal array
%NIDENT        Make 'identity' multi-way array
%INI           Initialization of loadings
%INITUCK       Initialization of loadings
%KRB           Khatri-Rao-Bro product
%NSHAPE        Rearrange a multi-way array
%NTIMES        Array multiplication
%
%-----------------------------------------------
%PLOTTING
%
%PLOTFAC       Plot the contents of Factors
%PFPLOT        Plot parafac model
%
%-----------------------------------------------
%MULTI-WAY TOOLS
%
%PARADEMO      PARAFAC demo
%TUCKDEMO      Tucker demo
%TWO2N         Conversion of indices between unfoldings and N-way arrays
%
%-----------------------------------------------
%ADDITIONAL FILES
%
%UNIMODALCROSSPRODUCTS   For unimodel regression
%CKRON                   Optimized Kronecker product
%CMATREP                 Optimized matrep
%COMPLPOL                Used in DTLD
%DERDIA3                 Used for core rotation
%DERSWD3                 Used for core rotation
%DERVAR3                 Used for core rotation
%GETINDXN
%GSM                     GS orthogonalization
%MISSMEAN                Mean of a matrix X with NaN's
%MISSMULT                Product of two matrices containing NaNs
%MISSSUM                 Sum of a matrix X with NaN's
%MONREG                  Monotone regression
%FASTNNLS                Fast version of built-in NNLS
%FNIPALS                 Nipals algorithm for PCA
%NONNEG1                 Alternative to NNLS
%NORMIT                  Normalize
%ULSR                    For unimodel regression
%UNIMODAL                Unimodal regression
%PFLS                    LS regression tool for parafac
%NSETDIFF                
%REFOLD3                 Refold an unfolded array
%SETNANS1                Fluorescence artifact treatment
%SETOPTS                 For setting options
%STDNAN                  Estimate std with NaN's
%
%
% Type
% >type readme.txt 
% for more help
```
