## Supplementary File S1 for "Robust latent-variable interpretation of *in vivo* regression models by nested resampling": corcond.m

```
function [Consistency,G,stdG,Target]=corcond(X,Factors,Weights,Plot);

%CORCOND Core consistency for PARAFAC model
%
% See also:
% 'unimodal' 'monreg' 'fastnnls'
%
% CORe CONsistency DIAgnostics (corcondia)
% Performs corcondia of a PARAFAC model and returns the cocote plot
% as well as the degree of consistency (100 % is max).
%
% Consistency=corcond(X,Factors,Weights,Plot);
% 
% INPUT
% X        : Data array 
% Factors  : Factors given in standard format as a cell array
% Weights  : Optional weights (otherwise skip input or give an empty array [])
% Plot     = 0 or not given => no plots are produced
%          = 1              => normal corcondia plot
%          = 2              => corcondia plot with standard deviations 
%
% OUTPUT
% The core consistency given as the percentage of variation in a Tucker3 core
% array consistent with the theoretical superidentity array. Max value is 100%
% Consistencies well below 70-90% indicates that either too many components
% are used or the model is otherwise mis-specified.
%

% Copyright (C) 1995-2006  Rasmus Bro & Claus Andersson
% Copenhagen University, DK-1958 Frederiksberg, Denmark,

% $ Version 1.02 $ Date 28. July 1998 $ Not compiled $
% $ Version 2.00 $ May 2001 $ Changed to array notation $ RB $ Not compiled $
% $ Version 2.01 $ Feb 2003 $ replaced regg with t3core when weights are used $ RB $ Not compiled $

DimX = size(X);
X = reshape(X,DimX(1),prod(DimX(2:end)));
Fac = size(Factors{1},2);

if nargin<4
  Plot=0;
end
if nargin<3
  Weights=0;
end

ord=length(DimX);
l_idx=0;
for i=1:ord
  l_idx=[l_idx sum(DimX(1:i))*Fac];
end


% Scale all loadings to same magnitude
magn=ones(Fac,1);
for i=1:ord
   L=Factors{i};
   for f=1:Fac
     magn(f)=magn(f)*norm(L(:,f));
     L(:,f)=L(:,f)/norm(L(:,f));
   end
   Factors{i}=L;
end
% Magn holds the singular value of each component. Scale each loading vector by 
% the cubic root (if three-way) so all loadings of a component have the same variance

magn = magn.^(1/ord);
for i=1:ord
   L=Factors{i};
   for f=1:Fac
     L(:,f)=L(:,f)*magn(f);
   end
   Factors{i}=L;
end


% Make diagonal array holding the magnitudes
Ident=nident(Fac,ord);
if Fac>1
   DimIdent=ones(1,ord)*Fac;
   Ident=nshape(reshape(Ident,DimIdent),ord);
end

% Make matrix of Kronecker product of all loadings expect the large; Z = kron(C,B ... )
  NewFac=[];
  NewFacNo=[];
  for i=ord:-1:1
    Z=Factors{i};
    % Check its of full rank or adjust core and use less columns
    rankZ=rank(Z);
    if rankZ<Fac
       %OLD out=Z(:,rankZ+1:Fac);Z=Z(:,1:rankZ);H=[[eye(rankZ)] pinv(Z)*out];Ident=H*Ident;
       [q,r]=qr(Z);
       Ident=r*Ident;
       Z=q;
       DimIdent(i)=size(r,1);
    end
    if i>1&Fac>1
      Ident=nshape(reshape(Ident,DimIdent([i:ord 1:i-1])),ord);
    end
    NewFac{i}=Z;
    NewFacNo=[rankZ NewFacNo];
  end
Factors=NewFac;
Fac=NewFacNo;
if nargin<3
  [G,stdG]=regg(reshape(X,DimX),Factors,Weights); %Doesn't work with weights
else
  G=t3core(reshape(X,DimX),Factors,Weights);
  stdG = G; % Arbitrary (not used)
end

DimG = size(G);
G = G(:);

 Ident=Ident(:);
 Target=Ident;
 [a,b]=sort(abs(Ident));
 b=flipud(b);
 Ident=Ident(b);
 GG=G(b);
 stdGG=stdG(b);
 bNonZero=find(Ident);
 bZero=find(~Ident);

 ssG=sum(G(:).^2);
 Consistency=100*(1-sum((Target-G).^2)/ssG);
 
 
 if Plot
    clf
    Ver=version;
    Ver=Ver(1);
    if Fac>1
       eval(['set(gcf,''Name'',''Diagonality test'');']);
       if Ver>4
          plot([Ident(bNonZero);Ident(bZero)],'y','LineWidth',3)
          hold on
          plot(GG(bNonZero),'ro','LineWidth',3)
          plot(length(bNonZero)+1:prod(Fac),GG(bZero),'gx','LineWidth',3)
          if Plot==2
            line([[1:length(G)];[1:length(G)]],[GG GG+stdGG]','LineWidth',1,'Color',[0 0 0])
            line([[1:length(G)];[1:length(G)]],[GG GG-stdGG]','LineWidth',1,'Color',[0 0 0])
          end
          hold off
          title(['Core consistency ',num2str(Consistency),'% (yellow target)'],'FontWeight','bold','FontSize',12)          
       else
          plot([Ident(bNonZero);Ident(bZero)],'y')
          hold on
          plot(GG(bNonZero),'ro')
          plot(length(bNonZero)+1:prod(Fac),GG(bZero),'gx')
          if Plot==2
            line([[1:length(G)];[1:length(G)]],[GG GG+stdGG]','LineWidth',1,'Color',[0 0 1])
            line([[1:length(G)];[1:length(G)]],[GG GG-stdGG]','LineWidth',1,'Color',[0 0 1])
          end
          hold off
          title(['Core consistency ',num2str(Consistency),'% (yellow target)'])
       end
       xlabel('Core elements (green should be zero/red non-zero)')
       ylabel('Core Size')
    else
       eval(['set(gcf,''Name'',''Diagonality test'');']);
       title(['Core consistency ',num2str(Consistency),'% (yellow target)'])
       xlabel('Core elements (green should be zero/red non-zero)')
       ylabel('Size')
       plot(GG(bNonZero),'ro')
       title(['Core consistency ',num2str(Consistency),'%'])
       xlabel('Core elements (red non-zero)')
       ylabel('Core Size')
    end
 end

G = reshape(G,DimG);

function [G,stdG]=regg(X,Factors,Weights);

%REGG Calculate Tucker core
%
% Calculate Tucker3 core

% $ Version 2.00 $ May 2001 $ Changed to array notation $ RB $ Not compiled $

DimX = size(X);
X = reshape(X,DimX(1),prod(DimX(2:end)));
Fac = size(Factors{1},2);

ord=length(DimX);
if ord<3
   disp(' ')
   disp(' !!Corcondia only applicable for three- and higher-way arrays!!')
   return
end

if length(Fac)==1
   for i=1:length(Factors)
      Fac(i) = size(Factors{i},2);
   end
end
vecX=X(:); % Vectorize X

% Make sure Weights are defined (as ones if none given)
if nargin<3
   Weights=ones(size(X));
end
if length(Weights(:))~=length(X(:));
   Weights=ones(size(X));
end
Weights=Weights(:);

% Set weights of missing elements to zero
id=find(isnan(vecX));
Weights(id)=zeros(size(id));
vecX(id)=zeros(size(id));

% Create Kronecker product of all but the last mode loadings
L2 = Factors{end-1};
L1 = Factors{end-2};
Z = kron(L2,L1);
for o=ord-3:-1:1
   Z = kron(Z,Factors{o});
end


% Make last mode loadings, L
L=Factors{end};

% We want to fit the model ||vecX - Y*vecG||, where Y = kron(L,Z), but 
% we calculate Y'Y and Y'vecX by summing over k
J=prod(DimX(1:ord-1));
Ytx = 0;
YtY = 0;
for k=1:DimX(ord)
   W=Weights((k-1)*J+1:k*J);
   WW=(W.^2*ones(1,prod(Fac)));
   Yk  = kron(L(k,:),Z);
   Ytx = Ytx + Yk'*(W.*vecX((k-1)*J+1:k*J));
   YtY = YtY + (Yk.*WW)'*Yk;
end

G=pinv(YtY)*Ytx;

if nargout>1
   se = (sum(vecX.^2) + G'*YtY*G -G'*Ytx);
   mse = se/(length(vecX)-length(G));
   stdG=sqrt(diag(pinv(YtY))*mse);
end
G = reshape(G,Fac);
```
