## Supplementary File S1 for "Robust latent-variable interpretation of *in vivo* regression models by nested resampling": coredian.m

```
function [DIA_Rel,DIA_Abs]=CoreDian(C)
%COREDIAN Calculates core diagonality
%
%[DIA_Rel,DIA_Abs]=CoreDian(C,Fac)
%
%DIA_Rel : Diagonality in percent
%DIA_Abs : Absolute sum of squares of
%          the diagonal elements 

% Copyright (C) 1995-2006  Rasmus Bro & Claus Andersson
% Copenhagen University, DK-1958 Frederiksberg, Denmark,
%
% $ Version 2.00 $ May 2001 $ Changed to array notation $ RB $ Not compiled $

Fac = size(C);
C = reshape(C,Fac(1),prod(Fac(2:end)));


c=size(Fac,2);
d=ones(1,c);
l=min(Fac);

C=C.^2;

DIA_Abs=0;
for j=1:l,
  [i,j]=getindxn(Fac,j*d);
  DIA_Abs = DIA_Abs + C(i,j);
end;

DIA_Rel=100*DIA_Abs/sum(sum(C));
```
