## Supplementary File S1 for "Robust latent-variable interpretation of *in vivo* regression models by nested resampling": coreswdn.m

```
function [SWD_Rel,SWD_Abs]=CoreSWDn(C)
%CORESWDN Calculates the 'core-slice-wise-diagonality'.
%
%
%[SWD_Rel,SWD_Abs]=CoreSWDn(C)
%
%Calculates the 'core-slice-wise-diagonality'.
%To make sense the core C should be quadratic over
%at least two modes. The last mode will always be used.
%
%[SWD_Rel,SWD_Abs]=CoreSWDn(C)
%
%SWD_Rel : Slice-wise diagonality in percent
%SWD_Abs : Absolute sum of squares of
%          the slice-wise diagonal elements 

% Copyright (C) 1995-2006  Rasmus Bro & Claus Andersson
% Copenhagen University, DK-1958 Frederiksberg, Denmark,
%

% $ Version 2.00 $ May 2001 $ Changed to array notation $ RB $ Not compiled $

W = size(C);
C = reshape(C,W(1),prod(W(2:end)));


c=size(W,2);
d=ones(1,c-1);

C=C.^2;

SWD_Abs=0;
if c==3,
   for k=1:W(3);
      tmp_1 = W(2)*(k-1);
      for j=1:W(1),
         ja = j + tmp_1;
         SWD_Abs = SWD_Abs + C(j,ja);
      end;
   end;
else
   for k=1:W(c);
      for j=1:W(1),
         [ia,ja]=gsetindxn(W,[j*d k]);
         SWD_Abs = SWD_Abs + C(ia,ja);
      end;
   end;
end;

SWD_Rel=100*SWD_Abs/sum(sum(C));
```
