## Supplementary File S1 for "Robust latent-variable interpretation of *in vivo* regression models by nested resampling": corevarn.m

```
function [Var_Rel,Var_Abs]=CoreVarn(C)
%COREVARN  Calculates the 'core-variance' 
%
%[Var_Rel,Var_Abs]=CoreVarn(C)
%Calculates the 'core-variance' using not means of
%variables(columns) but the avg of all elements in the matrix
%
%Var_Rel : Variance of squares in percent
%Var_Abs : Absolute variance of squares

% Copyright (C) 1995-2006  Rasmus Bro & Claus Andersson
% Copenhagen University, DK-1958 Frederiksberg, Denmark,
%
C=C.^2;
p=prod(size(C));
mn=sum(C(:))/p;

Var_Abs=sum(sum( (C-mn).^2) );

Var_Rel=100*Var_Abs/(p*(p-1)*mn^2);
```
