## Supplementary File S1 for "Robust latent-variable interpretation of *in vivo* regression models by nested resampling": demos.m

```
function tbxStruct=demos
% DEMOS Demo list for The N-way Toolbox for MATLAB

% Copyright (C) 1995-2006  Rasmus Bro & Claus Andersson
% Copenhagen University, DK-1958 Frederiksberg, Denmark,
%

if nargout==0, 
    demo toolbox 'The N-way Toolbox for MATLAB'; 
    return; 
end

tbxStruct.Name='The N-way Toolbox for MATLAB';
tbxStruct.Type='Toolbox';
tbxStruct.Help= {
' The N-way Toolbox for MATLAB contains state-of-the-art'
' tools for doing multi-way analysis in MATLAB.'};
tbxStruct.DemoList={
'PARAFAC Demo', 'parademo', '',
'Tucker Demo','tuckdemo', '',};
```
