## Supplementary File S1 for "Robust latent-variable interpretation of *in vivo* regression models by nested resampling": derdia3.m

```
function  dC=derdia3(C,n)

%DERDIA3 the derivative of the nth mode of the core 
%
%dC=derdia3(C,n)
%
%This function determines the derivative of the nth mode of
%the core rotation expression for the 3-way case w.r.t.
%maximization of the DIAGONALITY of the core

% Copyright (C) 1995-2006  Rasmus Bro & Claus Andersson
% Copenhagen University, DK-1958 Frederiksberg, Denmark,

W = size(C);
C = reshape(C,W(1),prod(W(2:end)));

dC=zeros(W(n),W(n));
W1=W(1);
W2=W(2);
W3=W(3);

if n==1,
   for a=1:W1,
      idxja = a + W2*(a-1);
      for b=1:W1,
         dC(a,b) = C(b,idxja)*C(a,idxja);
      end;
   end;
end;

if n==2,
   for a=1:W2,
      tmp_2 = W2*(a-1);
      idxja = a + tmp_2;
      for b=1:W2,
         idxjb = b + tmp_2;
         dC(a,b) = C(a,idxjb)*C(a,idxja);
      end;
   end;
end;           	

if n==3,
   for a=1:W3,
      idxja = a + W2*(a-1);
      for b=1:W3,
         idxjb = a + W2*(b-1);
         dC(a,b) = C(a,idxjb)*C(a,idxja);
      end;
   end;
end;    

%---------------------------------------------------------------------------------------
%function  dC=derdia3(C,W,n)
%
%function dC=derdia3(C,W,n)
%
%%This function determines the derivative of the nth mode of
%%the core rotation expression for the 3-way case w.r.t.
%%maximization of the DIAGONALITY of the core
%
%dC=zeros(W(n),W(n));
%
%if n==1,
%  for a=1:W(1),
%    for b=1:W(1),
%      [idxia idxja]=getindxn(W,[a a a]);
%      [idxib idxjb]=getindxn(W,[b a a]);
%      dC(a,b) = C(idxib,idxjb)*C(idxia,idxja);
%    end;
%  end;
%end;
% 
%if n==2,
%  for a=1:W(2),
%    for b=1:W(2),
%      [idxia idxja]=getindxn(W,[a a a]);
%      [idxib idxjb]=getindxn(W,[a b a]);
%      dC(a,b) = C(idxib,idxjb)*C(idxia,idxja);
%    end;
%  end;
%end;           	
%
%if n==3,
%  for a=1:W(3),
%    for b=1:W(3),
%      [idxia idxja]=getindxn(W,[a a a]);
%      [idxib idxjb]=getindxn(W,[a a b]);
%      dC(a,b) = C(idxib,idxjb)*C(idxia,idxja);
%    end;
%  end;
%end;
```
