## Supplementary File S1 for "Robust latent-variable interpretation of *in vivo* regression models by nested resampling": derswd3.m

```
cfunction  dC=derswd3(C,n)

%DERSWD3 for core rotations
%
%function dC=derswd3(C,W,n)
%
%This function determines the derivative of the nth mode of
%the core rotation expression for the 3-way case w.r.t.
%maximization of the slice-wise diagonality of the core
%It must hold that W(1)=W(2), otherwise rearrange the data array

% Copyright (C) 1995-2006  Rasmus Bro & Claus Andersson
% Copenhagen University, DK-1958 Frederiksberg, Denmark,
%
%

W = size(C);
C = reshape(C,W(1),prod(W(2:end)));
dC=zeros(W(n),W(n));
W1=W(1);
W2=W(2);
W3=W(3);

if n==1,
   tmp_1 = W(2)*([1:W3]-1);
   for a=1:W1,
      idxja = a + tmp_1;
      for b=1:W1,
         dC(a,b) = sum(C(b,idxja).*C(a,idxja));
      end;
   end;
end;

if n==2,
   tmp_2 = W2*([1:W3]-1);
   for a=1:W2,
      idxja = a + tmp_2;
      for b=1:W2,
         idxjb = b + tmp_2;
         dC(a,b) = sum(C(a,idxjb).*C(a,idxja));
      end;
   end;           	
end;

%----------------------------------------------------------------------------------
%function  dC=derswd3(C,W,n)
%
%%function dC=derswd3(C,W,n)
%
%%This function determines the derivative of the nth mode of
%%the core rotation expression for the 3-way case w.r.t.
%%maximization of the slice-wise diagonality of the core
%
%dC=zeros(W(n),W(n));
%
%if n==1,
%   for a=1:W(1),
%      for b=1:W(1),
%         dC(a,b)=0;
%         for i_n=1:W(3),
%            [idxia idxja]=getindxn(W,[a a i_n]);
%            [idxib idxjb]=getindxn(W,[b a i_n]);
%            dC(a,b) = dC(a,b) + C(idxib,idxjb)*C(idxia,idxja);
%         end;
%      end;
%   end;
%end;
%
%if n==2,
%   for a=1:W(2),
%      for b=1:W(2),
%         dC(a,b)=0;
%         for i_n=1:W(3),
%            [idxia idxja]=getindxn(W,[a a i_n]);
%            [idxib idxjb]=getindxn(W,[a b i_n]);
%            dC(a,b) = dC(a,b) + C(idxib,idxjb)*C(idxia,idxja);
%         end;
%      end;
%   end;           	
%end;
%
%if n==3, %Obsolete because dC=dC';
%   for a=1:W(3),
%      for b=1:W(3),
%         dC(a,b)=0;
%         for i=1:W(1),
%            [idxia idxja]=getindxn(W,[i i a]);
%            [idxib idxjb]=getindxn(W,[i i b]);
%            [a b idxia idxja idxib idxjb];
%            dC(a,b) = dC(a,b) + C(idxib,idxjb)*C(idxia,idxja);
%         end;
%      end;
%   end;
%end;
```
