## Supplementary File S1 for "Robust latent-variable interpretation of *in vivo* regression models by nested resampling": dervar3.m

```
function  dC=dervar3(C,n)

%DERVAR3 for core rotations
%
%function dC=dervar3(C,n)
%
%This function determines the derivative of the nth mode of
%the core rotation expression for the 3-way case w.r.t.
%maximization of the variance of the core
%
%This version has been optimized for speed, see below for a
%more easy to read scheme.

% Copyright (C) 1995-2006  Rasmus Bro & Claus Andersson
% Copenhagen University, DK-1958 Frederiksberg, Denmark,
%
% $ Version 2.00 $ May 2001 $ Changed to array notation $ RB $ Not compiled $

W = size(C);
C = reshape(C,W(1),prod(W(2:end)));

mc = mean(mean(C.^2));
E = (C.^2 - mc).*C;
dC = zeros(W(n),W(n));
W1 = W(1);
W2 = W(2);
W3 = W(3);

if n==1,
   for a=1:W1,
      for b=1:W1,
         
         dc=0;
         for j=1:W3,
            tmp_0 = W2*(j-1);
            idxja = [1:W2] + tmp_0;
            dc = dc + sum(C(b,idxja).*E(a,idxja));
         end;
         
         dC(a,b) = dc;
      end;
   end;
end;

if n==2,
   for a=1:W2,
      for b=1:W2,
         
         dc=0;
         for j=1:W3,
            tmp_1 = W2*(j-1);
            idxja = a + tmp_1;
            idxjb = b + tmp_1;
            dc = dc + sum(C(:,idxjb).*E(:,idxja));
         end;
         
         dC(a,b) = dc;
      end;
   end;
end;           	

if n==3,
   for a=1:W3,
      tmp_2 = W2*(a-1);
      for b=1:W3,
         tmp_3 = W2*(b-1);
         
         dc=0;
         for j=1:W2,
            idxja = j + tmp_2;
            idxjb = j + tmp_3;
            dc = dc + sum(C(:,idxjb).*E(:,idxja));
         end;
         
         dC(a,b) = dc;
      end;
   end;
end;    


%----------------------------------------------------------------------------------------
%function  dC=dervar3(C,W,n)
%
%%function dC=dervar3(C,W,n)
%
%%This function determines the derivative of the nth mode of
%%the core rotation expression for the 3-way case w.r.t.
%%maximization of the variance of the core
%
%mc=mean(mean(C.^2));
%dC=zeros(W(n),W(n));
%
%if n==1,
%   for a=1:W(1),
%      for b=1:W(1),
%         
%         dc=0;
%         for i=1:W(2),
%            for j=1:W(3),
%               [idxia idxja]=getindxn(W,[a i j]);
%               [idxib idxjb]=getindxn(W,[b i j]);
%               dc = dc + (C(idxia,idxja)^2 - mc)*C(idxib,idxjb)*C(idxia,idxja);
%            end;
%         end;
%         
%         dC(a,b) = dc;
%      end;
%   end;
%end;
%
%if n==2,
%   for a=1:W(2),
%      for b=1:W(2),
%         
%         dc=0;
%         for i=1:W(1),
%            for j=1:W(3),
%               [idxia idxja]=getindxn(W,[i a j]);
%               [idxib idxjb]=getindxn(W,[i b j]);
%               dc = dc + (C(idxia,idxja)^2 - mc)*C(idxib,idxjb)*C(idxia,idxja);
%            end;
%         end;
%         
%         dC(a,b) = dc;
%      end;
%   end;
%end;           	
%
%if n==3,
%   for a=1:W(3),
%      for b=1:W(3),
%         
%         dc=0;
%         for i=1:W(1),
%            for j=1:W(2),
%               [idxia idxja]=getindxn(W,[i j a]);
%               [idxib idxjb]=getindxn(W,[i j b]);
%               dc = dc + (C(idxia,idxja)^2 - mc)*C(idxib,idxjb)*C(idxia,idxja);
%            end;
%         end;
%         
%         dC(a,b) = dc;
%      end;
%   end;
%end;
```
