## Supplementary File S1 for "Robust latent-variable interpretation of *in vivo* regression models by nested resampling": dorrit.pdf

### Factors affecting 3-way modelling (PARAFAC) of fluorescence landscapes

Dorrit Baunsgaard
Food Technology
Department of Dairy and Food Science
The Royal Veterinary and Agricultural University
August 1999

**Preface**

This report present the results of an individual Ph.D.-course in food technology under the supervision of associate professor Rasmus Bro at the Department of Dairy and Food Science at the Royal Veterinary and Agricultural University in the summer 1999. The aim of the course is to examine physical and chemical factors in fluorescence landscape measurements that may influence the performance of the mathematical 3-way model PARAFAC.

### 1. Introduction

#### *1.1 Scope of the project*

The subject of this project is to examine more closely the physical (as opposed to mathematical) obstacles of analysing fluorescence landscape measurements with the use of the 3-way chemometric model, PARAFAC. Fluorescence landscapes of mixtures of four known fluorophores have been measured and PARAFAC has been used to model the fluorescence data. Different approaches have been used to examine the influence of the condition of the data on the model results.

In the report the background of the project is given with a short introduction to fluorescence measurements of mixtures as well as the trilinearity of fluorescence data conforming to the PARAFAC model. Subsequently a list is presented of physical and chemical factors influencing the fluorescence measurements and secondary the PARAFAC modelling. In the experimental part the samples, the fluorescence landscape measurements and the modelling methods are described. The results of the PARAFAC modelling of the data set are then presented and discussed. Finally, a conclusion is given and future projects are suggested.

#### *1.2 Fluorescence measurements of mixtures*

Fluorescence spectroscopy is often used as an analytical technique. Fluorescence has the advantage of selectivity and sensitivity in comparison with other spectroscopic analyses. A fluorophore is characterised as a molecule with a structure that can support an excited electron, which demands a conjugated double bonding structure often including an aromatic system or electron donating heteroatoms (N, O and S). This narrows the number of fluorophores as compared to chromophores (selectivity). The reason for the high sensitivity of fluorescence techniques is that the emission signal is measured above a low background level. This is inherently more sensitive than comparing two relatively large signals as in absorption spectroscopy. The sensitivity of fluorescence techniques is as much as 1000 times more sensitive than absorption spectroscopy. However, when measuring fluorescence of a complex mixture, the difficulties arise. Room temperature fluorescence spectra are relatively broad and featureless and the analysis of a number of components in a mixture will be frustrated by spectral overlaps. An improved way of analysing mixtures is to measure several emission spectra at different excitation wavelengths, thus creating an excitation-emission matrix (EEM) as a landscape that covers the total area of fluorescence. The presentation of data in an EEM format is a good way of visualising the different fluorophores in a sample, but apart from the possibility of characterising distinct peaks, it is difficult to unravel the individual fluorophores in an unknown sample.

##### *1.3 The trilinearity of fluorescence data applied to the PARAFAC model*

For a sample containing a single fluorophore, the elements of an EEM can be approximated as[1]:

(1) $M[\mathrm{j,k}] = \pi(\lambda_j)\kappa(\lambda_j)\Phi_f I_0(\lambda_k) 2.303\varepsilon(\lambda_k)cl$

Where $M[\mathrm{j,k}]$ represents the fluorescence intensity detected at wavelength $\lambda_j$ for excitation at $\lambda_k$; $\pi(\lambda_j)$ is the fraction of fluorescence photons emitted at wavelength $\lambda_j$; $\kappa(\lambda_j)$ expresses the wavelength dependence of the sensitivity of the analysing system on the emission side; $\Phi_f$ is the quantum yield of the fluorescence; $I_0(\lambda_k)$ is the intensity of the monochromatic incident light on the sample (including all effects on $I_0$ from the instrument on the excitation side); $2.303\varepsilon(\lambda_k)cl$ represents the optical density of the sample (the absorbance) and is the product of the molar absorption coefficient $\varepsilon(\lambda_k)$, the concentration $c$ of the fluorophore and the pathlength $l$ through the sample. Equation (1) assumes that the optical density is <<1 for all $\lambda_k$.

When several fluorophores (F) are present in a sample set, the fluorescence intensity can be described as a trilinear expression[2]:

(2) $$M[\mathrm{i, j, k}] = \sum_{\mathrm{f}}^{\mathrm{F}} c_f[\mathrm{i}]\ \pi_f[\mathrm{j}]\varepsilon_f[\mathrm{k}].$$

where $M[\mathrm{i, j, k}]$ represent the fluorescence intensity of the i'th sample detected at wavelength $\lambda_j$ when excited at wavelength $\lambda_k$;. This equation has simplified the terms in equation (1). $c_f[\mathrm{i}] = \Phi_f 2.303cl$ represents the wavelength independent factor of fluorophore f that contain all of the concentration dependence, the term $\pi_f[\mathrm{j}] = \pi(\lambda_j)\kappa(\lambda_j)$ contains all the emission dependent factors and $\varepsilon_f[\mathrm{k}] = I_0(\lambda_k)\varepsilon(\lambda_k)$ represents the excitation dependent factors.

The assumption that the absorbance has to be small still rules as well as there must be no light or energy transfer between the fluorophores.

The three-way model PARAFAC can be used to decompose fluorescence data, which meets the premises of equation (2). PARAFAC was proposed independently by Harshman[3] and Carroll and Chang[4]; the latter named it CANDECOMP. In the PARAFAC model, elements $x$[i, j, k] of a 3-way array **X** are expressed as sums of products of elements ($a_f$[i], $b_f$[j] and $\gamma_f$[k]) from 2-way loading matrices, **A**, **B** and **C**, according to the equation

$$(3) \qquad x[i,j,k] = \sum_{f=1}^{F} a_f[i] b_f[j] \gamma_f[k] + e[i,j,k]$$

where $e$[i,j,k] is the residual term. The parameters of the model are found by least squares fitting of the model.

Fluorescence EEM's of several samples form a 3-way matrix with $I$ samples, $J$ emission wavelengths and $K$ excitation wavelengths that fulfil the immediate requirements of the PARAFAC model. If the trilinearity of the matrix is true, the PARAFAC model can decompose the fluorescence data in F fluorophores identified by unique excitation and emission spectra as well as a sample profile relating the concentration distribution of each fluorophore. However, there is a limit to the number of fluorophores that can be decomposed, if the solutions of the model are expected to be unique, which is defined by Kruskal[5].

##### *1.4 Factors of the fluorescence EEM's and the PARAFAC model*

In reality fluorescence EEM's do not meet the terms of a trilinear model. The chemical conditions of the sample and the physical conditions of the measurements largely influence the recorded fluorescence data. Some of the most important influencing factors are listed below. Only factors affecting solution measurements are listed, since a solution of mixed fluorophores has been chosen as the model system for examining these influencing factors. Solid state measurements introduce a lot of extra perturbations on the recorded spectrum and it is easier to consider these extra effects when the basic problems are understood.

1. *Number of fluorophores*

It is important to consider the number of fluorophores that it is possible to decompose in a PARAFAC model. The more fluorophores, the bigger the risk that their fluorescence response may be perturbed and affect the measured landscapes. Effects such as the inner filter effects (see below) and the quantum yield of the different fluorophores has to be considered. If a fluorophore has a low quantum yield, the measured fluorescence of that compound might "disappear" in the more dominant responses of the other fluorophores, especially if they have spectra in the same wavelength area.

2. *Inner filter effect (concentration quenching)*

It is assumed in the above equations that the optical density of the sample is very low (<0.05). If not, an effect called the inner filter effect or concentration quenching of the sample may influence the measured fluorescence intensity. Considering one fluorophore in the sample, the emission spectrum is distorted by the self-absorption of the emission before reaching the detector reducing the measured fluorescence intensity in the wavelengths where the excitation and the emission spectrum of the fluorophore overlap. When several components are present in the sample, a high optical density of a chromophore can cause re-absorption of the emission of a fluorophore in wavelength areas, where the absorption spectrum of the former overlap the emission spectrum of the latter. The emission spectrum is then fully or partly distorted. The best way of avoiding the inner filter effect is to make sure that the concentration of all fluorophores is sufficiently low, which may be difficult if the sample contains unknown fluorophores.

3. *Temperature*

The fluorescence emission decay may be reduced due to enhanced non-radiative competitive decays (heat). If different fluorophores are not equally effected, the overall measured fluorescence response will change.

4. *pH*

If a change in pH changes the structural composition of a fluorophore, the fluorescence may be shifted in wavelengths or quenched. This effect can be used to change the relative proportions of the response of the fluorophores in a sample and perhaps counteract some of the other effects.

3. *Relation of fluorophores*

Most of the samples of interest are collected in "real life" and not as a model system mixed under controlled conditions in the laboratory. These samples can be very complex consisting of many fluorophores. Some of the fluorophores may be precursors to other fluorescent reaction products such as oxidation products, derivatisation products or thermal reaction products. These products often preserve or slightly change the fluorescent properties of the precursor. As an example, the tryptophan residue in a protein preserves its fluorescence but the spectral position is changing a little depending on the environment of the residue. These types of related fluorophores can be very difficult for the PARAFAC model to separate mathematically due to the often very similar spectral appearance.

5. *Light scattering*

The possibility of interference by scattered exciting light must always be considered in the PARAFAC modelling of fluorescence landscapes. Rayleigh and Raman scattering are the two important scattering effects when measuring the fluorescence of a solution without suspended particles. When a beam of light of a certain frequency passes through a transparent medium, the electrons of the molecules are set into forced vibration and emit a small proportion of scattered light of unchanged frequency, i.e. the Rayleigh scattering. During the scattering process, the vibrations of the nuclei of the atoms in the molecules can cause the nuclei to abstract part of the energy of the beam and convert it to vibrational energy. Part of the scattered light therefore carry energy at changed frequencies due to the nuclei vibrations, and this scattering is the Raman scattering. The Rayleigh scattering is thus appearing at the same wavelength in the emission spectrum as the excitation wavelength, whereas the Raman scattering peak is shifted towards higher wavelengths. The Raman scattering band is weak in intensity and will affect fluorescence landscapes with low fluorescence intensity. The Rayleigh scattering band can be strong in intensity depending on the excitation wavelength and will affect the measured fluorescence intensity, when the emission wavelength is close to the excitation wavelength. The fluorescence landscape is also influenced in the high wavelength side of the emission due to the second order Rayleigh scattering peak. The second order light from the instrument's grating monochromator is the cause of a second order fluorescence response and a second order Rayleigh scattering peak at twice the excitation wavelength.

#### 2. Experimental

##### *2.1 Choice of fluorophores*

In the previous chapter several factors affecting the fluorescence landscapes were listed. It is a very time consuming project to examine in detail all the listed effects, because the fluorescence landscape measurements as well as the PARAFAC modelling are rather time consuming. Therefore, a simple approach has been chosen, where four fluorophores with rather similar spectral properties are mixed in different concentrations. Hopefully, it is thus possible to have the fluorescence of one component influencing the others, which will strain the premises of the model.

**Table 1.** The spectral properties of the four fluorophores in water (pH ≈ 6). $\lambda_{ex}$,max is the wavelength of the excitation maximum, $\lambda_{em}$,max is the wavelength of the emission maximum, ε ($\lambda_{ex}$,max) is the molar absorption coefficient in the excitation max. and $\Phi_f$ is the fluorescence quantum yield.

| | $\lambda_{ex}$,max[a] (nm) | $\lambda_{em}$ ,max[a] (nm) | ε ($\lambda_{ex}$,max)[a] | $\Phi_f$[b] |
|---|---|---|---|---|
| L-phenylalanine | 255 | 285 | 180 | 0.02 |
| L-DOPA | 280 | 320 | 2680 | - |
| 1,4-dihydroxybenzene | 285 | 330 | 2550 | - |
| L-Tryptophan | 275 | 360 | 5500 | 0.14 |

[a] Experimentally determined in this project.

[b] From the literature[6]

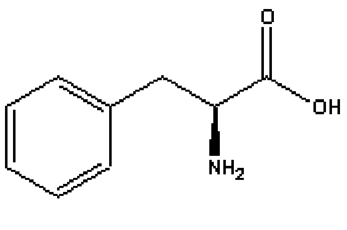

A

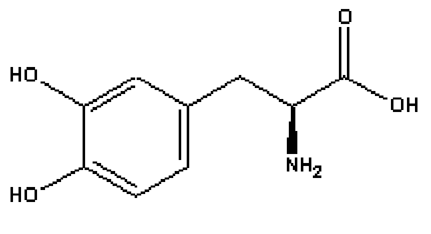

B

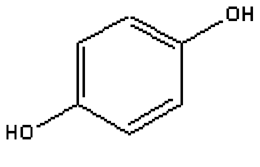

C

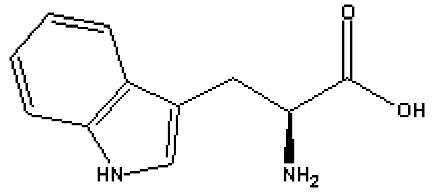

D

**Fig. 1** Molecular structure of the four fluorophores. A. L-phenylalanine, B. L-DOPA, C. 1,4-dihydroxybenzene and D. L-tryptophan.

They four chosen compounds are: L-phenylalanine, L- 3,4-dihydroxyphenylalanine (DOPA), 1,4-dihydroxybenzene and L-tryptophan. In Table 1 the spectral properties of the four fluorophores in aqueous solution are shown and in Fig. 1 the molecular structures are shown.

##### *2.2 The sample set*

**Table 2.** The concentrations of the four fluorophores in the measured samples.

| Sample no. | Hydroquinone ($10^{-6}$ M) | Tryptophan ($10^{-6}$ M) | Phenylalanine ($10^{-6}$ M) | Dopa ($10^{-6}$ M) |
|---|---|---|---|---|
| 1 | 275 | 0 | 0 | 0 |
| 2 | 56 | 0 | 0 | 0 |
| 3 | 28 | 0 | 0 | 0 |
| 4 | 0 | 25 | 0 | 0 |
| 5 | 0 | 16 | 0 | 0 |
| 6 | 0 | 8 | 0 | 0 |
| 7 | 0 | 0 | 5600 | 0 |
| 8 | 0 | 0 | 700 | 0 |
| 9 | 0 | 0 | 0 | 220 |
| 10 | 0 | 0 | 0 | 55 |
| 11 | 0 | 0 | 0 | 5 |
| 12 | 46 | 4 | 2800 | 18 |
| 13 | 17 | 2 | 4700 | 28 |
| 14 | 20 | 1 | 3200 | 8 |
| 15 | 10 | 4 | 3200 | 16 |
| 16 | 6 | 2 | 2800 | 28 |
| 17 | 3.5 | 1 | 350 | 20 |
| 18 | 3.5 | 0.5 | 175 | 20 |
| 19 | 3.5 | 0.25 | 700 | 10 |
| 20 | 1.75 | 4 | 1400 | 5 |
| 21 | 0.875 | 2 | 700 | 2.5 |
| 22 | 28 | 8 | 700 | 40 |
| 23 | 28 | 8 | 350 | 20 |
| 24 | 14 | 8 | 175 | 20 |
| 25 | 0.875 | 8 | 1400 | 2.5 |
| 26 | 1.75 | 8 | 700 | 5 |
| 27 | 3.5 | 2 | 700 | 80 |

A stock solution was made of each compound using Milli-Q water as the solvent. pH was measured for each sample but was not adjusted. Fluorescence landscapes were then measured of a series of samples, where the four components were mixed in different ratios. In Table 2 the concentrations of the four components of the different samples are shown. Considering the molar absorption coefficients in Table 1, it is clear that several of the concentrations used in the samples are above the limit of an optical density of 0.05. This is done to force the effects of concentration quenching on the measured fluorescent response. The concentration limits are: $2800*10^{-6}$ M (phenylalanine), $19*10^{-6}$ M (DOPA), $20*10^{-6}$ M (hydroquinone) and $9*10^{-6}$ M. However, the tryptophan concentration had to be kept below the limit because its high quantum yield made the concentrated samples totally dominate the whole wavelength area.

##### *2.2 Fluorescence measurements*

A Perkin-Elmer LS50 B fluorescence spectrometer was used to measure fluorescence landscapes using excitation wavelengths between 200-350 nm with 5 nm intervals. The emission wavelength range was 200-750 nm. Excitation and emission monochromator slit widths were set to 5 nm, respectively. Scan speed was 1500 nm/min.

##### *2.3 Fluorescence data preprocessing*

All fluorescence landscapes were converted to MATLAB ver. 5.3 using a MATLAB conversion program INCA (ver 1.4,). All data variables were removed during the conversion of the fluorescence data, which were not conforming to trilinear data (see chap.1.4). This include the Rayleigh scattering peaks, first and second order, but also all variables lower than the Rayleigh peak. The second order fluorescence data have also been removed, because the conformity of this area to the trilinearity of the PARAFAC model has not been fully examined. The variable areas concerned are treated as missing in the model, which means that they are not given a value, since they are not part of the defined model, but are iteratively replaced by model estimates during fitting[8]. In Fig. 2a a fluorescence landscape of one the measured samples is shown before any data reduction. Farthest to the right the trace of the first order Rayleigh peaks is seen. Then follows the true fluorescence response bounded on the left by the second order Rayleigh peak trace and on the left of that the second order emission response is shown.

Also the number of excitation and emission wavelengths were reduced to avoid areas where the four fluorophores had no or very little emission. The final sample matrix consisted of 22 samples, 74 emission wavelengths (263-482 nm) and 24 excitation wavelengths (200-315 nm). In Fig. 2b the reduced fluorescence landscape from 2a is shown including the missing areas.

##### *2.4 The PARAFAC algorithm*

The algorithm used in this report is described by Bro[7,8] and the implementation in MATLAB™ is available from The N-way Toolbox for MATLAB at the website http://www.models.kvl.dk. As validation of the models, core consistency diagnostic and split half analyses (if possible) have been used. The non-negativity constraint was applied to some of the models.

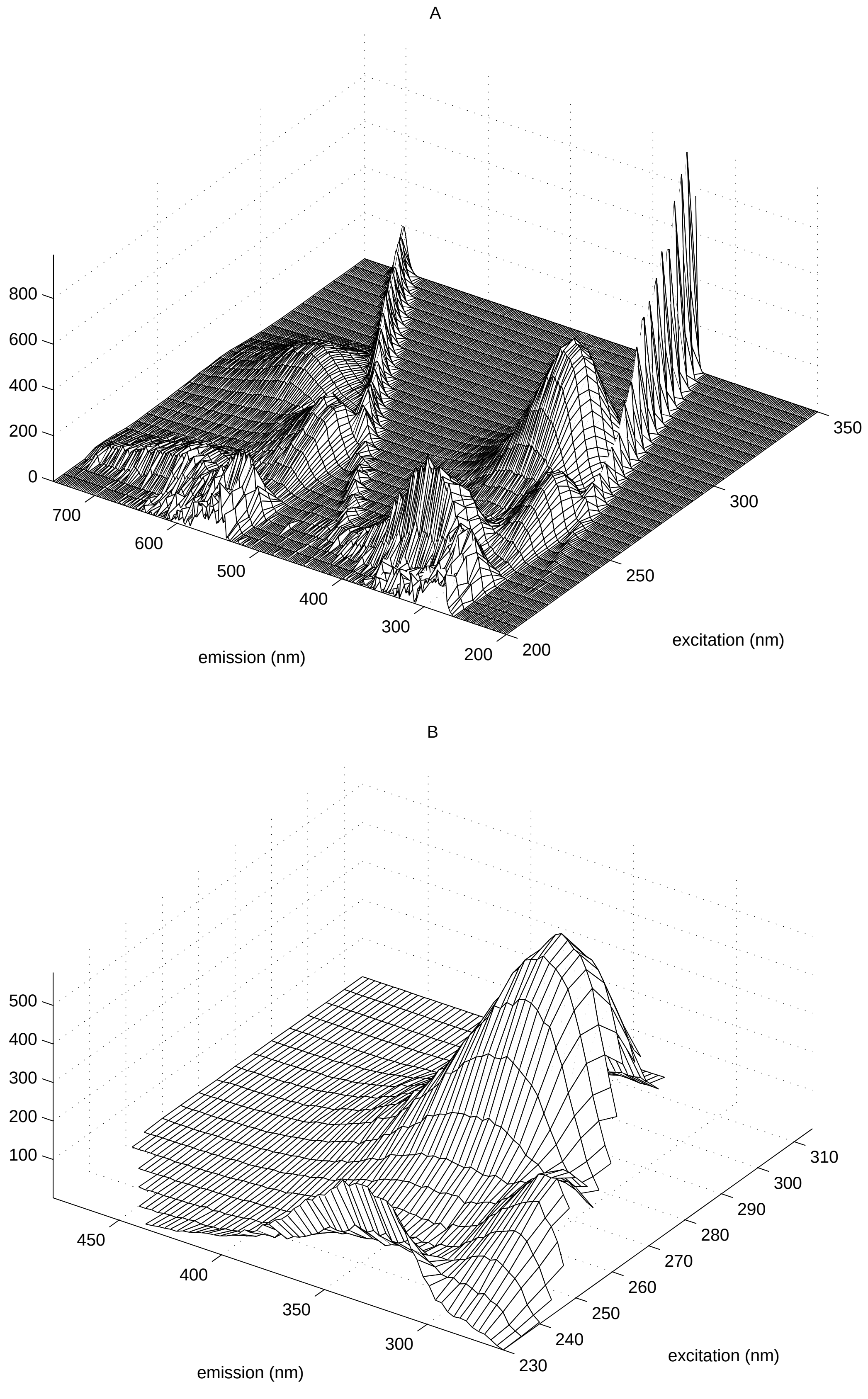

**Fig. 2** Fluorescence landscapes of a sample of mixed fluorophores before (top) and after (bottom) reduction of wavelengths and replacement of non-trilinear data with missing.

#### 3. Results and discussion

##### *3.1 The pure components*

All four fluorophores are modelled individually to check the performance of the model on these four fluorophores and to check the fluorescent purity of the solutions. Four 1-component unconstrained models are fitted using the pure samples measured for each compound, respectively (sample 1-11 in Table 2). In Fig. 3a-d the excitation and emission profiles of the four models are shown.

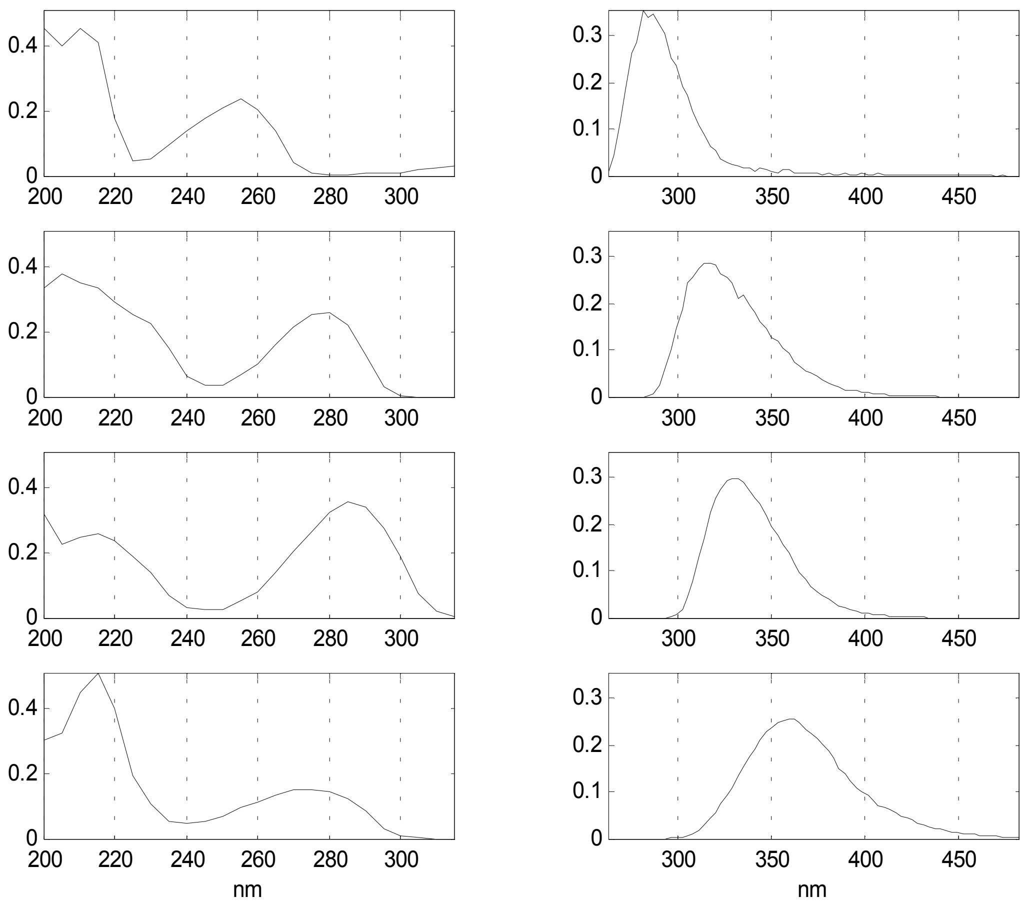

**Fig. 3** Excitation and emission profiles estimated from individual 1-component non-constrained PARAFAC models of the four compounds. From the top: phenylalanine, DOPA, hydroquinone and tryptophan. The spectral profiles are normalised, so that all variance is kept in the sample concentration mode.

All four one-component models were successfully estimated and the spectral profiles have an expected appearance and position. In the area between 200 and 230 nm in the excitation profiles, all four compounds show the extra band of the higher excited states, but the fluorescence landscapes turn out to be very noisy and distorted in the area. This makes the spectral resolution of the model in that part of the profiles difficult and introduce errors in the estimations. The low ultraviolet wavelength area of the measured spectra is very influenced by the condition of the xenon lamp in the instrument and this is probably causing the noise in this area.

The excitation profile of phenylalanine in Fig. 3a shows an estimated response in the area 290-315 nm, where there should not be any measured fluorescence. This might be caused by an impurity, but nothing is found in the fluorescence landscapes. Instead, the response in that area could be the instability of the model in this area due to a large proportion of missing data (see Fig. 2b).

A non-constrained four-component model of all the measured samples of the pure compounds is not so easily modelled as the four individual models. For five repeated models, two of them have a lower fit and a high number of iterations. This is indicative of a more difficult data set. Therefore, the pure sample set is modelled with non-negativity constraint on all three modes, since the excitation and emission wavelengths as well as the sample concentrations all have positive values. This model is shown in Fig. 4. Five repeated models now have the same fit and a low number of iterations. The profiles resemble the individually modelled profiles in Fig. 3. The phenylalanine excitation spectrum still shows some response in the 290-315 nm area due to missing data.

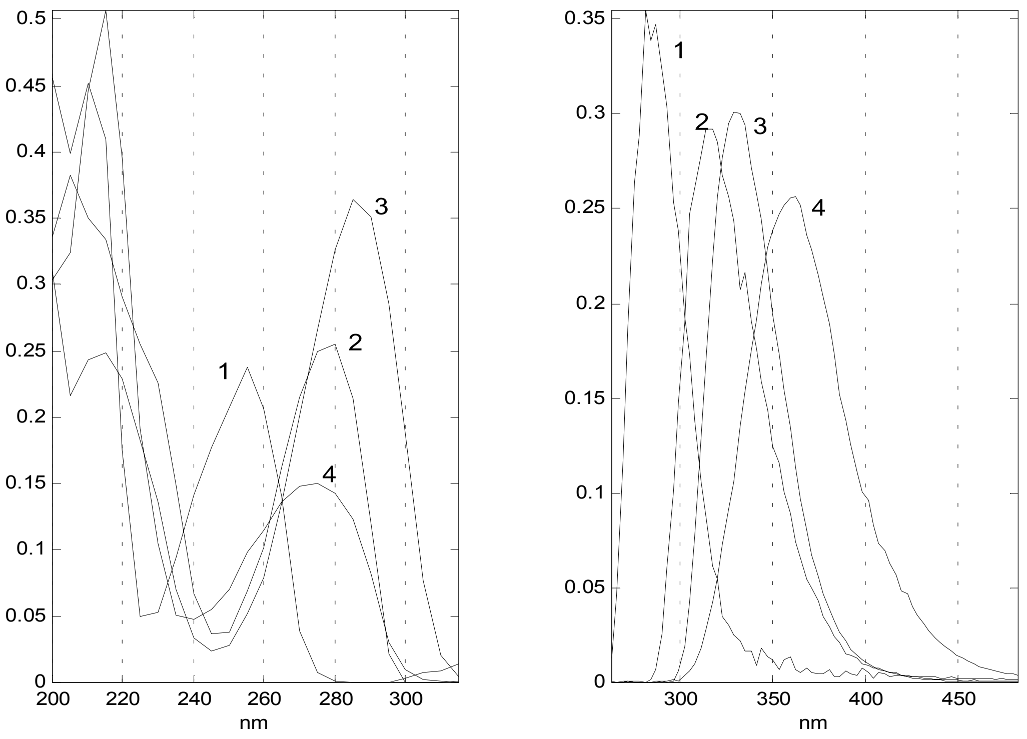

**Fig. 4** Excitation and emission profiles of a four-component non-negativity constraint model of all the pure samples. 1. Phenylalanine; 2. DOPA; 3. Hydroquinone; 4. Tryptophane.

##### *3.2 The models of mixtures*

From the results of modelling samples with only one fluorophore, it is decided, in order to achieve as stable data as possible, to omit the spectra measured in the excitation wavelength area from 200-230 nm and to constrain the models with non-negativity.

16 samples of mixed fluorophores have been measured (sample 12-27 in Table 2). The concentrations of the four compounds have been varied so as to obtain a data set that will explore the effects on the fluorescence measurements when there is fluorophores with high concentration dominating the measured fluorescence. A 4-component non-negativity constrained model of all the 16 mixed samples is fitted (Fig. 5).

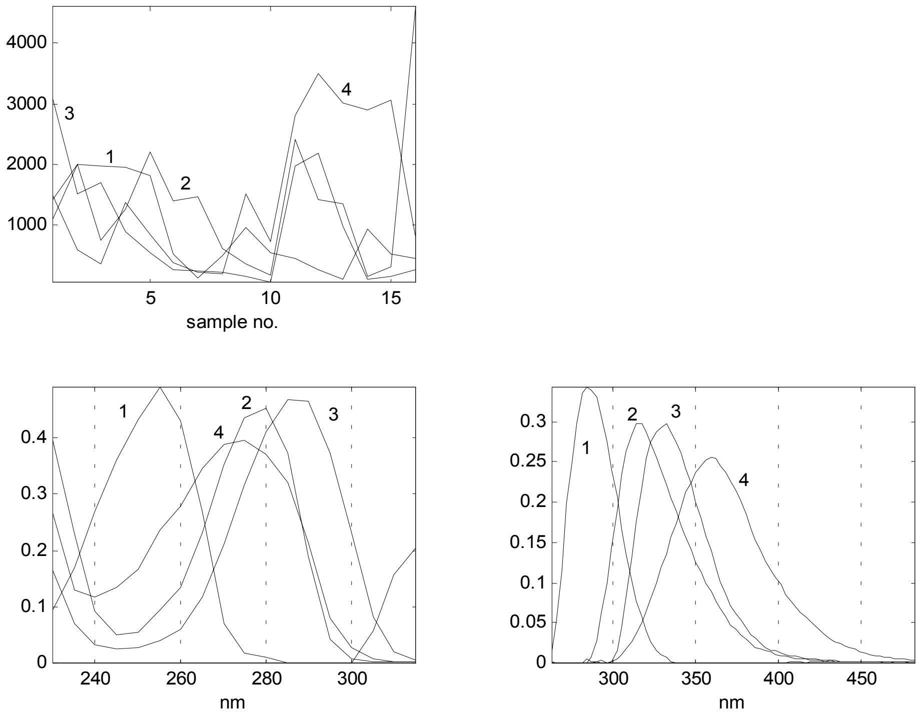

**Fig. 5** Sample concentration, excitation and emission profiles of a four-component non-negativity constraint model of all the mixed samples. The numbers of the spectral profiles are the same as in Fig. 4. The samples 1-16 of the model correspond to the samples 12-27 in Table 2.

Five repeated models only differ in the estimation of the 300-315 nm area in the excitation profile of the phenylalanine. Since the 200-230 nm area was removed, the missing data only covers 17 % of the fluorescence landscape, but it is still enough to destabilise the 300-315 nm area in the model. In this data set, the 300-315 nm area could easily be omitted from the data set without disturbing the model result. However, in fluorescence data covering a larger wavelength area with excitation stretching from the ultraviolet to the visible wavelength area, the missing area cannot be reduced to more than 30-40% and may have a considerable destabilising effect on the models[8].

The emission profiles in Fig. 5 are the same as in Fig. 4 except for a smoother shape probably due to more samples. Three- and five-component models were also generated. The three-component model had acceptable spectral profiles with a DOPA-hydroquinone hybrid, but the fit was significantly lowered with the four-component model. The five-component model had a core consistency of 3 % compared to 98 % for the four-component model. Even though these facts are no conclusive evidence, they do indicate that the four-component model is the appropriate. In Fig. 6 the results of a split half analyses are shown. Four subsets were generated, two by splitting the 16 mixed samples in two halves and two others by splitting the first two subsets and then combining the first half from each subset with each other and the second half with each other. Then a model of each subset is fitted and compared. As Fig. 6 show, the spectral profiles of the subsets are rather similar except for the 330-315 nm area in the excitation profiles.

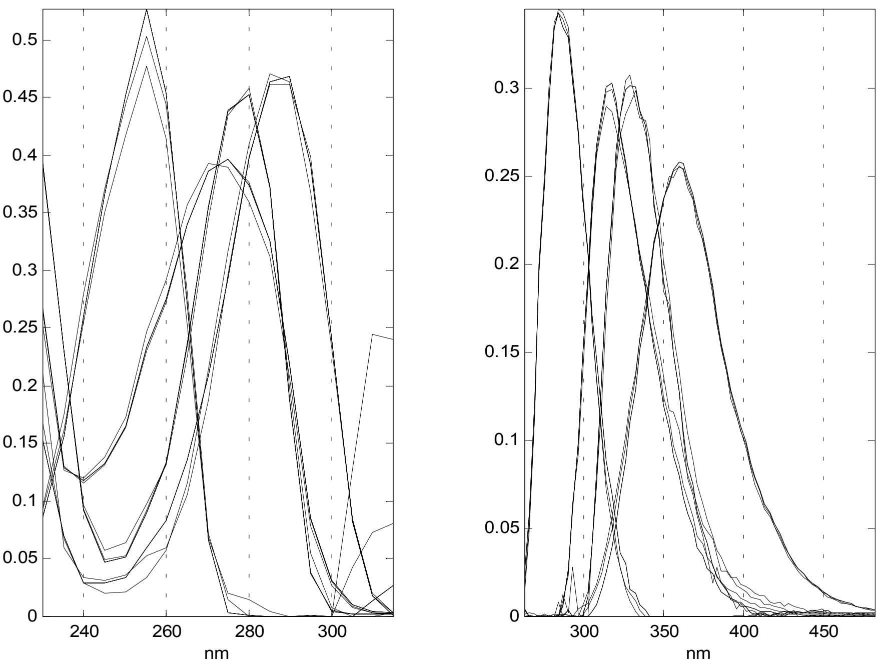

**Fig. 6** Comparison of excitation and emission profiles of four four-component non-negativity constraint models of subsets of the mixed sample set.

A comparison of the estimated sample concentrations for each of the four fluorophores with the concentrations in Table 2, scaling the respective values with the 1/(standard deviation), is shown in Fig. 7a-d.

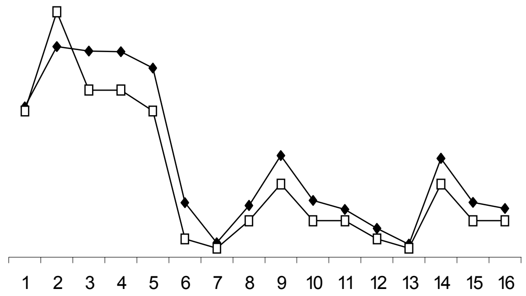

A. Phenylalanine

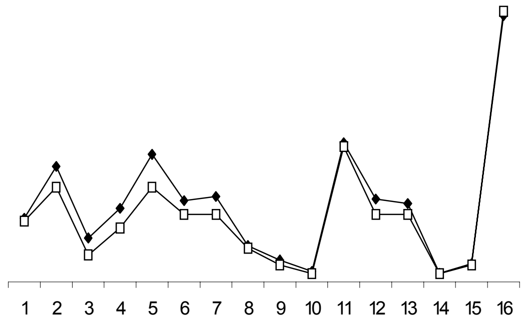

B. DOPA

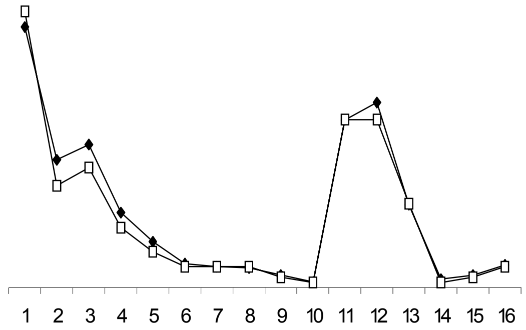

C. Hydroquinone

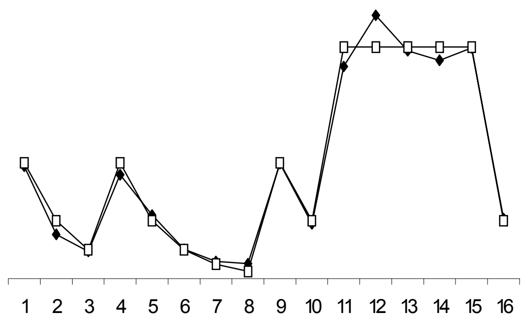

D. Tryptophan

**Fig. 7** Comparison of the estimated (♦) concentrations of the fluorophores modelled from all the mixed samples with the sample concentrations from Table 2 (□).

The trend in the estimations follow the sample concentrations rather well when considering sources of error such as the skill of the analyst. Especially the tryptophan estimations are very close to the true values, whereas the phenylalanine estimations in general differ the most. It might be suggested that this difference could be related to the fact that the fluorescence of phenylalanine is influenced the most and tryptophan the least by missing data due to their position in the landscape, which would ensure a more certain estimation of the tryptophan concentration.

The PARAFAC model estimations of true fluorescence spectra from a mixed sample set are quite impressive when considering the very similar spectral properties of DOPA and hydroquinone. On the other hand, the four fluorophores are all varied from high to low concentrations and the samples set must be considered as an rather ideal data set. It also appears that effects such as the inner filter effect have not had any real influence on the measurements.

In real life it is often difficult or impossible to vary the concentrations of analytes individually in a complex sample. To simulate a more difficult sample set, subsets of the mixed sample set are chosen, which will contain samples of very dominating versus very weak concentration of the fluorophores to strain the model.

A subset of four samples with low concentration of tryptophan was examined (sample 14 and 17-19, Table 2). In Fig. 8 the four-component model results are presented.

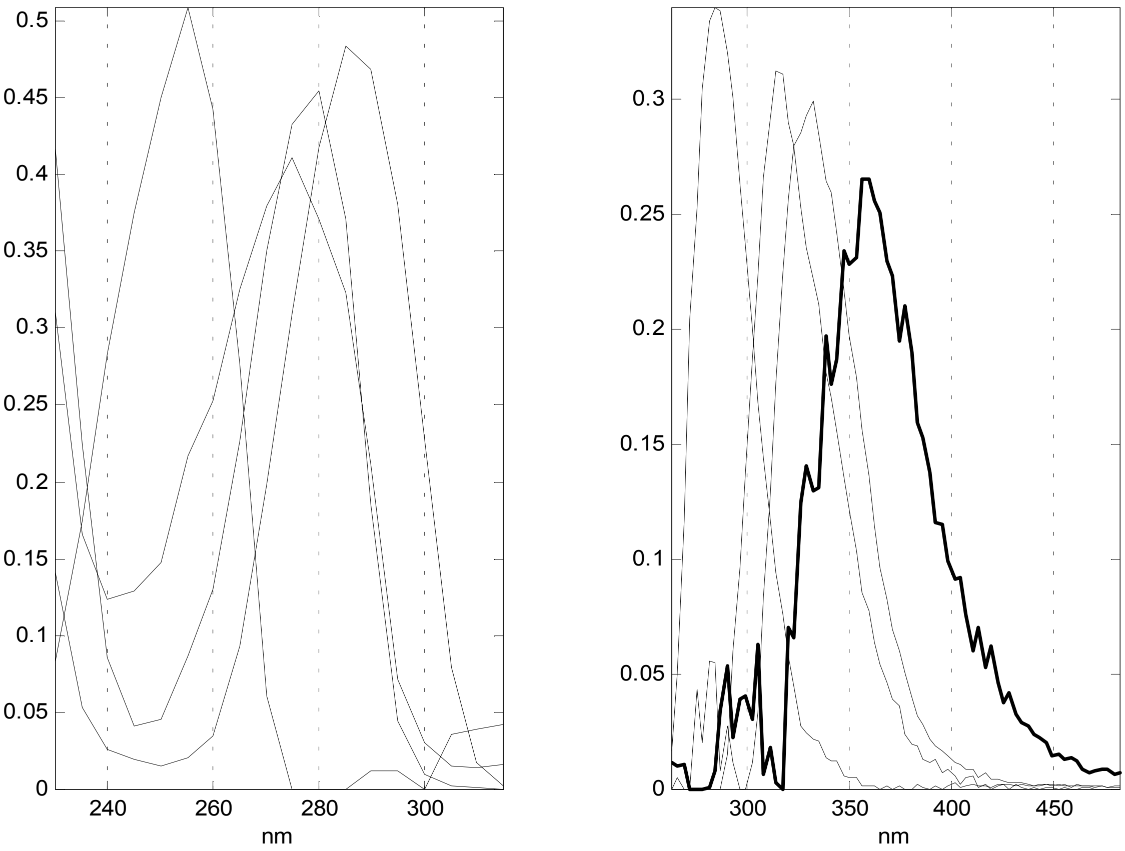

**Fig. 8** Excitation and emission profiles of a four-component non-negativity constraint model of a four-sample subset of mixed samples with low concentrations of tryptophan.

The excitation profile of tryptophan is well estimated considering the low concentrations. The emission profile has the right spectral position, but it is influenced by noise due to the very low fluorescence response. The profile would probably become smoother if a bigger sample set was modelled. The area 263-320 nm in the emission profiles of tryptophan and hydroquinone show extra "noise peaks", which is the "missing data area". The core consistency is rather low (37 %), but it is only 58 % for a three-component model, where the emission profile reveal a phenylalanine profile with a tail in the wavelength area of tryptophan. It is thus a difficult data set, which could probably be improved by more samples.

A subset of four samples, which are low in hydroquinone and DOPA (the samples 20-21 and 25-26 in Table 2), are also modelled. A four-component model was fitted, but five repeated runs showed problems with estimating the true DOPA and hydroquinone spectra (Fig. 9, top). The phenylalanine and tryptophan spectra, marked with bold lines in the plot, are estimated alike for in all five models, whereas DOPA and hydroquinone spectral estimations are very poor.

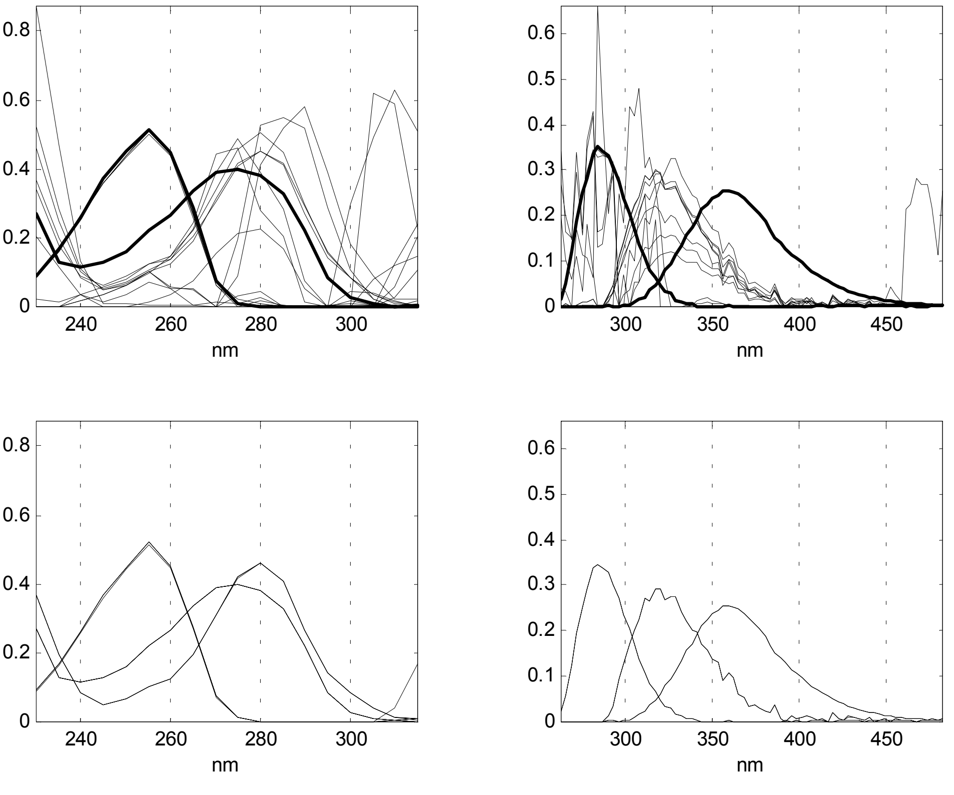

**Fig. 9** Comparison of excitation and emission profiles of five repeated models of a four-sample subset of mixed samples with low concentrations of DOPA and hydroquinone. Top: four-component models, phenylalanine and tryptophan profiles in bold. Bottom: three-component models.

Three-component models (Fig. 9, bottom) had a joint DOPA-hydroquinone profile, but the models converged very quickly in comparison and the core consistency was 97 % compared to 2 % for the four-component model. In a case like this there might

be the risk that the three-component model is chosen as the appropriate model, had the sample set consisted of unknown fluorophores.
It was also attempted to perturb four-component models using a sample set containing very low concentrations of phenylalanine, but in all the models, the phenylalanine profiles were well estimated. The fluorescence peak of phenylalanine is situated so far from the other three fluorophores in the fluorescence landscapes that it is possible to model the fluorophore in defiance of very low concentrations.

#### 4. Conclusion

It has been attempted to generate fluorescence landscapes of mixtures of four fluorophores with similar spectral properties (phenylalanine, DOPA, hydroquinone and tryptophan) that could strain the PARAFAC model so as to examine the effects on the modelling results. It proved to be possible to estimate true excitation and emission spectra of the fluorophores and a reasonable sample concentration profile of even small concentrations in mixed samples when modelling a data set with well distributed concentrations of each fluorophore. When modelling subsets of samples, where one of the fluorophores was only in low concentrations, tryptophan was difficult to model as an individual component. DOPA and hydroquinone, the two fluorophores most alike, failed to be estimated apart when both in low concentrations. Phenylalanine could be modelled in all sample subsets with low concentrations due to the position more apart from the others in the fluorescence landscape, but the large concentration of missing data in that area introduced instability in the estimations of phenylalanine, which affected the overall model performance. Having missing variables in the data set affects the model results more than expected when considering the low percentage (17 %) of missing in this data set as compared to the 40-60 % missing normally encountered with real data[8].

#### 5. Future work

The concentration quenching and the missing data effects were the two factors primarily studied in this fluorophore system. From the impressive goodness of the models of the four fluorophores, it must be concluded that even more complex systems are needed. In the missing data issue, the fluorescence wavelength area should be extended into the visible area; a small system of two ultraviolet and two visible fluorophores could be sufficient. The effect of concentration quenching of the fluorophores was not found in the models fitted here, because the need to obtain fluorescence spectra inside the detection range opposed the really high concentrations needed to explore the quenching effect. A model system containing a powerful chromophore with a concentration of almost total absorbance in the wavelength area of the fluorescence emission, which was gradually diluted, could be used to further examine this effect.

In the need to understand the problems of modelling “real life” samples[8], it can be argued that model systems of pure fluorophores may not fulfil the demands of a really difficult data set. The model systems are too pure and lack the relations of fluorophores as mentioned in chapter 1.4 as well as the influence by other components in the sample that may not fluoresce but are still a part of the matrix. One way of dealing with these problems would be to use “real life” sample with a fairly simple composition and a limited number of fluorophores, some or all of them identified. In this sample, the known fluorophores should then be varied at different concentrations and thus force some modelling problems on the measured fluorescence landscapes. A good example of a suitable “real life” sample is pure sugar in aqueous solution, which has been shown to contain a limited number of fluorophores successfully modelled, two of them identified as the amino acids tyrosine and tryptophan[8].

It is important that all the mentioned factors in chapter 1.4 are thoroughly examined. Appropriate fluorescence data sets should be chosen and probably only focus on one issue at a time, since the results in this project has shown that the sample set should be tailored to ensure the specific effect.
