## Supplementary File S1 for "Robust latent-variable interpretation of *in vivo* regression models by nested resampling": dtld.m

```
function [A,B,C,fit]=dtld(X,F,SmallMode);

%DTLD direct trilinear decomposition
%
% See also:
% 'gram', 'parafac'
%
%
% DIRECT TRILINEAR DECOMPOSITION
%
% calculate the parameters of the three-
% way PARAFAC model directly. The model
% is not the least-squares but will be close
% to for precise data with little model-error
%
% This implementation works with an optimal
% compression using least-squares Tucker3 fitting
% to generate two pseudo-observation matrices that
% maximally span the variation of all samples. per
% default the mode of smallest dimension is compressed
% to two samples, while the remaining modes are 
% compressed to dimension F.
% 
% For large arrays it is fastest to have the smallest
% dimension in the first mode
%
% INPUT
% [A,B,C]=dtld(X,F);
% X is the I x J x K array
% F is the number of factors to fit
% An optional parameter may be given to enforce which
% mode is to be compressed to dimension two
%

% Copyright (C) 1995-2006  Rasmus Bro & Claus Andersson
% Copenhagen University, DK-1958 Frederiksberg, Denmark,
%

% $ Version 2.00 $ May 2001 $ Changed to array notation $ RB $ Not compiled $
% $ Version 1.02 $ Date 28. July 1998 $ Not compiled $
% $ Version 1.03 $ Date 25. April 1999 $ Not compiled $

DimX = size(X);
X = reshape(X,DimX(1),prod(DimX(2:end)));

DontShowOutput = 1;

%rearrange X so smallest dimension is in first mode


if nargin<4
  [a,SmallMode] = min(DimX);
  X = nshape(reshape(X,DimX),SmallMode);
  DimX = DimX([SmallMode 1:SmallMode-1 SmallMode+1:3]);
  Fac   = [2 F F];
else
  X = nshape(reshape(X,DimX),SmallMode);
  DimX = DimX([SmallMode 1:SmallMode-1 SmallMode+1:3]);
  Fac   = [2 F F];
end
f=F;
if F==1;
  Fac   = [2 2 2];
  f=2;
end 


if DimX(1) < 2
  error(' The smallest dimension must be > 1')
end

if any(DimX(2:3)-Fac(2:3)<0)
  error(' This algorithm requires that two modes are of dimension not less the number of components')
end


% Compress data into a 2 x F x F array. Only 10 iterations are used since exact SL fit is insignificant; only obtaining good truncated bases is important
[Factors,Gt]=tucker(reshape(X,DimX),Fac,[0 0 0 0 NaN 10]);
% Convert to old format
Gt = reshape(Gt,size(Gt,1),prod(size(Gt))/size(Gt,1));

[At,Bt,Ct]=fac2let(Factors);

% Fit GRAM to compressed data
[Bg,Cg,Ag]=gram(reshape(Gt(1,:),f,f),reshape(Gt(2,:),f,f),F);

% De-compress data and find A


BB = Bt*Bg;
CC = Ct*Cg;
AA = X*pinv(krb(CC,BB)).';

if SmallMode == 1
  A=AA;
  B=BB;
  C=CC;
elseif SmallMode == 2 
  A=BB;
  B=AA;
  C=CC;
elseif SmallMode == 3
  A=BB;
  B=CC;
  C=AA;
end

fit = sum(sum(abs(X - AA*krb(CC,BB).').^2));
if ~DontShowOutput
  disp([' DTLD fitted raw data with a sum-squared error of ',num2str(fit)])
end
```
