## Supplementary File S1 for "Robust latent-variable interpretation of *in vivo* regression models by nested resampling": eemtimize.m

```
function result = eemtimize(X,maxlv);

%EEMTIMIZE 
% result = eemtimize(X,maxlv);
% result.stats(:,1) = % variance
% result.stats(:,2) = core consistency
% result.ids{f} included samples/variables for f-component model
% 
% IMPORTANT ASSUMPTIONS
% Only first order emission
% Emission >> Raman
% Rayleigh has been properly removed
% Expected to work best on many samples (more than 10 if similar complexity or better more than 30)

for i=1:maxlv
    disp(['DETERMINING A ',num2str(i),'-COMPONENT MODEL'])
    disp(' ')
    [ids{i},models{i}] = optimizemodel(X,i);
    stats(i,:) = [models{i}.detail.ssq.perc models{i}.detail.coreconsistency.consistency];
    disp(' ')
    disp(' ')
end

% Set negative coreconsistencies to zero
stats(find(stats(:,2)<0),2)=0;

% Plot results
subplot(2,1,1)
bar(stats(:,1))
axis tight
title('Percentage variance')
xlabel('Components')

subplot(2,1,2)
bar(stats(:,2))
axis tight
title('Coreconsistency')
xlabel('Components')
   
for i=1:maxlv
    disp(['Samples removed (',num2str(i),'): ',num2str(size(X,1)-sum(ids{i}{1}))'])
end

result.models=models;
result.ids = ids;
result.stats = stats;


function [id,model] = optimizemodel(X,lv);

% SETTINGS
maxnumber_repeats = 8; % Check up to 8 refitted parafac models each time when testing for local minima

% For each of the three criteria (randomness, Q, T2), the max value is
% taken relative to the (robust average) and the max of these three is
% considered. If this value is larger than a certain number (outlier_thres)
% then the corresponding variable/sample is removed.
outlier_thres = 4; % Corresponds to the number of 'standard deviations' away

% However, Q and T2 are further downweighted in order not to remove only
% slightly extreme samples
factor_random = 1;
factor_Q = 1.5;
factor_T2 = 1.5;


% INITIALIZE
dbg = 1; % Show diagnostic messages
if ~isa(X,'dataset');
    X = dataset(X);
end
for i=1:3
    if length(X.include{i})~=size(X,i)
        if dbg
            disp(['Include field in mode ',num2str(i),' modified to include everything']);
        end
        X.include{i}=1:size(X,i);
    end
end

opt = parafac('options');
opt.waitbar = 'off';
opt.display = 'off';
opt.plots = 'off';
for i=1:3
    opt.constraints{i}.nonnegativity = 1;
end
% Options for refitting parafac (use random initial values to chk for local
% minima
optrepeat = opt;
optrepeat.init = 3;


for i=1:3
    id{i} = logical(ones(size(X,i),1));
end

finished = 0;

while ~finished

    % FIT PARAFAC SOME TIMES TO CHECK FOR LOCAL MINIMA. START WITH THREE
    % AND MOVE FURTHER IF THE TWO BEST DO NOT HAVE SIMILAR FIT
    model = parafac(X(id{:}),lv,opt);
    % Check for local minima
    model2 = parafac(X(id{:}),lv,optrepeat);
    if model2.detail.ssq.residual<model.detail.ssq.residual
        second = model;
        model = model2;
        model2 = second;
    end
    model3 = parafac(X(id{:}),lv,optrepeat);
    if model3.detail.ssq.residual<model.detail.ssq.residual
        second = model;
        model = model3;
        model3 = second;
    end
    % Find the second best fitting model
    if model3.detail.ssq.residual<model2.detail.ssq.residual
        second = model3;
    else
        second = model2;
    end
    % Check if two best are the same
    if abs(model.detail.ssq.residual-second.detail.ssq.residual)/model.detail.ssq.residual>(100*optrepeat.stopcrit(1))
        format long
        if dbg
            [model.detail.ssq.residual;second.detail.ssq.residual]
            100*optrepeat.stopcrit(1)
            disp('Local minima detected. Will try to refit PARAFAC a few more times to find global minimum')
        end
        globalfound=0;
        while ~globalfound
            for j=4:maxnumber_repeats
                model2 = parafac(X(id{:}),lv,optrepeat);
                if model2.detail.ssq.residual<second.detail.ssq.residual
                    if model2.detail.ssq.residual<model.detail.ssq.residual
                        second=model;
                        model = model2;
                    else
                        second = model2;
                    end
                end
                if abs(model.detail.ssq.residual-second.detail.ssq.residual)/model.detail.ssq.residual<100*optrepeat.stopcrit(1)
                    if dbg
                        disp(['Found the minimum after ',num2str(j),' models'])
                    end
                    globalfound = 1; % Stop refitting - found two identical with lowest fit
                end
            end

        end
    end

    % WITH THE GIVEN SOLUTION, CALCULATE EXCITATION NOISINESS, T2 and Q
    % Residuals
    xhat = datahat(model);
    e = X.data(id{:})-xhat;

    % Find average randomness of emission spectra
    clear Y
    for i=1:size(e,3);
        E = squeeze(e(:,:,i));
        E = E(:,~isnan(sum(E)));
        d=diff(E');
        a=sum((d.*d));
        Y(:,i)= a(:);
    end

    devex = madc(Y);
    ave_devex = median(devex);
    size_randEM = devex/(factor_random*ave_devex);
    [m(1),num(1)] = max(size_randEM);

    % Find T2 sizes
    t2 = model.tsqs{1};
    t2 = t2/(factor_T2*median(t2));
    [m(2),num(2)] = max(t2);

    % Find Q sizes
    Q = model.ssqresiduals{1};
    Q = Q/(factor_Q*median(Q));
    [m(3),num(3)] = max(Q);

    [a,b]=max(m);
    if a>outlier_thres
        if b==1
            idd = find(id{3});
            disp(['Removing excitation ',num2str(idd(num(1))),' due to noisy behavior'])
            id{3}(idd(num(1)))=0;
        elseif b==2
            idd = find(id{1});
            disp(['Removing sample ',num2str(idd(num(2))),' due to high T2'])
            id{1}(idd(num(2)))=0;
        elseif b==3
            idd = find(id{1});
            disp(['Removing sample ',num2str(idd(num(3))),' due to high Q'])
            id{1}(idd(num(3)))=0;
        end
    else
        disp('Done with this one')
        finished = 1;
    end
end
```
