## Supplementary File S1 for "Robust latent-variable interpretation of *in vivo* regression models by nested resampling": explcore.m

```
function [Comb,ExplVariat]=explcore(G,n);

%EXPLCORE For interpretation of cores and arrays
%
%
% function [Comb,ExplVariat]=explcore(G,Fac,n);
% 'explcore.m'
%
% This algorithm requires access to:
% 'two2n.m' 
%
% [Comb,ExplVariat,ExplVarian]=explcore(G,n);
%
% G         : Core array from Tucker3 model
% n         : Show only the 'n' largest factor combinations.
%

% Copyright (C) 1995-2006  Rasmus Bro & Claus Andersson
% Copenhagen University, DK-1958 Frederiksberg, Denmark,
%

% $ Version 2.00 $ May 2001 $ Changed to array notation $ RB $ Not compiled $
% $ Version 1.03 $ Date 28. Oct 1999 $ Not compiled $ 'improved help'
% $ Version 1.02 $ Date 17. Sep 1998 $ Not compiled $

DimG = size(G);
G = reshape(G,DimG(1),prod(DimG(2:end)));
Fac = DimG;

if ~exist('n'),
    n=10;
end;

if n>length(G(:));
    n=length(G(:));
end;

C=length(Fac(1,:));
Par=zeros(1,C);

ssgunc=sum(G(:).^2);

fprintf('Col1: Number in list\n');
fprintf('Col2: Index to elements\n');
fprintf('Col3: Explained variation (sum of squares) of the core.\n'); 
fprintf('Col4: Core entry.\n'); 
fprintf('Col5: Sq. core entry.\n'); 


for l=1:n,
    [i j]=max(G(:).^2);
    
    [a b]=find(G==G(j));
    a=a(1);
    b=b(1);
    Par=two2n(Fac,[a b]);
    
    fprintf('%2i    ',l);
    fprintf('(');
    for c=1:C-1,
        fprintf('%2i,',Par(c));
    end;
    Comb(l,:)=Par;
    ExplVariat(l)=100*G(a,b).^2/ssgunc;
    fprintf('%2i) %15.5f%% %15.5f  %15.5f\n',Par(C),ExplVariat(l),G(a,b),G(a,b).^2);
    G(a,b)=0;

end; 
```
