## Supplementary File S1 for "Robust latent-variable interpretation of *in vivo* regression models by nested resampling": fac2let.m

```
function [A,B,C,D,E,F,G,H,I,J,K,L,M,N,O,P,Q,R,S,T,U,V,X,Y,Z]=fac2let(Factors,DimX);

%FAC2LET Convert 'Factors' to component matrices
%
%	
% [A,B,C]=fac2let(Factors);
% [A,B,C,D]=fac2let(Factors);
% [A,B,C,D,E]=fac2let(Factors);
%             .....
% [A,B,C,...,Z]=fac2let(Factors);
%
% This algorithm applies to the N-way case (2<N<25).
%

% Copyright (C) 1995-2006  Rasmus Bro & Claus Andersson
% Copenhagen University, DK-1958 Frederiksberg, Denmark,
%

% $ Version 1.02 $ Date 28. July 1998 $ Not compiled $
% $ Version 2.00 $ May 2001 $ Changed to array notation $ RB $ Not compiled $

Txt='ABCDEFGHIJKLMNOPQRSTUVXYZ';

if nargin==2
  order = length(DimX);
  F     = prod(length(Factors))/sum(DimX);
  for i=1:order
    start = sum(DimX(1:i-1))*F+1;
    endd = sum(DimX(1:i))*F;
    eval([Txt(i) ,'= reshape(Factors(',num2str(start),':',num2str(endd),'),',num2str(DimX(i)),',',num2str(F),');']);
  end
else
  for i = 1:length(Factors)
    eval([Txt(i) ,'= Factors{i};']);
  end
end
```
