## Supplementary File S1 for "Robust latent-variable interpretation of *in vivo* regression models by nested resampling": fastnnls.m

```
function [x,w] = fastnnls(XtX,Xty,tol)

%FASTNNLS Fast version of built-in NNLS
%	b = fastnnls(XtX,Xty) returns the vector b that solves X*b = y
%	in a least squares sense, subject to b >= 0, given the inputs
%       XtX = X'*X and Xty = X'*y.
%
%	A default tolerance of TOL = MAX(SIZE(X)) * NORM(X,1) * EPS
%	is used for deciding when elements of b are less than zero.
%	This can be overridden with b = fastnnls(X,y,TOL).
%
%	[b,w] = fastnnls(XtX,Xty) also returns dual vector w where
%	w(i) < 0 where b(i) = 0 and w(i) = 0 where b(i) > 0.
%
%
%	L. Shure 5-8-87 Copyright (c) 1984-94 by The MathWorks, Inc.
%
%  According to Bro & de Jong, J. Chemom, 1997, 11, 393-401

% Copyright (C) 1995-2006  Rasmus Bro & Claus Andersson
% Copenhagen University, DK-1958 Frederiksberg, Denmark,
%

% initialize variables


if nargin < 3
    tol = 10*eps*norm(XtX,1)*max(size(XtX));
end
[m,n] = size(XtX);
P = zeros(1,n);
Z = 1:n;
x = P';
ZZ=Z;
w = Xty-XtX*x;

% set up iteration criterion
iter = 0;
itmax = 30*n;

% outer loop to put variables into set to hold positive coefficients
while any(Z) & any(w(ZZ) > tol)
    [wt,t] = max(w(ZZ));
    t = ZZ(t);
    P(1,t) = t;
    Z(t) = 0;
    PP = find(P);
    ZZ = find(Z);
    nzz = size(ZZ);
    z(PP')=(Xty(PP)'/XtX(PP,PP)');
    z(ZZ) = zeros(nzz(2),nzz(1))';
    z=z(:);
% inner loop to remove elements from the positive set which no longer belong

    while any((z(PP) <= tol)) & iter < itmax

        iter = iter + 1;
        QQ = find((z <= tol) & P');
        alpha = min(x(QQ)./(x(QQ) - z(QQ)));
        x = x + alpha*(z - x);
        ij = find(abs(x) < tol & P' ~= 0);
        Z(ij)=ij';
        P(ij)=zeros(1,length(ij));
        PP = find(P);
        ZZ = find(Z);
        nzz = size(ZZ);
        z(PP)=(Xty(PP)'/XtX(PP,PP)');
        z(ZZ) = zeros(nzz(2),nzz(1));
        z=z(:);
    end
    x = z;
    w = Xty-XtX*x;
end

x=x(:);
```
