## Supplementary File S1 for "Robust latent-variable interpretation of *in vivo* regression models by nested resampling": fnipals.m

```
function [T,P]=fnipals(X,w,T)

%FNIPALS nipals algorithm for PCA
% 
% function [T,P]=fnipals(X,w,T)
%
% 'fnipals.m'
%
% This algorithm requires the presence of:
% 'missmean.m' 
%
% ----------------------------------------------------
%        Find eigenvectors according to NIPALS
% ----------------------------------------------------
%
% [T,P]=fnipals(X,w,T);
% [T,P]=fnipals(X,w);
%
% T is found so that X = T*P', s.t ||T||=1 and T'T=I
%
% X        : The matrix to be decomposed.
% w        : Number of factors to extract.
%            If w is high (perhaps>20) consider using SVD.
% T        : Initial guess of the solution, optional.
%            If T is not specified, a little time will
%            be used on finding orthogonal random 
%            starting values.
%
% You may want to calculate P afterwards by typing 'P=X*T'.
% Note that the T returned is orthonormal.
% Calculation of P is left of this implementation to save FLOP's.
% It handles missing values NaNs (very dispersed, less than 15%)
% If the problem is small enough you would prefer the SVD rather
% than NIPALS for finding T. NIPALS may be inaccurate when
% extracting too many factors, i.e., many more than the rank 
% of X. 

%scalar ConvLim WarnLim ItMax a b i

% $ Version 1.01 $ Date 18. June 1998 $ Not compiled $

% Copyright (C) 1995-2006  Rasmus Bro & Claus Andersson
% Copenhagen University, DK-1958 Frederiksberg, Denmark,
%


ConvLim=1e-12;
WarnLim=1e-4;
ConvLimMiss=100*ConvLim;
ItMax=100;

filename='fnipals.m';

[a b]=size(X);

if (w>a | w>b) | w<1,
    help(filename);
    error(['Error in ' filename ': Number of factors to extract is invalid!'])
end;

np=isnan(X);
MissingExist=any(np);

if ~exist('T'),
    T=orth(randn(a,w));
end;

if exist('P'),
    P=[];
end;

if ~MissingExist
    if (size(T) == [a w]),
        if a>b,
            P=X'*T;
            l2=Inf;
            Z=X'*X;
            for i=1:w,
                p=P(:,i);
                d=1;
                it=0;
                while (d>ConvLim) & (it<ItMax),
                    it=it+1;
                    p=Z*p;
                    l1=sqrt(p'*p);
                    p=p/l1;
                    d=(l1-l2)^2;
                    l2=l1;
                end;
                P(:,i)=sqrt(l1)*p;
                Z=Z-P(:,i)*P(:,i)';
                WarnLim=sqrt(l1)/1000;
                if it>=ItMax & d>WarnLim,
                    disp('FNIPALS, High-X: Iterated up to the ItMax limit!')
                    disp('FNIPALS, High-X: The solution has not converged!')
                end;
            end;
            T=X*P;
        else
            P=[];
            l2=Inf;
            Z=X*X';
            for i=1:w,
                t=T(:,i); 
                d=1;
                it=0;
                while (d>ConvLim) & (it<ItMax),
                    it=it+1;
                    t=Z*t;
                    l1=sqrt(t'*t);
                    t=t/l1;
                    d=(l1-l2).^2;
                    l2=l1;
                end;
                T(:,i)=sqrt(l1)*t;
                Z=Z-T(:,i)*T(:,i)';
                WarnLim=sqrt(l1)/1000;
                if it>=ItMax & d>WarnLim,
                    disp('FNIPALS, Wide-X: Iterated up to the ItMax limit!')
                    disp('FNIPALS, Wide-X: The solution has not converged!')
                end;
            end;
        end;
        T=gsm(T);
    else
        error(['Error in ' filename ': Number of factors to extract is invalid!'])
    end;
else
    MissIdx=find(np);
    [i j]=find(np);
    mnx=missmean(X)/2;
    mny=missmean(X')/2;
    n=size(i,1);
    for k=1:n,
        i_i=i(k);
        j_j=j(k);
        X(i_i,j_j) = mny(i_i) + mnx(j_j);
    end;
    mnz=(missmean(mnx)+missmean(mny))/2;
    
    ssmisold=sum(sum( X(MissIdx).^2 ));
    sstotold=sum(sum( X.^2 ));
    ssrealold=sstotold-ssmisold;
    iterate=1;
    while iterate
        
        if (size(T) == [a w]),
            if a>b,
                P=X'*T;
                l2=Inf;
                Z=X'*X;
                for i=1:w,
                    p=P(:,i);
                    d=1;
                    it=0;
                    while (d>ConvLim) & (it<ItMax),
                        it=it+1;
                        p=Z*p;
                        l1=sqrt(p'*p);
                        p=p/l1;
                        d=(l1-l2)^2;
                        l2=l1;
                    end;
                    P(:,i)=sqrt(l1)*p;
                    Z=Z-P(:,i)*P(:,i)';
                    WarnLim=sqrt(l1)/1000;
                    if it>=ItMax & d>WarnLim,
                        disp('FNIPALS, High-X: Iterated up to the ItMax limit!')
                        disp('FNIPALS, High-X: The solution has not converged!')
                    end;
                end;
                T=X*P;
            else
                P=[];
                l2=Inf;
                Z=X*X';
                for i=1:w,
                    t=T(:,i); 
                    d=1;
                    it=0;
                    while (d>ConvLim) & (it<ItMax),
                        it=it+1;
                        t=Z*t;
                        l1=sqrt(t'*t);
                        t=t/l1;
                        d=(l1-l2).^2;
                        l2=l1;
                    end;
                    T(:,i)=sqrt(l1)*t;
                    Z=Z-T(:,i)*T(:,i)';
                    WarnLim=sqrt(l1)/1000;
                    if it>=ItMax & d>WarnLim,
                        disp('FNIPALS, Wide-X: Iterated up to the ItMax limit!')
                        disp('FNIPALS, Wide-X: The solution has not converged!')
                    end;
                end;
            end;
            T=gsm(T);
        else
            error(['Error in ' filename ': Number of factors to extract is invalid!'])
        end;
        
        P=X'*T;
        Xm=T*P';
        X(MissIdx)=Xm(MissIdx);
        ssmis=sum(sum( Xm(MissIdx).^2 ));
        sstot=sum(sum( X.^2 ));
        ssreal=sstot-ssmis;
        if abs(ssreal-ssrealold)<ConvLim*ssrealold & abs(ssmis-ssmisold)<ConvLimMiss*ssmisold,
            iterate=0;
        end;
        ssrealold=ssreal;
        ssmisold=ssmis;   
    end;
end;
T=gsm(T);
```
