## Supplementary File S1 for "Robust latent-variable interpretation of *in vivo* regression models by nested resampling": fnnls.m

```
cfunction b = fastnnls(x,y,tol,b)

% $ Version 1.02 $ Date 28. July 1998 $ Not compiled $
%
% See also:
% 'unimodal' 'monreg' 'fastnnls'
%
%  FASTNNLS Fast non-negative least squares
%  The inputs are the matrix of predictor variables (x),
%  vector of predicted variable (y), and optional inputs
%  tolerance on the size of a regression coefficient that is
%  considered zero (tol), and initial guess for the regression
%  vector (b0). The output is the non-negatively constrained
%  least squares solution (b).
%
%  If tol is set to 0, the default tolerance will be used.
%  
%  FASTNNLS is fastest when a good estimate of the regression
%  vector is input. This eliminates much of the computation
%  involved in determining which coefficients will be nonzero
%  in the final regression vector. This makes it very useful
%  in alternating least squares routines. Note that the input
%  b0 must be a feasible (i.e. nonnegative) solution.
%
%  The FASTNNLS algorithm is based on the one developed by
%  Bro and de Jong, J. Chemometrics, Vol. 11, No. 5, 393-401, 1997
%
%I/O: b = fastnnls(x,y,tol,b0);
%
%See also: MCR, PARAFAC

% Copyright (C) 1995-2006  Rasmus Bro & Claus Andersson
% Copenhagen University, DK-1958 Frederiksberg, Denmark,
%
[m,n] = size(x);
if (nargin < 3 | tol == 0)
  tol = max(size(x))*norm(x,1)*eps;
end
if nargin < 4
  b = zeros(n,1);
end

p = logical(zeros(1,n));
p(find(b>0)) = ones(size(find(b>0)));
r = ~p;

sp = x(:,p)\y;
b(find(p)) = sp;
while min(sp) < 0
  b(find(b<0)) = zeros(size(find(b<0)));
  p = logical(zeros(1,n));
  p(find(b>0)) = ones(size(find(b>0)));
  r = ~p;
  sp = x(:,p)\y;
  b(find(p)) = sp;
end

w = x'*(y-x*b);
[wmax,ind] = max(w);
flag = 0;
while (wmax > tol & any(r))
  p(ind) = 1; 
  r(ind) = 0;
  sp = x(:,p)\y;
  while min(sp) < -tol
    tsp = zeros(n,1);
    tsp(find(p)) = sp;  
    fb = find(b);
    rat = b(fb)./(eps+(b(fb)-tsp(fb)));
    alpha = min(rat(rat>tol));
    b = b + alpha*(tsp-b);
    p = b > tol;
    r = ~p; 
    sp = x(:,p)\y;
  end
  b(find(p)) = sp;
  w = x'*(y-x*b);  
  [wmax,ind] = max(w);
  if p(ind)
    wmax = 0;
  end
end
```
