## Supplementary File S1 for "Robust latent-variable interpretation of *in vivo* regression models by nested resampling": getindxn.m

```
function [i,j]=getindxn(R,Idx);
%GETINDXN
%
%[i,j]=getindxn(R,Idx)


% Copyright (C) 1995-2006  Rasmus Bro & Claus Andersson
% Copenhagen University, DK-1958 Frederiksberg, Denmark,
%
l=size(Idx,2);

i=Idx(1);
j=Idx(2);

if l==3,
  j = j + R(2)*(Idx(3)-1);
 else
  for q = 3:l,
    j = j + prod(R(2:(q-1)))*(Idx(q)-1);
  end;
end;
```
