## Supplementary File S1 for "Robust latent-variable interpretation of *in vivo* regression models by nested resampling": gram.m

```
function [A,B,C]=gram(X1,X2,F);

%GRAM generalized rank annihilation method
%
% [A,B,C]=gram(X1,X2,F);
%
% cGRAM - Complex Generalized Rank Annihilation Method
% Fits the PARAFAC model directly for the case of a 
% three-way array with only two frontal slabs.
% For noise-free trilinear data the algorithm is exact.
% If input is not complex, similarity transformations
% are used for assuring a real solutions (Henk Kiers
% is thanked for providing the similarity transformations)
% 
% INPUTS:
% X1    : I x J matrix of data from observation one
% X2    : I x J matrix of data from observation two
% Fac   : Number of factors
% 
% OUTPUTS:
% A     : Components in the row mode (I x F)
% B     : Components in the column mode (J x F)
% C     : Weights for each slab; C(1,:) are the component 
%         weights for first slab such that the approximation
%         of X1 is equivalent to X1 = A*diag(C(1,:))*B.'
%

% Copyright (C) 1995-2006  Rasmus Bro & Claus Andersson
% Copenhagen University, DK-1958 Frederiksberg, Denmark,
%
% $ Version 1.02 $ Date 28. July 1998 $ Not compiled $
% $ Version 1.03 $ Date 22. February 1999 $ Not compiled $

  IsReal=0; % If complex data, complex solutions allowed.
  if all(isreal(X1))&all(isreal(X2))
     IsReal=1;
  end

  % Find optimal bases in F x F subspace
  [U,s,V]=svd(X1+X2);
  U=U(:,1:F);
  V=V(:,1:F);

  % Reduce to an F x F dimensional subspace
  S1=U'*X1*V;
  S2=U'*X2*V;

  % Solve eigenvalue-problem and sort according to size
  [k,l]=eig(S1\S2);
  l=diag(l);
  ii=abs(l)>eps;
  k=k(:,ii);
  l=l(ii);
  p=length(l);
  [l,ii]=sort(l);
  j=p:-1:1;
  l=l(j);
  l=diag(l);
  k=k(:,ii(j));
  k=k/norm(k);

  if IsReal % Do not allow complex solutions if only reals are considered
    T1=eye(F);
    T2=eye(F);
    [rhok,argk]=complpol(k);
    [rhol,argl]=complpol(diag(l));
    j=1;
    while j<=F
      if abs(imag(l(j,j)))<.00000001  % real eigenvalue
        if abs(imag(k(1,j)))>=.00000001 % complex eigenvector
          T1(j,j)=exp(i*argk(1,j));
        end;
      end;
      if abs(imag(l(j,j)))>=.00000001  % j-th and j+1-th are complex eigenvalues
        c=argk(1,j)+argk(1,j+1);
        T1(j,j)=exp(i*c/2);
        T1(j+1,j+1)=exp(i*c/2);
        T2(j:j+1,j:j+1)=[1 1;i -i];
        j=j+1;
      end;
      j=j+1;
    end;
    k=real(k/T1/T2);
    l=T2*T1*l/T1/T2;
    l=real(diag(diag(l)));
  end

  C(2,:)=ones(1,F);
  C(1,:)=diag(l)';
  A = U*S1*k;
  B=V/k';

C=(pinv(krb(B,A))*[X1(:) X2(:)]).';
```
