## Supplementary File S1 for "Robust latent-variable interpretation of *in vivo* regression models by nested resampling": gsm.m

```
function [E]=GSM(V);
%GSM orthogonalization
%
% [E]=GSM(V);
% GS   Gram-Schmidt Method for orthogonalisation
%      An orthonormal basis spanning the columns of V is returned in E.
% 
%      This algorithm does not use pivoting or any other
%      stabilization scheme. For a completely safe orthogonalization
%      you should use 'ORTH()' though is may take triple the time.
%      'GSM()' is optimized for speed and requies only minimum storage
%      during iterations. No check of rank is performed on V!
%

% Copyright (C) 1995-2006  Rasmus Bro & Claus Andersson
% Copenhagen University, DK-1958 Frederiksberg, Denmark,

[m n]=size(V);

%Allocate space for the basis
E=zeros(m,n);

%The first basis vector is taken directly from V
s=sqrt(sum(V(:,1).^2));
E(:,1)=V(:,1)/s;

%Find the other basis vectors as orthogonals to
%the already determined basis by projection
for k=2:n,
  f=V(:,k)-E(:,1:(k-1))*(E(:,1:(k-1))'*V(:,k));
  s=sqrt(sum(f.^2));
  if s<eps,
    E(:,k)=0*f;   %set to zeros
  else
    E(:,k)=f/s;   %normalize
  end;
end;
```
