## Supplementary File S1 for "Robust latent-variable interpretation of *in vivo* regression models by nested resampling": ini.m

```
function [Factors]=ini(X,Fac,MthFl,IgnFl)
%INI initialization of loadings
%
% function [Factors]=ini(X,Fac,MthFl,IgnFl)
%
% This algorithm requires access to:
% 'gsm' 'fnipals' 'missmult'
%
% ---------------------------------------------------------
%                    Initialize Factors 
% ---------------------------------------------------------
%
% [Factors]=ini(X,Fac,MthFl,IgnFl);
% [Factors]=ini(X,Fac,MthFl);
%
% X        : The multi-way data.
% Fac      : Vector describing the number of factors
%            in each of the N modes.
% MthFl    : Method flag indicating what kind of
%            factors you want to initiate Factors with:
%            '1' : Random values, orthogonal
%            '2' : Normalized singular vectors, orthogonal
% IgnFl    : This feature is only valid with MthFl==2.
%            If specified, these mode(s) will be ignored,
%            e.g. IgnFl=[1 5] or IgnFl=[3] will
%            respectively not initialize modes one and 
%            five, and mode three.
% Factors  : Contains, no matter what method, orthonormal
%            factors. This is the best general approach to
%            avoid correlated, hence ill-posed, problems.
%
% Note that it IS possible to initialize the factors to have
% more columns than rows, since this may be required by some
% PARAFAC models. If this is required, the 'superfluos' 
% columns will be random and orthogonal columns.
% This algorithm automatically arranges the sequence of the
% initialization to minimize time and memory consumption.
% Note, if you get a warning from NIPALS about convergence has
% not been reached, you can simply ignore this. With regards 
% to initialization this is not important as long as the
% factors being returned are in the range of the eigensolutions.

% $ Version 1.02 $ Date 30 Aug 1999 $ Not compiled $
% $ Version 1.0201 $ Date 21 Jan 2000 $ Not compiled $ RB removed orth of additional columns
% $ Version 2.00 $ May 2001 $ Changed to array notation $ RB $ Not compiled $

% Copyright (C) 1995-2006  Rasmus Bro & Claus Andersson
% Copenhagen University, DK-1958 Frederiksberg, Denmark,
%
format long
format compact

DimX = size(X);
X = reshape(X,DimX(1),prod(DimX(2:end)));

% Assign intermediaries
Show=0;
rand('seed',sum(100*clock));
MissingExist=any(isnan(X(:)));

% Initialize system variables
N=size(Fac,2);
if N==1,
    Fac=Fac*ones(1,size(DimX,2));
end;
N=size(Fac,2);

FIdx0=zeros(1,N);
FIdx1=zeros(1,N);
latest=1;
for c=1:N,
    if Fac(c)==-1,
        FIdx0(c)=0;
    else
        FIdx0(c)=latest;
        latest=latest+Fac(c)*DimX(c);
        FIdx1(c)=latest-1;
    end;
end;

% Check inputs
if ~exist('IgnFl'),
    IgnFl=[0];
end;

%Random values
if MthFl==1,
    for c=1:N,
        A=orth(rand( DimX(c) , min([Fac(c) DimX(c)]) ));
        %B=[A orth(rand(DimX(c),Fac(c)-DimX(c)))]; 
        B=[A rand(DimX(c),Fac(c)-DimX(c))]; 
        Factors(FIdx0(c):FIdx1(c))=B(:)';
    end;
    if Show>=1,
        fprintf('ini.m : Initialized using random values.\n');
    end;
else
    %Singular vectors
    Factors=rand(1,sum(~(Fac==-1).*DimX.*Fac));
    if MthFl==2 | MthFl==3 
        [A Order]=sort(Fac);
        RedData=X;
        CurDimX=DimX;
        for k=1:N,
            c=Order(k);
            if Fac(c)>0,
                for c1=1:c-1;
                    newi=CurDimX(c1+1);
                    newj=prod(CurDimX)/CurDimX(c1+1);
                    RedData=reshape(RedData',newi,newj);
                end;
                Op=0;
                if MissingExist | (Op==0 & Fac(c)<=5 & (50<min(size(RedData)) & min(size(RedData))<=120)),
                    %Need to apply NIPALS
                    t0=clock;
                    A=reshape(Factors(FIdx0(c):FIdx1(c)),DimX(c),Fac(c));
                    if MissingExist
                        MissIdx=find(isnan(RedData));
                        [A,P]=fnipals(RedData,min([Fac(c) DimX(c)]),A);
                        Xm=A*P';
                        RedData(MissIdx)=Xm(MissIdx);
                        MissingExist=0;
                     else
                        [A]=fnipals(RedData,min([Fac(c) DimX(c)]),A);
                    end;
                    B=[A orth(rand(DimX(c),Fac(c)-DimX(c)))];
                    Factors(FIdx0(c):FIdx1(c))=B(:)';
                    t1=clock;
                    if Show>=2,
                        disp(['ini.m: NIPALS used ' num2str(etime(t1,t0)) ' secs. on mode ' int2str(c)]),
                    end;
                    Op=1;
                end;
                if Op==0 & (120<min(size(RedData)) & min(size(RedData))<Inf),
                    %Need to apply Gram-Schmidt
                    t0=clock;
                    C=RedData*RedData';
                    A=reshape(Factors(FIdx0(c):FIdx1(c)),DimX(c),Fac(c));
                    for i=1:3,
                        A=gsm(C*A);
                    end;
                    B=[A orth(rand(DimX(c),Fac(c)-DimX(c)))];
                    Factors(FIdx0(c):FIdx1(c))=B(:)';
                    t1=clock;
                    if Show>=2,
                        disp(['ini.m: GS used ' num2str(etime(t1,t0)) ' secs. on mode ' int2str(c)]),
                    end;
                    Op=1;
                end;
                if Op==0 & (0<min(size(RedData)) & min(size(RedData))<=200),
                    %Small enough to apply SVD
                    t0=clock;
                    if max(size(RedData))<1000
                       [U S A]=svd(RedData',0);
                    else
                       [U S A]=svds(RedData');
                    end
                    A=A(:,1:min(size(A,2),min([Fac(c) DimX(c)])));
                    if size(A,2)<Fac(c)
                      A = [A rand(size(A,1),Fac(c)-size(A,2))];
                    end
                    n_ = Fac(c)- min([Fac(c) DimX(c)]);
                    if n_>0,
                       a = rand(DimX(c),n_);
                       if DimX(c)>=n_
                          a = orth(a);
                       else
                          a = orth(a')';
                       end;
                       B=[A a];
                    else 
                       Factors(FIdx0(c):FIdx1(c))=A(:)';
                    end;
                    
                    t1=clock;
                    if Show>=2,
                        disp(['ini.m: SVD used ' num2str(etime(t1,t0)) ' secs. on mode ' int2str(c)]),
                    end;
                    Op=1;
                end;
                CurDimX(c)=min([Fac(c) DimX(c)]);
                if MissingExist,
                    RedData=missmult(A',RedData);
                else
                    RedData=A'*RedData;
                end;
                %Examine if re-ordering is necessary
                if c~=1,
                    for c1=c:N,
                        if c1~=N,
                            newi=CurDimX(c1+1);
                            newj=prod(CurDimX)/newi;
                        else
                           newi=CurDimX(1);
                            newj=prod(CurDimX)/newi;
                        end;
                        RedData=reshape(RedData',newi,newj);
                    end;
                end;
            end;
        end;
        if Show>=1,
            fprintf('ini.m : Initialized using SVD and projection.\n');
        end;
    end;
end,
format
% Convert to new format
clear ff,id1 = 0;
for i = 1:length(DimX) 
   id2 = sum(DimX(1:i).*Fac(1:i));
   ff{i} = reshape(Factors(id1+1:id2),DimX(i),Fac(i));id1 = id2;
end
Factors = ff;
```
