## Supplementary File S1 for "Robust latent-variable interpretation of *in vivo* regression models by nested resampling": inituck.m

```
function [Factors]=inituck(X,Fac,MthFl,IgnFl)
%INITUCK initialization of loadings
%
% function [Factors]=inituck(X,Fac,MthFl,IgnFl)
%
% This algorithm requires access to:
% 'gsm' 'fnipals' 'missmult' 'missmean'
%
% ---------------------------------------------------------
%        Initialize Factors for the Tucker3 model
% ---------------------------------------------------------
%
% [Factors]=inituck(X,Fac,MthFl,IgnFl);
% [Factors]=inituck(X,Fac);
%
% X        : The multi-way data array.
% Fac      : Vector describing the number of factors
%            in each of the N modes.
% MthFl    : Method flag indicating what kind of
%            factors you want to initiate Factors with:
%            '1' : Random values, orthogonal
%            '2' : Normalized singular vectors, orthogonal
%            '3' : SVD with successive projections 
% IgnFl    : This feature is only valid with MthFl==2.
%            If specified, these mode(s) will be ignored,
%            e.g. IgnFl=[1 5] or IgnFl=[3] will
%            respectively not initialize modes one and 
%            five, and mode three.
% Factors  : Contains, no matter what method, orthonormal
%            factors. This is the best general approach to
%            avoid correlated, hence ill-posed, problems.
%
% The task of this initialization program is to find acceptable
% guesses to be used as starting point in the 'TUCKER.M' program.
% Note that it IS possible to initialize the factors to have
% more columns than rows, since this may be required by some
% models. If this is required, the 'superfluos' 
% columns will be random and orthogonal.
% This algorithm automatically arranges the sequence of the
% initialization to minimize time and memory consumption.
% If you get a warning from a NIPALS algorithm about convergence has
% not been reached, you can simply ignore this. With regards 
% to initialization this is not important as long as the
% factors being returned are in the range of the eigensolutions.

% Copyright (C) 1995-2006  Rasmus Bro & Claus Andersson
% Copenhagen University, DK-1958 Frederiksberg, Denmark,

% $ Version 2.00 $ May 2001 $ Changed to array notation $ RB $ Not compiled $
% $ Version 1.00 $ Date 24. May 1998 $ Not compiled $

format long
format compact
DimX = size(X);
X = reshape(X,DimX(1),prod(DimX(2:end)));


MissingExist=any(isnan(X(:)));

% Initialize system variables
N=size(Fac,2);
FIdx0=zeros(1,N);
FIdx1=zeros(1,N);
latest=1;
for c=1:N,
   if Fac(c)==-1,
      FIdx0(c)=0;
   else
      FIdx0(c)=latest;
      latest=latest+Fac(c)*DimX(c);
      FIdx1(c)=latest-1;
   end;
end;

% Check inputs
if ~exist('IgnFl'),
   IgnFl=[0];
end;

%Random values
if MthFl==1,
   for c=1:N,
      A=orth(rand( DimX(c) , min([Fac(c) DimX(c)]) ));
      B=[A orth(rand(DimX(c),Fac(c)-DimX(c)))]; 
      Factors(FIdx0(c):FIdx1(c))=B(:)';
   end;
end;

%Singular vectors
%Factors=rand(1,sum(~(Fac==-1).*DimX.*Fac)); %Matlab 4.2 compatibility
Factors=rand(1,sum((Fac~=-1).*DimX.*Fac)); %Matlab 4.2 compatibility
if MthFl==2 | MthFl==3 
   
   %Remove, in a fast way, the missing values by
   %approximations as means of columns and rows
   if MissingExist,
      [i j]=find(isnan(X));
      mnx=missmean(X)/3;
      mny=missmean(X')/3;
      n=size(i,1);
      for k=1:n,
         i_=i(k);
         j_=j(k);
         X(i_,j_) = mny(i_) + mnx(j_);
      end;
      mnz=(missmean(mnx)+missmean(mny))/2;
      p=find(isnan(X));
      X(p)=mnz;
   end;
   
   [A Order]=sort(Fac);
   RedData=X;
   CurDimX=DimX;
   for k=1:N,
      c=Order(k);
      if Fac(c)>0,
         for c1=1:c-1;
            newi=CurDimX(c1+1);
            newj=prod(CurDimX)/CurDimX(c1+1);
            RedData=reshape(RedData',newi,newj);
         end;
         Op=0;
         if Op==0 & Fac(c)<=5 & (10<min(size(RedData)) & min(size(RedData))<=120),
            %Need to apply NIPALS
            A=reshape(Factors(FIdx0(c):FIdx1(c)),DimX(c),Fac(c));
            A=fnipals(RedData,min([Fac(c) DimX(c)]),A);
            B=[A orth(rand(DimX(c),Fac(c)-DimX(c)))];
            Factors(FIdx0(c):FIdx1(c))=B(:)';
            Op=1;
         end;
         if Op==0 & (120<min(size(RedData)) & min(size(RedData))<Inf),
            %Need to apply Gram-Schmidt
            C=RedData*RedData';
            A=reshape(Factors(FIdx0(c):FIdx1(c)),DimX(c),Fac(c));
            for i=1:3,
               A=gsm(C*A);
            end;
            B=[A orth(rand(DimX(c),Fac(c)-DimX(c)))];
            Factors(FIdx0(c):FIdx1(c))=B(:)';
            Op=1;
         end;
         if Op==0 & (0<min(size(RedData)) & min(size(RedData))<=120),
            %Small enough to apply SVD
            [U S A]=svd(RedData',0);
            A=A(:,1:min([Fac(c) DimX(c)]));
            B=[A orth(rand(DimX(c),Fac(c)-DimX(c)))];
            Factors(FIdx0(c):FIdx1(c))=B(:)';
            Op=1;
         end;
         CurDimX(c)=min([Fac(c) DimX(c)]);
         RedData=A'*RedData;
         %Examine if re-ordering is necessary
         if c~=1,
            for c1=c:N,
               if c1~=N,
                  newi=CurDimX(c1+1);
                  newj=prod(CurDimX)/newi;
               else
                  newi=CurDimX(1);
                  newj=prod(CurDimX)/newi;
               end;
               RedData=reshape(RedData',newi,newj);
            end;
         end;
      end;
   end;
end;
format

% Convert to new format
clear ff
id1 = 0;

for i = 1:length(DimX) 
   
   if Fac(i)~=-1
      id2 = sum(DimX(1:i).*Fac(1:i).*(Fac(1:i)~=-1));
      ff{i} = reshape(Factors(id1+1:id2),DimX(i),Fac(i));
      id1 = id2;
   else
      ff{i}=[];
   end
end
Factors = ff;
```
