## Supplementary File S1 for "Robust latent-variable interpretation of *in vivo* regression models by nested resampling": kr.m

```
function AB = kr(A,B);
%KR Khatri-Rao product
%
% The Khatri - Rao product
% For two matrices with similar column dimension the khatri-Rao product
% is kr(A,B) = [kron(A(:,1),B(:,1)) .... kron(A(:,F),B(:,F))]
% 
% I/O AB = kr(A,B);
%
% kr(A,B) equals ppp(B,A) - where ppp is the triple-P product = 
% the parallel proportional profiles product which was originally 
% suggested in Bro, Ph.D. thesis, 1998

% Copyright (C) 1995-2006  Rasmus Bro & Claus Andersson
% Copenhagen University, DK-1958 Frederiksberg, Denmark,
%

% $ Version 1.02 $ Date 28. July 1998 $ Not compiled $
% $ Version 2.00 $ May 2001 $ Changed to array notation $ RB $ Not compiled $
% $ Version 2.01 $ May 2001 $ Error in helpfile - A and B reversed $ RB $ Not compiled $

disp('KR.M is obsolete and will be removed in future versions. ')
disp('use KRB.M instead.')


[I,F]=size(A);
[J,F1]=size(B);

if F~=F1
   error(' Error in kr.m - The matrices must have the same number of columns')
end

AB=zeros(I*J,F);
for f=1:F
   ab=B(:,f)*A(:,f).';
   AB(:,f)=ab(:);
end
```
