## Supplementary File S1 for "Robust latent-variable interpretation of *in vivo* regression models by nested resampling": maxvar3.m

```
function [Gv,ORot1,ORot2,ORot3]=maxvar3(G,MinRot,ConvLim,Options)
%MAXVAR3 maximize core squared variance
%
%[Gv,Ov1,Ov2,Ov3]=maxvar3(G,MinRot,ConvLim,Options)
%
%This m-file rotates a core to maximum sq. variance
%This algorithm uses inner re-iterations and
%includes a resampling-test for global maximum
%
%G           : Core to be rotated
%MinRot      : Minium number of iterations used to
%              check consistency of optimal value. {5}
%ConvLim     : Convergence limit, {1e-5}
%Options     : N-way TOOLBOX options list
%Gv          : Rotated core
%Ov1,Ov2,Ov3 : Orthogonal rotation matrices for the modes
%
%Upon rotation it holds that G = Ov1*Gv*ckron(Ov3',Ov2')
%
%[Gv,Ov1,Ov2,Ov3]=maxvar3(G)

% $ Version 2.00 $ May 2001 $ Changed to array notation $ RB $ Not compiled $

% Copyright (C) 1995-2006  Rasmus Bro & Claus Andersson
% Copenhagen University, DK-1958 Frederiksberg, Denmark,
%
W = size(G);
G = reshape(G,W(1),prod(W(2:end)));

format compact
rand('state',sum(100*clock));

if ~exist('MinRot') | isempty(MinRot) | MinRot<1,
   MinRot=5;
end;

if ~exist('ConvLim') | isempty(ConvLim) | ConvLim<eps,
   ConvLim=1e-5;
end;

if ~exist('Options') | isempty(Options),
   show=1;
end;

of0=corevarn(G);
O1=eye(W(1));
ORot1=O1;

show=2;

I2=eye(W(2));
O2=I2;
ORot2=O2;

I3=eye(W(3));
O3=I3;
ORot3=O3;

itmax=500;
itmin=3;
C=G;
of_max=0;
curr_of=of0;
in_conv_crit=1e-2;
Rot=0;
Rot_conv=0;

while Rot_conv==0,
   
   Rot=Rot+1;
   if Rot>=MinRot,
      Rot_conv=1;
   end;
   
   it=1;
   conv=0;
   
   ORot1=ORot1*O1;
   ORot2=ORot2*O2;
   ORot3=ORot3*O3;
   C=O1'*C*ckron(O3,O2);
   
   while conv==0,
      
      i_a=0;
      conv_a=0;
      while ~conv_a,

         i_a=i_a+1;
         dC=dervar3(reshape(C,W),1);
         [U D V]=svd(dC,0);
         O1=V*U';
         C=O1'*C;
         ORot1=ORot1*O1;
         
         of=corevarn(C);
         if show>1,
            fprintf('Maxvar3.m: Course %i, Mode %i, iteration %2i : {VOS=%10.6f%%}\n',Rot,1,i_a,of);
         end;
         if of-curr_of<in_conv_crit*curr_of,
            conv_a=1;
         end;
         curr_of=of;
      end;
      
      i_b=0;
      conv_b=0;
      while ~conv_b,
         i_b=i_b+1;
         
         dC=dervar3(reshape(C,W),2);
         [U D V]=svd(dC,0);
         O2=V*U';
         ORot2=ORot2*O2;
         C=C*ckron(I3,O2);
         
         of=corevarn(C);
         if show>1,
            fprintf('Maxvar3.m: Course %i, Mode %i, iteration %2i : {VOS=%10.6f%%}\n',Rot,2,i_b,of);
         end;
         if of-curr_of<in_conv_crit*curr_of,
            conv_b=1;
         end;
         curr_of=of;
      end;
      
      i_c=0;
      conv_c=0;
      while ~conv_c,
         i_c=i_c+1;
         
         dC=dervar3(reshape(C,W),3);
         [U D V]=svd(dC,0);
         O3=V*U';
         ORot3=ORot3*O3;
         C=C*ckron(O3,I2);
         
         of=corevarn(C);
         if show>1,
            fprintf('Maxvar3.m: Course %i, Mode %i, iteration %2i : {VOS=%10.6f%%}\n',Rot,3,i_c,of);
         end;
         if of-curr_of<in_conv_crit*curr_of,
            conv_c=1;
         end;
         curr_of=of;
      end;
      
      if itmin<it & of-of0<=of0*ConvLim,
         conv=1;
      end;
      
      of0=of;
      
      it=it+1;
      if it>itmax,
         conv=1;
         if show>0,
            fprintf('Maxvar3.m: Max. number of iterations exceeded.\n');
         end;
      end;
      
   end;
   
   O1=orth(rand(W(1),W(1))-0.5);  
   O2=orth(rand(W(2),W(2))-0.5);  
   O3=orth(rand(W(3),W(3))-0.5);  
   
   of_list(Rot)=of;
   ORot_list(Rot,:)=[reshape(ORot1,1,W(1).^2) reshape(ORot2,1,W(2).^2) reshape(ORot3,1,W(3).^2)]; 
   if 1< Rot & Rot>=MinRot,
      p=sort(of_list);
      if p(Rot)*0.99>p(Rot-1),
         Rot_conv=0;
         if show>0,
            fprintf('Maxvar3.m: More rotations must be performed to estimate the global maximum.\n');
            fprintf('%s\n',mat2str(p,7));
         end;
      end;
   end;
   
   Gv=C;
end;

[p i]=max(of_list);
ORot1=reshape(ORot_list(i,1:W(1)^2),W(1),W(1));
ORot2=reshape(ORot_list(i,(1+W(1)^2):(W(1)^2+W(2)^2)),W(2),W(2));
ORot3=reshape(ORot_list(i,(1+W(1)^2+W(2)^2):(W(1)^2+W(2)^2+W(3)^2)),W(3),W(3));
Gv=ORot1'*G*ckron(ORot3,ORot2);
format

Gv = reshape(Gv,W);
```
