## Supplementary File S1 for "Robust latent-variable interpretation of *in vivo* regression models by nested resampling": missmult.m

```
function [X]=missmult(A,B)

%MISSMULT product of two matrices containing NaNs
%
%[X]=missmult(A,B)
%This function determines the product of two matrices containing NaNs
%by finding X according to
%     X = A*B
%If there are columns in A or B that are pur missing values,
%then there will be entries in X that are missing too.
%
%The result is standardized, that is, corrected for the lower
%number of contributing terms.
%
%Missing elements should be denoted by 'NaN's


% Copyright (C) 1995-2006  Rasmus Bro & Claus Andersson
% Copenhagen University, DK-1958 Frederiksberg, Denmark,
%
%INBOUNDS
%REALONLY

[ia ja]=size(A);
[ib jb]=size(B);
X=zeros(ia,jb);

one_arry=ones(ia,1);
for j=1:jb,
   p=one_arry*B(:,j)';
   tmpMat=A.*p;
   X(:,j)=misssum(tmpMat')';
end;
```
