## Supplementary File S1 for "Robust latent-variable interpretation of *in vivo* regression models by nested resampling": misssum.m

```
function [mm]=misssum(X,def)
%MISSSUM sum of a matrix X with NaN's
%
%[mm]=misssum(X,def)
%
%This function calculates the sum of a matrix X.
%X may hold missing elements denoted by NaN's which
%are ignored.
%
%The result is standardized, that is, corrected for the lower
%number of contributing terms.
%
%Check that for no column of X, all values are missing

% Copyright (C) 1995-2006  Rasmus Bro & Claus Andersson
% Copenhagen University, DK-1958 Frederiksberg, Denmark,
%Insert zeros for missing, correct afterwards
missidx = isnan(X);
i = find(missidx);
if ~isempty(i),
    X(i) = zeros(size(i));
end;

%Find the number of real(non-missing objects)
if min(size(X))==1,
   n_real=length(X)-sum(missidx);
   weight=length(X);
else
   n_real=size(X,1)-sum(missidx);
   weight=size(X,1);
end

i=find(n_real==0);
if isempty(i) %All values are real and can be corrected
   mm=weight*sum(X)./n_real;
else %There are columns with all missing, insert missing
   n_real(i)=1;
   mm=weight*sum(X)./n_real;
   mm(i)=i + NaN;
end
```
