## Supplementary File S1 for "Robust latent-variable interpretation of *in vivo* regression models by nested resampling": monreg.m

```
function [b,B,AllBs]=monreg(x);

%MONREG monotone regression
%
% See also:
% 'unimodal' 'monreg' 'fastnnls'
%
%
% MONTONE REGRESSION
% according to J. B. Kruskal 64
%
% b     = min|x-b| subject to monotonic increase
% B     = b, but condensed
% AllBs = All monotonic regressions, i.e. AllBs(1:i,i) is the 
%         monotonic regression of x(1:i)
%
% Reference
% Bro and Sidiropoulos, "Journal of Chemometrics", 1998, 12, 223-247. 
%
% [b,B,AllBs]=monreg(x);

% Copyright (C) 1995-2006  Rasmus Bro & Claus Andersson
% Copenhagen University, DK-1958 Frederiksberg, Denmark,
%

I=length(x);
if size(x,2)==2
   B=x;
else
   B=[x(:) ones(I,1)];
end

   AllBs=zeros(I,I);
   AllBs(1,1)=x(1);
   i=1;
   while i<size(B,1)
      if B(i,1)>B(min(I,i+1),1)
          summ=B(i,2)+B(i+1,2);
          B=[B(1:i-1,:);[(B(i,1)*B(i,2)+B(i+1,1)*B(i+1,2))/(summ) summ];B(i+2:size(B,1),:)];
          OK=1;
          while OK
             if B(i,1)<B(max(1,i-1),1)
                summ=B(i,2)+B(i-1,2);
                B=[B(1:i-2,:);[(B(i,1)*B(i,2)+B(i-1,1)*B(i-1,2))/(summ) summ];B(i+1:size(B,1),:)];
                i=max(1,i-1);
             else
                OK=0;
             end
          end
          bInterim=[];
          for i2=1:i
             bInterim=[bInterim;zeros(B(i2,2),1)+B(i2,1)];
          end
          No=sum(B(1:i,2));
          AllBs(1:No,No)=bInterim;
      else
          i=i+1;
          bInterim=[];
          for i2=1:i
             bInterim=[bInterim;zeros(B(i2,2),1)+B(i2,1)];
          end
          No=sum(B(1:i,2));
          AllBs(1:No,No)=bInterim;
      end
  end

  b=[];
  for i=1:size(B,1)
    b=[b;zeros(B(i,2),1)+B(i,1)];
  end
```
