## Supplementary File S1 for "Robust latent-variable interpretation of *in vivo* regression models by nested resampling": ncosine.m

```
function [MultPhi,Phis] = ncosine(factor1,factor2);

%NCOSINE multiple cosine/Tuckers congruence coefficient
%
% [MultPhi,Phis] = ncosine(factor1,factor2,DimX,Fac);
%
% ----------------------INPUT---------------------
%
% factor1   = cell array with loadings of one model
% factor2   = cell array with loadings of one (other) model
%     If factor1 and factor2 are identical then
%        the multiple cosine of a given solution is
%          estimated; otherwise the similarity of the
%          two different solutions is given
%
% ----------------------OUTPUT---------------------
%
% MultPhi   Is the multiple cosine of the model
% Phis      Is the cosine between components in
%          individual component matrices arranged
%          as [PhiA;PhiB ...]

% $ Version 2.00 $ May 2001 $ Changed to array notation $ RB $ Not compiled $
% $ Version 1.02 $ Date 28. July 1998 $ Not compiled $
% Copyright (C) 1995-2006  Rasmus Bro & Claus Andersson
% Copenhagen University, DK-1958 Frederiksberg, Denmark,


% Convert to old format
Fac = size(factor1,2);
for i = 1:length(factor1)
   DimX(i) = size(factor1{i},1);
end

ff = [];
for f=1:length(factor1)
 ff=[ff;factor1{f}(:)];
end
factor1 = ff;

ff = [];
for f=1:length(factor2)
 ff=[ff;factor2{f}(:)];
end
factor2 = ff;


if length(factor1)~=length(factor2)
  error(' factor1 and factor2 must hold components of same sizes in NCOSINE.M')
end
ord=length(DimX);
l_idx=0;
Fac=length(factor1)/sum(DimX);
for o=1:ord
  l_idx=[l_idx sum(DimX(1:o))*Fac];
end
L1=reshape(factor1(1:DimX(1)*Fac),DimX(1),Fac);
L2=reshape(factor2(1:DimX(1)*Fac),DimX(1),Fac);
for f=1:Fac
  L1(:,f)=L1(:,f)/norm(L1(:,f));
  L2(:,f)=L2(:,f)/norm(L2(:,f));
end
%GT correction
Phis=L1'*L2;
%Previously: Phis=L2'*L2;
%End GT correction
MultPhi=Phis;

for i=2:ord
  L1=reshape(factor1(l_idx(i)+1:l_idx(i+1)),DimX(i),Fac);
  L2=reshape(factor2(l_idx(i)+1:l_idx(i+1)),DimX(i),Fac);
  for f=1:Fac
    L1(:,f)=L1(:,f)/norm(L1(:,f));
    L2(:,f)=L2(:,f)/norm(L2(:,f));
  end
  phi=(L1'*L2);
  MultPhi=MultPhi.*phi;
  Phis=[Phis;phi];
end
```
