## Supplementary File S1 for "Robust latent-variable interpretation of *in vivo* regression models by nested resampling": ncrossdecomp.m

```
function XvalResult = ncrossdecomp(Method,X,FacMin,FacMax,Segments,Cent,Show);

%NCROSSDECOMP crossvalidation of PARAFAC/Tucker/PCA
%
% See also:
% 'ncrossreg' 
%
% This file performs cross-validation of decomposition models
% PARAFAC, PCA, and Tucker. The cross-validation is performed
% such that part of the data are set to missing, the model is 
% fitted to the remaining data, and the residuals between fitted
% and true left-out elements is calculated. This is performed
% 'Segments' times such that all elements are left out once.
% The segments are chosen by taking every 'Segments' element of
% X(:), i.e. from the vectorized array. If X is of size 5 x 7, 
% and three segemnts are chosen ('Segments' = 3), then in the 
% first of three models, the model is fitted to the matrix
% 
% |x 0 0 x 0 0 x|
% |0 x 0 0 x 0 0|
% |0 0 x 0 0 x 0|
% |x 0 0 x 0 0 x|
% |0 x 0 0 x 0 0|
% 
% where x's indicate missing elements. After fitting the residuals
% in the locations of missing values are calculated. After fitting
% all three models, all residuals have been calculated.
% 
% Note that the number of segments must be chosen such that no columns
% or rows contain only missing elements (the algorithm will check this).
% Using 'Segments' = 7, 9, or 13 will usually achieve that.
% 
% I/O
% XvalResult = ncrossdecomp(Method,X,FacMin,FacMax,Segments,Cent,Show);
% 
% INPUT
% Method   : 'parafac', 'tucker', 'pca', or 'nipals'
%            For PCA the least squares model is calculated.
%            Thus, offsets and parameters are calculated in
%            a least squares sense unlike the method NIPALS,
%            which calculates the PCA model using an ad hoc 
%            approach for handling missing data (as in 
%            standard chemometric software).
% X        : Multi-way array of data 
% FacMin   : Lowest number of factors to use
% FacMax   : Highest number of factors (note that for Tucker only models 
%            with the same number of components in each mode are
%            calculated currently
% Segments : The number of segments to use. Try many!
% Cent     : If set of one, the data are centered across samples, 
%            i.e. ordinary centering. Note, however, that the centering
%            is not performed in a least squares sense but as preprocessing.
%            This is not optimal because the data have missing data because
%            of the way the elements are left out. This can give 
%            significantly lower fit than reasonable if you have few samples
%            or use few segments. Alternatively, you can center the data 
%            beforehand and perform cross-validation on the centered data
% Show     : If set to 0, no plot is given
%
% OUTPUT
% Structure XvalResult holding:
%           Fit: The fitted percentage of variation explaind (as a 
%                function of component number)
%          Xval: The cross-validated percentage of variation explaind
%                (as a function of component number)
%   FittedModel: The fitted model (as a function of component number)
%     XvalModel: The cross-validated model (as a function of component number)
% 
%  To visualize the output type "tucktest(XvalResult);"

% $ Version 1.0301 $ Date 28. June 1999 $ Not compiled $
% $ Version 2.00 $ May 2001 $ Changed to array notation $ RB $ Not compiled $
% $ Version 2.01 $ Mar 2002 $ Fixed error in segmentation check $ RB $ Not compiled $


% Copyright (C) 1995-2006  Rasmus Bro & Claus Andersson
% Copenhagen University, DK-1958 Frederiksberg, Denmark,
%

% uses NANSUM,

DimX = size(X);
ord = length(DimX);
X = reshape(X,DimX(1),prod(DimX(2:end)));

[I,J] = size(X);

if exist('Show')~=1
   Show = 1;
end


if strcmp(lower(Method(1:3)),'tuc')
   if length(FacMin)==1
      FacMin = ones(1,ord)*FacMin;
   elseif length(FacMin)~=ord
      error('Error in FacMin: When fitting Tucker models, the number of factors should be given for each mode')
   end
   if length(FacMax)==1
      FacMax = ones(1,ord)*FacMax;
   elseif length(FacMax)~=ord
      error('Error in FacMax: When fitting Tucker models, the number of factors should be given for each mode')
   end
end 


% Check if the selected segmentation works (does not produce rows/columns of only missing)
out = ones(I,J);
out(1:Segments:end)=NaN;
out2 = ones(size(X));
out(find(isnan(out2))) = NaN;
if any(sum(isnan(out))==I)
   error(' The chosen segmentation leads to columns of only missing elements')
elseif any(sum(isnan(out'))==J)
   error(' The chosen segmentation leads to rows of only missing elements')
end


if ~strcmp(lower(Method(1:3)),'tuc')
   
   XvalResult.Fit = zeros(FacMax,2)*NaN;
   XvalResult.Xval = zeros(FacMax,2)*NaN;
   for f = 1:FacMin-1
      XvalResult.XvalModel{f} =  'Not fitted';
      XvalResult.FittedModel{f} = 'Not fitted';
   end
   
   for f = FacMin:FacMax
      
      % Fitted model
      disp([' Total model - Comp. ',num2str(f),'/',num2str(FacMax)])
      [M,Mean,Param] = decomp(Method,X,DimX,f,1,Segments,Cent,I,J);
      Model = M + ones(I,1)*Mean';
      id    = find(~isnan(X));
      OffsetCorrectedData = X - ones(I,1)*Mean';
      XvalResult.Fit(f,:) = [100*(1 - sum( (X(id) - Model(id)).^2)/sum(OffsetCorrectedData(id).^2)) f];
      XvalResult.FittedModel{f} = Model;
      
      
      % Xvalidated Model of data
      ModelXval = zeros(I,J)*NaN;
      for s = 1:Segments
         disp([' Segment ',num2str(s),'/',num2str(Segments),' - Comp. ',num2str(f),'/',num2str(FacMax)])
         Xnow = X;
         Xnow(s:Segments:end) = NaN;
         [M,Mean] = decomp(Method,Xnow,DimX,f,s,Segments,Cent,I,J,Param);
         model = M + ones(I,1)*Mean';
         ModelXval(s:Segments:end) = model(s:Segments:end);
      end
      
      XvalResult.Xval(f,:) = [100*(1 - sum( (X(id) - ModelXval(id)).^2)/sum(OffsetCorrectedData(id).^2)) f];
      XvalResult.XvalModel{f} =  ModelXval;
      XvalResult.Factors(f)=f;

   end
   
else % Do Tucker model
   
   % Find all 
   PossibleNumber = [min(FacMin):max(FacMax)]'*ones(1,ord);
   possibleCombs = unique(nchoosek(PossibleNumber(:),ord),'rows');
   %remove useless
   f2 = [];
   for f1 = 1:size(possibleCombs,1)
      if (prod(possibleCombs(f1,:))/max(possibleCombs(f1,:)))<max(possibleCombs(f1,:)) % Check that the largest mode is larger than the product of the other
         f2 = [f2;f1];
      elseif any(possibleCombs(f1,:)>FacMax)  % Chk the model is desired,
         f2 = [f2;f1];
      end
   end
   possibleCombs(f2,:)=[];
   [f1,f2]=sort(sum(possibleCombs'));
   possibleCombs = [possibleCombs(f2,:) f1'];
   
   
   XvalResult.Fit = zeros(size(possibleCombs,1),ord+1)*NaN;
   XvalResult.Xval = zeros(size(possibleCombs,1),ord+1)*NaN;
   for f = 1:size(possibleCombs,1)
      XvalResult.XvalModel{f} =  'Not fitted';
      XvalResult.FittedModel{f} = 'Not fitted';
   end
   
   
   
   for f1 = 1:size(possibleCombs,1)
      
      % Fitted model
      disp([' Total model - Comp. ',num2str(possibleCombs(f1,1:end-1)),'/',num2str(FacMax)])
      [M,Mean,Param] = decomp(Method,X,DimX,possibleCombs(f1,1:end-1),1,Segments,Cent,I,J);
      Model = M + ones(I,1)*Mean';
      id    = find(~isnan(X));
      OffsetCorrectedData = X - ones(I,1)*Mean';
      XvalResult.Fit(f1,:) = [100*(1 - sum( (X(id) - Model(id)).^2)/sum(OffsetCorrectedData(id).^2)) possibleCombs(f1,1:end-1)];
      XvalResult.FittedModel{f1} = Model;
      
      
      % Xvalidated Model of data
      ModelXval = zeros(I,J)*NaN;
      for s = 1:Segments
         disp([' Segment ',num2str(s),'/',num2str(Segments),' - Comp. ',num2str(possibleCombs(f1,1:end-1)),'/',num2str(FacMax)])
         Xnow = X;
         Xnow(s:Segments:end) = NaN;
         [M,Mean] = decomp(Method,Xnow,DimX,possibleCombs(f1,1:end-1),s,Segments,Cent,I,J,Param);
         model = M + ones(I,1)*Mean';
         ModelXval(s:Segments:end) = model(s:Segments:end);
      end
      
      XvalResult.Xval(f1,:) = [100*(1 - sum( (X(id) - ModelXval(id)).^2)/sum(OffsetCorrectedData(id).^2)) possibleCombs(f1,1:end-1)];
      XvalResult.XvalModel{f1} =  ModelXval;
      XvalResult.Factors{f1}=possibleCombs(f1,1:end-1);
   end
   
end


if Show&FacMin-FacMax~=0
   if Method(1:3) == 'pca'
      Nam = 'PCA';
   elseif Method(1:3) == 'tuc'
      Nam = 'Tucker';
   elseif Method(1:3) == 'par'
      Nam = 'PARAFAC';
   elseif Method(1:3) == 'nip'
      Nam = 'NIPALS';
   end
   
   figure      
   save jjj
   if lower(Method(1:3))~='tuc'
      bar(FacMin:FacMax,[XvalResult.Fit(FacMin:FacMax,1) XvalResult.Xval(FacMin:FacMax,1)],.76,'grouped')
   else
      % extract the ones with lowest Xval fit (for each # total comp) for plotting
      fx = [];
      f5 =[];
      for f1 = 1:max(possibleCombs(:,end))
         f2 = find(possibleCombs(:,end)==f1);
         if length(f2)
            [f3,f4] = max(XvalResult.Xval(f2));
            f5 = [f5,f2(f4)];
         end
      end
      fx = [possibleCombs(f5,end) XvalResult.Fit(f5,1) XvalResult.Xval(f5,1)];
      bar(fx(:,1),fx(:,2:3),.76,'grouped')
      for f1 = 1:size(fx,1)
         f6=text(fx(f1,1),95,['[',num2str(possibleCombs(f5(f1),1:end-1)),']']);
         set(f6,'Rotation',270)
      end
   end
   
   g=get(gca,'YLim');
   set(gca,'YLim',[max(-20,g(1)) 100])
   legend('Fitted','Xvalidated',0)
   titl = ['Xvalidation results (',Nam,')'];
   if Cent
      titl = [titl ,' - centering'];
   else
      titl = [titl ,' - no centering'];
   end
   title(titl,'FontWeight','Bold')
   xlabel('Total number of components')
   ylabel('Percent variance explained')
end   


function [M,Mean,parameters] = decomp(Method,X,DimX,f,s,Segments,Cent,I,J,parameters);

Conv = 0;
it = 0;
maxit = 500;
% Initialize
if Cent
   Mean = nanmean(X)';
else
   Mean = zeros(J,1);
end

if lower(Method(1:3)) == 'par'
   Xc = reshape(X- ones(I,1)*Mean',DimX);
   if exist('parameters')==1
      fact = parafac(Xc,f,[1e-5 10 0 0 NaN maxit],[],parameters.fact);
   else
      fact = parafac(Xc,f,[1e-5 10 0 0 NaN maxit]);
   end
   M = reshape(nmodel(fact),DimX(1),prod(DimX(2:end)));
   parameters.fact=fact;
elseif lower(Method(1:3)) == 'tuc'
   Xc = reshape(X- ones(I,1)*Mean',DimX);
   if exist('parameters')==1
      [fact,G] = tucker(Xc,f,[1e-2 0 0 0 NaN maxit],[],[],parameters.fact,parameters.G);
   else
      [fact,G] = tucker(Xc,f,[1e-2 0 0 0 NaN maxit]);
   end
   parameters.fact=fact;
   parameters.G=G;
   M = reshape(nmodel(fact,G),DimX(1),prod(DimX(2:end)))      ;
elseif lower(Method) == 'nip'
   Xc = reshape(X- ones(I,1)*Mean',DimX(1),prod(DimX(2:end)));
   [t,p] = pcanipals(X- ones(I,1)*Mean',f,0);
   parameters.t=t;
   parameters.p=p;
   M = t*p';
elseif lower(Method) == 'pca'
   Xc = reshape(X- ones(I,1)*Mean',DimX(1),prod(DimX(2:end)));
   [t,p] = pcals(X- ones(I,1)*Mean',f,0);
   parameters.t=t;
   parameters.p=p;
   M = t*p';
else
   error(' Name of method not recognized') 
end
Fit = X - M - ones(I,1)*Mean';
Fit = sum(Fit(find(~isnan(X))).^2);

% Iterate
while ~Conv
   it     = it+1;
   FitOld = Fit;
   
   % Fit multilinear part
   Xcent = X - ones(I,1)*Mean';
   
   if Method(1:3) == 'par'
      fact = parafac(reshape(Xcent,DimX),f,[1e-2 0 0 0 NaN maxit],[],fact);
      M = reshape(nmodel(fact),DimX(1),prod(DimX(2:end)));
   elseif Method(1:3) == 'tuc'
      [fact,G] = tucker(reshape(Xcent,DimX),f,[1e-2 0 0 0 NaN maxit],[0 0 0],zeros(size(G)),fact,G);
      M = reshape(nmodel(fact,G),DimX(1),prod(DimX(2:end)));
   elseif Method == 'pca'
      [t,p] = pcals(Xcent,f,0,t,p,0);
      M = t*p';
   elseif Method == 'nip'
      [t,p] = pcanipals(Xcent,f,0);
      M = t*p';
   end
   
   % Find offsets
   if Cent
      x = X;
      mm=M+ones(I,1)*Mean';
      x(find(isnan(X)))=mm(find(isnan(X)));
      Mean = mean(x)';
   end
   
   
   %Find fit
   Fit = X - M - ones(I,1)*Mean';
   Fit = sum(Fit(find(~isnan(X))).^2);
   if abs(Fit-FitOld)/FitOld<1e-8 | it > 1500
      Conv = 1;
   end
   
end
disp([' Fit ',num2str(Fit),' using ',num2str(it),' it.'])


function [t,p] = pcals(X,F,cent,t,p,show);

%  LEAST SQUARES PCA WITH MISSING ELEMENTS
%  20-6-1999
% 
%  Calculates a least squares PCA model. Missing elements 
%  are denoted NaN. The solution is NOT nested, so one has
%  to calculate a new model for each number of components.


ShowMeFitEvery = 20;
MaxIterations  = 5;
[I,J]=size(X);
Xorig      = X;
Miss       = find(isnan(X));
NotMiss    = find(~isnan(X));
m          = t*p';
X(Miss)    = m(Miss);
ssX    = sum(X(NotMiss).^2);

Fit    = 3;
OldFit = 6;
it     = 0;

while abs(Fit-OldFit)/OldFit>1e-3 & it < MaxIterations;
   it      = it +1;
   OldFit  = Fit;
   
   [t,s,p] = svds(X,F);
   t       = t*s;
   
   Model   = t*p';
   X(Miss) = Model(Miss);
   Fit     = sum(sum( (Xorig(NotMiss) - Model(NotMiss)).^2));
   
   if ~rem(it,ShowMeFitEvery)&show
      disp(['    Fit after ',num2str(it),' it. :',num2str(RelFit),'%'])
   end
   
end


function [t,p,Mean] = pcanipals(X,F,cent);

% NIPALS-PCA WITH MISSING ELEMENTS
% cent: One if centering is to be included, else zero


[I,J]=size(X);
rand('state',sum(100*clock))

Xorig      = X;
Miss       = isnan(X);
NotMiss    = ~isnan(X);
ssX    = sum(X(find(NotMiss)).^2);
Mean   = zeros(1,J);
if cent
   Mean    = nanmean(X);
end
X      = X - ones(I,1)*Mean;

t=[];
p=[];

for f=1:F
   Fit    = 3;
   OldFit = 6;
   it     = 0;
   T      = rand(I,1);
   P      = rand(J,1);
   Fit    = 2;
   FitOld = 3;
   
   while abs(Fit-FitOld)/FitOld>1e-7 & it < 100;
      FitOld  = Fit;
      it      = it +1;
      
      for j = 1:J
        try
         id=find(NotMiss(:,j));
         if length(id)==0
            id,end

         P(j) = T(id)'*X(id,j)/(T(id)'*T(id));
       catch
         P(j) = 0;
       end
      end
      P = P/norm(P);
      
      for i = 1:I
         id=find(NotMiss(i,:));
         T(i) = P(id)'*X(i,id)'/(P(id)'*P(id));
      end
      
      Fit = X-T*P';
      Fit = sum(Fit(find(NotMiss)).^2);
   end
   t = [t T];
   p = [p P];
   X = X - T*P';
   
end

function Xc = nanmean(X)

if isempty(X)
   Xc = NaN;
   return
end

i = isnan(X);
j = find(i);
i = sum(i);
X(j) = 0;
Num = size(X,1)-i;
Xc = sum(X);
i = find(Num);
Xc(i) = Xc(i)./Num(i);
Xc(find(~Num))=NaN;
```
