## Supplementary File S1 for "Robust latent-variable interpretation of *in vivo* regression models by nested resampling": ncrossreg2.m

```
function [XValResult,Model]=ncrossreg(X,y,MaxFac,Centering,SegmentsID);
%NCROSSREG cross-validation of regression model
%
% See also:
% 'ncrossdecomp'
%
% CROSS-VALIDATION OF BI- & MULTILINEAR REGRESSION MODELS
% Performs cross-validation of
% - NPLS  Input multi-way array X
% - PLS   Input two-way X
%
% The data are by default centered across the first mode, but no scaling
% is applied (this must be done beforehand)
%
% I/O
% [XValResult,Model]=ncrossreg(X,y,MaxFac,Centering);
%
% INPUT
% X         : X array
% y         : y array
% MaxFac    : Maximal number of factors (from one to MaxFac factors will be investigated)
%
% OPTIONAL INPUT
% Centering : If not zero, centering is performed on every segment
% SegmentsID: Optional binary matrix. Rows as rows in X and i'th column defines i'th segment
%             Rows in i'th column set to one are left out at
%
% OUTPUT
% XValResult
%  Structured array with the following elements
%  ypred    : Cross-validated predictions
%  ssX_Xval : Cross-validated sum of squares of residuals in X (f'th element for f-component model)
%  ssX_Fit  : Fitted sum of squares of residuals in X (f'th element for f-component model)
%  ssY_Xval : Cross-validated sum of squares of residuals in Y (f'th element for f-component model)
%  ssY_Fit  : Fitted sum of squares of residuals in Y (f'th element for f-component model)
%  Percent  : Structured array holding Xexp and Yexp, each with fitted and X-validated % Variance captured.
%  PRESS    : Predicted REsidual Sum of Squares in Y
%  RMSEP    : Root Mean Square Error of Prediction (cross-validation)
%
% Model
%  Structured array holding the NPLS model


% $ Version 1.0301 $ Date 26. June 1999
% $ Version 1.0302 $ Date 1. January 2000
% $ Version 2.00 $ May 2001 $ Changed to array notation $ RB $ Not compiled $
% $ Version 2.01 $ jan 2003 $ Added option for skipping centering and added percentages in output $ RB $ Not compiled $
% $ Version 2.02 $ Sep 2003 $ Fixed bug in non-center option $ RB $ Not compiled $

% Copyright (C) 1995-2006  Rasmus Bro & Claus Andersson
% Copenhagen University, DK-1958 Frederiksberg, Denmark,
%

I = size(X,1);

DimX = size(X);
DimY = size(y);
X = reshape(X,DimX(1),prod(DimX(2:end)));
y = reshape(y,DimY(1),prod(DimY(2:end)));

Ypred     = zeros([MaxFac DimY]);
Ex        = zeros([MaxFac DimX]);
Ey        = zeros([MaxFac DimY]);

if nargin<4
  Centering = 1;
elseif isempty(Centering)
  Centering = 1;
end

%%%%%%%%%%%%%%%%%
%%MAKE SEGMENTS%%
%%%%%%%%%%%%%%%%%
if exist('SegmentsID')~=1
  SegmentsID = MakeSegments(I);
end

%%%%%%%%%%%%%%%%%%%
%%MAKE SUB-MODELS%%
%%%%%%%%%%%%%%%%%%%

for i=1:size(SegmentsID,2)
  In = find(~SegmentsID(:,i));
  Out = find(SegmentsID(:,i));
  % Offsets
  if Centering
    Mx = nanmean(X(In,:));
    My = nanmean(y(In,:));
  else
    Mx = zeros(1,prod(DimX(2:end)));
    My = zeros(1,prod(DimY(2:end)));
  end
  %Centered data
  Xc = X(In,:)-repmat(Mx,length(In),1);
  yc = y(In,:)-repmat(My,length(In),1);

  %   %Centered data
  %  Xc = X(In,:)-ones(length(In),1)*Mx;
  %  yc = y(In,:)-ones(length(In),1)*My;

  % Calculate model
  DimXc = DimX;DimXc(1)=size(Xc,1);
  Xc = reshape(Xc,DimXc);
  DimYc = DimY;DimYc(1)=size(yc,1);
  yc = reshape(yc,DimYc);
  [Xfactors,Yfactors,Core,B] = npls(Xc,yc,MaxFac,NaN);


  %Predict left-out samples
  for f=1:MaxFac
    Xc = X(Out,:)-ones(length(Out),1)*Mx;
    DimXc = DimX;
    DimXc(1)=size(Xc,1);
    Xc = reshape(Xc,DimXc);
    [ypr,T,ssx,Xres] = npred(Xc,f,Xfactors,Yfactors,Core,B,NaN);
    Ex(f,Out,:)    = reshape(Xres,DimXc(1),prod(DimXc(2:end)));
    Ypred(f,Out,:) = ypr+ones(length(Out),1)*My;
    Ypredf = squeeze(Ypred(f,:,:));
    if size(y,2) == 1
      YpredfOut=Ypredf(Out);
    else
      YpredfOut=Ypredf(Out,:);
    end
    %size(Ey(f,Out,:)),size(y(Out,:)),size(YpredfOut)
    if size(y,2)==1
      Ey(f,Out,:)    = squeeze(y(Out,:))'-squeeze(YpredfOut);
    else
      Ey(f,Out,:)    = squeeze(y(Out,:))-squeeze(YpredfOut);
    end
  end
end

if Centering
  Mx = nanmean(X(In,:));
  My = nanmean(y(In,:));
else
  Mx = zeros(1,prod(DimX(2:end)));
  My = zeros(1,prod(DimY(2:end)));
end
%Centered data
Xc = X-repmat(Mx,size(X,1),1);
yc = y-repmat(My,size(y,1),1);

%%Centered data
%Xc = X-ones(I,1)*Mx;
%yc = y-ones(I,1)*My;
[Xfactors,Yfactors,Core,B,ypred,ssx,ssy] = npls(reshape(Xc,DimX),reshape(yc,DimY),MaxFac,NaN);
Model.Xfactors = Xfactors;
Model.Yfactors = Yfactors;
Model.Core     = Core;
Model.B        = B;
Model.Yfitted  = ypred;
Model.MeanX    = Mx;
Model.MeanY    = My;

sseX_fit  = ssx(2:end,1);
sseY_fit  = ssy(2:end,1);
for f=1:MaxFac
  id=find(~isnan(Ex(f,:)));sseX_xval(f) = sum(Ex(f,id).^2);
  id=find(~isnan(Ey(f,:)));sseY_xval(f) = sum(Ey(f,id).^2);
  PRESS(f) = sum(Ey(f,id).^2);
end
RMSEP = sqrt(PRESS/I);

Xval = [sseX_fit sseX_xval'];
Yval = [sseY_fit sseY_xval'];
Xval = 100*(1-Xval/sum(Xc(find(~isnan(X))).^2));
Yval = 100*(1-Yval/sum(yc(find(~isnan(y))).^2));

XValResult.ypred       = Ypred;
XValResult.ssX_Xval    = sseX_xval;
XValResult.ssX_Fit     = sseX_fit';
XValResult.ssY_Xval    = sseY_xval;
XValResult.ssY_Fit     = sseY_fit';
XValResult.Percent.Xexp = Xval;
XValResult.Percent.Yexp = Yval;
XValResult.PRESS       = PRESS;
XValResult.RMSEP       = RMSEP;
XValResult.DefSegments = sparse(SegmentsID);

disp('  ')
disp('       Percent Variance Captured by N-PLS Model   ')
disp('  ')
disp('           -----X-Block-----    -----Y-Block-----')
disp('   LV #    Fitted     Xval      Fitted     Xval       RMSEP')
disp('   ----    -------   -------    -------   -------   ---------')
format = '   %3.0f     %6.2f    %6.2f     %6.2f    %6.2f    %6.4f';
for f = 1:MaxFac
  tab = sprintf(format,[f Xval(f,:) Yval(f,:) RMSEP(f)]);
  disp(tab)
end
disp('  ')


function SegmentsID = MakeSegments(I);

% XvalMeth=questdlg('Which type of validation do you want to perform (ENTER => full Xval)?','Choose validation','Full X-validation','Segmented','Prespecified','Full X-validation');
XvalMeth = 'Full X-validation';

switch XvalMeth
  case 'Full X-validation'
    SegmentsID = speye(I);

  case 'Segmented'
    prompt={'Enter the number of segments:'};
    eval(['def={''',num2str(min(I,max(3,round(I/7)))),'''};'])
    dlgTitle='Number of segments';
    lineNo=1;
    answer=inputdlg(prompt,dlgTitle,lineNo,def);
    NumbSegm=eval(answer{1});

    % Make sure the number of segments is OK
    while NumbSegm<2|NumbSegm>I
      prompt={'INCONSISTENT NUMBER CHOSEN (must be > 1 and <= samples)'};
      eval(['def={''',num2str(min(I,max(3,round(I/7)))),'''};'])
      dlgTitle='Number of segments';
      lineNo=1;
      answer=inputdlg(prompt,dlgTitle,lineNo,def);
      NumbSegm=eval(answer{1})
      NumbSegm<2|NumbSegm>I
    end

    XvalSegm=questdlg('How should segments be chosen?','Choose segmentation','111222333...','123123123...','Random','123123123...');
    switch XvalSegm

      case '111222333...'
        SegmentsID = sparse(I,NumbSegm);
        NumbInEachSegm = floor(I/NumbSegm);
        Additional = I-NumbInEachSegm*NumbSegm;
        currentsample = 1;
        for i=1:NumbSegm
          if i <=Additional
            add = NumbInEachSegm+1;
          elseif i<NumbSegm
            add = NumbInEachSegm;
          else
            add = I-currentsample+1;
          end
          SegmentsID(currentsample:currentsample+add-1,i)=1;
          currentsample = currentsample + add;
        end
      case '123123123...'
        SegmentsID = sparse(I,NumbSegm);
        NumbInEachSegm = floor(I/NumbSegm);
        for i=1:NumbSegm
          SegmentsID(i:NumbSegm:end,i)=1;
        end
      case 'Random'
        % Make nonrandom and then randomize order
        SegmentsID = sparse(I,NumbSegm);
        NumbInEachSegm = floor(I/NumbSegm);
        for i=1:NumbSegm
          SegmentsID(i:NumbSegm:end,i)=1;
        end
        rand('state',sum(100*clock)) %Randomize randomizer
        [a,b] = sort(rand(I,1))
        SegmentsID = SegmentsID(b,:);
    end

  case 'Prespecified'
    prompt={'Enter the name of the file defining the subsets'};
    def={'SegmentsID'};
    dlgTitle='Import definition';
    lineNo=1;
    answer=inputdlg(prompt,dlgTitle,lineNo,def);
    SegmentsID=eval(answer{1});

end % switch

function y = nanmean(x)
%NANMEAN Average or mean ignoring NaNs.
%   NANMEAN(X) returns the average treating NaNs as missing values.
%   For vectors, NANMEAN(X) is the mean value of the non-NaN
%   elements in X.  For matrices, NANMEAN(X) is a row vector
%   containing the mean value of each column, ignoring NaNs.
%
%   See also NANMEDIAN, NANSTD, NANMIN, NANMAX, NANSUM.

%   Copyright (c) 1993-98 by The MathWorks, Inc.
%   $Revision: 2.8 $  $Date: 1997/11/29 01:45:53 $

if isempty(x) % Check for empty input.
  y = NaN;
  return
end

% Replace NaNs with zeros.
nans = isnan(x);
i = find(nans);
x(i) = zeros(size(i));

if min(size(x))==1,
  count = length(x)-sum(nans);
else
  count = size(x,1)-sum(nans);
end

% Protect against a column of all NaNs
i = find(count==0);
count(i) = ones(size(i));
y = sum(x)./count;
y(i) = i + NaN;
```
