## Supplementary File S1 for "Robust latent-variable interpretation of *in vivo* regression models by nested resampling": neye.m

```
function G=neye(Fac);
% NEYE  Produces a super-diagonal array
%
%function G=neye(Fac);
%
% $ Version 2.00 $ May 2001 $ Changed to array notation $ RB $ Not compiled $
% $ Version 1.00 $ Date 5. Aug. 1998 $ Not compiled $
%
% This algorithm requires access to:
% 'getindxn'
%
% See also:
% 'parafac' 'maxvar3' 'maxdia3'
%
% ---------------------------------------------------------
%             Produces a super-diagonal array
% ---------------------------------------------------------
%	
% G=neye(Fac);
%
% Fac      : A row-vector describing the number of factors
%            in each of the N modes. Fac must be a 1-by-N vector. 
%            Ex. [3 3 3] or [2 2 2 2]

% Copyright (C) 1995-2006  Rasmus Bro & Claus Andersson
% Copenhagen University, DK-1958 Frederiksberg, Denmark,
%

N=size(Fac,2);
if N==1,
   fprintf('Specify ''Fac'' as e vector to define the order of the core, e.g.,.\n')
   fprintf('G=eyecore([2 2 2 2])\n')
end;

G=zeros(Fac(1),prod(Fac(2:N)));

for i=1:Fac(1),
   [gi,gj]=getindxn(Fac,ones(1,N)*i);
   G(gi,gj)=1;
end;

G = reshape(G,Fac);
```
