## Supplementary File S1 for "Robust latent-variable interpretation of *in vivo* regression models by nested resampling": nident.m

```
function id = nident(J,order,mag);

% NIDENT make 'identity' multi-way array
%
% Make 'identity' array of order given by "order" and dimension J.
% id = nident(J,order);
% if extra input vector, mag, is given the j'th superdiagonal will be
% equal to mag(j)

% Copyright (C) 1995-2006  Rasmus Bro & Claus Andersson
% Copenhagen University, DK-1958 Frederiksberg, Denmark,
%

% $ Version 2.00 $ May 2001 $ Changed to array notation $ RB $ Not compiled $
% $ Version 1.02 $ Date 28. July 1998 $ Not compiled $

if nargin<3
  mag=ones(J,1);
end

id=zeros(J^order,1);
for f=1:J
  idd=f;
  for i=2:order
    idd=idd+J^(i-1)*(f-1);
  end
  id(idd)=mag(f);
end
id  = reshape(id,ones(1,order)*J);
```
