## Supplementary File S1 for "Robust latent-variable interpretation of *in vivo* regression models by nested resampling": nmodel.m

```
function [Xm]=nmodel(Factors,G,Om);

%NMODEL make model of data from loadings
%
% function [Xm]=nmodel(Factors,G,Om);
%
% This algorithm requires access to:
% 'neye.m'
%
%
% [Xm]=nmodel(Factors,G,Om);
%
% Factors  : The factors in a cell array. Use any factors from 
%            any model. 
% G        : The core array. If 'G' is not defined it is assumed
%            that a PARAFAC model is being established.
%            Use G = [] in the PARAFAC case.
% Om       : Oblique mode.
%            'Om'=[] or 'Om'=0, means that orthogonal
%                   projections are requsted. (default)
%            'Om'=1 means that the factors are oblique.  
%            'Om'=2 means that the ortho/oblique is solved automatically.  
%                   This takes a little additional time.
% Xm       : The model of X.
%
% Using the factors as they are (and the core, if defined) the general N-way model
% is calculated. 


% $ Version 2.00 $ May 2001 $ Changed to array notation $ RB $ Not compiled $
% $ Version 1.02 $ Date 17. Apr 1999 $ Not compiled $
% Copyright (C) 1995-2006  Rasmus Bro & Claus Andersson
% Copenhagen University, DK-1958 Frederiksberg, Denmark,
%

for i = 1:length(Factors);
   DimX(i)=size(Factors{i},1);
end
i = find(DimX==0);
for j = 1:length(i)
   DimX(i(j)) = size(G,i(j));
end


if nargin<2, %Must be PARAFAC
   Fac=size(Factors{1},2);
   G=[];
else
   for f = 1:length(Factors)
      if isempty(Factors{f})
         Fac(f) = -1;
      else
         Fac(f) = size(Factors{f},2);
      end;
   end
end

if ~exist('Om')
    Om=[];
end;

if isempty(Om)
    Om=0;
end;

if size(Fac,2)==1,
    Fac=Fac(1)*ones(1,size(DimX,2));
end;
N=size(Fac,2);

if size(DimX,2)>size(Fac,2),
    Fac=Fac*ones(1,size(DimX,2));
end;  
N=size(Fac,2);

Fac_orig=Fac;
i=find(Fac==-1);
if ~isempty(i)
    Fac(i)=zeros(1,length(i));
    Fac_ones(i)=ones(1,length(i));
end;
DimG=Fac;
i=find(DimG==0);
DimG(i)=DimX(i);

if isempty(G),
   G=neye(DimG);
end;   
G = reshape(G,size(G,1),prod(size(G))/size(G,1));

% reshape factors to old format
ff = [];
for f=1:length(Factors)
 ff=[ff;Factors{f}(:)];
end
Factors = ff;


if DimG(1)~=size(G,1) | prod(DimG(2:N))~=size(G,2),

    help nmodel

    fprintf('nmodel.m   : ERROR IN INPUT ARGUMENTS.\n');
    fprintf('             Dimension mismatch between ''Fac'' and ''G''.\n\n');
    fprintf('Check this : The dimensions of ''G'' must correspond to the dimensions of ''Fac''.\n');
    fprintf('             If a PARAFAC model is established, use ''[]'' for G.\n\n');
    fprintf('             Try to reproduce the error and request help at\n');
    return;
end;

if sum(DimX.*Fac) ~= length(Factors),
    help nmodel
    fprintf('nmodel.m   : ERROR IN INPUT ARGUMENTS.\n');
    fprintf('             Dimension mismatch between the number of elements in ''Factors'' and ''DimX'' and ''Fac''.\n\n');
    fprintf('Check this : The dimensions of ''Factors'' must correspond to the dimensions of ''DimX'' and ''Fac''.\n');
    fprintf('             You may be using results from different models, or\n');
    fprintf('             You may have changed one or more elements in ''Fac'' or ''DimX'' after ''Factors'' have been calculated.\n\n');
    fprintf('             Read the information above for information on arguments.\n');
    return;
end;

FIdx0=cumsum([1 DimX(1:N-1).*Fac(1:N-1)]);
FIdx1=cumsum([DimX.*Fac]);

if Om==0,
    Orthomode=1;
end;

if Om==1,
    Orthomode=0;
end;

if Om==2,
    Orthomode=1;
    for c=1:N,
        if Fac_orig(c)~=-1,
            A=reshape(Factors(FIdx0(c):FIdx1(c)),DimX(c),Fac(c));
            AA=A'*A;
            ssAA=sum(sum(AA.^2));
            ssdiagAA=sum(sum(diag(AA).^2));
            if abs(ssAA-ssdiagAA) > 100*eps;
                Orthomode=0;
            end;
        end;
    end;
end;

if Orthomode==0,
    Zmi=prod(abs(Fac_orig(2:N)));
    Zmj=prod(DimX(2:N));
    Zm=zeros(Zmi,Zmj);
    DimXprodc0 = 1;
    Facprodc0 = 1;
    Zm(1:Facprodc0,1:DimXprodc0)=ones(Facprodc0,DimXprodc0);
    for c=2:N,
        if Fac_orig(c)~=-1,
            A=reshape(Factors(FIdx0(c):FIdx1(c)),DimX(c),Fac(c));
            DimXprodc1 = DimXprodc0*DimX(c);
            Facprodc1 = Facprodc0*Fac(c);
            Zm(1:Facprodc1,1:DimXprodc1)=ckron(A',Zm(1:Facprodc0,1:DimXprodc0));
            DimXprodc0 = DimXprodc1;
            Facprodc0 = Facprodc1;
        end;
    end;
    if Fac_orig(1)~=-1,
        A=reshape(Factors(FIdx0(1):FIdx1(1)),DimX(1),Fac(1));
        Xm=A*G*Zm;
    else 
        Xm=G*Zm;
    end;
elseif Orthomode==1,
    CurDimX=DimG;
    Xm=G;
    newi=CurDimX(2);
    newj=prod(CurDimX)/CurDimX(2);
    Xm=reshape(Xm',newi,newj);
    for c=2:N,
        if Fac_orig(c)~=-1,
            A=reshape(Factors(FIdx0(c):FIdx1(c)),DimX(c),Fac(c));
            Xm=A*Xm;
            CurDimX(c)=DimX(c);
        else
            CurDimX(c)=DimX(c);
        end;
        if c~=N,
            newi=CurDimX(c+1);
            newj=prod(CurDimX)/CurDimX(c+1);
        else,
				newi=CurDimX(1);
            newj=prod(CurDimX)/CurDimX(1);
        end;
        Xm=reshape(Xm',newi,newj);
    end;
    if Fac_orig(1)~=-1,
        A=reshape(Factors(FIdx0(1):FIdx1(1)),DimX(1),Fac(1));
        Xm=A*Xm;
    end;
end;    

Xm = reshape(Xm,DimX);
```
