## Supplementary File S1 for "Robust latent-variable interpretation of *in vivo* regression models by nested resampling": nonneg.m

```
function [F,SS]=nonneg(X,M,F,Options);
%NONNEG1 alternative to NNLS
%
%function [F,SS]=nonneg(X,M,F,Options);
%
% 'nonneg.m'
% $ Version 0.01 $ Date 6. Aug. 1997 $ Not compiled $
%
% This algorithm requires access to:
% ''
%
% ---------------------------------------------------------
%             Fast! Non-negativity Regression
% ---------------------------------------------------------
%	
% [F,SS]=nonneg(X,M,F,Options);
% [F,SS]=nonneg(X,M);
%
% X        : Matrix of regressors.
% M        : Matrix of regressands.
% F        : The non-negativity constrained solution matrix.
% Options  : Type 'help Options' at the prompt.
%
% Given X and M this algorithm solves for the optimal 
% F in a least squares sense, using that
%      X = F*M 
% in the problem
%      min ||X-F*M||, s.t. F>=0, for given X and M.
%
% This version does not accept missing values.

% Copyright (C) 1995-2006  Rasmus Bro & Claus Andersson
% Copenhagen University, DK-1958 Frederiksberg, Denmark,
%

format long
format compact
Show=0;

SS=[];

[aX bX]=size(X);
[aM bM]=size(M);
aF=aX;
bF=aM;
w=bF;

Xc=X;
W=eye(w);
itimax=100;
itomax=100;
SSimax=1e-10;
SSomax=1e-12;
SSiOld=realmax;
SSoOld=realmax;
%Initialize F
if ~exist('F')
   F=X*M'/(M*M');
   FOld=F;
   I=find(F<0);
   FOld(I)=0;
   SSiOld=sum(sum( (F-FOld).^2 ));
   F=FOld;
end;
FOld=F;
SSX=sum(sum(X.^2));
MMT=M*M';
XMT=X*M';
InvMMT=1./diag(MMT);

ito=0;
convo=0;
while ~convo,
   ito=ito+1;
   convi=0;
   iti=0;
   while ~convi,
      iti=iti+1;
      
      %Iterate on variables for non-negativity
      for i=1:w,
         W(i,i)=0;
         f=XMT(:,i)-F*W*MMT(:,i);
         f=InvMMT(i)*f;
         I=find(f<0);
         if ~isempty(I),
             %f(I)=0;
             f(I)=zeros(1,length(I));
         end;
         F(:,i)=f;
         W(i,i)=1;
      end;
      
      %Estimate error now
      SSi=sum(sum( (F-FOld).^2 ));
      if SSi < sum(sum( FOld.^2 ))*SSimax | iti>itimax,
         convi=1;
      end;
      FOld=F;

   end;
   
   %Estimate error on the transformed LS problem
   SSo=sum(sum( (XMT-F*MMT).^2 ));
   if (SSoOld-SSo)/SSX<SSomax,
      convo=1;
   end;
   if ito>itomax,
      convo=1;
   end;
   SSoOld=SSo;
end;
format
```
