## Supplementary File S1 for "Robust latent-variable interpretation of *in vivo* regression models by nested resampling": normit.m

```
function [Y,D]=normit(X)
%NORMIT normalize
%
%[Y,D]=normit(X);
%Normalizes X, such that X  = Y *D
%It also applies that    X' = D'*Y'
%Always give X column wise


% Copyright (C) 1995-2006  Rasmus Bro & Claus Andersson
% Copenhagen University, DK-1958 Frederiksberg, Denmark,
%

[a b]=size(X);

Y=zeros(a,b);

SS=sqrt(sum(X.^2));
for i=1:b,
  Y(:,i)=X(:,i)./SS(i);
end;

if nargout==2,
  D=(Y'*Y)\(Y'*X);
end;
```
