## Supplementary File S1 for "Robust latent-variable interpretation of *in vivo* regression models by nested resampling": npls.m

```
function [Xfactors,Yfactors,Core,B,ypred,ssx,ssy,reg] = npls(X,Y,Fac,show);

%NPLS multilinear partial least squares regression
%
% See also:
% 'parafac' 'tucker'
%
%
% MULTILINEAR PLS  -  N-PLS
%
% INPUT
% X        Array of independent variables
% Y        Array of dependent variables
% Fac      Number of factors to compute
% 
% OPTIONAL
% show	   If show = NaN, no outputs are given
%
%
% OUTPUT
% Xfactors Holds the components of the model of X in a cell array.
%          Use fac2let to convert the parameters to scores and
%          weight matrices. I.e., for a three-way array do
%          [T,Wj,Wk]=fac2let(Xfactors);
% Yfactors Similar to Xfactors but for Y
% Core     Core array used for calculating the model of X
% B        The regression coefficients from which the scores in
%          the Y-space are estimated from the scores in the X-
%          space (U = TB);
% ypred    The predicted values of Y for one to Fac components
%          (array with dimension Fac in the last mode)
% ssx      Variation explained in the X-space.
%          ssx(f+1,1) is the sum-squared residual after first f factors.
%          ssx(f+1,2) is the percentage explained by first f factors.
% ssy      As above for the Y-space
% reg      Cell array with regression coefficients for raw (preprocessed) X
%
%
% AUXILIARY
%
% If missing elements occur these must be represented by NaN.
%
%
% [Xfactors,Yfactors,Core,B,ypred,ssx,ssy,reg] = npls(X,y,Fac);
% or short
% [Xfactors,Yfactors,Core,B] = npls(X,y,Fac);
%

% Copyright (C) 1995-2006  Rasmus Bro & Claus Andersson
% Copenhagen University, DK-1958 Frederiksberg, Denmark,


% $ Version 1.02 $ Date July 1998 $ Not compiled $
% $ Version 1.03 $ Date 4. December 1998 $ Not compiled $ Cosmetic changes
% $ Version 1.04 $ Date 4. December 1999 $ Not compiled $ Cosmetic changes
% $ Version 1.05 $ Date July 2000 $ Not compiled $ error caused weights not to be normalized for four-way and higher
% $ Version 1.06 $ Date November 2000 $ Not compiled $ increase max it and decrease conv crit to better handle difficult data
% $ Version 2.00 $ May 2001 $ Changed to array notation $ RB $ Not compiled $
% $ Version 2.01 $ June 2001 $ Changed to handle new core in X $ RB $ Not compiled $
% $ Version 2.02 $ January 2002 $ Outputs all predictions (1 - LV components) $ RB $ Not compiled $
% $ Version 2.03 $ March 2004 $ Changed initialization of u $ RB $ Not compiled $
% $ Version 2.04 $ Jan 2005 $ Modified sign conventions of scores and loads $ RB $ Not compiled $
% $ Version 3.00 $ Aug 2007 $ Serious error in sign switch fixed $ RB $ Not compiled $
% $ Version 3.01 $ Sept 2012 $ Reverted to original initialization of u $ KJ $ Important for proper interpretation of microarray model

if nargin==0
   disp(' ')
   disp(' ')
   disp(' THE N-PLS REGRESSION MODEL')
   disp(' ')
   disp(' Type <<help npls>> for more info')
   disp('  ')
   disp(' [Xfactors,Yfactors,Core,B,ypred,ssx,ssy] = npls(X,y,Fac);')
   disp(' or short')
   disp(' [Xfactors,Yfactors,Core,B] = npls(X,y,Fac);')
   disp(' ')
   return
elseif nargin<3
   error(' The inputs X, y, and Fac must be given')
end


if ~exist('show')==1|nargin<4
   show=1;
end

maxit=120;

DimX = size(X);
X = reshape(X,DimX(1),prod(DimX(2:end)));
ordX = length(DimX);if ordX==2&size(X,2)==1;ordX = 1;end
DimY = size(Y);
Y = reshape(Y,DimY(1),prod(DimY(2:end)));
ordY = length(DimY);if ordY==2&size(Y,2)==1;ordY = 1;end


[I,Jx]=size(X);
[I,Jy]=size(Y);

missX=0;
missy=0;
MissingX = 0;
MissingY = 0;
if any(isnan(X(:)))|any(isnan(Y(:)))
   if any(isnan(X(:)))
      MissingX=1;
   else
      MissingX=0;
   end
   if any(isnan(Y(:)))
      MissingY=1;
   else
      MissingY=0;
   end
   if show~=0&~isnan(show)
      disp(' ')
      disp(' Don''t worry, missing values will be taken care of')
      disp(' ')
   end
   missX=abs(1-isnan(X));
   missy=abs(1-isnan(Y));
end
crit=1e-10;
B=zeros(Fac,Fac);
T=[];
U=[];
Qkron =[];
if MissingX
   SSX=sum(sum(X(find(missX)).^2));
else
   SSX=sum(sum(X.^2));
end
if MissingY
   SSy=sum(sum(Y(find(missy)).^2));
else
   SSy=sum(sum(Y.^2));
end
ssx=[];
ssy=[];
Xres=X;
Yres=Y;
xmodel=zeros(size(X));
Q=[];
W=[];

for num_lv=1:Fac
   
   %init
   
   
%    u=rand(DimX(1),1); %Old version **IMPORTANT CHANGE FOR GENE EXPRESSION MODEL**
   if size(Yres,2)==1
     u = Yres;
   else
     [u] = pcanipals(Yres,1,0);   
   end
   
   t=rand(DimX(1),1);
   tgl=t+2;it=0;
   while (norm(t-tgl)/norm(t))>crit&it<maxit
      tgl=t;
      it=it+1;
      
      % w=X'u
      [wloads,wkron] = Xtu(X,u,MissingX,missX,Jx,DimX,ordX); %Why not run on Xres?
      
      % t=Xw
      if MissingX
         for i=1:I,
            m = find(missX(i,:));
            t(i)=X(i,m)*wkron(m)/(wkron(m)'*wkron(m)); %Why not Xres?
         end
      else
         t=X*wkron; %Why not Xres?
      end
      
      % w=X'u
      [qloads,qkron] = Xtu(Yres,t,MissingY,missy,Jy,DimY,ordY);
      % u=yq
      if MissingY
         for i=1:I
            m = find(missy(i,:));
            u(i)=Yres(i,m)*qkron(m)/(qkron(m)'*qkron(m));
         end
      else
         u=Yres*qkron;
      end
      
      if all(sign(u)<0) %Note that sign switching of y scores and weights must occur within the regression and not after to preserve the integrity of the model
          qloads{1} = -qloads{1};
          % Unfold solution
            if length(qloads)==1
               qkron = qloads{1};
            else
               qkron = kron(qloads{end},qloads{end-1});
               for o = ordY-3:-1:1
                  qkron = kron(qkron,qloads{o});
               end
            end
          % u=yq
          if MissingY
             for i=1:I
                m = find(missy(i,:));
                u(i)=Yres(i,m)*qkron(m)/(qkron(m)'*qkron(m));
             end
          else
             u=Yres*qkron;
          end
      end
   end
   
   
%    % Fix signs
%    [Factors] = signswtch({t,wloads{:}},X);
%    t = Factors{1};
%    wloads = Factors(2:end);
%    % Fix signs
%    [Factors] = signswtch({u,qloads{:}},X);
%    u = Factors{1};
%    qloads = Factors(2:end);
   
%    % Arrange t scores so they positively correlated with u
%    cc = corrcoef([t u]);
%    if sign(cc(2,1))<0
%      t = -t;
%      wloads{1}=-wloads{1};
%    end   

   % If all t scores are negative, invert results
   if all(sign(t)<0)
      t = -t;
      wloads{1} = -wloads{1};
   end
   
   T=[T t];
   for i = 1:ordX-1
      if num_lv == 1
         W{i} = wloads{i};
      else
         W{i} = [W{i} wloads{i}];
      end
   end
   U=[U u];
   for i = 1:max(ordY-1,1)
      if num_lv == 1
         Q{i} = qloads{i};
      else
         Q{i} = [Q{i} qloads{i}];
      end
   end
   Qkron = [Qkron qkron];
  
   % Make core arrays
   if ordX>1
      Xfac{1}=T;Xfac(2:ordX)=W;
      Core{num_lv} = calcore(reshape(X,DimX),Xfac,[],0,1); %Why not Xres?
   else
      Core{num_lv} = 1;
   end
%   if ordY>1
%      Yfac{1}=U;Yfac(2:ordY)=Q;
%      Ycore{num_lv} = calcore(reshape(Y,DimY),Yfac,[],0,1);
%   else
%      Ycore{num_lv} = 1;
%   end
   
   
   B(1:num_lv,num_lv)=inv(T'*T)*T'*U(:,num_lv);
   
   if Jy > 1
      if show~=0&~isnan(show)
         disp(' ') 
         fprintf('number of iterations: %g',it);
         disp(' ')
      end
   end
   
   % Make X model
   if ordX>2
      Wkron = kron(W{end},W{end-1});
   else
      Wkron = W{end};
   end
   for i = ordX-3:-1:1
      Wkron = kron(Wkron,W{i});
   end
   if num_lv>1
      xmodel=T*reshape(Core{num_lv},num_lv,num_lv^(ordX-1))*Wkron';
   else
      xmodel = T*Core{num_lv}*Wkron';
   end
   
   % Make Y model   
 %  if ordY>2
 %     Qkron = kron(Q{end},Q{end-1});
 %  else
 %     Qkron = Q{end};
 %  end
 %  for i = ordY-3:-1:1
 %     Qkron = kron(Qkron,Q{i});
 %  end
 %  if num_lv>1
 %     ypred=T*B(1:num_lv,1:num_lv)*reshape(Ycore{num_lv},num_lv,num_lv^(ordY-1))*Qkron';
 %  else
 %     ypred = T*B(1:num_lv,1:num_lv)*Ycore{num_lv}*Qkron';
 %  end
 ypred=T*B(1:num_lv,1:num_lv)*Qkron';
 Ypred(:,num_lv) = ypred(:); % Vectorize to avoid problems with different orders and the de-vectorize later on
   
   Xres=X-xmodel; 
   Yres=Y-ypred;
   if MissingX
      ssx=[ssx;sum(sum(Xres(find(missX)).^2))];
   else
      ssx=[ssx;sum(sum(Xres.^2))];
   end
   if MissingY
      ssy=[ssy;sum(sum((Y(find(missy))-ypred(find(missy))).^2))];
   else
      ssy=[ssy;sum(sum((Y-ypred).^2))];
   end
end
ypred = reshape(Ypred',[size(Ypred,2) DimY]);
ypred = permute(ypred,[2:ordY+1 1]);

ssx= [ [SSX(1);ssx] [0;100*(1-ssx/SSX(1))]];
ssy= [ [SSy(1);ssy] [0;100*(1-ssy/SSy(1))]];

if show~=0&~isnan(show)
   disp('  ')
   disp('   Percent Variation Captured by N-PLS Model   ')
   disp('  ')
   disp('   LV      X-Block    Y-Block')
   disp('   ----    -------    -------')
   ssq = [(1:Fac)' ssx(2:Fac+1,2) ssy(2:Fac+1,2)];
   format = '   %3.0f     %6.2f     %6.2f';
   for i = 1:Fac
      tab = sprintf(format,ssq(i,:)); disp(tab)
   end
end

Xfactors{1}=T;
for j = 1:ordX-1
   Xfactors{j+1}=W{j};
end

Yfactors{1}=U;
for j = 1:max(ordY-1,1)
   Yfactors{j+1}=Q{j};
end

% Calculate regression coefficients that apply directly to X
  if nargout>7
    if length(DimY)>2
      error(' Regression coefficients are only calculated for models with vector Y or multivariate Y (not multi-way Y)')
    end
      R = outerm(W,0,1);
    for iy=1:size(Y,2)
      if length(DimX) == 2
        dd = [DimX(2) 1];
      else
        dd = DimX(2:end);
      end
      for i=1:Fac
        sR = R(:,1:i)*B(1:i,1:i)*diag(Q{1}(iy,1:i));
        ssR = sum( sR',1)';
        reg{iy,i} = reshape( ssR ,dd);
      end       
    end
    
  end


function [wloads,wkron] = Xtu(X,u,Missing,miss,J,DimX,ord);


% w=X'u
if Missing
   for i=1:J
      m = find(miss(:,i));
      if (u(m)'*u(m))~=0
        ww=X(m,i)'*u(m)/(u(m)'*u(m));
      else
        ww=X(m,i)'*u(m);
      end
      if length(ww)==0
         w(i)=0;
      else
         w(i)=ww;
      end
   end
else
   w=X'*u;
end

% Reshape to array
if length(DimX)>2
   w_reshaped=reshape(w,DimX(2),prod(DimX(3:length(DimX))));
else
   w_reshaped = w(:);
end


% Find one-comp decomposition
if length(DimX)==2
   wloads{1} = w_reshaped/norm(w_reshaped);
   
elseif length(DimX)==3&~any(isnan(w_reshaped))
   [w1,s,w2]=svd(w_reshaped);
   wloads{1}=w1(:,1);
   wloads{2}=w2(:,1);
else
   wloads=parafac(reshape(w_reshaped,DimX(2:length(DimX))),1,[0 2 0 0 NaN]');
   for j = 1:length(wloads);
      wloads{j} = wloads{j}/norm(wloads{j});
   end
end

% Apply sign convention
for i = 1:length(wloads)
   sq = (wloads{i}.^2).*sign(wloads{i});
   wloads{i} = wloads{i}*sign(sum(sq));
end


% Unfold solution
if length(wloads)==1
   wkron = wloads{1};
else
   wkron = kron(wloads{end},wloads{end-1});
   for o = ord-3:-1:1
      wkron = kron(wkron,wloads{o});
   end
end


function [Factors,it,err,corcondia]=parafac(X,Fac,Options,const,OldLoad,FixMode,Weights);

% PARAFAC multiway parafac model
%
% See also:
% 'npls' 'tucker' 'dtld' 'gram'
%
%
%     ___________________________________________________
%
%                  THE PARAFAC MODEL
%     ___________________________________________________
% 
% [Factors,it,err,corcondia,Weights] = parafac(X,Fac,Options,const,OldLoad,FixMode,Weights);
%
% or skipping optional in/outputs
%
% Factors = parafac(X,Fac);
%
% Algorithm for computing an N-way PARAFAC model. Optionally
% constraints can be put on individual modes for obtaining 
% orthogonal, nonnegative, or unimodal solutions. The algorithm
% also handles missing data. For details of PARAFAC 
% modeling see R. Bro, Chemom. Intell. Lab. Syst., 1997.
%
% Several possibilities exist for speeding up the algorithm. 
% Compressing has been incorporated, so that large arrays can be
% compressed by using Tucker (see Bro & Andersson, Chemom. 
% Intell. Lab. Syst., 1998).
% Another acceleration method incorporated here is to 
% extrapolate the individual loading elements a number of 
% iterations ahead after a specified number of iterations.
%
% A temporary MAT-file called TEMP.mat is saved for every 
% 50 iterations. IF the computer breaks down or the model 
% seems to be good enough, one can break the program and 
% load the last saved estimate. The loadings in TEMP.MAT
% are given a cell array as described below and can be 
% converted to A, B, C etc. by FAC2LET.M
% 
% All loading vectors except in first mode are normalized, 
% so that all variance is kept in the first mode (as is 
% common in two-way PCA). The components are arranged as
% in PCA. After iterating, the most important component is
% made the first component etc.
%
%
%
% ----------------------INPUT---------------------
%
% X          X is the input array, which can be from three- to N-way (also
%            twoway if the third mode is interpreted as a onedimensional
%            mode). 
%
% Fac		    No of factors/components sought.
%
%
% ----------------OPTIONAL INPUT---------------------
%
% Options    Optional parameters. If not given or set to zero or [], 
%            defaults will be used. If you want Options(5) to be 2 and
%            not change others, simply write Options(5)=2. Even if Options
%            hasn't been defined Options will contain zeros except its
%            fifth element.
%
%            Options(1) - Convergence criterion
%            The relative change in fit for which the algorithm stops.
%            Standard is 1e-6, but difficult data might require a lower value.
%  
%            Options(2) - Initialization method
%            This option is ignored if PARAFAC is started with old values.
%            If no default values are given the default Options(2) is 0.
%            The advantage of using DTLD or SVD for initialization is that
%            they often provide good starting values. However, since the 
%            initial values are then fixed, repeating the fitting will give
%            the exact same solution. Therefore it is not possible to substantiate
%            if a local minimum has been reached. To avoid that use an initialization
%            based on random values (2).
%
%            0  = fit using DTLD/GRAM for initialization (default if three-way and no missing)
%            1  = fit using SVD vectors for initialization (default if higher than three-way or missing)
%            2  = fit using random orthogonalized values for initialization
%            10 = fit using the best-fitting models of several models
%            fitted using a few iterations
%
%            Options(3) - Plotting options
%            2=produces several graphical outputs (loadings shown during iterations)
%            1=as above, but graphics only shown after convergence
%            0=no plots
%
%            Options(4) - Not user-accesible
% 
%            Options(5) - How often to show fit
%            Determines how often the deviation between the model and the data
%            is shown. This is helpful for adjusting the output to the number
%            of iterations. Default is 10. If showfit is set to NaN, almost no
%            outputs are given 
%
%            Options(6) - Maximal number of iterations
%            Maximal number of iterations allowed. Default is 2500.
%
% const      A vector telling type of constraints put on the loadings of the
%            different modes. Same size as DimX but the i'th element tells
%            what constraint is on that mode.
%            0 => no constraint,
%            1 => orthogonality
%            2 => nonnegativity
%            3 => unimodality (and nonnegativitiy)
%            If const is not defined, no constraints are used.
%            For no constraints in a threeway problem const = [0 0 0]
%
% OldLoad    If initial guess of the loadings is available. OldLoad should be
%            given a cell array where OldLoad{1}=A,OldLoad{2}=B etc.
%
% FixMode    FixMode is a binary vector of same sixe as DimX. If 
%            FixMode(i) = 1 => Mode i is fixed (requires old values given)
%            FixMode(i) = 0 => Mode i is not fixed hence estimated
%            Ex.: FixMode = [0 1 1] find the scores of a data set given the loadings.
%            When some modes are fixed, the numbering of the components will 
%            also be fixed. Normally components are sorted according to variance
%            as in PCA, but this will not be performed if some modes are fixed.
%
% Weights    If a matrix of the same size as X is given, weighted regression
%            is performed using the weights in the matrix Weights. 
%
% ---------------------OUTPUT---------------------
%
% Factors    PARAFAC estimate of loadings in one matrix. For a 3 component
%            solution to a 4 x 3 x 3 array the loadings A, B & C will be
%            stored in a 3 element cell vector:
%            Factors{1}=A,
%            Factors{2}=B
%            Factors{3}=C
%            etc.
%
%            Use FAC2LET.M for converting to "normal" output.
%
% it         Number of iterations used. Can be helpful for checking if the algorithm
%            has converged or simply hit the maximal number of iterations (default 2500).
%
% err        The fit of the model = the sum of squares of errors (not including missing
%            elements).
%
% Corcondia  Core consistency test. Should ideally be 100%. If significantly below
%            100% the model is not valid
%
%
%
% OTHER STUFF
%  
%  Missing values are handled by expectation maximization only. Set all 
%  missing values to NaN
%
%  COMMAND LINE (SHORT)
%
%  Factors = parafac(X,Fac);
%

% Copyright, 1998 - 
% This M-file and the code in it belongs to the holder of the
% copyrights and is made public under the following constraints:
% It must not be changed or modified and code cannot be added.
% The file must be regarded as read-only. Furthermore, the
% code can not be made part of anything but the 'N-way Toolbox'.
% In case of doubt, contact the holder of the copyrights.
%
% Rasmus Bro
% Chemometrics Group, Food Technology
% Department of Food and Dairy Science
% Royal Veterinary and Agricultutal University
% Rolighedsvej 30, DK-1958 Frederiksberg, Denmark
% Phone  +45 35283296
% Fax    +45 35283245
%

% $ Version 1.03 $ Date 1. October   1998 $ Not compiled $ Changed sign-convention because of problems with centered data
% $ Version 1.04 $ Date 18. February 1999 $ Not compiled $ Removed auxiliary line
% $ Version 1.06 $ Date 1. December  1999 $ Not compiled $ Fixed bug in low fit error handling
% $ Version 1.07 $ Date 17. January  2000 $ Not compiled $ Fixed bug in nnls handling so that the algorithm is not stopped until nonnegative appear
% $ Version 1.08 $ Date 21. January  2000 $ Not compiled $ Changed init DTLD so that primarily negative loadings are reflected if possible
% $ Version 1.09 $ Date 30. May 2000 $ Not compiled $ changed name noptioPF to noptiopf
% $ Version 2.00 $ May 2001 $ Changed to array notation $ RB $ Not compiled $
% $ Version 2.001 $ June 2001 $ Fixed error in weighted regression $ RB $ Not compiled $

NumbIteraInitia=20;

if nargin==0
   disp(' ')
   disp(' ')
   disp(' THE PARAFAC MODEL')
   disp(' ')
   disp(' Type <<help parafac>> for more info')
   disp('  ')
   disp(' [Factors,it,err,Corcondia] = parafac(X,Fac,Options,const,OldLoad,FixMode,Weights);')
   disp(' or short')
   disp(' Factors = parafac(X,Fac);')
   disp(' ')
   disp(' Options=[Crit Init Plot NotUsed ShowFit MaxIt]')
   disp(' ')
   disp(' ')
   disp(' EXAMPLE:')
   disp(' To fit a four-component PARAFAC model to X of size 6 x 2 x 200 x 3 type')
   disp(' Factors=parafac(X,4)')
   disp(' and to obtain the scores and loadings from the output type')
   disp(' [A,B,C,D]=fac2let(Factors);')
   return
elseif nargin<2
   error(' The inputs X, and Fac must be given')
end

DimX = size(X);
X = reshape(X,DimX(1),prod(DimX(2:end)));

if nargin<3
   load noptiopf
   OptionsDefault=Options;
else
   % Call the current Options OptionsHere and load default to use if some of the current settings should be default
   Options=Options(:);
   I=length(Options);
   if I==0
      Options=zeros(8,1);
   end
   I=length(Options);
   if I<8
      Options=[Options;zeros(8-I,1)];
   end
   OptionsHere=Options;
   load noptiopf
   OptionsDefault=Options;
   Options=OptionsHere;
end

if ~exist('OldLoad')==1
   OldLoad=0;
elseif length(OldLoad)==0
   OldLoad=0;
end

% Convergence criteria
if Options(1,1)==0
   Options(1,1)=OptionsDefault(1,1);
end
crit=Options(1);

% Initialization
if ~any(Options(2))
   Options(2)=OptionsDefault(2);
end
Init=Options(2);

% Interim plotting
Plt=Options(3,1);
if ~any([0 1 2]==Plt)
   error(' Options(3,1) - Plotting - not set correct; must be 0,1, or 2')
end

if Options(5,1)==0
   Options(5,1)=OptionsDefault(5,1);
end
showfit=Options(5,1);
if isnan(showfit)
   showfit=-1;
end
if showfit<-1|round(showfit)~=showfit
   error(' Options(5,1) - How often to show fit - not set correct; must be positive integer or -1')
end

if Options(6,1)==0
   Options(6,1)=OptionsDefault(6,1);
   maxit=Options(6,1);
elseif Options(6)>0&round(Options(6))==Options(6)
   maxit=Options(6,1);
else
   error(' Options(6,1) - Maximal number of iterations - not set correct; must be positive integer')
end

ShowPhi=0; % Counter. Tuckers congruence coef/Multiple cosine/UUC shown every ShowPhiWhen'th time the fit is shown
ShowPhiWhen=10;
MissConvCrit=1e-4; % Convergence criterion for estimates of missing values
NumberOfInc=0; % Counter for indicating the number of iterations that increased the fit. ALS algorithms ALLWAYS decrease the fit, but using outside knowledge in some sense (approximate equality or iteratively reweighting might cause the algorithm to diverge

% INITIALIZE 
if showfit~=-1
   disp(' ') 
   disp(' PRELIMINARY')
   disp(' ')
end
ord=length(DimX);

if showfit~=-1
   disp([' A ',num2str(Fac),'-component model will be fitted'])
end

if exist('const')~=1
   const=zeros(size(DimX));
elseif length(const)~=ord
   const=zeros(size(DimX));
   if showfit~=-1
      disp(' Constraints are not given properly')
   end
end

if showfit~=-1
   for i=1:ord
      if const(i)==0
         disp([' No constraints on mode ',num2str(i)])
      elseif const(i)==1
         disp([' Orthogonality on mode ',num2str(i)])
      elseif const(i)==2
         disp([' Nonnegativity on mode ',num2str(i)])
      elseif const(i)==3
         disp([' Unimodality on mode ',num2str(i)])
      end
   end
end

% Check if orthogonality required on all modes
if max(max(const))==1
   if min(min(const))==1,disp(' ')
      disp(' Not possible to orthogonalize all modes in this implementation.')
      error(' Contact the authors for further information')
   end
end

if exist('FixMode')==1
   if length(FixMode)~=ord
      FixMode = zeros(1,ord);
   end
else
   FixMode = zeros(1,ord);
end

if showfit~=-1
   if any(FixMode)
      disp([' The loadings of mode : ',num2str(find(FixMode(:)')),' are fixed']) 
   end
end
if exist('Weights')~=1
   Weights=[];
end

% Display convergence criterion
if showfit~=-1
   disp([' The convergence criterion is ',num2str(crit)]) 
end

% Define loading as one ((r1*r2*r3*...*r7)*Fac x 1) vector [A(:);B(:);C(:);...].
% The i'th loading goes from lidx(i,1) to lidx(i,2)
lidx=[1 DimX(1)*Fac];
for i=2:ord
   lidx=[lidx;[lidx(i-1,2)+1 sum(DimX(1:i))*Fac]];
end

% Check if weighted regression required
if size(Weights,1)==size(X,1)&prod(size(Weights))/size(X,1)==size(X,2)
    Weights = reshape(Weights,size(Weights,1),prod(size(Weights))/size(X,1));
   if showfit~=-1
      disp(' Given weights will be used for weighted regression')
   end
   DoWeight=1;
else
   if showfit~=-1
      disp(' No weights given')
   end
   DoWeight=0;
end

% Make idx matrices if missing values
if any(isnan(X(:)))
   MissMeth=1;
else
   MissMeth=0;
end
if MissMeth
   id=sparse(find(isnan(X)));
   idmiss2=sparse(find(~isnan(X)));
   if showfit~=-1
      disp([' ', num2str(100*(length(id)/prod(DimX))),'% missing values']);
      disp(' Expectation maximization will be used for handling missing values')
   end
   SSX=sum(sum(X(idmiss2).^2)); % To be used for evaluating the %var explained
   % If weighting to zero should be used
   % Replace missing with mean values or model estimates initially
   if length(OldLoad)==sum(DimX)*Fac
      model=nmodel(OldLoad);
      model = reshape(model,DimX);
      X(id)=model(id);
   else
      meanX=mean(X(find(~isnan(X))));
      meanX=mean(meanX);
      X(id)=meanX*ones(size(id));
   end
else
   if showfit~=-1
      disp(' No missing values')
   end
   SSX=sum(sum(X.^2)); % To be used for evaluating the %var explained
end

% Check if weighting is tried used together with unimodality or orthogonality
if any(const==3)|any(const==1)
   if DoWeight==1
      disp(' ')
      disp(' Weighting is not possible together with unimodality and orthogonality.')
      disp(' It can be done using majorization, but has not been implemented here')
      disp(' Please contact the authors for further information')
      error
   end
end

% Acceleration
acc=-5;     
do_acc=1;   % Do acceleration every do_acc'th time
acc_pow=2;  % Extrapolate to the iteration^(1/acc_pow) ahead
acc_fail=0; % Indicate how many times acceleration have failed 
max_fail=4; % Increase acc_pow with one after max_fail failure
if showfit~=-1
   disp(' Line-search acceleration scheme initialized')
end

% Find initial guesses for the loadings if no initial values are given

% Use old loadings
if length(OldLoad)==ord % Use old values
   if showfit~=-1
      disp(' Using old values for initialization')
   end
   Factors=OldLoad;
   % Use DTLD
elseif Init==0
   if min(DimX)>1&ord==3&MissMeth==0
      if showfit~=-1
         disp(' Using direct trilinear decomposition for initialization')
      end
      [A,B,C]=dtld(reshape(X,DimX),Fac);
      A=real(A);B=real(B);C=real(C);
      % Check for signs and reflect if appropriate
      for f=1:Fac
         if sign(sum(A(:,f)))<0
            if sign(sum(B(:,f)))<0
               B(:,f)=-B(:,f);
               A(:,f)=-A(:,f);
            elseif sign(sum(C(:,f)))<0
               C(:,f)=-C(:,f);
               A(:,f)=-A(:,f);
            end
         end
         if sign(sum(B(:,f)))<0
            if sign(sum(C(:,f)))<0
               C(:,f)=-C(:,f);
               B(:,f)=-B(:,f);
            end
         end
      end
      Factors{1}=A;Factors{2}=B;Factors{3}=C;

   else
      if showfit~=-1
         disp(' Using singular values for initialization')
      end
      Factors=ini(X,Fac,2);
   end
   
   % Use SVD 
elseif Init==1
   if showfit~=-1
      disp(' Using singular values for initialization')
   end
   Factors=ini(X,Fac,2);
   
   % Use random (orthogonal)
elseif Init==2
   if showfit~=-1
      disp(' Using orthogonal random for initialization')
   end
   Factors=ini(X,Fac,1);
   
elseif Init==3
   error(' Initialization option set to three has been changed to 10')
   
   % Use several small ones of the above
elseif Init==10
   if showfit~=-1
      disp(' Using several small runs for initialization')
   end
   Opt=Options;
   Opt(5) = NaN;
   Opt(6) = NumbIteraInitia;
   Opt(2) = 0;
   ERR=[];
   [Factors,it,err] = parafac(X,Fac,Opt,const,[],[],Weights);
   ERR = [ERR;err];
   Opt(2) = 1;
   [F,it,Err] = parafac(X,Fac,Opt,const,[],[],Weights);
   ERR=[ERR;Err];
   if Err<err
      Factors=F;
      err=Err;
   end
   Opt(2)=2;
   for rep=1:3
      [F,it,Err]=parafac(X,Fac,Opt,const,[],[],Weights);
      ERR=[ERR;Err];
      if Err<err
         Factors=F;
         err=Err;
      end
   end
   if showfit~=-1
      disp(' ')
      disp(' Obtained fit-values')
      disp([' Method   Fit'])
      disp([' DTLD     ',num2str(ERR(1))])
      disp([' SVD      ',num2str(ERR(2))])
      disp([' RandOrth ',num2str(ERR(3))])
      disp([' RandOrth ',num2str(ERR(4))])
      disp([' RandOrth ',num2str(ERR(5))])
   end
else
   error(' Problem in PARAFAC initialization - Not set correct')
end
% Convert to old format
ff = [];
for f=1:length(Factors)
 ff=[ff;Factors{f}(:)];
end
Factors = ff;

% ALTERNATING LEAST SQUARES
err=SSX;
f=2*crit;
it=0;
connew=2;conold=1; % for missing values
ConstraintsNotRight = 0; % Just to ensure that iterations are not stopped if constraints are not yet fully imposed

if showfit~=-1
   disp(' ')
   disp(' Sum-of-Squares   Iterations  Explained')
   disp(' of residuals                 variation')
end

while ((f>crit) | (norm(connew-conold)/norm(conold)>MissConvCrit) | ConstraintsNotRight) & it<maxit
   conold=connew; % for missing values
   it=it+1;
   acc=acc+1; 
   if acc==do_acc;
      Load_o1=Factors;
   end
   if acc==do_acc+1;
      acc=0;Load_o2=Factors;
      Factors=Load_o1+(Load_o2-Load_o1)*(it^(1/acc_pow));
      % Convert to new format
      clear ff,id1 = 0;
      for i = 1:length(DimX) 
         id2 = sum(DimX(1:i).*Fac);ff{i} = reshape(Factors(id1+1:id2),DimX(i),Fac);id1 = id2;
      end
      model=nmodel(ff);
      model = reshape(model,DimX(1),prod(DimX(2:end)));
      
      if MissMeth
         connew=model(id);
         errX=X-model;
         if DoWeight==0
            nerr=sum(sum(errX(idmiss2).^2));
         else
            nerr=sum(sum((Weights(idmiss2).*errX(idmiss2)).^2));
         end
      else
         if DoWeight==0
            nerr=sum(sum((X-model).^2));
         else
            nerr=sum(sum((X.*Weights-model.*Weights).^2));
         end
      end
      if nerr>err
         acc_fail=acc_fail+1;
         Factors=Load_o2;
         if acc_fail==max_fail,
            acc_pow=acc_pow+1+1;
            acc_fail=0;
            if showfit~=-1
               disp(' Reducing acceleration');
            end
         end
      else
         if MissMeth
            X(id)=model(id);
         end
      end
   end
   
   
   if DoWeight==0
      for ii=ord:-1:1
         if ii==ord;
            i=1;
         else
            i=ii+1;
         end
         idd=[i+1:ord 1:i-1];
         l_idx2=lidx(idd,:);
         dimx=DimX(idd);
         if ~FixMode(i)
            L1=reshape(Factors(l_idx2(1,1):l_idx2(1,2)),dimx(1),Fac);
            if ord>2
               L2=reshape(Factors(l_idx2(2,1):l_idx2(2,2)),dimx(2),Fac);
               Z=krb(L2,L1);
            else
               Z = L1;
            end
            for j=3:ord-1
               L1=reshape(Factors(l_idx2(j,1):l_idx2(j,2)),dimx(j),Fac);
               Z=krb(L1,Z);
            end
            ZtZ=Z'*Z;
            ZtX=Z'*X';
            OldLoad=reshape(Factors(lidx(i,1):lidx(i,2)),DimX(i),Fac);
            L=pfls(ZtZ,ZtX,DimX(i),const(i),OldLoad,DoWeight,Weights);
            Factors(lidx(i,1):lidx(i,2))=L(:);
         end
         x=zeros(prod(DimX([1:ii-1 ii+1:ord])),DimX(ii));  % Rotate X so the current last mode is the first
         x(:)=X;
         X=x';
      end
   else
      for ii=ord:-1:1
         if ii==ord;
            i=1;
         else
            i=ii+1;
         end
         idd=[i+1:ord 1:i-1];
         l_idx2=lidx(idd,:);
         dimx=DimX(idd);
         if ~FixMode(i)
            L1=reshape(Factors(l_idx2(1,1):l_idx2(1,2)),dimx(1),Fac);
            if ord>2
               L2=reshape(Factors(l_idx2(2,1):l_idx2(2,2)),dimx(2),Fac);
               Z=krb(L2,L1);
            else
               Z = L1;
            end
            for j=3:ord-1
               L1=reshape(Factors(l_idx2(j,1):l_idx2(j,2)),dimx(j),Fac);
               Z=krb(L1,Z);
            end
            OldLoad=reshape(Factors(lidx(i,1):lidx(i,2)),DimX(i),Fac);
            L=pfls(Z,X,DimX(i),const(i),OldLoad,DoWeight,Weights);
            Factors(lidx(i,1):lidx(i,2))=L(:);
         end
         x=zeros(prod(DimX([1:ii-1 ii+1:ord])),DimX(ii));
         x(:)=X;
         X=x';
         x(:)=Weights;
         Weights=x';
      end
   end
   
   % EVALUATE SOFAR
   % Convert to new format
   clear ff,id1 = 0;
   for i = 1:length(DimX) 
      id2 = sum(DimX(1:i).*Fac);
      ff{i} = reshape(Factors(id1+1:id2),DimX(i),Fac);
      id1 = id2;
   end
   model=nmodel(ff);
   model = reshape(model,DimX(1),prod(DimX(2:end)));
   if MissMeth  % Missing values present
      connew=model(id);
      X(id)=model(id);
      errold=err;
      errX=X-model;
      if DoWeight==0
         err=sum(sum(errX(idmiss2).^2));
      else
         err=sum(sum((Weights(idmiss2).*errX(idmiss2)).^2));
      end
   else
      errold=err;
      if DoWeight==0
         err=sum(sum((X-model).^2));
      else
         err=sum(sum((Weights.*(X-model)).^2));
      end
   end
   
   if err<1000*eps, % Getting close to the machine uncertainty => stop
      disp(' WARNING')
      disp(' The misfit is approaching the machine uncertainty')
      disp(' If pure synthetic data is used this is OK, otherwise if the')
      disp(' data elements are very small it might be appropriate ')
      disp(' to multiply the whole array by a large number to increase')
      disp(' numerical stability. This will only change the solution ')
      disp(' by a scaling constant')
      f = 0;
   else
      f=abs((err-errold)/err);
      if f<crit % Convergence: then check that constraints are fulfilled
         if any(const==2)|any(const==3) % If nnls or unimodality imposed
            for i=1:ord % Extract the 
               if const(i)==2|const(i)==3 % If nnls or unimodality imposed
                  Loadd = Factors(sum(DimX(1:i-1))*Fac+1:sum(DimX(1:i))*Fac);
                  if any(Loadd<0)
                     ConstraintsNotRight=1;
                  else
                     ConstraintsNotRight=0;
                  end
               end
            end
         end
      end
   end
   
   if it/showfit-round(it/showfit)==0
      if showfit~=-1,
         ShowPhi=ShowPhi+1;
         if ShowPhi==ShowPhiWhen,
            ShowPhi=0;
            if showfit~=-1,
               disp(' '),
               disp('    Tuckers congruence coefficient'),
               % Convert to new format
               clear ff,id1 = 0;
               for i = 1:length(DimX) 
                  id2 = sum(DimX(1:i).*Fac);ff{i} = reshape(Factors(id1+1:id2),DimX(i),Fac);id1 = id2;
               end
               [phi,out]=ncosine(ff,ff);
               disp(phi),
               if MissMeth
                  fprintf(' Change in estim. missing values %12.10f',norm(connew-conold)/norm(conold));
                  disp(' ')
                  disp(' ')
               end
               disp(' Sum-of-Squares   Iterations  Explained')
               disp(' of residuals                 variation')
            end
         end
         if DoWeight==0
            PercentExpl=100*(1-err/SSX);
         else
            PercentExpl=100*(1-sum(sum((X-model).^2))/SSX);
         end
         fprintf(' %12.10f       %g        %3.4f    \n',err,it,PercentExpl);
         if Plt==2
            % Convert to new format
            clear ff,id1 = 0;
            for i = 1:length(DimX) 
               id2 = sum(DimX(1:i).*Fac);ff{i} = reshape(Factors(id1+1:id2),DimX(i),Fac);id1 = id2;
            end
            pfplot(reshape(X,DimX),ff,Weights',[0 0 0 0 0 0 0 1]);
            drawnow
         end
      end
   end
   
   
   
   % Make safety copy of loadings and initial parameters in temp.mat
   if it/50-round(it/50)==0
      save temp Factors
   end
   
   % JUDGE FIT
   if err>errold
      NumberOfInc=NumberOfInc+1;
   end
   
end % while f>crit


% CALCULATE TUCKERS CONGRUENCE COEFFICIENT
if showfit~=-1 & DimX(1)>1
   disp(' '),disp('   Tuckers congruence coefficient')
   % Convert to new format
   clear ff,id1 = 0;
   for i = 1:length(DimX) 
      id2 = sum(DimX(1:i).*Fac);ff{i} = reshape(Factors(id1+1:id2),DimX(i),Fac);id1 = id2;
   end
   [phi,out]=ncosine(ff,ff);
   disp(phi)
   disp(' ')
   if max(max(abs(phi)-diag(diag(phi))))>.85
      disp(' ')
      disp(' ')
      disp(' WARNING, SOME FACTORS ARE HIGHLY CORRELATED.')
      disp(' ')
      disp(' You could decrease the number of components. If this')
      disp(' does not help, try one of the following')
      disp(' ')
      disp(' - If systematic variation is still present you might')
      disp('   wanna decrease your convergence criterion and run')
      disp('   one more time using the loadings as initial guess.')
      disp(' ')
      disp(' - Or use another preprocessing (check for constant loadings)')
      disp(' ')
      disp(' - Otherwise try orthogonalising some modes,')
      disp(' ')
      disp(' - Or use Tucker3/Tucker2,')
      disp(' ')
      disp(' - Or a PARAFAC with some modes collapsed (if # modes > 3)')
      disp(' ')
   end
end


% SHOW FINAL OUTPUT

if DoWeight==0
   PercentExpl=100*(1-err/SSX);
else
   PercentExpl=100*(1-sum(sum((X-model).^2))/SSX);
end
if showfit~=-1
   fprintf(' %12.10f       %g        %3.4f \n',err,it,PercentExpl);
   if NumberOfInc>0
      disp([' There were ',num2str(NumberOfInc),' iterations that increased fit']);
   end
end


% POSTPROCES LOADINGS (ALL VARIANCE IN FIRST MODE)
A=reshape(Factors(lidx(1,1):lidx(1,2)),DimX(1),Fac);
for i=2:ord
   B=reshape(Factors(lidx(i,1):lidx(i,2)),DimX(i),Fac);
   for ff=1:Fac
      A(:,ff)=A(:,ff)*norm(B(:,ff));
      B(:,ff)=B(:,ff)/norm(B(:,ff));
   end
   Factors(lidx(i,1):lidx(i,2))=B(:);
end
Factors(lidx(1,1):lidx(1,2))=A(:);
if showfit~=-1
   disp(' ')
   disp(' Components have been normalized in all but the first mode')
end

% PERMUTE SO COMPONENTS ARE IN ORDER AFTER VARIANCE DESCRIBED (AS IN PCA) IF NO FIXED MODES
if ~any(FixMode)
   A=reshape(Factors(lidx(1,1):lidx(1,2)),DimX(1),Fac);
   [out,order]=sort(diag(A'*A));
   order=flipud(order);
   A=A(:,order);
   Factors(lidx(1,1):lidx(1,2))=A(:);
   for i=2:ord
      B=reshape(Factors(lidx(i,1):lidx(i,2)),DimX(i),Fac);
      B=B(:,order);
      Factors(lidx(i,1):lidx(i,2))=B(:);
   end  
   if showfit~=-1
      disp(' Components have been ordered according to contribution')
   end
elseif showfit ~= -1
   disp(' Some modes fixed hence no sorting of components performed')
end

% APPLY SIGN CONVENTION IF NO FIXED MODES


%  FixMode=1
if ~any(FixMode)&~(any(const==2)|any(const==3))
   Sign = ones(1,Fac);
   for i=ord:-1:2
      A=reshape(Factors(lidx(i,1):lidx(i,2)),DimX(i),Fac);
      Sign2=ones(1,Fac);
      for ff=1:Fac
         [out,sig]=max(abs(A(:,ff)));
         Sign(ff) = Sign(ff)*sign(A(sig,ff));
         Sign2(ff) = sign(A(sig,ff));
      end
      A=A*diag(Sign2);
      Factors(lidx(i,1):lidx(i,2))=A(:);
   end 
   A=reshape(Factors(lidx(1,1):lidx(1,2)),DimX(1),Fac);
   A=A*diag(Sign);
   Factors(lidx(1,1):lidx(1,2))=A(:);
   if showfit~=-1
      disp(' Components have been reflected according to convention')
   end
   
end 

% TOOLS FOR JUDGING SOLUTION
if nargout>3      
   x=X;
   if MissMeth
      x(id)=NaN*id;
   end
   % Convert to new format
   clear ff,id1 = 0;
   for i = 1:length(DimX) 
      id2 = sum(DimX(1:i).*Fac);ff{i} = reshape(Factors(id1+1:id2),DimX(i),Fac);id1 = id2;
   end
   corcondia=corcond(reshape(x,DimX),ff,Weights,1);
end

if Plt==1|Plt==2
   % Convert to new format
   clear ff,id1 = 0;
   for i = 1:length(DimX) 
      id2 = sum(DimX(1:i).*Fac);ff{i} = reshape(Factors(id1+1:id2),DimX(i),Fac);id1 = id2;
   end

   pfplot(reshape(X,DimX),ff,Weights,ones(1,8));
end

% Show which criterion stopped the algorithm
if showfit~=-1
   if ((f<crit) & (norm(connew-conold)/norm(conold)<MissConvCrit))
      disp(' The algorithm converged')
   elseif it==maxit
      disp(' The algorithm did not converge but stopped because the')
      disp(' maximum number of iterations was reached')
   elseif f<eps
      disp(' The algorithm stopped because the change in fit is now')
      disp(' smaller than the machine uncertainty.')
   else
      disp(' Algorithm stopped for some mysterious reason')
   end
end

% Convert to new format
clear ff,id1 = 0;
for i = 1:length(DimX) 
   id2 = sum(DimX(1:i).*Fac);ff{i} = reshape(Factors(id1+1:id2),DimX(i),Fac);id1 = id2;
end
Factors = ff;


function [A,B,C,fit]=dtld(X,F,SmallMode);

%DTLD direct trilinear decomposition
%
% See also:
% 'gram', 'parafac'
%
% Copyright, 1998 - 
% This M-file and the code in it belongs to the holder of the
% copyrights and is made public under the following constraints:
% It must not be changed or modified and code cannot be added.
% The file must be regarded as read-only. Furthermore, the
% code can not be made part of anything but the 'N-way Toolbox'.
% In case of doubt, contact the holder of the copyrights.
%
% Rasmus Bro
% Chemometrics Group, Food Technology
% Department of Food and Dairy Science
% Royal Veterinary and Agricultutal University
% Rolighedsvej 30, DK-1958 Frederiksberg, Denmark
% Phone  +45 35283296
% Fax    +45 35283245
%
%
%
% DIRECT TRILINEAR DECOMPOSITION
%
% calculate the parameters of the three-
% way PARAFAC model directly. The model
% is not the least-squares but will be close
% to for precise data with little model-error
%
% This implementation works with an optimal
% compression using least-squares Tucker3 fitting
% to generate two pseudo-observation matrices that
% maximally span the variation of all samples. per
% default the mode of smallest dimension is compressed
% to two samples, while the remaining modes are 
% compressed to dimension F.
% 
% For large arrays it is fastest to have the smallest
% dimension in the first mode
%
% INPUT
% [A,B,C]=dtld(X,F);
% X is the I x J x K array
% F is the number of factors to fit
% An optional parameter may be given to enforce which
% mode is to be compressed to dimension two
%
% Copyright 1998
% Rasmus Bro, KVL
%

% $ Version 2.00 $ May 2001 $ Changed to array notation $ RB $ Not compiled $
% $ Version 1.02 $ Date 28. July 1998 $ Not compiled $
% $ Version 1.03 $ Date 25. April 1999 $ Not compiled $

DimX = size(X);
X = reshape(X,DimX(1),prod(DimX(2:end)));

DontShowOutput = 1;

%rearrange X so smallest dimension is in first mode


if nargin<4
  [a,SmallMode] = min(DimX);
  X = nshape(reshape(X,DimX),SmallMode);
  DimX = DimX([SmallMode 1:SmallMode-1 SmallMode+1:3]);
  Fac   = [2 F F];
else
  X = nshape(reshape(X,DimX),SmallMode);
  DimX = DimX([SmallMode 1:SmallMode-1 SmallMode+1:3]);
  Fac   = [2 F F];
end
f=F;
if F==1;
  Fac   = [2 2 2];
  f=2;
end 


if DimX(1) < 2
  error(' The smallest dimension must be > 1')
end

if any(DimX(2:3)-Fac(2:3)<0)
  error(' This algorithm requires that two modes are of dimension not less the number of components')
end


% Compress data into a 2 x F x F array. Only 10 iterations are used since exact SL fit is insignificant; only obtaining good truncated bases is important
[Factors,Gt]=tucker(reshape(X,DimX),Fac,[0 0 0 0 NaN 10]);
% Convert to old format
Gt = reshape(Gt,size(Gt,1),prod(size(Gt))/size(Gt,1));

[At,Bt,Ct]=fac2let(Factors);

% Fit GRAM to compressed data
[Bg,Cg,Ag]=gram(reshape(Gt(1,:),f,f),reshape(Gt(2,:),f,f),F);

% De-compress data and find A


BB = Bt*Bg;
CC = Ct*Cg;
AA = X*pinv(krb(CC,BB)).';

if SmallMode == 1
  A=AA;
  B=BB;
  C=CC;
elseif SmallMode == 2 
  A=BB;
  B=AA;
  C=CC;
elseif SmallMode == 3
  A=BB;
  B=CC;
  C=AA;
end

fit = sum(sum(abs(X - AA*krb(CC,BB).').^2));
if ~DontShowOutput
  disp([' DTLD fitted raw data with a sum-squared error of ',num2str(fit)])
end


function mwa = outerm(facts,lo,vect)

if nargin < 2
  lo = 0;
end
if nargin < 3
  vect = 0;
end
order = length(facts);
if lo == 0
  mwasize = zeros(1,order);
else
  mwasize = zeros(1,order-1);
end
k = 0;
for i = 1:order
  if i ~= lo
    [m,n] = size(facts{i});
    k = k + 1;
    mwasize(k) = m;
    if k > 1
      if nofac ~= n
        error('All orders must have the same number of factors')
      end
    else
      nofac = n;
    end
  end
end
mwa = zeros(prod(mwasize),nofac);

for j = 1:nofac
  if lo ~= 1
    mwvect = facts{1}(:,j);
    for i = 2:order
	  if lo ~= i
        %mwvect = kron(facts{i}(:,j),mwvect);
		mwvect = mwvect*facts{i}(:,j)';
		mwvect = mwvect(:);
	  end
    end
  elseif lo == 1
    mwvect = facts{2}(:,j);
	for i = 3:order
      %mwvect = kron(facts{i}(:,j),mwvect);
	  mwvect = mwvect*facts{i}(:,j)';
	  mwvect = mwvect(:);
	end
  end
  mwa(:,j) = mwvect;
end
% If vect isn't one, sum up the results of the factors and reshape
if vect ~= 1
  mwa = sum(mwa,2);
  mwa = reshape(mwa,mwasize);
end


function [t,p,Mean,Fit,RelFit] = pcanipals(X,F,cent);

% NIPALS-PCA WITH MISSING ELEMENTS
% 20-6-1999
%
% Calculates a NIPALS PCA model. Missing elements 
% are denoted NaN. The solution is nested
% 
% Comparison for data with missing elements
% NIPALS : Nested    , not least squares, not orthogonal solutoin
% LSPCA  : Non nested, least squares    , orthogonal solution
% 
% I/O
% [t,p,Mean,Fit,RelFit] = pcanipals(X,F,cent);
% 
% X   : Data with missing elements set to NaN
% F   : Number of componets
% cent: One if centering is to be included, else zero
% 
% Copyright
% Rasmus Bro
% KVL 1999
%
%

[I,J]=size(X);
if any(sum(isnan(X))==I)|any(sum(isnan(X)')==J)
   error(' One column or row only contains missing')
end

Xorig      = X;
Miss       = isnan(X);
NotMiss    = ~isnan(X);

ssX    = sum(X(find(NotMiss)).^2);

Mean   = zeros(1,J);
if cent
   Mean    = nanmean(X);
end
X      = X - ones(I,1)*Mean;

t=[];
p=[];

for f=1:F
   Fit    = 3;
   OldFit = 6;
   it     = 0;
   
   T      = nanmean(X')';
   P      = nanmean(X)';
   Fit    = 2;
   FitOld = 3;
   
   while abs(Fit-FitOld)/FitOld>1e-7 & it < 1000;
      FitOld  = Fit;
      it      = it +1;
      
      for j = 1:J
         id=find(NotMiss(:,j));
         P(j) = T(id)'*X(id,j)/(T(id)'*T(id));
      end
      P = P/norm(P);
      
      for i = 1:I
         id=find(NotMiss(i,:));
         T(i) = P(id)'*X(i,id)'/(P(id)'*P(id));
      end
      
      Fit = X-T*P';
      Fit = sum(Fit(find(NotMiss)).^2);
   end
   t = [t T];
   p = [p P];
   X = X - T*P';

end

Model   = t*p' + ones(I,1)*Mean;
Fit     = sum(sum( (Xorig(find(NotMiss)) - Model(find(NotMiss))).^2));
RelFit  = 100*(1-Fit/ssX);

function y = nanmean(x)
if isempty(x) % Check for empty input.
    y = NaN;
    return
end
nans = isnan(x);
i = find(nans);
x(i) = zeros(size(i));

if min(size(x))==1,
  count = length(x)-sum(nans);
else
  count = size(x,1)-sum(nans);
end
i = find(count==0);
count(i) = ones(size(i));
y = sum(x)./count;
y(i) = i + NaN;
```
