## Supplementary File S1 for "Robust latent-variable interpretation of *in vivo* regression models by nested resampling": npred.m

```
function [ypred,T,ssX,Xres]=npred(X,Fac,Xfactors,Yfactors,Core,B,show)

%NPRED prediction with NPLS model
%
% See also:
% 'npls' 'testreg'
%
%
% Predict Y for a new set of samples using an N-PLS model
%
% [ypred,T,ssx,Xres] = npred(X,Fac,Xfactors,Yfactors,Core,B,show);
%
% INPUT
% X        The array to be predicted
% Fac      Number of factors to use in prediction
% Xfactors Parameters of the calibration model of X (incl. scores) in a cell array
% Yfactors Parameters of the calibration model of Y (incl. scores) in a cell array
% Core     Core array used for calculating the model of X
% B        Regression matrix of calibration model
%
% OUTPUT
% ypred    the predictions of y
% T        is the scores of the new samples
% ssX      sum-of-squares of x residuals
%          ssX(1,1)  sum-of-squares of X
%          ssX(2,1)  sum-of-squares of residuals
%          ssX(1,2)  percentage variation explained after 0 component (=0)
%          ssX(2,2)  percentage variation explained after Fac component
%
% Xres     is the residuals of X
%


% Copyright (C) 1995-2006  Rasmus Bro & Claus Andersson
% Copenhagen University, DK-1958 Frederiksberg, Denmark,


% Convert to old notation

Dimx = size(X);
X = reshape(X,Dimx(1),prod(Dimx(2:end)));

DimX = size(Xfactors{1},1);
for j = 2:length(Xfactors)
   DimX(j) = size(Xfactors{j},1);
end
DimY = size(Yfactors{1},1);
for j = 2:length(Yfactors)
   DimY(j) = size(Yfactors{j},1);
end


if nargin==0
  disp(' ')
  disp(' NPRED')
  disp('[ypred,T,ssx,Xres] = npred(X,Fac,Xfactors,Yfactors,Core,B,show);')
  disp(' ')
  return
end

ord=length(DimX);


if ~exist('show')==1
  show=1;
end

maxit=20;


[I,not]=size(X);
if any(isnan(X(:)))
  miss=1-isnan(X);
  Missing=1;
else
  Missing=0;
end

Xres=X;
T=zeros(I,Fac);

W = [];
for f = 1:Fac
   w = Xfactors{end}(:,f);
   for o = length(DimX)-1:-1:2
      w = kron(w,Xfactors{o}(:,f));
   end
   W = [W w];
end
Q = [];
for f = 1:Fac
   q = Yfactors{end}(:,f);
   for o = length(DimY)-1:-1:2
      q = kron(q,Yfactors{o}(:,f));
   end
   Q = [Q q];
end


for f=1:Fac
   if Missing
      for i=1:I
         m = find(miss(i,:));
         T(i,f)=Xres(i,m)*W(m,f)/(W(m,f)'*W(m,f));
      end
   else
      T(:,f)=Xres*W(:,f);
   end
   if f==Fac
      Wkron = Xfactors{end}(:,1:Fac);
      for o = length(DimX)-1:-1:2
         Wkron = kron(Wkron,Xfactors{o}(:,1:Fac));
      end
      Xres=Xres-T(:,1:Fac)*reshape(Core{Fac},Fac,Fac^(length(DimX)-1))*Wkron';
   end
end

ypred=T*B(1:Fac,1:Fac)*Q(:,1:Fac)';

ssx=sum(Xres(find(~isnan(Xres))).^2);
ssX=sum(X(find(~isnan(Xres))).^2);
ssX=[ssX 0;ssx 100*(1-ssx/ssX)];

Xres = reshape(Xres,[size(X,1) DimX(2:end)]);
```
