## Supplementary File S1 for "Robust latent-variable interpretation of *in vivo* regression models by nested resampling": nprocess.m

```
function [Xnew,mX,sX]=nprocess(X,Cent,Scal,mX,sX,reverseProc,show,usemse);

%NPROCESS pre and postprocessing of multiway arrays
%
%
% CENTERING AND SCALING OF N-WAY ARRAYS
%
% This m-file works in two ways
%    I.  Calculate center and scale parameters and preprocess data
%    II. Use given center and scale parameters for preprocessing data
% 
% %%% I. Calculate center and scale parameters %%%
% 
%     [Xnew,Means,Scales]=nprocess(X,Cent,Scal);
% 
%     INPUT
%     X       Data array
%     Cent    is binary row vector with as many elements as DimX.
%             If Cent(i)=1 the centering across the i'th mode is performed
%             I.e cnt = [1 0 1] means centering across mode one and three.
%     Scal    is defined likewise. Scal(i)=1, means scaling to standard  
%             deviation one within the i'th mode
% 
%     OUTPUT
%     Xnew    The preprocessed data
%     mX      Sparse vector holding the mean-values 
%     sX      Sparse vector holding the scales
%
% %%% II. Use given center and scale parameters %%%
% 
%     Xnew=nprocess(X,Cent,Scal,mX,sX,reverse);
% 
%     INPUT
%     X       Data array
%     Cent    is binary row vector with as many elements as DimX.
%             If Cent(i)=1 the centering across the i'th mode is performed
%             I.e Cent = [1 0 1] means centering across mode one and three.
%     Scal    is defined likewise. Scal(i)=1, means scaling to standard  
%             deviation one within the i'th mode
%     mX      Sparse vector holding the mean-values 
%     sX      Sparse vector holding the scales
%     reverse Optional input
%             if reverse = 1 normal preprocessing is performed (default)
%             if reverse = -1 inverse (post-)processing is performed
%
%     OUTPUT
%     Xnew    The preprocessed data
%
% For convenience this m-file does not use iterative 
% preprocessing, which is necessary for some combinations of scaling
% and centering. Instead the algorithm first standardizes the modes
% successively and afterwards centers. The prior standardization ensures
% that the individual variables are on similar scale (this might be slightly
% disturbed upon centering - unlike for two-way data).
%
% The full I/O for nprocess is
% [Xnew,mX,sX]=nprocess(X,Cent,Scal,mX,sX,reverse,show,usemse);
% where show set to zero avoids screen output and where usemse set 
% to one uses RMSE instead of STD for scaling (more appropriate 
% in some settings)

% Copyright, 1998 - 
%
% Rasmus Bro
% Chemometrics Group, Food Technology
% Department of Food and Dairy Science
% Royal Veterinary and Agricultutal University
% Rolighedsvej 30, DK-1958 Frederiksberg, Denmark
% Phone  +45 35283296
% Fax    +45 35283245
%


% $ Version 1.03 $ Date 6. May 1998 $ Drastic error in finding scale parameters corrected $ Not compiled $
% $ Version 1.031 $ Date 25. January 2000 $ Error in scaling part $ Not compiled $
% $ Version 1.032 $ Date 28. January 2000 $ Minor bug$ Not compiled $
% $ Version 1.033 $ Date 14. April 2001 $ Incorrect backscaling fixed.
% $ Version 2.00 $ May 2001 $ Changed to array notation $ RB $ Not compiled $
% $ Version 2.00 $ May 2001 $ rewritten by Giorgio Tomasi $ RB $ Not compiled $
% $ Version 2.01 $ Feb 2002 $ Fixed errors occuring with one-slab inputs $ RB $ Not compiled $
% $ Version 2.02 $ Oct 2003 $ Added possibility for sclaing with RMSE $ RB $ Not compiled $


% $ Version 1.03 $ Date 6. May 1998 $ Drastic error in finding scale parameters corrected $ Not compiled $

% Copyright (C) 1995-2006  Rasmus Bro & Claus Andersson
% Copenhagen University, DK-1958 Frederiksberg, Denmark,
%

%
% CENTERING AND SCALING OF N-WAY ARRAYS
%
% This m-file works in two ways
%    I.  Calculate center and scale parameters and preprocess data
%    II. Use given center and scale parameters for preprocessing data
% 
% %%% I. Calculate center and scale parameters %%%
% 
%     [Xnew,Means,Scales]=nprocess(X,DimX,Cent,Scal);
% 
%     INPUT
%     X       Data array
%     DimX    Size of X
%     Cent    is binary row vector with as many elements as DimX.
%             If Cent(i)=1 the centering across the i'th mode is performed
%             I.e cnt = [1 0 1] means centering across mode one and three.
%     Scal    is defined likewise. Scal(i)=1, means scaling to standard  
%             deviation one within the i'th mode
% 
%     OUTPUT
%     Xnew    The preprocessed data
%     mX      Sparse vector holding the mean-values 
%     sX      Sparse vector holding the scales
%
% %%% II. Use given center and scale parameters %%%
% 
%     Xnew=nprocess(X,DimX,Cent,Scal,mX,sX);
% 
%     INPUT
%     X       Data array
%     DimX    Size of X
%     Cent    is binary row vector with as many elements as DimX.
%             If Cent(i)=1 the centering across the i'th mode is performed
%             I.e Cent = [1 0 1] means centering across mode one and three.
%     Scal    is defined likewise. Scal(i)=1, means scaling to standard  
%             deviation one within the i'th mode
%     mX      Sparse vector holding the mean-values 
%     sX      Sparse vector holding the scales
%     reverse Optional input
%             if reverse = 1 normal preprocessing is performed (default)
%             if reverse = -1 inverse (post-)processing is performed
%
%     OUTPUT
%     Xnew    The preprocessed data
%
% For convenience this m-file does not use iterative 
% preprocessing, which is necessary for some combinations of scaling
% and centering. Instead the algorithm first standardizes the modes
% successively and afterwards centers. The prior standardization ensures
% that the individual variables are on similar scale (this might be slightly
% disturbed upon centering - unlike for two-way data).
%
%	Copyright
%	Rasmus Bro 1997
%	Denmark
%
ord  = ndims(X);
DimX = size(X);
Xnew = X;

if nargin<3
   error(' Three input arguments must be given')
end

if nargin==4
   error(' You must input both mX and sX even if you are only doing centering')
end

if nargin<8
  usemse=0;
end

if ~exist('mX','var')
   mX = [];
end
if ~exist('sX','var')
   sX = [];
end

MODE = isa(mX,'cell')&isa(sX,'cell');

if ~(exist('show')==1)
   show=1;
end

if ~exist('reverseProc')==1
   reverseProc=1;
end

if ~any([1 -1]==reverseProc)
   error( 'The input <<reverse>> must be one or minus one')
end

if show~=-1
   if ~MODE
      disp(' Calculating mean and scale and processing data')
   else
      if reverseProc==1
         disp(' Using given mean and scale values for preprocessing data')
      elseif reverseProc==-1
         disp(' Using given mean and scale values for postprocessing data')
      end
   end
end

for i=1:ndims(X)
   Inds{i} = ones(size(Xnew,i),1);
end
Indm = repmat({':'},ndims(Xnew) - 1,1); 

out=0;
if ~MODE
   mX = cell(ord,1);
   sX = cell(ord,1);
end
Ord2Patch = [2,1;1,2];
if reverseProc == 1
   %Standardize
   for j = ord:-1:1
      o = [j 1:j-1 j+1:ord];
      if Scal(j)
         if show~=-1
            disp([' Scaling mode ',num2str(j)])
         end
         if ~MODE
           if ~usemse
             sX{j} = (stdnan(nshape(Xnew,j)')').^-1;
           else
             sX{j} = (rmsenan(nshape(Xnew,j)')').^-1;
           end
         end
         Xnew = Xnew.*ipermute(sX{j}(:,Inds{o(2:end)}),o);
      end
   end
   %Center
   for j = ord:-1:1
      o = [1:j-1 j+1:ord,j];
      if Cent(j)
         if show~=-1
            if ~MODE
               disp([' Centering mode ',num2str(j)])
            else
               disp([' Subtracting off-sets in mode ',num2str(j)])
            end
         end
         if ~MODE
            if ord ~= 2
               mmm = nshape(Xnew,j);
               if min(size(mmm))==1
                  mmm = mmm;
               else
                  mmm = missmean(mmm);
               end
               mX{j} = reshape(mmm,DimX(o(1:end-1)));
            else
               mX{j} = reshape(missmean(nshape(Xnew,j)),DimX(o(1)),1);
            end
         end
         size(Xnew) 
         size(ipermute(mX{j}(Indm{:},Inds{j}),o))
         Xnew = Xnew - ipermute(mX{j}(Indm{:},Inds{j}),o);
      end
   end
   
else

   %Center
   for j = 1:ord
      
      if Cent(j)
         if show~=-1
            disp([' Adding off-sets in mode ',num2str(j)])
         end
         Xnew = Xnew + ipermute(mX{j}(Indm{:},Inds{j}),[1:j-1 j+1:ord,j]);
      end
   end
   %Standardize
   for j = 1:ord
      o = [1:j-1 j+1:ord];
      if Scal(j)
         if show~=-1
            disp([' Rescaling back to original domain in mode ',num2str(j)])
         end
         Xnew = Xnew ./ ipermute(sX{j}(:,Inds{o}),[j o]);
      end
   end
   
end

function st=rmsenan(X);

%RMSENAN estimate RMSE with NaN's
%
% Estimates the RMSE of each column of X
% when there are NaN's in X.
%
% Columns with only NaN's get a standard deviation of zero


% $ Version 1.02 $ Date 28. July 1998 $ Not compiled $
% 
%
% Copyright, 1998 - 
% This M-file and the code in it belongs to the holder of the
% copyrights and is made public under the following constraints:
% It must not be changed or modified and code cannot be added.
% The file must be regarded as read-only. Furthermore, the
% code can not be made part of anything but the 'N-way Toolbox'.
% In case of doubt, contact the holder of the copyrights.
%
% Rasmus Bro
% Chemometrics Group, Food Technology
% Department of Food and Dairy Science
% Royal Veterinary and Agricultutal University
% Rolighedsvej 30, DK-1958 Frederiksberg, Denmark
%

[I,J]=size(X);

st=[];
for j=1:J
  id=find(~isnan(X(:,j)));
  if length(id)
    st=[st sqrt(mean(X(id,j).^2))];
  else
    st=[st 0];
  end
end
```
