## Supplementary File S1 for "Robust latent-variable interpretation of *in vivo* regression models by nested resampling": nsetdiff.m

```
function [v]=nsetdiff(A,B);
%NSETDIFF
%
%[v]=nsetdiff(A,B);
%Slow setdiff by CA, 1998

% Copyright (C) 1995-2006  Rasmus Bro & Claus Andersson
% Copenhagen University, DK-1958 Frederiksberg, Denmark,
AS=sort(A);
len_AS=length(AS);
for i=1:len_AS-1,
    for j=i+1:len_AS,
        if AS(i)==AS(j),
            AS(i)=NaN;
        end;
    end
end;
I=find(isnan(AS));
if ~isempty(I)
    AS(I)=[];
end;


BS=sort(B);
len_BS=length(BS);
for i=1:len_BS-1,
    for j=i+1:len_BS,
        if BS(i)==BS(j),
            BS(i)=NaN;
        end;
    end
end;
I=find(isnan(BS));
if ~isempty(I)
    BS(I)=[];
end;


len_AS=length(AS);
len_BS=length(BS);
if len_AS >= len_BS
    for i=1:len_AS,
        for j=1:len_BS,
            if AS(i)==BS(j),
                AS(i)=NaN;
            end;
        end;
    end;
    I=find(isnan(AS));
    if ~isempty(I)
        AS(I)=[];
    end;
    v=AS;
else
    for i=1:len_BS,
        for j=1:len_AS,
            if BS(i)==AS(j),
                BS(i)=NaN;
            end;
        end;
    end;
    I=find(isnan(BS));
    if ~isempty(I)
        BS(I)=[];
    end;
    v=BS;
end;
```
