## Supplementary File S1 for "Robust latent-variable interpretation of *in vivo* regression models by nested resampling": nshape.m

```
function varargout=nshape(X,f);

%NSHAPE rearrange a multi-way array
%
% [Xf,DimXf] = nshape(X,f);
%
% Refolds an N-way array so that Xf is X with index
% f as row-index, and the remaining in succesive order. For an 
% I x J x K x L four-way array this means X1 is I x JKL, X2 is
% J x IKL, X3 is K x IJL, and X4 is L x IJK
%
%
%    K  _______             
%      /      /|           1      J     2·J    J·K
%     /______/ |         1  _____________________
%    |      |  |           |      |      |      |
%    |      | /    -->     |      |      |      |        f = (Mode) 1 (same as original array)
% I  |______|/          I  |______|______|______|
%           J
%
%                          1      I     2·I    K·I
%                        1  _____________________
%                          |      |      |      |
%                  -->     |      |      |      |        f = (Mode) 2
%                        J |______|______|______|
%
%  
%                          1      I     2·I    I·J
%                        1  _____________________
%                          |      |      |      |
%                  -->     |      |      |      |        f = (Mode) 3
%                        K |______|______|______|
%
%
% f can also indicate the order (meaning the sequence) of the modes
% [Xf,DimXf] = nshape(X,[3 2 1 4]);
% will return Xf as K x JIL
%
% If the last input is not given all rearrangements are given.
% For a fourway array this would read
% [X1,X2,X3,X4]=nshape(X);
%

% Copyright (C) 1995-2006  Rasmus Bro & Claus Andersson
% Copenhagen University, DK-1958 Frederiksberg, Denmark,


% $ Version 1.03 $ Date 18. July 1999 $ Not compiled $
% $ Version 1.031 $ Date 18. July 1999 $ Error in help figure and now outputs new DimX $ Not compiled $
% $ Version 2.0 $ Jan 2002 $ Not compiled $ Improved speed and added permute functionality Giorgio Tomasi


ord       = ndims(X);
DimX      = size(X);
varargout = [];
if nargin < 2
   f     = 0;
   do_it = ones(1,nargout);
else
   if length(f) == 1
      do_it = [1:ord] == f;
   end
end
if length(f) == 1
   for i = 1:ord
      if do_it(i)
         varargout{end+1} = reshape(permute(X,[i 1:i-1 i+1:ord]),DimX(i),prod(DimX([1:i-1 i+1:ord])));
      end
   end
   if nargin == 2
      varargout{2} = [DimX(f) DimX([1:f-1 f+1:ord])];
   end
else
   if length(f)==ord
      DimX         = DimX(f);
      varargout{1} = reshape(permute(X,f),DimX(1),prod(DimX(2:end)));
      if nargin == 2
         varargout{2} = DimX;
      end
   else
      error(['f can either be the dimension to be put first or',char(10),...
            'a vector containing the new order of the dimensions']);
   end
end
```
