## Supplementary File S1 for "Robust latent-variable interpretation of *in vivo* regression models by nested resampling": ntimes.m

```
function product = ntimes(X,Y,modeX,modeY);


%NTIMES Array multiplication
%
% X*Y is the array/matrix product of X and Y. These are multiplied across the 
% modeX mode/dimension of X and modeY mode/dimension of Y. The number of levels of X in modeX
% must equal the number of levels in modeY of Y.
%
% The product will be an array of order two less than the sum of the orders of A and B
% and thus works as a straightforward extension of the matrix product. The order
% of the modes in the product are such that the X-modes come first and then the
% Y modes. 
%
% E.g. if X is IxJ and Y is KxL and I equals L, then 
% ntimes(X,Y,1,2) will yield a JxK matrix equivalent to the matrix product
% X'*Y'. If X is IxJxK and Y is LxMxNxO with J = N then 
% ntimes(X,Y,2,3) yields a product of size IxKxLxMxO
% 
%I/O: product = NTIMES(X,Y,modeX,modeY) 
% 
%See also TIMES,MTIMES.

% Copyright (C) 1995-2006  Rasmus Bro & Claus Andersson
% Copenhagen University, DK-1958 Frederiksberg, Denmark,
%
orderX = ndims(X);
orderY = ndims(Y);
xsize  = size(X);
ysize  = size(Y);

X = permute(X,[modeX 1:modeX-1 modeX+1:orderX]);
Y = permute(Y,[modeY 1:modeY-1 modeY+1:orderY]);
xsize2  = size(X);
ysize2  = size(Y);

if size(X,1)~=size(Y,1)
   error(' The number of levels must be the same in the mode across which multiplications is performed')
end

%multiply the matricized arrays
product = reshape(X,xsize2(1),prod(xsize2(2:end)))'*reshape(Y,ysize2(1),prod(ysize2(2:end)));
%reshape to three-way structure
product = reshape(product,[xsize2(2:end) ysize2(2:end)]);
```
