## Supplementary File S1 for "Robust latent-variable interpretation of *in vivo* regression models by nested resampling": parademo.m

```
%parademo.m
%
% REQUIRES THE DATA SETS !!!!!
%

% Copyright (C) 1995-2006  Rasmus Bro & Claus Andersson
% Copenhagen University, DK-1958 Frederiksberg, Denmark,
%


close all
clear all
echo off
home
echo on
load claus;
echo off

disp(' ')
disp(' Fluorescence measurements ideally follow the trilinear PARAFAC')
disp(' model. We will use a simple data set of 5 mixtures of three')
disp(' amino-acids (trp,phe & tyr) to show how the pure spectra and')
disp(' concentrations can be found with parafac')
disp(' ')
disp(' First lets look at the raw data:')
disp(' Press any key to continue')
pause

echo on
figure(1);
for i=1:5,
   subplot(3,2,i)
   sample = squeeze(X(i,:,:));
   mesh(ExAx,EmAx,sample);
   title(['Raw data - sample ',num2str(i)]);
   xlabel('Excitation [nm]')
   ylabel('Emission [nm]')
   axis([ExAx(1) ExAx(end) EmAx(1) EmAx(end) 0 1000]);
   grid on
   drawnow
end;
echo off

disp(' ')
disp(' Press any key to continue')
pause
close all
home

disp(' ')
disp(' The data will be slightly reduced to save computation time. This is done')
disp(' by reducing the 2. and 3. emission and excitation mode:')
disp(' ')

echo on
X = X(:,1:5:end,1:2:end);
EmAx = EmAx(1:5:end);
ExAx = ExAx(1:5:end);
size(X)
echo off

disp(' ')
disp(' Press any key to continue')
pause
close all
home


disp(' ')
disp(' PARAFAC may be fitted to the data, but in order to investigate how')
disp(' many components are needed we will use the tool pftest to get an')
disp(' indication. We use 1 to 5 components because we assume that appr.')
disp(' 3 are reasonable but want to check that. We fit models from 1 to 5 ')
disp(' components and fit each three times. This way we can check that the')
disp(' is the same every time we fit, say a four-component model. If it is')
disp(' it is an indication that we have used too many components or that the')
disp('  data are difficult to fit')
disp(' ')
disp(' This process will take some time, but afterwards a plot is produced')
disp(' which is often very instructive to look at')
disp(' ')
disp(' ')
disp(' Press any key to continue')
disp(' ')
pause
close all
home

echo on
[ssX,Corco] = pftest(3,X,5,[0 0 0 0 NaN]);
echo off 
disp(' ')
disp(' Indeed, the fit values seem to indicate that three components are ')
disp(' suitable, whereas the core consistency seems to point to a possible')
disp(' fourth component. This fourth component must be weak though considering')
disp(' the low increase in fit. If looking into the 3- and 4-component models')
disp(' it is realized that the fourth component is reflecting Rayleigh scatter.')
disp(' The problem could have been avoided if we remove emission below ')
disp(' excitation (by setting these elements to NaN), but this is not pursued')
disp(' here. Instead, we will fit a three-component parafac model.')
disp(' ')
disp(' We will turn the plotting on, so that a graphical output is produced')
disp(' ') 
disp(' Press any key to continue')
pause
close all
home

echo on
model = parafac(X,3,[0 0 2]);
echo off 

disp(' ') 
disp(' Press any key to continue')
pause
close all
home

disp(' Using the function fac2let, we can get scores and loading out')
disp(' ')
echo on
[A,B,C] = fac2let(model);
echo off
disp(' ')
disp(' Scores')
disp(A)
disp(' ')
disp(' Concentrations')
disp(y)
disp(' ')
disp(readme)
disp(' ')

disp(' Scrutinize the scores and compare to the concentrations to see')
disp(' the relation between the two. Each score correspond to a specific')
disp(' component up to scale and permutation')
disp(' ')
disp(' END OF  PARADEMO')
```
