## Supplementary File S1 for "Robust latent-variable interpretation of *in vivo* regression models by nested resampling": parafac.m

```
function [Factors,it,err,corcondia]=parafac(X,Fac,Options,const,OldLoad,FixMode,Weights);

% PARAFAC multiway parafac model
%
% See also:
% 'npls' 'tucker' 'dtld' 'gram'
%
%
%     ___________________________________________________
%
%                  THE PARAFAC MODEL
%     ___________________________________________________
%
% [Factors,it,err,corcondia] = parafac(X,Fac,Options,const,OldLoad,FixMode,Weights);
%
% or skipping optional in/outputs
%
% Factors = parafac(X,Fac);
%
% Algorithm for computing an N-way PARAFAC model. Optionally
% constraints can be put on individual modes for obtaining
% orthogonal, nonnegative, or unimodal solutions. The algorithm
% also handles missing data. For details of PARAFAC
% modeling see R. Bro, Chemom. Intell. Lab. Syst., 1997.
%
% Several possibilities exist for speeding up the algorithm.
% Compressing has been incorporated, so that large arrays can be
% compressed by using Tucker (see Bro & Andersson, Chemom.
% Intell. Lab. Syst., 1998).
% Another acceleration method incorporated here is to
% extrapolate the individual loading elements a number of
% iterations ahead after a specified number of iterations.
%
% A temporary MAT-file called TEMP.mat is saved for every
% 50 iterations. IF the computer breaks down or the model
% seems to be good enough, one can break the program and
% load the last saved estimate. The loadings in TEMP.MAT
% are given a cell array as described below and can be
% converted to A, B, C etc. by FAC2LET.M typing
% [A,B,C]=fac2let(Factors,size(X));
%
% All loading vectors except in first mode are normalized,
% so that all variance is kept in the first mode (as is
% common in two-way PCA). The components are arranged as
% in PCA. After iterating, the most important component is
% made the first component etc.
%
%
%
% ----------------------INPUT---------------------
%
% X          X is the input array, which can be from three- to N-way (also
%            twoway if the third mode is interpreted as a onedimensional
%            mode).
%
% Fac        No of factors/components sought.
%
%
% ----------------OPTIONAL INPUT---------------------
%
% Options    Optional parameters. If not given or set to zero or [],
%            defaults will be used. If you want Options(5) to be 2 and
%            not change others, simply write Options(5)=2. Even if Options
%            hasn't been defined Options will contain zeros except its
%            fifth element.
%
%            Options(1) - Convergence criterion
%            The relative change in fit for which the algorithm stops.
%            Standard is 1e-6, but difficult data might require a lower value.
%
%            Options(2) - Initialization method
%            This option is ignored if PARAFAC is started with old values.
%            If no default values are given the default Options(2) is 0.
%            The advantage of using DTLD or SVD for initialization is that
%            they often provide good starting values. However, since the
%            initial values are then fixed, repeating the fitting will give
%            the exact same solution. Therefore it is not possible to substantiate
%            if a local minimum has been reached. To avoid that use an initialization
%            based on random values (2).
%
%            0  = fit using DTLD/GRAM for initialization (default if
%                                 three-way and no missing and if sizes are
%                                 largere than number of factors at least
%                                 in two modes)
%            1  = fit using SVD vectors for initialization (default if higher than three-way or missing)
%            2  = fit using random orthogonalized values for initialization
%            10 = fit using the best-fitting models of several models
%            fitted using a few iterations
%
%            Options(3) - Plotting options
%            0 = no plots
%            1 = produces several graphical outputs
%            2 = produces several graphical outputs (loadings also shown during iterations)
%            3 = as 2 but no core consistency check (very slow for large arrays and/or many components)
%
%            Options(4) - Scaling
%            0 or 1 = default scaling (columns in mode one carry the variance)
%            2      = no scaling applied (hence fixed elements will not be modified
%
%            Options(5) - How often to show fit
%            Determines how often the deviation between the model and the data
%            is shown. This is helpful for adjusting the output to the number
%            of iterations. Default is 10. If showfit is set to NaN, almost no
%            outputs are given
%
%            Options(6) - Maximal number of iterations
%            Maximal number of iterations allowed. Default is 2500.
%
% const      A vector telling type of constraints put on the loadings of the
%            different modes. Same size as DimX but the i'th element tells
%            what constraint is on that mode.
%            0 => no constraint,
%            1 => orthogonality
%            2 => nonnegativity
%            3 => unimodality (and nonnegativitiy)
%            4 => L1 fitting (will be imposed in all modes)
%            5 => L1 fitting and nonnegativity (will be imposed in all modes)
%            If const is not defined, no constraints are used.
%            For no constraints in a threeway problem const = [0 0 0]
%
% OldLoad    If initial guess of the loadings is available. OldLoad should be
%            given a cell array where OldLoad{1}=A,OldLoad{2}=B etc.
%
% FixMode    FixMode is a binary vector of same sixe as DimX. If
%            FixMode(i) = 1 => Mode i is fixed (requires old values given)
%            FixMode(i) = 0 => Mode i is not fixed hence estimated
%            Ex.: FixMode = [0 1 1] find the scores of a data set given the loadings.
%            When some modes are fixed, the numbering of the components will
%            also be fixed. Normally components are sorted according to variance
%            as in PCA, but this will not be performed if some modes are fixed.
%
% Weights    If a matrix of the same size as X is given, weighted regression
%            is performed using the weights in the matrix Weights. Statistically
%            the weights will usually contain the inverse error standard
%            deviation of the particular element
%
% ---------------------OUTPUT---------------------
%
% Factors    PARAFAC estimate of loadings in one matrix. For a 3 component
%            solution to a 4 x 3 x 3 array the loadings A, B & C will be
%            stored in a 3 element cell vector:
%            Factors{1}=A,
%            Factors{2}=B
%            Factors{3}=C
%            etc.
%
%            Use FAC2LET.M for converting to "normal" output or simply extract the
%            components as e.g. A = Factors{1};
%
% it         Number of iterations used. Can be helpful for checking if the algorithm
%            has converged or simply hit the maximal number of iterations (default 2500).
%
% err        The fit of the model = the sum of squares of errors (not including missing
%            elements).
%
% Corcondia  Core consistency test. Should ideally be 100%. If significantly below
%            100% the model is not valid
%
%
%
% OTHER STUFF
%
%  Missing values are handled by expectation maximization only. Set all
%  missing values to NaN
%
%  COMMAND LINE (SHORT)
%
%  Factors = parafac(X,Fac);
%

% Copyright (C) 1995-2006  Rasmus Bro & Claus Andersson
% Copenhagen University, DK-1958 Frederiksberg, Denmark,

% $ Version 1.03 $ Date 1. October   1998 $ Not compiled $ Changed sign-convention because of problems with centered data
% $ Version 1.04 $ Date 18. February 1999 $ Not compiled $ Removed auxiliary line
% $ Version 1.06 $ Date 1. December  1999 $ Not compiled $ Fixed bug in low fit error handling
% $ Version 1.07 $ Date 17. January  2000 $ Not compiled $ Fixed bug in nnls handling so that the algorithm is not stopped until nonnegative appear
% $ Version 1.08 $ Date 21. January  2000 $ Not compiled $ Changed init DTLD so that primarily negative loadings are reflected if possible
% $ Version 1.09 $ Date 30. May 2000 $ Not compiled $ changed name noptioPF to noptiopf
% $ Version 2.00 $ May 2001 $ Changed to array notation $ RB $ Not compiled $
% $ Version 2.001 $ June 2001 $ Fixed error in weighted regression $ RB $ Not compiled $
% $ Version 2.002 $ Jan 2002 $ Fixed scaling problem due to non-identifiability of DTLD(QZ) by scaling and normalizing after each iteration $ RB $ Not compiled $
% $ Version 2.003 $ Jan 2002 $ Fixed negative solutions when nonneg imposed $ RB $ Not compiled $
% $ Version 2.004 $ Jan 2002 $ Changed initialization when many components used $ RB $ Not compiled $
% $ Version 2.005 $ Jan 2002 $ Changed absolute fit criterion (approacing eps) into relative sse/ssx$ RB $ Not compiled $
% $ Version 2.006 $ Jan 2002 $ Fixed post-scaling when fixed loadings $ RB $ Not compiled $
% $ Version 2.01 $ Jan 2003 $ Removed corcondia for two-way data (doesn't work) and fixed a bug for data with dimension 2 $ RB $ Not compiled $
% $ Version 2.011 $ feb 2003 $ Added an option (4) for not post scaling components $ RB $ Not compiled $
% $ Version 2.10  $ jan 2004 $ Fixed a plotting error occuring when fitting model to old data $ RB $ Not compiled $
% $ Version 2.11  $ jan 2004 $ Fixed that PCA can be fitted $ RB $ Not compiled $
% $ Version 2.12  $ Jul 2004 $ Fixed initialization bug $ RB $ Not compiled $
% $ Version 2.13 $ Jan 2005 $ Modified sign conventions of scores and loads $ RB $ Not compiled $
% $ Version 2.14 $ Feb 2006 $ Fixed bug in sign-swicth when loadings are fixed $ RB $ Not compiled $
% $ Version 2.15 $ Aug 2007 $ ALS scheme substantially improved using linesearch acc. to Rajih, Comon, Harshman 2007 $ RB $ Not compiled $
% $ Version 2.16 $ Jun 2010 $ Fixed that scaling of component is not done in fixed modes $ RB $ Not compiled $

NumbIteraInitia=20;
%accel_pattern='none';
%accel_pattern='shifted'; % Use every-iteration linesearch
accel_pattern='paired'; % Use every second iteration linesearch
if nargin==0
    disp(' ')
    disp(' ')
    disp(' THE PARAFAC MODEL')
    disp(' ')
    disp(' Type <<help parafac>> for more info')
    disp('  ')
    disp(' [Factors,it,err,Corcondia] = parafac(X,Fac,Options,const,OldLoad,FixMode,Weights);')
    disp(' or short')
    disp(' Factors = parafac(X,Fac);')
    disp(' ')
    disp(' Options=[Crit Init Plot NotUsed ShowFit MaxIt]')
    disp(' ')
    disp(' ')
    disp(' EXAMPLE:')
    disp(' To fit a four-component PARAFAC model to X of size 6 x 2 x 200 x 3 type')
    disp(' Factors=parafac(X,4)')
    disp(' and to obtain the scores and loadings from the output type')
    disp(' [A,B,C,D]=fac2let(Factors);')
    return
elseif nargin<2
    error(' The inputs X, and Fac must be given')
end

DimX = size(X);
X = reshape(X,DimX(1),prod(DimX(2:end)));

nonneg_obeyed = 1; % used to check if noneg is ok

if nargin<3
    load noptiopf
    OptionsDefault=Options;
else
    % Call the current Options OptionsHere and load default to use if some of the current settings should be default
    Options=Options(:);
    I=length(Options);
    if I==0
        Options=zeros(8,1);
    end
    I=length(Options);
    if I<8
        Options=[Options;zeros(8-I,1)];
    end
    OptionsHere=Options;
    load noptiopf
    OptionsDefault=Options;
    Options=OptionsHere;
end

if ~exist('OldLoad')==1
    OldLoad=0;
elseif length(OldLoad)==0
    OldLoad=0;
end

% Convergence criteria
if Options(1,1)==0
    Options(1,1)=OptionsDefault(1,1);
end
crit=Options(1);


% Initialization
if ~any(Options(2))
    Options(2)=OptionsDefault(2);
end
Init=Options(2);

% Interim plotting
Plt=Options(3,1);
if ~any([0 1 2 3]==Plt)
    error(' Options(3,1) - Plotting - not set correct; must be 0,1,2 or 3')
end

if Options(5,1)==0
    Options(5,1)=OptionsDefault(5,1);
end
showfit=Options(5,1);
if isnan(showfit)
    showfit=-1;
end
if showfit<-1|round(showfit)~=showfit
    error(' Options(5,1) - How often to show fit - not set correct; must be positive integer or -1')
end

if Options(6,1)==0
    Options(6,1)=OptionsDefault(6,1);
    maxit=Options(6,1);
elseif Options(6)>0&round(Options(6))==Options(6)
    maxit=Options(6,1);
else
    error(' Options(6,1) - Maximal number of iterations - not set correct; must be positive integer')
end

ShowPhi=0; % Counter. Tuckers congruence coef/Multiple cosine/UUC shown every ShowPhiWhen'th time the fit is shown
ShowPhiWhen=10;
MissConvCrit=1e-4; % Convergence criterion for estimates of missing values
NumberOfInc=0; % Counter for indicating the number of iterations that increased the fit. ALS algorithms ALLWAYS decrease the fit, but using outside knowledge in some sense (approximate equality or iteratively reweighting might cause the algorithm to diverge

% INITIALIZE
if showfit~=-1
    disp(' ')
    disp(' PRELIMINARY')
    disp(' ')
end
ord=length(DimX);

if showfit~=-1
    disp([' A ',num2str(Fac),'-component model will be fitted'])
end

if exist('const')~=1
    const=zeros(size(DimX));
elseif length(const)~=ord
    if length(DimX)==2 & length(const)==3
        const = const(1:2);
    elseif ~iscell(const)
        const=zeros(size(DimX));
        if showfit~=-1
            disp(' Constraints are not given properly')
        end
    end
end

if showfit~=-1
    for i=1:ord
        if ~iscell(const)
            if const(i)==0
                disp([' No constraints on mode ',num2str(i)])
            elseif const(i)==1
                disp([' Orthogonality on mode ',num2str(i)])
            elseif const(i)==2
                disp([' Nonnegativity on mode ',num2str(i)])
            elseif const(i)==3
                disp([' Unimodality on mode ',num2str(i)])
            elseif any(const==4)
                disp([' L1 fitting in mode ',num2str(i)])
            elseif any(const==5)
                disp([' L1 and nonnegativity fitting in mode ',num2str(i)])
            end
        else
            disp([' compressed nonnegativity in mode ',num2str(i)])
        end
    end
end

% Check if orthogonality required on all modes
DoingPCA= 0;
if ~iscell(const)
    if max(max(const))==1
        if min(min(const))==1,
            if length(DimX)>2
                disp(' ')
                disp(' Not possible to orthogonalize all modes in this implementation.')
                error(' Contact the authors for further information')
            else
                const = [1 0]; % It's ok for PCA but do in one mode to get LS and then orthogonalize afterwards
                DoingPCA = 1;
            end
        end
    end
end

if exist('FixMode')==1
    if length(FixMode)~=ord
        FixMode = zeros(1,ord);
    end
else
    FixMode = zeros(1,ord);
end

if showfit~=-1
    if any(FixMode)
        disp([' The loadings of mode : ',num2str(find(FixMode(:)')),' are fixed'])
    end
end
if exist('Weights')~=1
    Weights=[];
end

% Display convergence criterion
if showfit~=-1
    disp([' The convergence criterion is ',num2str(crit)])
end

% Define loading as one ((r1*r2*r3*...*r7)*Fac x 1) vector [A(:);B(:);C(:);...].
% The i'th loading goes from lidx(i,1) to lidx(i,2)
lidx=[1 DimX(1)*Fac];
for i=2:ord
    lidx=[lidx;[lidx(i-1,2)+1 sum(DimX(1:i))*Fac]];
end

% Check if weighted regression required
if size(Weights,1)==size(X,1)&prod(size(Weights))/size(X,1)==size(X,2)
    Weights = reshape(Weights,size(Weights,1),prod(size(Weights))/size(X,1));
    if showfit~=-1
        disp(' Given weights will be used for weighted regression')
    end
    DoWeight=1;
else
    if showfit~=-1
        disp(' No weights given')
    end
    DoWeight=0;
end

% Make idx matrices if missing values
if any(isnan(X(:)))
    MissMeth=1;
else
    MissMeth=0;
end
if MissMeth
    id=sparse(find(isnan(X)));
    idmiss2=sparse(find(~isnan(X)));
    if showfit~=-1
        disp([' ', num2str(100*(length(id)/prod(DimX))),'% missing values']);
        disp(' Expectation maximization will be used for handling missing values')
    end
    SSX=sum(sum(X(idmiss2).^2)); % To be used for evaluating the %var explained
    % If weighting to zero should be used
    % Replace missing with mean values or model estimates initially
    %Chk format ok.
    dimisok = 1;
    if length(OldLoad)==length(DimX)
        for i=1:length(DimX)
            if ~all(size(OldLoad{i})==[DimX(i) Fac])
                dimisok = 0;
            end
        end
    else
        dimisok = 0;
    end
    if dimisok
        model=nmodel(OldLoad);
        model = reshape(model,DimX);
        X(id)=model(id);
    else
        meanX=mean(X(find(~isnan(X))));
        meanX=mean(meanX);
        X(id)=meanX*ones(size(id));
    end
else
    if showfit~=-1
        disp(' No missing values')
    end
    if iscell(const)
        SSX=sum(sum(X.^2)); % To be used for evaluating the %var explained
    elseif ~(any(const==4)|any(const==5))
        SSX=sum(sum(X.^2)); % To be used for evaluating the %var explained
    else
        SSX=sum(abs(X(:)));
    end
end

% Check if weighting is tried used together with unimodality or orthogonality
if ~iscell(const)
    if any(const==3)|any(const==1)
        if DoWeight==1
            disp(' ')
            disp(' Weighting is not possible together with unimodality and orthogonality.')
            disp(' It can be done using majorization, but has not been implemented here')
            disp(' Please contact the authors for further information')
            error
        end
    end
end

% Acceleration
acc=-5;
do_acc=1;   % Do acceleration every do_acc'th time
acc_pow=2;  % Extrapolate to the iteration^(1/acc_pow) ahead
acc_fail=0; % Indicate how many times acceleration have failed
max_fail=4; % Increase acc_pow with one after max_fail failure
if showfit~=-1
    disp(' Line-search acceleration scheme initialized')
end

% Find initial guesses for the loadings if no initial values are given

% Use old loadings
if length(OldLoad)==ord % Use old values
    if showfit~=-1
        disp(' Using old values for initialization')
    end
    Factors=OldLoad;
    % Use DTLD
elseif Init==0
    if min(DimX)>1&ord==3&MissMeth==0
        if sum(DimX<Fac)<2
            if showfit~=-1
                disp(' Using direct trilinear decomposition for initialization')
            end
            try
                [A,B,C]=dtld(reshape(X,DimX),Fac);
            catch
                A = rand(DimX(1),Fac);B = rand(DimX(2),Fac);C = rand(DimX(3),Fac);
            end
        else
            if showfit~=-1
                disp(' Using random values for initialization')
            end
            for i=1:length(DimX)
                Factors{i}=rand(DimX(i),Fac);
            end
            A = Factors{1};B=Factors{2};C = Factors{3};
        end
        A=real(A);B=real(B);C=real(C);
        % Check for signs and reflect if appropriate
        for f=1:Fac
            if sign(sum(A(:,f)))<0
                if sign(sum(B(:,f)))<0
                    B(:,f)=-B(:,f);
                    A(:,f)=-A(:,f);
                elseif sign(sum(C(:,f)))<0
                    C(:,f)=-C(:,f);
                    A(:,f)=-A(:,f);
                end
            end
            if sign(sum(B(:,f)))<0
                if sign(sum(C(:,f)))<0
                    C(:,f)=-C(:,f);
                    B(:,f)=-B(:,f);
                end
            end
        end
        Factors{1}=A;Factors{2}=B;Factors{3}=C;
        
    else
        if showfit~=-1
            disp(' Using singular values for initialization')
        end
        try
            Factors=ini(reshape(X,DimX),Fac,2);
        catch
            Factors=[];
            for i=1:length(DimX);
                l = rand(DimX(i),Fac);
                Factors{i} =l;
            end
            if showfit~=-1
                disp(' Oops sorry - ended up with random instead')
            end
            
        end
    end
    
    % Use SVD
elseif Init==1
    if all(DimX>=Fac)
        if showfit~=-1
            disp(' Using singular values for initialization')
        end
        try
            Factors=ini(reshape(X,DimX),Fac,2);
            catchs
            Factors=[];
            for i=1:length(DimX);
                l = rand(DimX(i),Fac);
                Factors = [Factors;l(:)];
            end
        end
    else
        if showfit~=-1
            disp(' Using random values for initialization')
        end
        for i=1:length(DimX)
            Factors{i}=rand(DimX(i),Fac);
        end
    end
    
    % Use random (orthogonal)
elseif Init==2
    if showfit~=-1
        disp(' Using orthogonal random for initialization')
    end
    Factors=ini(reshape(X,DimX),Fac,1);
    
elseif Init==3
    error(' Initialization option set to three has been changed to 10')
    
    % Use several small ones of the above
elseif Init==10
    if showfit~=-1
        disp(' Using several small runs for initialization')
    end
    Opt=Options;
    Opt(5) = NaN;
    Opt(6) = NumbIteraInitia;
    Opt(2) = 0;
    ERR=[];
    [Factors,it,err] = parafac(reshape(X,DimX),Fac,Opt,const,[],[],Weights);
    ERR = [ERR;err];
    Opt(2) = 1;
    [F,it,Err] = parafac(reshape(X,DimX),Fac,Opt,const,[],[],Weights);
    ERR=[ERR;Err];
    if Err<err
        Factors=F;
        err=Err;
    end
    Opt(2)=2;
    for rep=1:3
        [F,it,Err]=parafac(reshape(X,DimX),Fac,Opt,const,[],[],Weights);
        ERR=[ERR;Err];
        if Err<err
            Factors=F;
            err=Err;
        end
    end
    if showfit~=-1
        disp(' ')
        disp(' Obtained fit-values')
        disp([' Method   Fit'])
        disp([' DTLD     ',num2str(ERR(1))])
        disp([' SVD      ',num2str(ERR(2))])
        disp([' RandOrth ',num2str(ERR(3))])
        disp([' RandOrth ',num2str(ERR(4))])
        disp([' RandOrth ',num2str(ERR(5))])
    end
else
    error(' Problem in PARAFAC initialization - Not set correct')
end

% Check for signs and reflect if appropriate

for f=1:Fac
    for m=1:ord-1
        if sign(sum(Factors{m}(:,f)<0)) & FixMode(m)==0
            contin=1;
            for m2 = m+1:ord
                if contin & FixMode(m2)==0
                    if sign(sum(Factors{m2}(:,f)<0))
                        Factors{m}(:,f)=-Factors{m}(:,f);
                        Factors{m2}(:,f)=-Factors{m2}(:,f);
                        contin=0;
                    end
                end
            end
        end
    end
end
% Convert to old format


if iscell(Factors)
    ff = [];
    for f=1:length(Factors)
        ff=[ff;Factors{f}(:)];
    end
    Factors = ff;
end


% ALTERNATING LEAST SQUARES
err=SSX;
f=2*crit;
it=0;
connew=2;conold=1; % for missing values
ConstraintsNotRight = 0; % Just to ensure that iterations are not stopped if constraints are not yet fully imposed

if showfit~=-1
    disp(' ')
    if iscell(const)
        disp(' Sum-of-Squares   Iterations  Explained')
        disp(' of residuals                 variation')
    elseif any(const==4)|any(const==5)
        disp(' Sum-of-absolute   Iterations  Explained')
        disp(' residuals                     variation')
    else        
        disp(' Sum-of-Squares   Iterations  Explained')
        disp(' of residuals                 variation')
    end
end


while (((f>crit) | (norm(connew-conold)/norm(conold)>MissConvCrit) | ConstraintsNotRight) & it<maxit)|~ nonneg_obeyed
    conold=connew; % for missing values
    it=it+1;
    acc=acc+1;
    
    if strcmp(accel_pattern,'paired') % this chooses Bro standard two-steps version
        if it==1
            startiter=-acc+1;
            disp([' LS paired accel will be done every other time after iter ' ...
                int2str(startiter)])
        end
        
        if acc==do_acc;
            Load_o1=Factors;
        end
        if acc==do_acc+1;
            acc=0;Load_o2=Factors;
            Factors=Load_o1+(Load_o2-Load_o1)*(it^(1/acc_pow));
            % Convert to new format
            clear ff,id1 = 0;
            for i = 1:length(DimX)
                id2 = sum(DimX(1:i).*Fac);ff{i} = reshape(Factors(id1+1:id2),DimX(i),Fac);id1 = id2;
            end
            model=nmodel(ff);
            model = reshape(model,DimX(1),prod(DimX(2:end)));
            %G(it)=err;
            
            if MissMeth
                connew=model(id);
                errX=X-model;                
                if iscell(const)
                    nerr=sum(sum(errX(idmiss2).^2));
                elseif DoWeight==0&~(any(const==4)|any(const==5))
                    nerr=sum(sum(errX(idmiss2).^2));
                elseif any(const==4)|any(const==5)
                    nerr=sum(sum(abs(errX(idmiss2(:)))));
                else
                    nerr=sum(sum((Weights(idmiss2).*errX(idmiss2)).^2));
                end
            else
                if iscell(const)
                    nerr=sum(sum((X-model).^2));
                elseif DoWeight==0&~(any(const==4)|any(const==5))
                    nerr=sum(sum((X-model).^2));
                elseif any(const==4)|any(const==5)
                    nerr=sum(sum(abs(X(:)-model(:))));
                else
                    nerr=sum(sum((X.*Weights-model.*Weights).^2));
                end
            end
            if nerr>err
                acc_fail=acc_fail+1;
                Factors=Load_o2;
                if acc_fail==max_fail,
                    acc_pow=acc_pow+1+1;
                    acc_fail=0;
                    if showfit~=-1
                        disp(' Reducing acceleration');
                    end
                end
            else
                if MissMeth
                    X(id)=model(id);
                end
            end
        end
    elseif strcmp(accel_pattern,'shifted')% this chooses harshman's every-iter way
        if acc==do_acc-1;
            Load_o2=Factors;
            disp([' LS shifted accel will be done every time after iter ' int2str(it)])
        elseif acc==do_acc;
            Load_o1=Load_o2; % shift the one that was latest back to first
            acc=0;Load_o2=Factors; % latest factors become the newer set
            Factors=Load_o1+(Load_o2-Load_o1)*(it^(1/acc_pow));
            % Convert to new format
            clear ff,id1 = 0;
            for i = 1:length(DimX)
                id2 = sum(DimX(1:i).*Fac);ff{i} = reshape(Factors(id1+1:id2),DimX(i),Fac);id1 = id2;
            end
            model=nmodel(ff);
            model = reshape(model,DimX(1),prod(DimX(2:end)));
            %G(it)=err;
            
            if MissMeth
                connew=model(id);
                errX=X-model;
                if iscell(const)
                    nerr=sum(sum(errX(idmiss2).^2));
                elseif (DoWeight==0&~(any(const==4)|any(const==5)))
                    nerr=sum(sum(errX(idmiss2).^2));
                elseif any(const==4)|any(const==5)
                    nerr=sum(sum(abs(errX(idmiss2(:)))));
                else
                    nerr=sum(sum((Weights(idmiss2).*errX(idmiss2)).^2));
                end
            else
                if iscell(const)
                    nerr=sum(sum((X-model).^2));
                elseif(DoWeight==0&~(any(const==4)|any(const==5)))
                    nerr=sum(sum((X-model).^2));
                elseif any(const==4)|any(const==5)
                    nerr=sum(sum(abs(X(:)-model(:))));
                else
                    nerr=sum(sum((X.*Weights-model.*Weights).^2));
                end
            end
            if nerr>err
                acc_fail=acc_fail+1;
                Factors=Load_o2;
                if acc_fail==max_fail,
                    acc_pow=acc_pow+1+1;
                    acc_fail=0;
                    if showfit~=-1
                        disp(' Reducing acceleration');
                    end
                end
            else
                if MissMeth
                    X(id)=model(id);
                end
            end
        end
    elseif strcmp(accel_pattern,'none')
        % do nothing
    else
        error('accel_pattern should be ''shifted'', ''paired'', or ''none''.')
    end
    
    %   if acc==do_acc; %     Load_o1=Factors;    %   end,    %   if,     %   acc==do_acc+1;    %     acc=0;Load_o2=Factors;    %     Factors=Load_o1+(Load_o2-Load_o1)*(it^(1/acc_pow));    %     %,    %     %     Convert to new format,    %     clear ff,id1 = 0;    %     for i,    %     = 1:length(DimX),    %       id2 = sum(DimX(1:i).*Fac);ff{i} = reshape(Factors(id1+1:id2),DimX(i),Fac);id1 = id2;    %     end,    %     model=nmodel(ff);    %     model = reshape(model,DimX(1),prod(DimX(2:end)));    %     if MissMeth,    %       connew=model(id);    %       errX=X-model;    %       if DoWeight==0,    %,    %,    %       nerr=sum(sum(errX(idmiss2).^2));    %       else,    %         nerr=sum(sum((Weights(idmiss2).*errX(idmiss2)).^2));    %       end,    %     else,    %       if DoWeight==0,    %         nerr=sum(sum((X-model).^2));    %       else,    %    %         nerr=sum(sum((X.*Weights-model.*Weights).^2));    %       end,    %     end,    %     if nerr>err,    %       acc_fail=acc_fail+1;    %       Factors=Load_o2;    %       if acc_fail==max_fail,    %         acc_pow=acc_pow+1+1;    %         acc_fail=0;    %         if showfit~=-1,    %,     %         disp(' Reducing acceleration');    %         end,    %,    %         end,    %     else,    %       if MissMeth,    %         X(id)=model(id);    %       end,    %     end,    %   end
    if ~iscell(const)
        if DoWeight==0&~(any(const==4)|any(const==5))
            for ii=ord:-1:1
                if ii==ord;
                    i=1;
                else
                    i=ii+1;
                end
                idd=[i+1:ord 1:i-1];
                l_idx2=lidx(idd,:);
                dimx=DimX(idd);
                if ~FixMode(i)
                    L1=reshape(Factors(l_idx2(1,1):l_idx2(1,2)),dimx(1),Fac);
                    if ord>2
                        L2=reshape(Factors(l_idx2(2,1):l_idx2(2,2)),dimx(2),Fac);
                        Z=krb(L2,L1);
                    else
                        Z = L1;
                    end
                    for j=3:ord-1
                        L1=reshape(Factors(l_idx2(j,1):l_idx2(j,2)),dimx(j),Fac);
                        Z=krb(L1,Z);
                    end
                    ZtZ=Z'*Z;
                    ZtX=Z'*X';
                    OldLoad=reshape(Factors(lidx(i,1):lidx(i,2)),DimX(i),Fac);
                    L=pfls(ZtZ,ZtX,DimX(i),const(i),OldLoad,DoWeight,Weights);
                    %L=(pinv(ZtZ+eye*600)*ZtX)';
                    Factors(lidx(i,1):lidx(i,2))=L(:);
                end
                x=zeros(prod(DimX([1:ii-1 ii+1:ord])),DimX(ii));  % Rotate X so the current last mode is the first
                x(:)=X;
                X=x';
            end
        elseif DoWeight
            for ii=ord:-1:1
                if ii==ord;
                    i=1;
                else
                    i=ii+1;
                end
                idd=[i+1:ord 1:i-1];
                l_idx2=lidx(idd,:);
                dimx=DimX(idd);
                if ~FixMode(i)
                    L1=reshape(Factors(l_idx2(1,1):l_idx2(1,2)),dimx(1),Fac);
                    if ord>2
                        L2=reshape(Factors(l_idx2(2,1):l_idx2(2,2)),dimx(2),Fac);
                        Z=krb(L2,L1);
                    else
                        Z = L1;
                    end
                    for j=3:ord-1
                        L1=reshape(Factors(l_idx2(j,1):l_idx2(j,2)),dimx(j),Fac);
                        Z=krb(L1,Z);
                    end
                    OldLoad=reshape(Factors(lidx(i,1):lidx(i,2)),DimX(i),Fac);
                    L=pfls(Z,X,DimX(i),const(i),OldLoad,DoWeight,Weights);
                    Factors(lidx(i,1):lidx(i,2))=L(:);
                end
                x=zeros(prod(DimX([1:ii-1 ii+1:ord])),DimX(ii));
                x(:)=X;
                X=x';
                x(:)=Weights;
                Weights=x';
            end
        elseif (any(const==4)|any(const==5))
            for ii=ord:-1:1
                if ii==ord;
                    i=1;
                else
                    i=ii+1;
                end
                idd=[i+1:ord 1:i-1];
                l_idx2=lidx(idd,:);
                dimx=DimX(idd);
                if ~FixMode(i)
                    L1=reshape(Factors(l_idx2(1,1):l_idx2(1,2)),dimx(1),Fac);
                    if ord>2
                        L2=reshape(Factors(l_idx2(2,1):l_idx2(2,2)),dimx(2),Fac);
                        Z=krb(L2,L1);
                    else
                        Z = L1;
                    end
                    for j=3:ord-1
                        L1=reshape(Factors(l_idx2(j,1):l_idx2(j,2)),dimx(j),Fac);
                        Z=krb(L1,Z);
                    end
                    OldLoad=reshape(Factors(lidx(i,1):lidx(i,2)),DimX(i),Fac);
                    if any(const==5);
                        L=plae(X',Z,OldLoad',1)';
                    else
                        L=plae(X',Z,OldLoad',0)';
                    end
                    Factors(lidx(i,1):lidx(i,2))=L(:);
                end
                x=zeros(prod(DimX([1:ii-1 ii+1:ord])),DimX(ii));
                x(:)=X;
                X=x';
            end
        end
        
    else
        for ii=ord:-1:1
            if ii==ord;
                i=1;
            else
                i=ii+1;
            end
            idd=[i+1:ord 1:i-1];
            l_idx2=lidx(idd,:);
            dimx=DimX(idd);
            if ~FixMode(i)
                L1=reshape(Factors(l_idx2(1,1):l_idx2(1,2)),dimx(1),Fac);
                if ord>2
                    L2=reshape(Factors(l_idx2(2,1):l_idx2(2,2)),dimx(2),Fac);
                    Z=krb(L2,L1);
                else
                    Z = L1;
                end
                for j=3:ord-1
                    L1=reshape(Factors(l_idx2(j,1):l_idx2(j,2)),dimx(j),Fac);
                    Z=krb(L1,Z);
                end
                OldLoad=reshape(Factors(lidx(i,1):lidx(i,2)),DimX(i),Fac);
                L=compresnonneg(X',Z,OldLoad,const{i},it);
                Factors(lidx(i,1):lidx(i,2))=L(:);
            end
            x=zeros(prod(DimX([1:ii-1 ii+1:ord])),DimX(ii));  % Rotate X so the current last mode is the first
            x(:)=X;
            X=x';
        end
    end
    
    
    
    % POSTPROCES LOADINGS (ALL VARIANCE IN FIRST MODE)
    if ~any(FixMode)
        A=reshape(Factors(lidx(1,1):lidx(1,2)),DimX(1),Fac);
        for i=2:ord
            B=reshape(Factors(lidx(i,1):lidx(i,2)),DimX(i),Fac);
            for ff=1:Fac
                A(:,ff)=A(:,ff)*norm(B(:,ff));
                B(:,ff)=B(:,ff)/norm(B(:,ff));
            end
            Factors(lidx(i,1):lidx(i,2))=B(:);
        end
        Factors(lidx(1,1):lidx(1,2))=A(:);
    end
    % APPLY SIGN CONVENTION IF NO FIXED MODES
    %  FixMode=1
    if ~iscell(const)
    if ~any(FixMode)&~(any(const==2)|any(const==3))
        Sign = ones(1,Fac);
        for i=ord:-1:2
            A=reshape(Factors(lidx(i,1):lidx(i,2)),DimX(i),Fac);
            Sign2=ones(1,Fac);
            for ff=1:Fac
                [out,sig]=max(abs(A(:,ff)));
                Sign(ff) = Sign(ff)*sign(A(sig,ff));
                Sign2(ff) = sign(A(sig,ff));
            end
            A=A*diag(Sign2);
            Factors(lidx(i,1):lidx(i,2))=A(:);
        end
        A=reshape(Factors(lidx(1,1):lidx(1,2)),DimX(1),Fac);
        A=A*diag(Sign);
        Factors(lidx(1,1):lidx(1,2))=A(:);
    end
    end
    % Check if nonneg_obeyed
    if ~iscell(const)
    for i=1:ord
        if const(i)==2|const(i)==3
            A=reshape(Factors(lidx(i,1):lidx(i,2)),DimX(i),Fac);
            if any(A(:))<0
                nonneg_obeyed=0;
            end
        end
    end
    end
    
    % EVALUATE SOFAR
    % Convert to new format
    clear ff,id1 = 0;
    for i = 1:length(DimX)
        id2 = sum(DimX(1:i).*Fac);
        ff{i} = reshape(Factors(id1+1:id2),DimX(i),Fac);
        id1 = id2;
    end
    model=nmodel(ff);
    model = reshape(model,DimX(1),prod(DimX(2:end)));
    if MissMeth  % Missing values present
        connew=model(id);
        X(id)=model(id);
        errold=err;
        errX=X-model;
        if iscell(const)
            err=sum(sum(errX(idmiss2).^2));
        elseif (DoWeight==0&~(any(const==4)|any(const==5)))
            err=sum(sum(errX(idmiss2).^2));
        elseif any(const==4)|any(const==5)
            err=sum(sum(abs(errX(idmiss2(:)))));
        else
            err=sum(sum((Weights(idmiss2).*errX(idmiss2)).^2));
        end
    else
        errold=err;
        if iscell(const)
            err=sum(sum((X(:)-model(:)).^2));
        elseif (DoWeight==0&~(any(const==4)|any(const==5)))
            err=sum(sum((X-model).^2));
        elseif any(const==4)|any(const==5)
            err=sum(sum(abs(X(:)-model(:))));
        else
            err=sum(sum((Weights.*(X-model)).^2));
        end
    end
    if err/SSX<1000*eps, % Getting close to the machine uncertainty => stop
        disp(' WARNING')
        disp(' The misfit is approaching the machine uncertainty')
        disp(' If pure synthetic data is used this is OK, otherwise if the')
        disp(' data elements are very small it might be appropriate ')
        disp(' to multiply the whole array by a large number to increase')
        disp(' numerical stability. This will only change the solution ')
        disp(' by a scaling constant')
        f = 0;
    else
        f=abs((err-errold)/err);
        if f<crit % Convergence: then check that constraints are fulfilled
            if ~iscell(const)
            if any(const==2)|any(const==3) % If nnls or unimodality imposed
                for i=1:ord % Extract the
                    if const(i)==2|const(i)==3 % If nnls or unimodality imposed
                        Loadd = Factors(sum(DimX(1:i-1))*Fac+1:sum(DimX(1:i))*Fac);
                        if any(Loadd<0)
                            ConstraintsNotRight=1;
                        else
                            ConstraintsNotRight=0;
                        end
                    end
                end
            end
            end
        end
    end
    
    if it/showfit-round(it/showfit)==0
        if showfit~=-1,
            ShowPhi=ShowPhi+1;
            if ShowPhi==ShowPhiWhen,
                ShowPhi=0;
                if showfit~=-1,
                    disp(' '),
                    disp('    Tuckers congruence coefficient'),
                    % Convert to new format
                    clear ff,id1 = 0;
                    for i = 1:length(DimX)
                        id2 = sum(DimX(1:i).*Fac);ff{i} = reshape(Factors(id1+1:id2),DimX(i),Fac);id1 = id2;
                    end
                    [phi,out]=ncosine(ff,ff);
                    disp(phi),
                    if MissMeth
                        fprintf(' Change in estim. missing values %12.10f',norm(connew-conold)/norm(conold));
                        disp(' ')
                        disp(' ')
                    end
                    disp(' Sum-of-Squares   Iterations  Explained')
                    disp(' of residuals                 variation')
                end
            end
            if iscell(const)
                PercentExpl=100*(1-(sum(sum((X-model).^2))/SSX));
            elseif (DoWeight==0&~(any(const==4)|any(const==5)))
                PercentExpl=100*(1-err/SSX);
            elseif any(const==4)|any(const==5)
                PercentExpl=100*(1-sum(abs((X(:)-model(:))))/SSX);
            else
                PercentExpl=100*(1-(sum(sum((X-model).^2))/SSX));
            end
            fprintf(' %12.10f       %g        %3.4f    \n',err,it,PercentExpl);
            if Plt==2|Plt==3
                % Convert to new format
                clear ff,id1 = 0;
                for i = 1:length(DimX)
                    id2 = sum(DimX(1:i).*Fac);ff{i} = reshape(Factors(id1+1:id2),DimX(i),Fac);id1 = id2;
                end
                pfplot(reshape(X,DimX),ff,Weights',[0 0 0 0 0 0 0 1]);
                drawnow
            end
        end
    end
    
    % Make safety copy of loadings and initial parameters in temp.mat
    if it/50-round(it/50)==0
        save temp Factors
    end
    
    % JUDGE FIT
    if err>errold
        NumberOfInc=NumberOfInc+1;
    end
    % POSTPROCESS. IF PCA on two-way enforce orth in both modes.
    
end % while f>crit

if DoingPCA
    A=reshape(Factors(lidx(1,1):lidx(1,2)),DimX(1),Fac);
    B=reshape(Factors(lidx(i,1):lidx(i,2)),DimX(i),Fac);
    [u,s,v]=svd(A*B',0);
    A = u(:,1:size(A,2))*s(1:size(A,2),1:size(A,2));
    B = u(:,1:size(B,2));
    Factors = [A(:);B(:)];
end


% CALCULATE TUCKERS CONGRUENCE COEFFICIENT
if showfit~=-1 & DimX(1)>1
    disp(' '),disp('   Tuckers congruence coefficient')
    % Convert to new format
    clear ff,id1 = 0;
    for i = 1:length(DimX)
        id2 = sum(DimX(1:i).*Fac);ff{i} = reshape(Factors(id1+1:id2),DimX(i),Fac);id1 = id2;
    end
    [phi,out]=ncosine(ff,ff);
    disp(phi)
    disp(' ')
    if max(max(abs(phi)-diag(diag(phi))))>.85
        disp(' ')
        disp(' ')
        disp(' WARNING, SOME FACTORS ARE HIGHLY CORRELATED.')
        disp(' ')
        disp(' You could decrease the number of components. If this')
        disp(' does not help, try one of the following')
        disp(' ')
        disp(' - If systematic variation is still present you might')
        disp('   wanna decrease your convergence criterion and run')
        disp('   one more time using the loadings as initial guess.')
        disp(' ')
        disp(' - Or use another preprocessing (check for constant loadings)')
        disp(' ')
        disp(' - Otherwise try orthogonalising some modes,')
        disp(' ')
        disp(' - Or use Tucker3/Tucker2,')
        disp(' ')
        disp(' - Or a PARAFAC with some modes collapsed (if # modes > 3)')
        disp(' ')
    end
end


% SHOW FINAL OUTPUT
if iscell(const)
    PercentExpl=100*(1-sum(sum((X-model).^2))/SSX);
elseif DoWeight==0&~(any(const==4)|any(const==5))
    PercentExpl=100*(1-err/SSX);
elseif any(const==5)
    PercentExpl=100*(1-sum(abs(X(:)-model(:)))/SSX);
else
    PercentExpl=100*(1-sum(sum((X-model).^2))/SSX);
end
if showfit~=-1
    fprintf(' %12.10f       %g        %3.4f \n',err,it,PercentExpl);
    if NumberOfInc>0
        disp([' There were ',num2str(NumberOfInc),' iterations that increased fit']);
    end
end

% POSTPROCES LOADINGS (ALL VARIANCE IN FIRST MODE)
if Options(4)==0|Options(4)==1
    A=reshape(Factors(lidx(1,1):lidx(1,2)),DimX(1),Fac);
    for i=2:ord
        if ~FixMode(i)
            B=reshape(Factors(lidx(i,1):lidx(i,2)),DimX(i),Fac);
            for ff=1:Fac
                A(:,ff)=A(:,ff)*norm(B(:,ff));
                B(:,ff)=B(:,ff)/norm(B(:,ff));
            end
            Factors(lidx(i,1):lidx(i,2))=B(:);
        end
    end
    Factors(lidx(1,1):lidx(1,2))=A(:);
    if showfit~=-1
        disp(' ')
        disp(' Components have been normalized in all but the first mode')
    end
end

% PERMUTE SO COMPONENTS ARE IN ORDER AFTER VARIANCE DESCRIBED (AS IN PCA) IF NO FIXED MODES
if ~any(FixMode)
    A=reshape(Factors(lidx(1,1):lidx(1,2)),DimX(1),Fac);
    [out,order]=sort(diag(A'*A));
    order=flipud(order);
    A=A(:,order);
    Factors(lidx(1,1):lidx(1,2))=A(:);
    for i=2:ord
        B=reshape(Factors(lidx(i,1):lidx(i,2)),DimX(i),Fac);
        B=B(:,order);
        Factors(lidx(i,1):lidx(i,2))=B(:);
    end
    if showfit~=-1
        disp(' Components have been ordered according to contribution')
    end
elseif showfit ~= -1
    disp(' Some modes fixed hence no sorting of components performed')
end


% TOOLS FOR JUDGING SOLUTION
if nargout>3
    x=X;
    if MissMeth
        x(id)=NaN*id;
    end
    % Convert to new format
    clear ff,id1 = 0;
    for i = 1:length(DimX)
        id2 = sum(DimX(1:i).*Fac);ff{i} = reshape(Factors(id1+1:id2),DimX(i),Fac);id1 = id2;
    end
    any(isnan(x(:)))
    any(isnan(ff{1}(:)))
    any(isnan(ff{2}(:)))
    any(isnan(ff{3}(:)))
    corcondia=corcond(reshape(x,DimX),ff,Weights,0);
end

% APPLY SIGN CONVENTION IF NO FIXED MODES
%  FixMode=1
if ~iscell(const)
if ~any(FixMode)&~(any(const==2)|any(const==3))
    Sign = ones(1,Fac);
    for i=ord:-1:2
        A=reshape(Factors(lidx(i,1):lidx(i,2)),DimX(i),Fac);
        Sign2=ones(1,Fac);
        for ff=1:Fac
            [out,sig]=max(abs(A(:,ff)));
            Sign(ff) = Sign(ff)*sign(A(sig,ff));
            Sign2(ff) = sign(A(sig,ff));
        end
        A=A*diag(Sign2);
        Factors(lidx(i,1):lidx(i,2))=A(:);
    end
    A=reshape(Factors(lidx(1,1):lidx(1,2)),DimX(1),Fac);
    A=A*diag(Sign);
    Factors(lidx(1,1):lidx(1,2))=A(:);
    
    %   % Instead of above, do signs so as to make them as "natural" as possible
    %   Factors = signswtch(Factors,reshape(X,DimX));
    %   DIDN't WORK (TOOK AGES FOR 7WAY DATA)
    
    
    if showfit~=-1
        disp(' Components have been reflected according to convention')
    end
end
end

% Convert to new format
clear ff,id1 = 0;
for i = 1:length(DimX)
    id2 = sum(DimX(1:i).*Fac);ff{i} = reshape(Factors(id1+1:id2),DimX(i),Fac);id1 = id2;
end
Factors = ff;


if Plt==1|Plt==2|Plt==3
    %   if Fac<6&Plt~=3&order>2&ord>2
    if Fac<6&Plt~=3&ord>2
        pfplot(reshape(X,DimX),ff,Weights,ones(1,8));
    else
        pfplot(reshape(X,DimX),ff,Weights,[1 1 0 1 1 1 1 1]);
        if ord>2
            disp(' Core consistency plot not shown because it requires large memory')
            disp(' It can be made writing pfplot(X,Factors,[Weights],[0 0 1 0 0 0 0 0]');
        else
            disp(' Core consistency not applicable for two-way data')
        end
    end
end

% Show which criterion stopped the algorithm
if showfit~=-1
    if ((f<crit) & (norm(connew-conold)/norm(conold)<MissConvCrit))
        disp(' The algorithm converged')
    elseif it==maxit
        disp(' The algorithm did not converge but stopped because the')
        disp(' maximum number of iterations was reached')
    elseif f<eps
        disp(' The algorithm stopped because the change in fit is now')
        disp(' smaller than the machine uncertainty.')
    else
        disp(' Algorithm stopped for some mysterious reason')
    end
end

function swloads = signswtch(loads,X);

%SIGNSWTCH switches sign of multilinear models so that signs are in
%accordance with majority of data
%
%
% I/O swloads = signswtch(loads,X);
%
% Factors must be a cell with the loadings. If Tucker or NPLS, then the
% last element of the cell must be the core array

try % Does not work in older versions of matlab
    warning('off','MATLAB:divideByZero');
end
sizeX=size(X);
order = length(sizeX);
for i=1:order;
    F(i) = size(loads{i},2);
end


if isa(X,'dataset')% Then it's a SDO
    inc=X.includ;
    X = X.data(inc{:});
end


% Compare centered X with center loading vector

if length(loads)==order % PARAFAC
    % go through each component and then update in the end
    for m = 1:order % For each mode determine the right sign
        for f=1:F(1) % one factor at the time
            s=[];
            a = loads{m}(:,f);
            x = permute(X,[m 1:m-1 m+1:order]);
            for i=1:size(x(:,:),2); % For each column
                id = find(~isnan(x(:,i)));
                if length(id)>1
                    try
                        c = corrcoef(x(id,i),a(id));
                    catch
                        disp('Oops - something wrong in signswtch - please send a note to')
                        whos
                    end
                    if isnan(c(2,1))
                        s(i)=0;
                    else
                        s(i) = c(2,1)*length(id); % Weigh correlation by number of elements so many-miss columns don't influence too much
                    end
                else
                    s(i) = 0;
                end
            end
            S(m,f) = sum(s);
        end
    end
    
    % Use S to switch signs. If the signs of S (for each f) multiply to a
    % positive number the switches are performed. If not, the mode of the
    % negative one with the smallest absolute value is not switched.
    
    for f = 1:F(1)
        if sign(prod(S(:,f)))<1 % Problem: make the smallest negative positive to avoid switch of that
            id = find(S(:,f)<0);
            [a,b]=min(abs(S(id,f)));
            S(id(b(1)),f)=-S(id(b(1)),f);
        end
    end
    % Now ok, so switch what needs to be switched
    for f = 1:F(1)
        for m = 1:order
            if sign(S(m,f))<1
                loads{m}(:,f)=-loads{m}(:,f);
            end
        end
    end
     
elseif length(loads)==(order+1) % NPLS/Tucker
    
    % go through each mode and update and correct core accordinglu
    for m = 1:order % For each mode determine the right sign
        for f=1:F(m) % one factor at the time
            a = loads{m}(:,f);
            x = permute(X,[m 1:m-1 m+1:order]);
            for i=1:size(x(:,:),2); % For each column
                id = find(~isnan(x(:,i)));
                if length(id)>1
                    c = corrcoef(x(id,i),a(id));
                    if isnan(c(2,1))
                        s(i)=0;
                    else
                        s(i) = c(2,1)*length(id); % Weigh correlation by number of elements so many-miss columns don't influence too much
                    end
                else
                    s(i) = 0;
                end
            end
            if sum(s) < 0
                % turn around
                loads{m}(:,f) = -loads{m}(:,f);
                
                % Then switch the core accordingly
                G = loads{order+1};
                G = permute(G,[m 1:m-1 m+1:order]);
                sizeG = size(G);
                G = reshape(G,sizeG(1),prod(sizeG)/sizeG(1));
                G(f,:) = -G(f,:);
                G = reshape(G,sizeG);
                G = ipermute(G,[m 1:m-1 m+1:order]);
                loads{order+1} = G;
            end
        end
    end
    
    
else
    error('Unknown model type in SIGNS.M')
end

swloads = loads;

function S = plae(X,A,Sold,nonnegativity);

% Least Absolute Error ``pseudoinverse'':
% iteratively solves the problem of min wrt S sum(sum(abs(X-A*S)))
% monotonically convergent in terms of LAE,
% but not guaranteed in general to find globally opt soln.
% ALL parameters are real-valued
% uses wmf.m
% complexity per complete iteration is order of F*N*I*logI
% where S is FxN, and A is IxF
% Assumes A is tall and full rank
% N. Sidiropoulos, April 12, 2000
% RB 2005, Added nonnegativity

[I,F]=size(A);
[I,N]=size(X);

if nargin<3
    S = pinv(A)*X; % use LS-solution as good initialization
else
    S = Sold;
end
LAE = sum(sum(abs(X-A*S)));
SMALLNUMBER = eps*10;
MAXNUMITER = 30;

%fprintf('LAE = %12.10f\n',LAE);
LAEold = 2*LAE;
LAEinit = LAE;
it     = 0;

while abs((LAE-LAEold)/LAEold) > SMALLNUMBER & it < MAXNUMITER & LAE > 10*eps
    it=it+1;
    LAEold=LAE;
    
    % update elements of S one by one:
    
    for f=1:F,
        Y = X - (A*S - A(:,f)*S(f,:));
        for n=1:N,
            S(f,n) = wmf(Y(:,n)./A(:,f),A(:,f));
        end
        if nonnegativity
            S(f,find(S(f,:)<0)) = 0;
        end
    end
    
    % compute new LAE:
    
    LAE = sum(sum(abs(X-A*S)));
    %fprintf('LAE = %12.10f\n',LAE);
    
end % while loop

function a = wmf(x,w);

% weighted median filter with non-negative real weights
% input (vector x) and output (scalar a) are both real-valued
% minimize sum_i abs(w(i)) * abs(x(i) - a)
% N. Sidiropoulos, April 12, 2000
% Ref: cf. e.g., Yang et al, IEEE Trans. Signal Proc. 43(3):591-592, Mar. 1995
% Complexity is NlogN, N=length(x), due to the sorting operation

[s,p] = sort(x);
absw = abs(w);
sw = absw(p);
t = 0.5*sum(absw);
N=length(x);
psum=0;
for n=N:-1:1,
    psum = psum + sw(n);
    if (psum >= t)
        a = s(n);
        break;
    end
end


function B=compresnonneg(X,Z,Bold,V,it)

% Find B minimizing ||X-Z*B'|| subject to V*B>=0
% B=compresnonneg(X,Z,V);

[I,J]=size(X);
[Iz,F]=size(Z);
[BJ,~]=size(Bold);
[VI,VJ]=size(V);
z=sparse(I*J,F*J);
for i=1:J;
    z((i-1)*Iz+1:i*Iz,(i-1)*F+1:i*F)=Z;
end

VV=sparse(VI*BJ,VJ*F);
co=0;
for i=1:VI;
    for f=1:F
        co=co+1;
        VV(co,f:F:end)=V(i,:);
    end
end

if it<10 % Impose constraints slowly during first ten iteration
    mx = -max(X(:));
    mx=mx*(10/(10-it));
    thrs=repmat(mx,size(VV,1),1); 
else
    thrs=sparse(size(VV,1),1);
end
B = lsi2(z'*z,z'*X(:),VV,thrs);
B=reshape(B,F,BJ)';

function x = lsi2(EE,Ef,G,h);
% x = lsi2(EE,Ef,G,h)
% Least squares with inequality constraints.
% Solves the following problem:
%       min || Ex - f || subject to Gx >= h
% Here input: EE = E'E and Ef=E'f

R=chol(EE);
RR=inv(R);
EEinv=RR*RR';
GG=G*RR;
hh=h-G*EEinv*Ef;
z=ldp(GG,hh);
if ~any(isnan(z))
    x=RR*z+EEinv*Ef;
else
    x=NaN;
end

function b=ldp(G,h);

% find min||b|| subject to Gb=>h
% 
% From Lawson & Hanson 74, p. 165
%
% Copyright 1998 
% Rasmus Bro
%


[I,J]=size(G);
E=[G';h'];
f=[zeros(J,1);1];
uhat=fastnnls(E'*E,E'*f);
r=E*uhat-f;

if norm(r)==0
  disp(' No solution to LDP problem')
elseif r(J+1)~=0
  b=-r(1:J)/r(J+1);
else
  b=NaN;
end
```
