## Supplementary File S1 for "Robust latent-variable interpretation of *in vivo* regression models by nested resampling": pfls.m

```
function load=pfls(ZtZ,ZtX,dimX,cons,OldLoad,DoWeight,W);

%PFLS
%
% See also:
% 'unimodal' 'monreg' 'fastnnls'
%
% 
% Calculate the least squares estimate of
% load in the model X=load*Z' => X' = Z*load'
% given ZtZ and ZtX
% cons defines if an unconstrained solution is estimated (0)
% or an orthogonal (1), a nonnegativity (2), or a unimodality (3)
%
%
% Used by PARAFAC.M

% $ Version 1.02 $ Date 28. July 1998 $ Not compiled $
% Apr 2002 - Fixed error in weighted ls $ rb

% Copyright (C) 1995-2006  Rasmus Bro & Claus Andersson
% Copenhagen University, DK-1958 Frederiksberg, Denmark,
%


if ~DoWeight

  if cons==0 % No constr
    %load=((Z'*Z)\Z'*Xinuse)';
    load=(pinv(ZtZ)*ZtX)';
  
  elseif cons==1 % Orthogonal loadings acc. to Harshman & Lundy 94
    load=ZtX'*(ZtX*ZtX')^(-.5);

  elseif cons==2 % Nonnegativity constraint
    load=zeros(size(OldLoad));
    for i=1:dimX
       load(i,:)=fastnnls(ZtZ,ZtX(:,i))';
%       if min(load(i,:))<-eps*1000
%          load(i,:)=OldLoad(i,:);
%       end
    end

  elseif cons==3 % Unimodality & NNLS
     load=OldLoad;
     F=size(OldLoad,2);
     if F>1
       for i=1:F
        ztz=ZtZ(i,i);
        ztX=ZtX(i,:)-ZtZ(i,[1:i-1 i+1:F])*load(:,[1:i-1 i+1:F])';
        beta=(pinv(ztz)*ztX)';
        load(:,i)=ulsr(beta,1);
       end
     else
       beta=(pinv(ZtZ)*ZtX)';
       load=ulsr(beta,1);
     end
  end

elseif DoWeight
  Z=ZtZ;
  X=ZtX;
  if cons==0 % No constr
    load=OldLoad;
    one=ones(1,size(Z,2));
    for i=1:dimX
      ZW=Z.*(W(i,:).^2'*one);
      %load(i,:)=(pinv(Z'*diag(W(i,:))*Z)*(Z'*diag(W(i,:))*X(i,:)'))';
      load(i,:)=(pinv(ZW'*Z)*(ZW'*X(i,:)'))';
    end

  elseif cons==2 % Nonnegativity constraint
    load=OldLoad;
    one=ones(1,size(Z,2));
    for i=1:dimX
      ZW=Z.*(W(i,:).^2'*one);
      load(i,:)=fastnnls(ZW'*Z,ZW'*X(i,:)')';
    end

  elseif cons==1
    disp(' Weighted orthogonality not implemented yet')
    disp(' Please contact the authors for further information')
    error

  elseif cons==3
    disp(' Weighted unimodality not implemented yet')
    disp(' Please contact the authors for further information')
    error

  end

end


% Check that NNLS and alike do not intermediately produce columns of only zeros
if cons==2|cons==3
  if any(sum(load)==0)  % If a column becomes only zeros the algorithm gets instable, hence the estimate is weighted with the prior estimate. This should circumvent numerical problems during the iterations
    load = .9*load+.1*OldLoad;
  end
end
```
