## Supplementary File S1 for "Robust latent-variable interpretation of *in vivo* regression models by nested resampling": pfplot.m

```
function pfplot(X,Factors,Weights,Option);
%PFPLOT plot parafac model
%
% See also:
% 'parafac'
%
%
% pfplot(X,Factors,Weights,Option);
% Different aspects for evaluation of the solution.
%
% Option # = 1
% 1	NOT ACCESIBLE
% 2	NOT ACCESIBLE
% 3	DIAGONALITY PLOT
% 4	PLOTS OF RESIDUAL VARIANCE
% 5	PLOTS OF LEVERAGE
% 6	RESIDUALS (STANDARD DEVIATION) VERSUS LEVERAGE
% 7	NORMAL PROBABILITY PLOT
% 8	LOADING PLOT
% 
% You HAVE to input all four inputs. If you have no weights, just input [].
% The last input must be an 8-vector with ones if you want the plot and
% zeros else. E.g.
%
% pfplot(X,factors,[],[0 0 1 0 0 0 0 1]);
%
% to have the diagonality and the loading plot
%

% $ Version 1.02 $ Date 28. July 1998 $ Not compiled $
% $ Version 1.03 $ Date 6. October 1999 $ Changed to handle missing values correctly$
% $ Version 2.00 $ May 2001 $ Changed to array notation $ RB $ Not compiled $

% Copyright (C) 1995-2006  Rasmus Bro & Claus Andersson
% Copenhagen University, DK-1958 Frederiksberg, Denmark,
%


warning off 

DimX = size(X);
X = reshape(X,DimX(1),prod(DimX(2:end)));

% Convert to old format
NewLoad = Factors;
ff = [];
for f=1:length(Factors)
  ff=[ff;Factors{f}(:)];
end
Factors = ff;


factors = Factors;
ord=length(DimX);
Fac=length(factors)/sum(DimX);
lidx(1,:)=[1 DimX(1)*Fac];
for i=2:ord
  lidx=[lidx;[lidx(i-1,2)+1 sum(DimX(1:i))*Fac]];
end
if Option(3)==1
  % ESTIMATE DIAGONALITY OF T3-CORE
  diagonality=corcond(reshape(X,DimX),NewLoad,Weights,1);
end
model=nmodel(NewLoad);
model = reshape(model,DimX(1),prod(DimX(2:end)));
if Option(4)==1
  % PLOTS OF RESIDUAL VARIANCE
  figure,eval(['set(gcf,''Name'',''Residual variance'');']);
  aa=ceil(sqrt(ord));bb=ceil(ord/aa);
  for i=1:ord
    r=nshape(reshape(X-model,DimX),i)';
    varian=stdnan(r).^2;
    subplot(aa,bb,i)
    plot(varian)
    if DimX(i)<30
      hold on
      plot(varian,'r+')
    end
    eval(['xlabel(''Mode ', num2str(i),''');']);
    ylabel('Residual variance');
  end
end
if Option(5)==1
  % PLOTS OF LEVERAGE
  figure
  eval(['set(gcf,''Name'',''Leverage'');']);
  aa=ceil(sqrt(ord));
  bb=ceil(ord/aa);
  for i=1:ord
    A=reshape(factors(lidx(i,1):lidx(i,2)),DimX(i),Fac);
    lev=diag(A*pinv(A'*A)*A');
    subplot(aa,bb,i)
    if std(lev)>eps
      plot(lev+100*eps,'+')
      for j=1:DimX(i)
        text(j,lev(j),num2str(j))
      end
    else
      warning('Leverage is constant')
    end
    eval(['xlabel(''Mode ', num2str(i),''');']);
    ylabel('Leverage');
  end
end
if Option(6)==1
  % RESIDUALS (STANDARD DEVIATION) VERSUS LEVERAGE
  figure
  eval(['set(gcf,''Name'',''Residuals vs. Leverages'');']);
  aa=ceil(sqrt(ord));
  bb=ceil(ord/aa);
  for i=1:ord
    subplot(aa,bb,i)
    A=reshape(factors(lidx(i,1):lidx(i,2)),DimX(i),Fac);
    lev=diag(A*pinv(A'*A)*A')'+100*eps;
    r=nshape(reshape(X-model,DimX),i)';
    stand=stdnan(r);
    if std(lev)>eps
      plot(lev,stand,'+')
      for j=1:DimX(i)
        text(lev(j),stand(j),num2str(j))
      end
      eval(['xlabel(''Leverage in mode ', num2str(i),''');']);
      ylabel('Standard deviation');
    else
      warning('Leverage is constant')
    end
  end
end
if Option(7)==1
  % NORMAL PROBABILITY PLOT
  if exist('normplot')
    disp(' ')
    disp(' Normal probability plots are time-consuming')
    disp(' They are made in the statistics toolbox though, so we can''t change that!')
    figure,
    eval(['set(gcf,''Name'',''Normal probability of residuals'');']);
    aa=ceil(sqrt(ord));
    bb=ceil(ord/aa);
    r=nshape(reshape(X-model,DimX),i)';
    r=r(:);
    normplot(r(find(~isnan(r))))
  end
end
if Option(8)==1
  % LOADING PLOT
  if sum(Option)>1
    figure
  end
  eval(['set(gcf,''Name'',''Loadings'');']);
  aa=ceil(sqrt(ord));
  bb=ceil(ord/aa);
  for i=1:ord
    subplot(aa,bb,i)
    A=reshape(factors(lidx(i,1):lidx(i,2)),DimX(i),Fac);
    plot(A)
    eval(['xlabel(''Mode ', num2str(i),''');']);
    ylabel('Loading');
  end
end
drawnow
```
