## Supplementary File S1 for "Robust latent-variable interpretation of *in vivo* regression models by nested resampling": pftest.m

```
function [ssX,Corco,It,Factors] = pftest(NumRep,X,Fac,Options,const,OldLoad,Weights);

%PFTEST find the number of PARAFAC components
%
% See also:
% 'corcondia'
%
%
% TEST HOW MANY COMPONENTS TO USE IN PARAFAC MODEL
% 
% Used for testing the appropriate number of components in a PARAFAC
% model. Input the appropriate input options (see PARAFAC for help) as
% well as the number of replicate fittings to use (NumRep). For one to Fac 
% component models are fitted each NumRep times.
% Three measures are output in three matrices, where the f'th row hold the values
% for the f-component model and each column correspond to a specific replicate run
%
% ssX:   THE SUM-SQUARED ERROR
%
%        What-To-Look-For:
%        look for sudden change as in a Scree-plot (often difficult)
%        and look for sudden increase in number of local minima (replicate
%        points for one component are not identical). This is often a good
%        indication that noise is being modeled.
%
% Corco: CORE CONSISTENCY DIAGNOSTIC (CORCONDIA)
%
%        What-To-Look-For:
%        CORCONDIA is a percentage below or equal to 100%. A value of 80-100% 
%        means that the model is valid, while a value below, say 40% means that
%        the model is not valid. A value between 40 and 80% means that the model
%        is probably valid but somehow difficult to estimate, e.g., due to 
%        slight misspecification or correlations. The Corcondia will mostly
%        decrease with number of components but very sharply where the correct
%        number of components are exceeded. Hence, the appropriate number of
%        components is the model with the highest number of components and a
%        valid CORCONDIA
%
% It:    NUMBER OF ITERATIONS
%   
%        What-To-Look-For:
%        A sudden increase in the number of iterations needed
%        suggests that too many components may be used (but could also
%        just be due to a difficult problem).
%
% I/O FORMAT as PARAFAC with an additional parameter, NumRep, for the number
% of replicate runs
%
% [ssX,Corco,It] = pftest(NumRep,X,Fac,Options,const,OldLoad,Weights);
%
% Note that plots are only produced in Matlab ver. > 5.2

% $ Version 2.12 $ Jan 2004 $ Loadings can be viewed by pressing circles $ RB $ Not compiled $
% $ Version 2.11 $ Jan 2004 $ Added iterations in the output/plot $ RB $ Not compiled $
% $ Version 2.01 $ Feb 2003 $ Fixed error in IO $ RB $ Not compiled $
% $ Version 2.00 $ May 2001 $ Changed to array notation $ RB $ Not compiled $
% $ Version 1.02 $ Date 28. July 1998 $ Not compiled $

% Copyright (C) 1995-2006  Rasmus Bro & Claus Andersson
% Copenhagen University, DK-1958 Frederiksberg, Denmark,
%

if nargin==0
  disp(' ')
  disp(' PFTEST for determining the number of components')
  disp(' ')
  disp(' I/O      : [ssX,Corco] = pftest(NumRep,X,Fac,Options,const,OldLoad,Weights);')
  disp(' I/O short: [ssX,Corco] = pftest(NumRep,X,Fac);')
  disp(' ')
  return
end

if nargin<4
  Options=[];
end
if nargin<5
  const=[];
end
if nargin<6
  OldLoad=[];
end
if nargin<7
  FixMode=[];
end
if nargin<8
  Weights=[];
end

ssX=zeros(Fac,NumRep);
Corco=zeros(Fac,NumRep);


FactorsOut=[];
for f=1:Fac
  Options(2)=0; % DTLD init
  [Factors,it,err,corcondia]=parafac(X,f,Options,const,OldLoad,FixMode,Weights);
  ssX(f,1)=err; 
  BestErr=err;
  Corco(f,1)=corcondia;
  FactorsNow=Factors;
  FactorsAll{f,1}=Factors;
  It(f,1)=it;
  
  Options(2)=1; % SVD init
  [Factors,it,err,corcondia]=parafac(X,f,Options,const,OldLoad,FixMode,Weights);
  ssX(f,2)=err;
  Corco(f,2)=corcondia;
  if err<BestErr
    FactorsNow=Factors;
    BestErr=err;
  end
  It(f,2)=it;
  FactorsAll{f,2}=Factors;
  
  Options(2)=2; % random init
  for rep=3:NumRep
    [Factors,it,err,corcondia]=parafac(X,f,Options,const,OldLoad,FixMode,Weights);
    ssX(f,rep)=err;
    Corco(f,rep)=corcondia;
    if err<BestErr
      FactorsNow=Factors;
      BestErr=err;
    end
    FactorsAll{f,rep}=Factors;
    It(f,rep)=it;
  end

  FactorsOut=[FactorsOut;FactorsNow];
end

% Only plot if MATLAB version >= 5
VER = version;
if VER(1)~='4'
  delta=.25/NumRep; % shift points a little horizontally for appearance
  figure
  set(gcf,'userdata',FactorsAll);
  subplot(3,1,1)
  for r=1:NumRep
    for f=1:Fac
      gg=plot(f+(r-1)*delta,ssX(f,r), ...
        'MarkerEdgeColor','k','MarkerFaceColor','r', ...
        'LineWidth',2,'Marker','o','LineStyle','none', ...
        'MarkerSize',8);
      set(gg,'userdata',[f r]);
      set(gg,'ButtonDownFcn',['fac=get(gcf,''userdata'');f=get(gco,''userdata'');figure,plotfac(fac{f(1),f(2)}),set(gcf,''name'',[''Comp# '',num2str(f(1)),'' - Repl# '',num2str(f(2))]);']);
      hold on
    end
  end
  axis([1-.1 Fac+1 0-.05*max(ssX(:)) 1.05*max(ssX(:)) ])
  set(gca,'XTick',[1:Fac])
  ylabel('Residual sum of squares','FontWeight','bold')
  title('PARAFAC TEST - press a circle in top plot to see loadings ','FontWeight','bold')
  hold off
  subplot(3,1,2)
  for r=1:NumRep
    plot(1+(r-1)*delta:1:Fac+(r-1)*delta,Corco(:,r), ...
      'MarkerEdgeColor','k','MarkerFaceColor','r', ...
      'LineWidth',2,'Marker','o','LineStyle','none', ...
      'MarkerSize',8)
    hold on
  end
  hold off
  MinCo=min(Corco(:));
  axis([1 Fac+1 min([MinCo 0]) 100 ]);
  set(gca,'XTick',[1:Fac])
  ylabel('Core consistency','FontWeight','bold')

  subplot(3,1,3)
  for r=1:NumRep
    plot(1+(r-1)*delta:1:Fac+(r-1)*delta,It(:,r), ...
      'MarkerEdgeColor','k','MarkerFaceColor','r', ...
      'LineWidth',2,'Marker','o','LineStyle','none', ...
      'MarkerSize',8)
    hold on
  end
  axis([1 Fac+1 0 max(it(:)) ])
  set(gca,'XTick',[1:Fac])
  ylabel('Numb. of iterations','FontWeight','bold')
  hold off
  xlabel('Number of components','FontWeight','bold')
  
end

if nargout>3
  disp(' ')
  whic = input([' For which number of components (1:',num2str(Fac),') do you want the parameters : ']);
  Factors=FactorsOut(whic,:);
end
```
