## Supplementary File S1 for "Robust latent-variable interpretation of *in vivo* regression models by nested resampling": plotfac.m

```
function []=plotfac(Factors,H,Axis0,Axis1,Axis2);
%PLOTFAC Plot the contents of Factors
%	
% plotfac(Factors,[[,H],Axis1,Axis2,Axis3]);
% 
% Use this function to plot the factors that are stored in
% the row-vector Factors. Afterwards you can assign
% you own titles a.s.o. If H is supplied, the factors
% in the first mode are scaled with the diagonal
% elements. If Limits is provided as a 2-column matrix
% with lower (1st col) and upper (2nd col) boundaries
% for each mode. Specify 'NaN' for unknown.

% $ Version 2.00 $ May 2001 $ Changed to array notation $ RB $ Not compiled $
% $ Version 1.02 $ Date 11. October 1999 $ Not compiled $

% Copyright (C) 1995-2006  Rasmus Bro & Claus Andersson
% Copenhagen University, DK-1958 Frederiksberg, Denmark,
%

cmapp=[
    1.0000	0       0;
    0	0.6250	1.0000;
    1.0000  0.7500	0;
    0.5000  1.0000	0;
    0.1250	0	1.0000;
    0    	1.0000	0.2500;
    0.8750	0	1.0000;
    1.0000	0	0.3750;		
    0    	1.0000	1.0000;
    1.0000	0.3750	0;
    0    	0.2500	1.0000;
    0.8750	1.0000	0;
    0.5000	0	1.0000;
    0.1250	1.0000	0;
    1.0000	0	0.7500;
    0	1.0000	0.6250];
 
for f = 1:length(Factors)
   Fac(f) = size(Factors{f},2);
   DimX(f) = size(Factors{f},1);
end

% Convert to old format
ff = [];
for f=1:length(Factors)
 ff=[ff;Factors{f}(:)];
end
Factors = ff;
 
if length(Fac)==1,
    Fac=Fac*ones(size(DimX));
end;

Fac_orig=Fac;
Fac(find(Fac==-1))=0;
CC_=length(DimX(1,:));
if ~exist('H') | isempty(H),
    H=eye(Fac(1));
end;
if ~exist('Axis0') | isempty(Axis0),
    Axis0=NaN*ones(CC_,2);
end;
if ~exist('Axis1') | isempty(Axis1),
    Axis1=NaN*ones(CC_,2);
end;
if ~exist('Axis2') | isempty(Axis2),
    Axis2=NaN*ones(CC_,2);
end;

ColLst='mcrgbky';
FIdx0=cumsum([1 DimX(1:CC_-1).*Fac(1:CC_-1)]);
FIdx1=cumsum([DimX.*Fac]);
set(gcf,'DefaultAxesColorOrder',cmapp);
set(gcf,'PaperType','A4');
screen = get(0, 'ScreenSize');
width = screen(3);
height = screen(4);
w_width = round(width/2);
w_height = round(height/2);
set(gcf,'Position',[width-w_width-15 height-w_height-75 w_width w_height]);
sa=1:CC_;
i=find(Fac_orig==-1);
if ~isempty(i),
    sa(i)=[];
end;
nCC_=size(sa,2);
for c=sa,
    t=reshape(Factors(FIdx0(c):FIdx1(c)),DimX(c),Fac(c));
    if c==1,
        t=t*H;
    end;
    tmax=max(max(t));
    tmin=min(min(t));
    elong=max([abs(tmax) abs(tmin)])*0.01;
    tmax=tmax+elong;
    tmin=tmin-elong;
    if tmax-tmin<=eps,
        tmax=tmin+1;
    end;
    if nCC_>1,
        subplot(floor((nCC_+1)/2),2,c);
    end;
    switch c;
        case 1,
            Limits=Axis0;
        case 2,
            Limits=Axis1;
        case 3,
            Limits=Axis2;
        end;
    if any(isnan(Limits)),
        xmin=1;
        xmax=size(t,1);
        plot([xmin:((xmax-xmin)/(size(t,1)-1)):xmax],t);
        axis([xmin xmax tmin tmax]);
    else
        if size(Limits,2) == 2,
            xmin=Limits(1);
            xmax=Limits(end);
            plot([xmin:((xmax-xmin)/(size(t,1)-1)):xmax],t);
            axis([xmin xmax tmin tmax]);
        else
            xmin=min(Limits);
            xmax=max(Limits);
            plot(Limits,t);
            axis([xmin xmax tmin tmax]);
        end;
    end;
    if exist('mode_strings'),
        str=deblank(mode_strings(c,:));
    else
        str=['Mode #' num2str(c)'];
    end;
    title(str);
    grid;
    
    drawnow;
end;

h4 = uicontrol('Parent',gcf,'Units','normalized','Style','text','Tag','StaticText1','HandleVisibility','off','HorizontalAlignment','center','Position', ...
        [0.01 .001 0.8 0.03],'FontSize',8,'foregroundcolor','blue','handlevisibility','on', ...
      'String','Component order (red,blue,yellow,green,lightblue,lightgreen,purple)','fontname','verdana');
```
