## Supplementary File S1 for "Robust latent-variable interpretation of *in vivo* regression models by nested resampling": ppp.m

```
cfunction AB=ppp(A,B);

% $ Version 1.02 $ Date 28. July 1998 $ Not compiled $
% The parallel proportional profiles product - triple-P product
% For two matrices with similar column dimension the triple-P product
% is ppp(A,B) = [kron(B(:,1),A(:,1) .... kron(B(:,F),A(:,F)]
% 
% AB = ppp(A,B);
%
% NB. This file is obsolete. Use kr.m instead but not that it takes
% inputs oppositely

% Copyright (C) 1995-2006  Rasmus Bro & Claus Andersson
% Copenhagen University, DK-1958 Frederiksberg, Denmark,
%

disp('PPP.M is obsolete and will be removed in future versions. ')
disp('use KRB.M instead. Note that krb(B,A) = ppp(A,B)')

[I,F]=size(A);
[J,F1]=size(B);

if F~=F1
   error(' Error in ppp.m - The matrices must have the same number of columns')
end

AB=zeros(I*J,F);
for f=1:F
   ab=A(:,f)*B(:,f).';
   AB(:,f)=ab(:);
end
```
