## Supplementary File S1 for "Robust latent-variable interpretation of *in vivo* regression models by nested resampling": readme.txt

IMPORTANT NOTES ON THE N-WAY TOOLBOX ver. 3.20

There are some important details that are necessary to know in order to be able to use the N-way toolbox properly. These are given here - please read carefully before using the toolbox.


CONDITIONS
Copyright (C) 1995-2012  Rasmus Bro & Claus Andersson
Copenhagen University, DK-1958 Frederiksberg, Denmark,

for more details. 
Preferably refer to

C. A. Andersson and R. Bro. The N-way Toolbox for MATLAB. Chemom.Intell.Lab.Syst. 52 (1):1-4, 2000.

or alternatively 

The N-way Toolbox for MATLAB ver. 3.20, http://www.models.life.ku.dk/
R. Bro & C. A. Andersson
Faculty of Life Sciences
Copenhagen University
DK-1958 Frederiksberg
Denmark


WHERE DOES THE TOOLBOX WORK?
The toolbox has been tested on matlab 7.3 in Windows XP/Vista/7 only. The toolbox uses features that are not compatible with matlab 4.x, so if you have matlab 4.x you should use version 1.04 of this toolbox instead. 


SETTING UP THE TOOLBOX
In order to install the toolbox, simply (extract and) copy the files to a directory (e.g. NWAY). After copying all files, go to the 'update' homepage in order to see if newer versions of individual files are available. Copy these files indiviually overwriting the old files. Make sure that the updates are copied after the main files. 

Make sure that the path ../nway is included in MATLAB's path. If you have e.g. the PLS_toolbox, some files are named identically. This may cause problems depending on which functions you use. If you want to use e.g. the parafac function from the N-way toolbox, you have to ensure that either the path to the N-way toolbox appears before the path to the PLS_toolbox or that you run matlab from the nway directory.

In order to get help on what files are present in the toolbox type <<help nway>> at the matlab command line (if nway is the name of the directory where you have the files.


DATA INPUT
Unlike, older matlab 4 compatible versions of this toolbox, the data are input directly as multi-way arrays. Hence, there is no need for the DimX used earlier for defining the size of the array. If you have a 10x8x100 array, X, that is held in a 10x(8*100) matrix, i.e. the old matrix format, you can convert to
a three-way array by

X = reshape(X,10,8,100);

This is the format in which the data must be input to the functions.


MODEL OUTPUT
Also the output has changed in most cases since version 1. With the use of cell arrays, it is much easier to handle the output of a varying number of component matrices. Let the components of a three-way parafac model is held in a cell e.g. called Factors; e.g. arising from the call of a four-component model

Factors = parafac(X,3);

Then the first mode loadings are held in Factors{1}:

A = Factors{1};
B = Factors{2};
C = Factors{3};

For a Tucker model such as

[Factors,G]=tucker(X,[3 3 2]);

the components are found similarly and G will be a 3x3x2 array.

For a tri-PLS2 model (three-way X, two-way Y) two component sets are defined

Xfactors - T=Xfactors{1}, Wj=Xfactors{2}, Wk = Xfactors{3}
Yfactors - U=Yfactors{1}, Q=Yfactors{2}

Instead of using the cell notation, it is possible to use the  M-file FAC2LET (factors to letters) to extract components; e.g.

[A,B,C] = fac2let(Factors);


MISSING DATA
For all algorithms the same flag is used for missing elements, namely NaN. If you have a data set, X, where missing elements are, e.g.,  designated by the number -9999, you can easily modify the data as

X(find(X==-9999))=NaN*find(X==-9999);


SUPPORT
We are interested in and dependent on feedback from the users. If you have problems running the toolbox please supply screendumps as well as version number of the toolbox, MATLAB, and operating system before contacting us. We will do the utmost to help overcoming the problems. In the rare event that the support required is very time-consuming we will have to charge for this service.

The authors may be contacted by email:

 (primarily Tucker and application/helper programs)
 (primarily PARAFAC/N-PLS)
