## Supplementary File S1 for "Robust latent-variable interpretation of *in vivo* regression models by nested resampling": refold3.m

```
function [Y]=Refold3(X,R);
%REFOLD3
%
%[Y]=Refold3(X,R);
%
%If you have confused the way you unfolded
%your data, you can mend it by this function

% Copyright (C) 1995-2006  Rasmus Bro & Claus Andersson
% Copenhagen University, DK-1958 Frederiksberg, Denmark,

Y=zeros(size(X));
Olis=zeros(1,R(3)*R(2));
lis=[0 kron([1:R(2)],R(3))];
lis=lis(1:R(2));
for k=1:R(3),
   Olis(1,(k-1)*R(2)+1:k*R(2))=k + lis;
end;
Y=X(1:R(1),Olis);
```
