## Supplementary File S1 for "Robust latent-variable interpretation of *in vivo* regression models by nested resampling": runMRBpls.m

```
%Run PLS on MRB AngII data
%Alex Caulk
%Yale University
%February 2018

%--------------------------------------------------------------------------
%Matt's data ordered in matrix as MRBa2(i,j,k)
%
%Rows (i) 1-4: ATA, DTA, SAA, IAA
%
%Columns (j) variables
%1: Systolic BP
%2: OD
%3: H
%4: L
%5: od systolic
%6: ir systolic
%7: h systolic
%8: estimated in vivo axial stretch
%9: systolic circ stress
%10: systolic ax stress
%11: systolic circ stiff
%12: systolic ax stiff
%13: systolic struct stiff
%14: systolic stored energy
%15: distensibility
%16: medial cross-sectional area
%17: adventitial cross-sectional area
%18: medial elastin area fraction
%19: medial SMC area fraction
%20: medial collagen area fraction
%21: medial GAG area fraction
%22: adventitial elastin area fraction
%23: adventitial collagen area fraction
%24: medial CD3 cell density
%25: medial CD45 cell density
%26: medial CD68 cell density
%27: adventitial CD3 cell density
%28: adventitial CD45 cell density
%29: adventitial CD68 cell density
%30: medial MMP2 area fraction
%31: medial MMP12 area fraction
%32: medial MMP13 area fraction
%33: adventitial MMP2 area fraction
%34: adventitial MMP12 area fraction
%35: adventitial MMP13 area fraction
%
%Columns (k) 1-6 time points: 0,4,7,14,21,28 days AngII infusion
%--------------------------------------------------------------------------

clear
close all
clc

load('MRB_AngII.mat')
vessel = {'ATA','DTA','SAA','IAA'};
times = {'0d','4d','7d','14d','21d','28d'};
vars = {'BP','OD','H','L','od','ir','h','lziv','tqq','tzz','Cqqqq','Czzzz',...
        'Cth','W','Dist','CSAm','CSAa','eAFm','SMCAFm','cAFm','GAGAFm',...
        'eAFa','cAFa','CD3m','CD45m','CD68m','CD3a','CD45a','CD68a',...
        'MMP2m','MMP12m','MMP13m','MMP2a','MMP12a','MMP13a'};

%--------------------------------------------------------------------------
%Choose vessel regions to evaluate to set i indices
%--------------------------------------------------------------------------

ves_choice = listdlg('PromptString','Select Vessels to Include in Evaluation','SelectionMode','multiple','ListString',vessel);

%--------------------------------------------------------------------------
%Choose data for X matrix to set j indices of X
%--------------------------------------------------------------------------

varX_choice = listdlg('PromptString','Select Variables for X Matrix','SelectionMode','multiple','ListString',vars);

%--------------------------------------------------------------------------
%Choose data for Y matrix to set j indices of Y
%--------------------------------------------------------------------------

vars_leftover = vars;
vars_leftover(varX_choice) = []; %Remove choices for X to avoid repeat data in X and Y matrices
varY_choice = listdlg('PromptString','Select Variables for Y Matrix','SelectionMode','multiple','ListString',vars_leftover);

%--------------------------------------------------------------------------
%Choose time points for evaluation to set k indices
%--------------------------------------------------------------------------

time_choice = listdlg('PromptString','Select Times to Include in Evaluation','SelectionMode','multiple','ListString',times);

%--------------------------------------------------------------------------
%run PLS on selected data
%--------------------------------------------------------------------------

X = MRBa2(ves_choice,varX_choice,time_choice);
Y = MRBa2(ves_choice,varY_choice,time_choice);

for j = 1:size(X,2)
    for k = 1:size(X,3)
        X_norm(:,j,k) = (X(:,j,k)-mean(X(:,j,k)))/std(X(:,j,k));
    end
end

for j = 1:size(Y,2)
    for k = 1:size(Y,3)
        Y_norm(:,j,k) = (Y(:,j,k)-mean(Y(:,j,k)))/std(Y(:,j,k));
    end
end

% [T,P,U,Q,b,W] = multiPLS(X_data,Y_data);
[Xfactors,Yfactors,Core,B,ypred,ssx,ssy] = npls(X_norm,Y_norm,3,NaN);
```
