## Supplementary File S1 for "Robust latent-variable interpretation of *in vivo* regression models by nested resampling": SetNaNs1.m

```
function XCorr=SetNaNs1(X,ExMin,ExMax,EmMin,EmMax,WidthMin,WidthMax,Value,MaxOrder,Mode);
%SETNANS1 fluorescence artifact treatment
%
%XCorr=SetNaNs1(X,ExMin,ExMax,EmMin,EmMax,WidthMin,WidthMax,Value,MaxOrder,Mode);
%
%This algorithm takes a fluorescence landscape with
%excitation ranges and emission ranges defined by
%ExMin, ExMax, EmMin and EmMax. Given these specifications
%the algorithm identifies the Rayleigh-peak and replaces
%a certain number of neighboring wavelengths at each side
%of the peak with the 'Value' multiplied by
%the existing value in the range, - use NaN to set missing.
%
%Generally the peaks broaden as the excitation wavelength increase.
%The 'WidthMin' is the width of the Rayleigh peaks at the lowest
%excitation wavelength and 'WidthMax' is the peakwidth at the highest
%excitation wavelength. Giv also the width of the 2nd order Rayleigh
%peak of the first spectrum as 'Width2nd' since 2nd order scattering may be
%much wider than first order. The algorithm replaces up to and including
%scattering of 'MaxOrder' order starting from the 1st (no 0th order).
%'Mode'=1 uses the lowest emission wavelength as low limit, hence
%removing all emission wavelengths on the low side of the 0th order
%Rayleigh peak. 'Mode'=2 removes only the peak itself.
%
%The landscape is contained in the matrix X and each row in the matrix
%represents one emission spectrum. Hence, X has dimensions (a x b) where
%a is the number of excitation lines, and b is the number
%of measured emission wavelengths.
%
%                                  b
%          -------------------------
%          | 1st emission spectrum |
%          | 2nd emission spectrum |
%          |         ....          |
%          | ath emission spectrum |
%        a -------------------------
%
% %XCorr=SetNaNs1(X,ExMin,ExMax,EmMin,EmMax,WidthMin,WidthMax,Value,MaxOrder,Mode);


% Copyright (C) 1995-2006  Rasmus Bro & Claus Andersson
% Copenhagen University, DK-1958 Frederiksberg, Denmark,

XCorr=X;
[a b]=size(X);

exax=ExMin:(ExMax-ExMin)/(a-1):ExMax;
emax=EmMin:(EmMax-EmMin)/(b-1):EmMax;

if WidthMin==WidthMax,
   Width=WidthMin*ones(1,a);
else
   Width=WidthMin:(WidthMax-WidthMin)/(a-1):WidthMax;
end;

WFactor=[1 1.4 1.7 1.9 2];

for order=1:MaxOrder,
   
   for excount=1:a,
      
      exc=order*exax(excount);
      
      W = WFactor(order)*Width(excount);
      
      rangemin = max(find(emax<=(exc-W)));
      rangemax = min(find(emax>=(exc+W)));
      
      if (exc-W) < EmMin,
         rangemin = 1;
      end;
      
      if (exc-W) > EmMax,
         rangemin = 0;
      end;
      
      if (exc+W) < EmMin,
         rangemax = 0;
      end;
      
      if (exc+W) > EmMax,
         rangemax = b;
      end;
      
      if ~(rangemax*rangemin==0),
         if Mode==1 & order==1;
            rangemin = 1;
         end;
         XCorr(excount,rangemin:rangemax)=Value*X(excount,rangemin:rangemax);
         [rangemin rangemax];
      end;
      
   end;
end;
```
