## Supplementary File S1 for "Robust latent-variable interpretation of *in vivo* regression models by nested resampling": setopts.m

```
function Options=setopts(Model);
%SETOPTS
%
%Options=setopts(Model);
%

% Copyright (C) 1995-2006  Rasmus Bro & Claus Andersson
% Copenhagen University, DK-1958 Frederiksberg, Denmark,


assgn=0;

if strcmp(upper(Model),'TUCKER'),
   Options=zeros(9,3);
   Options(1,1:2)=[1e-7 1e-8];
   Options(2,1)=0;
   Options(3,1)=1;
   Options(4,1)=1;
   Options(5,1)=1;
   Options(6,1:2)=[1000 50];
   Options(7,1:2)=[1 2];
   Options(8,1)=0;
   Options(9,1:2)=[1 1];
   Options(10,1)=160;
   save('noptions_t.mat','-v4','Options')
   assgn=1;
end;

if strcmp(upper(Model),'PARAFAC'),
   Options=zeros(9,3);
   Options(1,1:2)=[1e-6 1e-6];
   Options(2,1)=0;
   Options(3,1)=1;
   Options(4,1)=1;
   Options(5,1)=5;
   Options(6,1:2)=[1000 1000];
   Options(7,1:2)=[1 2];
   Options(8,1)=0;
   Options(9,1:2)=[1 1];
   Options(10,1)=60;
   save('noptions_pf.mat','-v4','Options')
   assgn=1;
end;

if ~assgn,
   Options=zeros(9,3);
   Options(1,1:2)=[1e-6 1e-6]; %Relative convergence criteria
   Options(2,1)=0;
   Options(3,1)=1;
   Options(4,1)=1;
   Options(5,1)=5;
   Options(6,1:2)=[1000 1000]; %Max. number of inner and outer iterations.
   Options(7,1:2)=[1 2];
   Options(8,1)=0;
   Options(9,1:2)=[1 1];  %Flags for amount of printout. 0=minimum, 1=medium, 2=full.
   Options(10,1)=60; %Parameters for saving temporary results
   fprintf('Identifier in ''Model'' not found, ''Options'' will be subdefault.\n')
end;

%---------------------------------------------
% Options definition's
%
%   Options=zeros(9,7);
%   Options(1,1:2)=[1e-6 1e-6]; %Relative convergence criteria
%   Options(2,1)=0; %Initialization
%   Options(3,1)=1; %
%   Options(4,1)=1;
%   Options(5,1)=5;
%   Options(6,1:2)=[1000 1000]; %Max. number of inner and outer iterations.
%   Options(7,1:2)=[1 2];
%   Options(8,1)=0;
%   Options(9,1:2)=[1 1];  %Flags for amount of printout. 0=minimum, 1=medium, 2=full.
%   Options(10,1)=60; %Parameters for saving temporary results
```
