## Supplementary File S1 for "Robust latent-variable interpretation of *in vivo* regression models by nested resampling": stdnan.m

```
function st=stdnan(X);

%STDNAN estimate std with NaN's
%
% Estimates the standard deviation of each column of X
% when there are NaN's in X.
%
% Columns with only NaN's get a standard deviation of zero


% $ Version 1.02 $ Date 28. July 1998 $ Not compiled $

% Copyright (C) 1995-2006  Rasmus Bro & Claus Andersson
% Copenhagen University, DK-1958 Frederiksberg, Denmark,

[I,J]=size(X);

st=[];
for j=1:J
  id=find(~isnan(X(:,j)));
  if length(id)
    st=[st std(X(id,j))];
  else
    st=[st 0];
  end
end
```
