## Supplementary File S1 for "Robust latent-variable interpretation of *in vivo* regression models by nested resampling": t3core.m

```
function G=T3core(X,Load,Weights,NonNeg);
%T3CORE calculate Tucker core
%
% G=T3core(X,Load,Weights,NonNeg);
% Calculate a Tucker3 core given X, the loadings, Load
% in vectorized format and optionally Weights. Missing NaN, NonNeg = 1 => nonnegativity


% Copyright (C) 1995-2006  Rasmus Bro & Claus Andersson
% Copenhagen University, DK-1958 Frederiksberg, Denmark,
%


% $ Version 2.00 $ May 2001 $ Changed to array notation $ RB $ Not compiled $


for i = 1:length(Load)
   Fac(i) = size(Load{i},2);
end

DimX = size(X);
X = reshape(X,DimX(1),prod(DimX(2:end)));

% Convert to old format
ff = [];
for f=1:length(Load)
 ff=[ff;Load{f}(:)];
end
Load = ff;

ord=length(DimX);

if length(Fac)==1
   Fac=ones(1,ord)*Fac;
end

l_idx=0;
for i=1:ord
   l_idx=[l_idx sum(DimX(1:i).*Fac(1:i))];
end

if exist('Weights')~=1,
   Weights=0;
elseif isempty(Weights)
    Weights = 0;
end

if exist('NonNeg')~=1,
   NonNeg=0;
end


if any(isnan(X(:)))
   id=find(isnan(X));
   M=ones(size(X));
   M(id)=zeros(size(id));
   X(id)=1000000*ones(size(id));
   if prod(size(Weights))==prod(DimX)
      %Modified by GT
      Weights = reshape(Weights,DimX(1),prod(DimX(2:end)));
      %End GT
      Weights = Weights.*M;
   else
      Weights=M;
   end
end


if prod(size(Weights))==prod(DimX) % Use weighted approach
   LL=reshape(Load(l_idx(1)+1:l_idx(2)),DimX(1),Fac(1));
   xtz=zeros(1,prod(Fac));
   ztz=zeros(prod(Fac),prod(Fac));
   for i=1:DimX(1)
      L1=reshape(Load(l_idx(ord)+1:l_idx(ord+1)),DimX(ord),Fac(ord));
      L2=reshape(Load(l_idx(ord-1)+1:l_idx(ord)),DimX(ord-1),Fac(ord));
      Z=kron(L1,L2);
      for ii=ord-2:-1:2
         L=reshape(Load(l_idx(ii)+1:l_idx(ii+1)),DimX(ii),Fac(ii));
         Z=kron(Z,L);
      end
      Z=kron(Z,LL(i,:));
      ztz=ztz+(Z.*(Weights(i,:)'*ones(1,size(Z,2)) ))'*Z;
      xtz=xtz+(X(i,:).*Weights(i,:))*Z;
   end
   if NonNeg==1;
      G=fastnnls(ztz,xtz');
   else
      G=pinv(ztz)*xtz';
   end
   
else % No weighting

   ztz=zeros(prod(Fac),prod(Fac));
   
   L1=reshape(Load(l_idx(ord)+1:l_idx(ord+1)),DimX(ord),Fac(ord));
   L1tL1=L1'*L1;
   L2=reshape(Load(l_idx(ord-1)+1:l_idx(ord)),DimX(ord-1),Fac(ord-1));
   L2tL2=L2'*L2;
   ztz=kron(L1tL1,L2tL2);
   for o=ord-2:-1:1,
      L=reshape(Load(l_idx(o)+1:l_idx(o+1)),DimX(o),Fac(o));
      LtL=L'*L;
      ztz=kron(ztz,LtL);
   end
   
   xtz=zeros(prod(Fac),1);
   F=ones(ord,1);
   F(1)=0;
   for f=1:prod(Fac)
      F(1)=F(1)+1;
      for ff=1:ord-1
         if F(ff)==Fac(ff)+1;
            F(ff+1)=F(ff+1)+1;
            F(ff)=1;
         end
         % F runs through all combinations of factors
      end
      L=reshape(Load(l_idx(1)+1:l_idx(2),:),DimX(1),Fac(1));
      cc=L(:,F(1))'*X;
      for j=ord:-1:3,
         ccc=zeros(prod(DimX(2:j-1)),DimX(j));
         ccc(:)=cc;
         cc=ccc';
         L=reshape(Load(l_idx(j)+1:l_idx(j+1),:),DimX(j),Fac(j));
         cc=L(:,F(j))'*cc;
      end
      L=reshape(Load(l_idx(2)+1:l_idx(2+1),:),DimX(2),Fac(2));
      cc=L(:,F(2))'*cc';
      xtz(f)=cc;
   end
   if NonNeg==1,
      G=fastnnls(ztz,xtz);
   else
      G=pinv(ztz)*xtz;
   end
end

G = reshape(G,Fac);
```
