## Supplementary File S1 for "Robust latent-variable interpretation of *in vivo* regression models by nested resampling": tuckdemo.m

```
%tuckdemo.m
%
% Made for the Self-Study application
%
% REQUIRES THE DATA SETS !!!!!
%

% Copyright (C) 1995-2006  Rasmus Bro & Claus Andersson
% Copenhagen University, DK-1958 Frederiksberg, Denmark,
%


close all
clear all
load('data\dataset1');
load('data\dataset1res');

fprintf('\n 1 Pak ### Inspect raw data - press a key..\n');pause
figure(1);set(gcf,'Position',[-1 31 804 534]);
for i=1:28,
   m=reshape(X(i,:),R(2),R(3));
   mesh(EmAx,ExAx,m);
   title(['Raw data. Time ' int2str(i)]);
   xlabel('Excitation [nm]')
   ylabel('Emission [nm]')
   axis([EmAx(1) EmAx(311) ExAx(1) ExAx(20) 0 1000]);
   grid on
   drawnow
end;

fprintf('\n 2 Pak ### Inspect calibration data - press a key..\n');pause
%Remove wrong/disturbing observations
for i=1:28,
   m=reshape(X(i,:),R(2),R(3));
   mn=SetNaNs1(m,ExAx(1),ExAx(20),EmAx(1),EmAx(311),18,20,NaN,2,0);
   mnr=cmatrep(mn,'gt',999.99,NaN);
   mesh(EmAx,ExAx,mnr);
   title(['Corrected data. Time ' int2str(i)]);
   xlabel('Excitation [nm]')
   ylabel('Emission [nm]')
   axis([EmAx(1) EmAx(311) ExAx(1) ExAx(20) 0 1000]);
   grid on
   drawnow
end;

fprintf('\n 3 Pak ### Calculate all possible 1-4 models and list - press a key\n');pause

for i=1:37,
   fprintf(' %2i  [%i %i %i]  %f \n',i,list(i,:));
end;

fprintf('\n 4 Pak ### Calculate all possible 1-4 models and list sorted and plot - press a key\n');pause

[i j]=sort(list(:,4));
lists=list(j,:);
for i=1:37,
   fprintf(' %2i  [%i %i %i]  %f \n',i,lists(i,:));
end;

figure(1);set(gcf,'Position',[-1 31 804 534]);
plot(lists(:,4));
grid on
ylabel('Explained variation');
xlabel('Model ranking');

fprintf('\n 5 Pak ### Calculate a [1 3 3] model, list core and facplot - press a key\n');pause

format short
format compact
W=[1 3 3];
load('data\dataset3res') %[Factors,G,XExpl,Xf]=tucker(X,R,W);
int2str(G)
figure(1);set(gcf,'Position',[-1 31 804 534]);

% Convert factors to new format
clear ff,id1 = 0;
for i = 1:length(DimX) 
   id2 = sum(DimX(1:i).*W(1:i));
   ff{i} = reshape(Factors(id1+1:id2),DimX(i),W(i));
   id1 = id2;
end
Factors = ff;
plotfac(Factors,[],[1 28;250 440;250 560]);drawnow;legend('1','2','3')
set(gcf,'Position',[-1 31 804 534]);

fprintf('\n 6 Pak ### Continue with a [3 3 3] model, list core and facplot - press a key\n');pause

W=[3 3 3];
disp('[3 3 3] Before rotation')
int2str(G)
figure(1);set(gcf,'Position',[-1 31 804 534]);
plotfac(Factors,[],[1 28;250 440;250 560]);legend('1','2','3')
[A B C]=fac2let(Factors);

fprintf('\n 7 Pak ### Continue with a [3 3 3] model with VOS rotation and facplot  - press a key\n');pause

disp('[3 3 3] After rotation to optimal variance of squares')
G = reshape(G,W);
[Gv,Ov1,Ov2,Ov3]=maxvar3(G,1e-5);
Gv = reshape(Gv,size(Gv,1),size(Gv,2)*size(Gv,3));
int2str(Gv)
Av=A*Ov1;Bv=B*Ov2;Cv=C*Ov3;SSres=sum(sum( (X-Av*Gv*kron(Cv',Bv')).^2 ));
figure(2);set(gcf,'Position',[-1 31 804 534]);
plotfac({Av;Bv;Cv},[],[1 28;250 440;250 560]);legend('1','2','3')

fprintf('\n 8 Pak ### The [3 3 3] model with DIA rotation and facplot - press a key\n');pause

disp('[3 3 3] After rotation to optimal diagonality')
[Gd,Od1,Od2,Od3]=maxdia3(G,1e-5);
Gd = reshape(Gd,size(Gd,1),size(Gd,2)*size(Gd,3));
int2str(Gd)
Ad=A*Od1;Bd=B*Od2;Cd=C*Od3;SSres=sum(sum( (X-Ad*Gd*kron(Cd',Bd')).^2 ));
figure(3);set(gcf,'Position',[-1 31 804 534]);
plotfac({Ad; Bd;Cd},[],[1 28;250 440;250 560]);legend('1','2','3')

fprintf('\n 9 Pak ### Compare cores  - press a key\n');pause

fprintf('\nNot rotated - Fig 1\n');disp(int2str(G));
fprintf('\nVOS rotated - Fig 2\n');disp(int2str(Gv));
fprintf('\nDIA rotated - Fig 3\n');disp(int2str(Gd));

format
```
