## Supplementary File S1 for "Robust latent-variable interpretation of *in vivo* regression models by nested resampling": tucker.m

```
function [Factors,G,ExplX,Xm]=tucker(X,Fac,Options,ConstrF,ConstrG,Factors,G);
%TUCKER multi-way tucker model
%
% function [Factors,G,ExplX,Xm]=tucker(X,Fac[,Options[,ConstrF,[ConstrG[,Factors[,G]]]]]);
%
% Change: True LS unimodality now supported.
%
% This algorithm requires access to:
% 'fnipals' 'gsm' 'inituck' 'calcore' 'nmodel' 'nonneg' 'setopts' 'misssum'
% 'missmean' 't3core'
%
% See also:
% 'parafac' 'maxvar3' 'maxdia3' 'maxswd3'
%
% ---------------------------------------------------------
%             The general N-way Tucker model
% ---------------------------------------------------------
%
% [Factors,G,ExplX,Xm]=tucker(X,Fac,Options,ConstrF,ConstrG,Factors,G);
% [Factors,G,ExplX,Xm]=tucker(X,Fac);
%
% INPUT
% X        : The multi-way data array.
% Fac      : Row-vector describing the number of factors
%            in each of the N modes. A '-1' (minus one)
%            will tell the algorithm not to estimate factors
%            for this mode, yielding a Tucker2 model.
%            Ex. [3 2 4]
%
% OPTIONAL INPUT
% Options  : See parafac.
% ConstrF  : Constraints that must apply to 'Factors'.
%            Define a row-vector of size N that describes how
%            each mode should be treated.
%            '0' orthogonality (default)
%            '1' non-negativity
%            '2' unconstrained
%            '4' unimodality and non-negativity.
%            E.g.: [0 2 1] yields ortho in first mode, uncon in the second
%            and non-neg in the third mode.
%            Note: The algorithm uses random values if there are no
%            non-negative components in the iteration intermediates. Thus,
%            if non-negativity is applied, the iterations may be
%            non-monotone in minor sequences.
% ConstrG  : Constraints that must apply to 'G'.
%            '[]' or '0' will not constrain the elements of 'G'.
%            To define what core elements should be allowed, give a core that
%            is 1 (one) on all active positions and zero elsewhere - this boolean
%            core array must have the same dimensions as defined by 'Fac'.
%
% OUTPUT
% Factors  : A row-vector containing the solutions.
% G        : Core array that matches the dimensions defined by 'Fac'.
% ExplX    : Fraction of variation (sums of squares explained)
% Xm       : Xhat (the model of X)
%
% This algorithm applies to the general N-way case, so
% the array X can have any number of dimensions. The
% principles of 'projections' and 'systematic unfolding
% methodology (SUM)' are used in this algorithm to provide
% a fast approach - also for larger data arrays. This
% algorithm can handle missing values if denoted
% by NaN's. It can also be used to make TUCKER2/1 models by
% properly setting the elements of 'Fac' to -1.
%
% Note: When estimating a Tucker model on data using non-orthogonal factors,
%       the sum of square of the core may differ between models of the
%       same dataset. This is in order since the factors may
%       thus be correlated. However, the expl. var. should always be the same.
%

% $ Version 2.01  $ May 2011 $ Fixed problem with Tucker2 calculations & missing 'Option'$ CA $ Not compiled $
% $ Version 2.003 $ Jan 2002 $ Fixed problem with length of factors under special conditions $ CA $ Not compiled $
% $ Version 2.002 $ Jan 2002 $ Fixed reshaping of old input G $ RB $ Not compiled $
% $ Version 2.001 $ July 2001 $ Changed problem with checking if Factors exist (should check if it exists in workspace specifically)$ RB $ Not compiled $
% $ Version 2.00  $ May 2001 $ Changed to array notation $ RB $ Not compiled $
% $ Version 1.12  $ Date 14. Nov. 1999 $ Not compiled $
%
%
% Copyright, 1998-, 2011-
%
% Claus A. Andersson
% Chemometrics Group - Life Sciences
% University of Copenhagen
% Rolighedsvej 30, DK-1958 Frederiksberg, Denmark
% //
%

DimX = size(X);
X = reshape(X,DimX(1),prod(DimX(2:end)));
FacNew = Fac;
FacNew(find(FacNew==-1)) = DimX(find(FacNew==-1));

format long
format compact
dbg=0;

if nargin==0,
    help('tucker.m');
    error(['Error calling ''tucker.m''. Since no input arguments were given, the ''help'' command was initiated.'])
    return;
end;
if nargin<2,
    help('tucker.m');
    error(['Error calling ''tucker.m''. At least two (2) input arguments must be given. Read the text above.'])
    return;
end;
if size(Fac,2)==1,
    help('tucker.m');
    error(['Error calling ''tucker.m''. ''Fac'' must be a row-vector.'])
end;

% Initialize system variables
N=size(Fac,2);
Fac_orig=Fac;
finda=find(Fac==-1);
if ~isempty(finda),
    Fac(finda)=zeros(size(finda));
end;
FIdx0=cumsum([1 DimX(1:N-1).*Fac(1:N-1)]);
FIdx1=cumsum([DimX.*Fac]);
pmore=30;
pout=0;
Xm=[];
MissingExist=any(isnan(X(:)));
if MissingExist,
    IdxIsNans=find(isnan(X));
end;
SSX=misssum(misssum(X.^2));

if exist('Options')~=1,
    Options_=[0];
else
    Options_=Options;
end;
load noptiot3.mat;
i=find(Options_);
Options(i)=Options_(i);
if isnan(Options(5)),
    prlvl = 0;
else
    prlvl = 1;
end;
Options12=Options(1);
Options11=Options12*10;
Options21=Options(2);
Options31=Options(3);
Options41=Options(4);
Options51=Options(5);
Options61=Options(6);
Options71=Options(7);
Options81=Options(8);
Options91=Options(9);
Options101=Options(10);

if ~exist('ConstrF'),
    ConstrF=[];
end;
if isempty(ConstrF),
    ConstrF=zeros(size(DimX));
end;
if ConstrF==0 ,
    ConstrF=zeros(size(DimX));
end;

if ~exist('ConstrG')
    ConstrG=[];
end;
if isempty(ConstrG),
    ConstrG=0;
end;

if exist('Factors')~=1,
    Factors=[];
end;

if ~(exist('G')==1),
    G=[];
else
    G=reshape(G,size(G,1),prod(size(G))/size(G,1));
end;

%Give a status/overview
if prlvl>0,
    fprintf('\n\n');
    fprintf('=================   RESUME  &  PARAMETERS   ===================\n');
    fprintf('Array                 : %i-way array with dimensions (%s)\n',N,int2str(DimX));
    if any(Fac==0),
        fprintf('Model                 : (%s) TUCKER2 model\n',int2str(Fac));
    else
        fprintf('Model                 : (%s) TUCKER3 model\n',int2str(Fac));
    end;
end

%Mth initialization
txt1=str2mat('derived by SVD (orthogonality constrained).');
txt1=str2mat(txt1,'derived by NIPALS (orthogonality constrained).');
txt1=str2mat(txt1,'derived by Gram-Schmidt (orthogonality constrained).');
txt1=str2mat(txt1,'This mode is not compressed/calculated, i.e., TUCKER2 model.');
txt1=str2mat(txt1,'derived by non-negativity least squares.');
txt1=str2mat(txt1,'derived by unconstrained simple least squares.');
txt1=str2mat(txt1,'unchanged, left as defined in input ''Factors''.');
txt1=str2mat(txt1,'derived by unimodality constrained regression.');
MethodO=1;
for k=1:N,
    UpdateCore(k)=1;
    if ConstrF(k)==0,
        if Fac(k)>0,
            if 0<DimX(k) & DimX(k)<=180,
                Mth(k)=1;
            end;
            if 180<DimX(k) & DimX(k)<=Inf,
                Mth(k)=3;
            end;
            if Fac(k)<=6 & 180<DimX(k),
                Mth(k)=2;
            end;
        end;
        UpdateWithPinv(k)=1; %Update with the L-LS-P-w/Kron approach
        CalcOrdinar(k)=1;
    end;
    if ConstrF(k)==1,
        Mth(k)=5; %nonneg
        MethodO=2; %use the flexible scheme
        UpdateCore(k)=1; %Update the core in this mode
        CalcOrdinar(k)=1;
    end;
    if ConstrF(k)==2,
        Mth(k)=6; %uncon
        MethodO=2; %use the flexible scheme
        UpdateCore(k)=1; %Update the core in this mode
        CalcOrdinar(k)=1;
    end;
    if ConstrF(k)==3,
        Mth(k)=7; %unchanged
        MethodO=2;
        UpdateCore(k)=1; %Update the core in this mode
        CalcOrdinar(k)=1;
    end;
    if ConstrF(k)==4,
        Mth(k)=8; %unimod
        MethodO=2; %use the flexible scheme
        UpdateCore(k)=1; %Update the core in this mode
        CalcOrdinar(k)=1;
    end;
    if Fac_orig(k)==-1
        Mth(k)=4;
        UpdateCore(k)=0; %Do not update core for this mode
        CalcOrdinar(k)=1;
    end;
    if Options91>=1,
        if prlvl>0,
            if Mth(k)~=4,
                fprintf('Mode %i                : %i factors %s\n',k,Fac(k),txt1(Mth(k),:));
            else
                fprintf('Mode %i                : %s\n',k,txt1(Mth(k),:));
            end;
        end;
    end;
end;

UserFactors=1;
if isempty(Factors),
    UserFactors=0;
else
    ff = [];
    for f=1:length(Factors)
        if ~all(size(Factors{f})==[DimX(f),Fac(f)]), %%
            Factors{f}=rand(DimX(f),Fac(f));%% Added by CA, 27-01-2002
        end;%%
        ff=[ff;Factors{f}(:)];
    end
    OrigFactors=Factors;
    Factors = ff;
end;

usefacinput=0;
if MissingExist,
    if ~UserFactors
        [i j]=find(isnan(X));
        mnx=missmean(X)/3;
        mny=missmean(X')/3;
        n=size(i,1);
        for k=1:n,
            i_=i(k);
            j_=j(k);
            X(i_,j_) = mny(i_) + mnx(j_);
        end;
        mnz=(missmean(mnx)+missmean(mny))/2;
        p=find(isnan(X));
        X(p)=mnz;
    else
        usefacinput=1;
        % Convert to new format
        clear ff,id1 = 0;
        for i = 1:length(DimX)
            id2 = sum(DimX(1:i).*Fac(1:i));ff{i} = reshape(Factors(id1+1:id2),DimX(i),Fac(i));id1 = id2;
        end
        Fact = ff;
        Xm=nmodel(Fact,reshape(G,Fac_orig));
        Xm = reshape(Xm,DimX(1),prod(DimX(2:end)));
        X(IdxIsNans)=Xm(IdxIsNans);
    end;
    SSMisOld=sum(sum( X(IdxIsNans).^2 ));
    SSMis=SSMisOld;
end;

% Initialize the Factors by some method
UserFactors=1;
if isempty(Factors),
    Factors=inituck(reshape(X,DimX),Fac_orig,2,[]);
    
    % Convert to old factors
    ff = [];
    for f=1:length(Factors)
        ff=[ff;Factors{f}(:)];
    end
    Factors = ff;
    UserFactors=0;
end;

% Initialize the core
Core_uncon=0;
Core_nonneg=0;
Core_cmplex=0;
Core_const=0;
G_cons=ConstrG;
if all(ConstrG(:)==1),
    ConstrG=0;
end;
if ConstrG==0,
    % Convert to new format
    clear ff,id1 = 0;
    for i = 1:length(DimX)
        id2 = sum(DimX(1:i).*Fac(1:i));ff{i} = reshape(Factors(id1+1:id2),DimX(i),Fac(i));id1 = id2;
    end
    Fact = ff;
    G=calcore(reshape(X,DimX),Fact,[],1,MissingExist);
    G = reshape(G,size(G,1),prod(size(G))/size(G,1));
    Core_uncon=1;
elseif prod(size(ConstrG)==[Fac(1) prod(Fac(2:N))]),
    tmpM2=1;
    for k=1:N;
        if Mth(k)==4,
            tmpM1=eye(DimX(k));
        else
            tmpM1=reshape(Factors(FIdx0(k):FIdx1(k)),DimX(k),Fac(k));
        end;
        tmpM2=ckron(tmpM2,tmpM1);
    end
    w=ConstrG(:);
    fwz=find(w==1);
    fwnn=find(w==2);
    fwfix=find(w==3);
    G=zeros(prod(Fac),1);
    G(fwz)=tmpM2(:,fwz)\X(:);
    enda=size(Fac,2); %!
    G=reshape(G,Fac(1),prod(Fac(2:enda)));
    Core_cmplex=1;
    MethodO=2;
    UpdateCore=2*ones(1,N);
elseif ConstrG==2,
    % Convert to new format
    clear ff,id1 = 0;
    for i = 1:length(DimX)
        id2 = sum(DimX(1:i).*Fac(1:i));ff{i} = reshape(Factors(id1+1:id2),DimX(i),Fac(i));id1 = id2;
    end
    Fact = ff;
    G=calcore(reshape(X,DimX),Fact,[],1,MissingExist);
    G = reshape(G,size(G,1),prod(size(G))/size(G,1));
    G(find(G<0))=0;
    Core_nonneg=1;
elseif ConstrG==3,
    Core_const=1;
    UpdateCore=0*UpdateCore; %%%Added by CA, 27-01-2002
end;

if prlvl>0
    fprintf('Type of algorithm     : ');
    if MethodO==1,
        fprintf('Orthogonal projections.\n');
    elseif MethodO==2,
        fprintf('Flexible scheme.\n');
    end;
    fprintf('Core                  : ');
    if Core_cmplex==1 & Core_nonneg==0,
        fprintf('Unconstrained composite/sparse core.\n');
    elseif Core_cmplex==1 & Core_nonneg==1,
        fprintf('Non-negativity constrained and composite/sparse core.\n');
    elseif Core_const==1,
        fprintf('Fixed core.\n');
    else
        fprintf('Full unconstrained core.\n');
    end;
end;

if prlvl>0,
    if MissingExist,
        if Options91>=1,
            fprintf('Missing data          : Yes, 2 active loops (expectation maximization).\n');
            fprintf('                        %i values (%.2f%%) out of %i are NaNs/missing.\n',prod(size(IdxIsNans)),100*prod(size(IdxIsNans))/prod(size(X)),prod(size(X)));
            if usefacinput==0,
                fprintf('                        Missing values initialized from column and row means.\n');
            else
                fprintf('                        Missing values initialized from model based on the given input.\n');
            end;
            fprintf('Convergence crit. 1   : %.5g (relative) sum of sq. core elements (corrected for missing values).\n',Options12);
            fprintf('Convergence crit. 2   : %.5g (relative) sum of sq. of pred. missing values.\n',Options11);
            fprintf('Iteration limit       : %i is the maximum number of overall iterations.\n',Options61);
        end;
    else
        if Options91>=1,
            fprintf('Missing data          : No, 1 active loop.\n');
            fprintf('Convergence crit. 1   : %.5g (relative) sum of sq. core elements.\n',Options12);
            fprintf('Iteration limit       : %i is the maximum number of overall iterations.\n',Options61);
        end;
    end;
    fprintf('\n');
    if MissingExist,
        str1=' Iter. 1  |  Corrected sum of   |  Sum of sq. miss.  |   Expl.  ';
        str2='     #    |  sq. core elements  |       values       |  var. [%]';
        fprintf('%s\n',str1);
        fprintf('%s\n\n',str2);
    else
        str1=' Iter. 1  |       Sum of        |   Expl.  ';
        str2='     #    |  sq. core elements  |  var. [%]';
        fprintf('%s\n',str1);
        fprintf('%s\n\n',str2);
    end;
end;
Conv_true=0;
if MethodO==1, %Can use the faster projection technique
    SSGOld=0;
    Converged2=0;
    it1=0;
    itlim1=0;
    t0=clock;
    while ~Converged2,
        Converged1=0;
        while ~Converged1,
            it1=it1+1;
            % Iterate over the modes
            for c=1:N,
                %Compress the data by projections
                if Mth(c)~=4,
                    CurDimX=DimX;
                    RedData=X;
                    for k=1:N;
                        if k~=c,
                            if Mth(k)~=4,
                                kthFactor=reshape(Factors(FIdx0(k):FIdx1(k)),DimX(k),Fac(k));
                                RedData=kthFactor'*RedData;
                                CurDimX(k)=Fac(k);
                            else
                                RedData=RedData;
                            end,
                        end,
                        if k~=N,
                            newi=CurDimX(k+1);
                            newj=prod(CurDimX)/newi;
                        else
                            newi=CurDimX(1);
                            newj=prod(CurDimX)/newi;
                        end;
                        RedData=reshape(RedData',newi,newj);
                    end;
                    %Reshape to the proper unfolding
                    for k=1:(c-1);
                        if k~=c,
                            newi=CurDimX(k+1);
                            newj=prod(CurDimX)/CurDimX(k+1);
                        else,
                            newi=CurDimX(1);
                            newj=prod(CurDimX)/CurDimX(1);
                        end;
                        RedData=reshape(RedData',newi,newj);
                    end;
                    %Find a basis in the projected space
                    %...using the robust SVD
                    if Mth(c)==1,
                        if MissingExist,
                            [U S V]=svd(RedData',0);
                            cthFactor=V(:,1:Fac(c));
                        else
                            [U S V]=svd(RedData',0);
                            cthFactor=V(:,1:min(Fac(c),size(V,2)));
                            if size(cthFactor,2)<Fac(c)
                                cthFactor = [cthFactor rand(size(cthFactor,1),Fac(c)-size(cthFactor,2))];
                            end
                        end;
                    end;
                    %...using the fast NIPALS
                    if Mth(c)==2,
                        if MissingExist,
                            [cthFactor]=fnipals(RedData,Fac(c),reshape(Factors(FIdx0(c):FIdx1(c)),DimX(c),Fac(c)));
                        else
                            [cthFactor]=fnipals(RedData,Fac(c),reshape(Factors(FIdx0(c):FIdx1(c)),DimX(c),Fac(c)));
                        end;
                    end;
                    %...using simplified continuous Gram-Schmidt orthogonalization
                    if Mth(c)==3,
                        if MissingExist,
                            TempMat=RedData*RedData';
                            cthFactor=reshape(Factors(FIdx0(c):FIdx1(c)),DimX(c),Fac(c));
                            for i=1:2,
                                [cthFactor]=gsm(TempMat*cthFactor);
                            end;
                        else
                            TempMat=RedData*RedData';
                            cthFactor=reshape(Factors(FIdx0(c):FIdx1(c)),DimX(c),Fac(c));
                            for i=1:2,
                                [cthFactor]=gsm(TempMat*cthFactor);
                            end;
                        end;
                    end;
                    %...this is void (no compression for this mode)
                    if Mth(c)==4,
                    end;
                    %...this is void (Keep factors unchanged)
                    if Mth(c)==7,
                    end;
                    %Update the 'Factors' with the current estimates
                    if Mth(c)~=4 & Mth(c)~=7
                        Factors(FIdx0(c):FIdx1(c))=cthFactor(:)';
                    end;
                end;
            end;
            
            if ~Core_const & Core_uncon==1 & Core_nonneg==0,
                % Convert to new format
                clear ff,id1 = 0;
                for i = 1:length(DimX)
                    id2 = sum(DimX(1:i).*Fac(1:i));ff{i} = reshape(Factors(id1+1:id2),DimX(i),Fac(i));id1 = id2;
                end
                Fact = ff;
                G=calcore(reshape(X,DimX),Fact,[],1,MissingExist);
                G = reshape(G,size(G,1),prod(size(G))/size(G,1));
            elseif Core_nonneg==1,
                g=T3core(reshape(X,DimX),Fac,Factors(:),0,1);
                G=reshape(g,Fac(1),prod(Fac(2:N)));
            else
                tmpM2=1;
                for k=1:N;
                    if Mth(k)==4,
                        tmpM1=eye(DimX(k));
                    else
                        tmpM1=reshape(Factors(FIdx0(k):FIdx1(k)),DimX(k),Fac(k));
                    end;
                    tmpM2=ckron(tmpM2,tmpM1);
                end
                G=G(:);
                G(fwz)=tmpM2(:,fwz)\X(:);
                enda=size(Fac,2);
                G=reshape(G,Fac(1),prod(Fac(2:enda)));
            end;
            
            SSG=sum(sum(G.^2));
            if MissingExist,
                SSG=SSG-SSMis;
            end;
            if abs(SSG-SSGOld)<Options12*SSGOld,
                Converged1=1;
            end;
            if it1>=Options61,
                itlim1=1;
                Converged1=1;
                Converged2=1;
            end;
            SSGOld=SSG;
            js=0;
            %Save on time count
            if Options101>0 & (etime(clock,t0)>Options101),
                save('temp.mat','Factors','G','DimX','Fac');
                t0=clock;
                js=1;
            end;
            %Save on iteration count
            %if (Options101<0) & (mod(it1,abs(Options101))==0),
            keval=it1/Options51;
            if (Options101<0) & ( abs( keval - floor(keval) ) <=eps),
                save('temp.mat','Factors','G','DimX','Fac');
                js=1;
            end;
            %if mod(it1,Options51)==0 | it1==1  | js==1,
            keval=it1/Options51;
            if (abs( keval - floor(keval) ) <=eps) | it1==1  | js==1, %Matlab 4.2 comp.
                % Convert to new format
                clear ff,id1 = 0;
                for i = 1:length(DimX)
                    if Fac(i)
                        id2 = sum(DimX(1:i).*Fac(1:i).*(Fac(1:i)~=0));
                        ff{i} = reshape(Factors(id1+1:id2),DimX(i),Fac(i));id1 = id2;
                    else
                        ff{i}=[];
                    end
                end
                Fact = ff;
                Xm=nmodel(Fact,reshape(G,FacNew));
                Xm = reshape(Xm,DimX(1),prod(DimX(2:end)));
                
                if MissingExist,
                    X(IdxIsNans)=Xm(IdxIsNans);
                    SSMis=sum(sum( Xm(IdxIsNans).^2 ));
                    if abs(SSMis-SSMisOld)<Options11*SSMisOld,
                        Converged2=1;
                    end;
                    SSMisOld=SSMis;
                else
                    Converged2=1;
                end;
                ExplX=100*(1-sum(sum((X-Xm).^2))/SSX);
                pout=pout+1;
                if pout>pmore,
                    if prlvl > 0,
                        fprintf('%s\n',str1);
                        fprintf('%s\n',str2);
                    end;
                    pout=0;
                end;
                
                if prlvl>0,
                    if MissingExist,
                        fprintf(' %6i       %14.3f     %14.3f         %8.4f',it1,SSG,SSMis,ExplX);
                    else
                        fprintf(' %6i        %14.3f      %8.4f',it1,SSG,ExplX);
                    end;
                    if js,
                        fprintf(' - saved to ''temp.mat'' \n')
                    else
                        fprintf('\n')
                    end;
                end;
            end;
        end; %Inner loop
    end; %Outer loop
    if prlvl>0,
        if itlim1==0,
            fprintf('   Stopped. Convergence criteria reached.\n');
        else
            fprintf('   Stopped. Iteration limits reached in model and expectation loops.\n');
        end;
        if MissingExist,
            fprintf(' %6i       %14.3f     %14.3f         %8.4f',it1,SSG,SSMis,ExplX);
        else
            fprintf(' %6i        %14.3f      %8.4f',it1,SSG,ExplX);
        end;
    end;
    if Options101~=0,
        save('temp.mat','Factors','G','DimX','Fac');
        if prlvl>0,
            fprintf(' - saved to ''temp.mat'' \n')
        end;
    else
        if prlvl>0,
            fprintf('\n')
        end;
    end;
    
elseif MethodO==2, %Must use slower but more general schemes
    
    SSGOld=0;
    OldFactors=0*Factors;
    Converged2=0;
    it1=0;
    t0=clock;
    itlim1=0;
    while ~Converged2,
        Converged1=0;
        while ~Converged1,
            it1=it1+1;
            %Iterate over the modes
            for c=1:N,
                faclist1=[N:-1:1 N:-1:1];
                faclist=faclist1(N-c+2:N-c+2+(N-2));
                tmpM2=1;
                tmpM2Pinv=1;
                for k=1:N-1;
                    if Mth(faclist(k))==4,
                        tmpM1=eye(DimX(faclist(k)));
                    else
                        tmpM1=reshape(Factors(FIdx0(faclist(k)):FIdx1(faclist(k))),DimX(faclist(k)),Fac(faclist(k)));
                    end;
                    if CalcOrdinar(c)==1,
                        tmpM2=ckron(tmpM2,tmpM1');
                    end
                end
                %Estimate Factors for the cth way
                %...this is void (no compression for this mode)
                if any(Mth(c)==[1 2 3]),
                    tmpM4=G*tmpM2;
                    if MissingExist,
                        %cthFactor=(X*tmpM4')/((tmpM4*X'*X*tmpM4')^(1/2));
                        [U S V]=svd(tmpM4*X'*X*tmpM4',0);
                        mS=min(size(S));
                        Sm=S;
                        Sm(1:mS,1:mS)=diag(1./sqrt(diag(S(1:mS,1:mS))));
                        pinvm=U*Sm*V';
                        cthFactor=X*tmpM4'*pinvm;
                    else
                        %cthFactor=(X*tmpM4')/((tmpM4*X'*X*tmpM4')^(1/2));
                        [U S V]=svd(tmpM4*X'*X*tmpM4',0);
                        mS=min(size(S));
                        Sm=S;
                        Sm(1:mS,1:mS)=diag(1./sqrt(diag(S(1:mS,1:mS))));
                        pinvm=U*Sm*V';
                        cthFactor=X*tmpM4'*pinvm;
                    end;
                end;
                if Mth(c)==4,
                end;
                if Mth(c)==5, %Nonneg
                    tmpM3=reshape(Factors(FIdx0(c):FIdx1(c)),DimX(c),Fac(c));
                    if it1==1,
                        %tmpM3(find(tmpM3<0))=0;
                        i__=find(tmpM3<0);
                        if ~isempty(i__),
                            tmpM3(i__)=zeros(1,length(i__));
                        end;
                    end;
                    if CalcOrdinar(c) == 1,
                        tmpM4=G*tmpM2;
                        if dbg, ss1=sum(sum( (X-tmpM3*tmpM4).^2 ));end;
                        [cthFactor,SS]=nonneg(X,tmpM4,tmpM3);
                        if dbg, ss2=sum(sum( (X-cthFactor*tmpM4).^2 ));
                            fprintf('Nonng report (Ordi) %15.8d  %15.8d\n',ss1,ss2); end;
                    end;
                end;
                if Mth(c)==6, %Uncon
                    cthFactor=reshape(Factors(FIdx0(c):FIdx1(c)),DimX(c),Fac(c));
                    if CalcOrdinar(c) == 1,
                        M_=G*tmpM2;
                        if dbg, ss1=sum(sum( (X-cthFactor*G*tmpM2).^2 ));end;
                        cthFactor=X/M_;
                        if dbg, ss2=sum(sum( (X-cthFactor*G*tmpM2).^2 ));
                            fprintf('Uncon report (Ordi) %15.8d  %15.8d\n',ss1,ss2);end;
                    end;
                end;
                if Mth(c)==7, %Not updated
                end;
                if Mth(c)==8, %Unimodality
                    tmpM3=reshape(Factors(FIdx0(c):FIdx1(c)),DimX(c),Fac(c));
                    if it1==1,
                        %tmpM3(find(tmpM3<0))=0;
                        i__=find(tmpM3<0);
                        if ~isempty(i__),
                            tmpM3(i__)=zeros(1,length(i__));
                        end;
                    end;
                    if CalcOrdinar(c) == 1,
                        tmpM4=G*tmpM2;
                        if dbg, ss1=sum(sum( (X-tmpM3*tmpM4).^2 ));end;
                        cthFactor = unimodal(tmpM4',X');
                        if dbg, ss2=sum(sum( (X-cthFactor*tmpM4).^2 ));fprintf('Unimod report (Ordi) %15.8d  %15.8d\n',ss1,ss2); end;
                    end;
                end;
                
                
                %Update 'Factors' with the current estimates
                if Mth(c)~=4 & Mth(c)~=7
                    cn=sum(cthFactor.^2);
                    for i=1:Fac(c);
                        if it1<=10 & cn(i)<eps,
                            cthFactor(:,i)=rand(size(cthFactor,1),1);
                        end;
                    end;
                    if Core_const,
                        if c>1
                            cthFactor=normit(cthFactor);
                        end;
                    else
                        cthFactor=normit(cthFactor);
                    end;
                    Factors(FIdx0(c):FIdx1(c))=cthFactor(:)';
                end;
                
                %Estimate the new core after each SUBiteration
                CoreupdatedInner=0;
                if ~Core_const
                    if UpdateCore(c)==1,
                        tmpM3=reshape(Factors(FIdx0(c):FIdx1(c)),DimX(c),Fac(c));
                        if CalcOrdinar(c) == 1,
                            if dbg, ss1=sum(sum( (X-tmpM3*G*tmpM2).^2 ));end;
                            G=tmpM3\(X/tmpM2);
                            if dbg, ss2=sum(sum( (X-tmpM3*G*tmpM2).^2 ));
                                fprintf('Core report  (Ordi) %15.8d  %15.8d\n',ss1,ss2);end;
                        end;
                        CoreupdatedInner=1;
                    end;
                    if UpdateCore(c)==2,
                        G=zeros(Fac(c),prod(Fac(nsetdiff(1:N,c))));
                        tmpM2=1;
                        for k=[(c-1):-1:1 N:-1:c];
                            if Mth(k)==4,
                                tmpM1=eye(DimX(k));
                            else
                                tmpM1=reshape(Factors(FIdx0(k):FIdx1(k)),DimX(k),Fac(k));
                            end;
                            tmpM2=ckron(tmpM2,tmpM1);
                        end
                        G=G(:);
                        fwz=find(G_cons==1);
                        G(fwz)=tmpM2(:,fwz)\X(:);
                        G=reshape(G,Fac(c),prod(Fac(nsetdiff(1:N,c))));
                        CoreupdatedInner=1;
                    end;
                end;
                
                %Reshape to the next unfolding
                if c~=N,
                    newix=DimX(c+1);
                    newjx=prod(DimX)/DimX(c+1);
                    newig=FacNew(c+1);
                    newjg=prod(FacNew)/FacNew(c+1);
                else
                    newix=DimX(1);
                    newjx=prod(DimX)/DimX(1);
                    newig=FacNew(1);
                    newjg=prod(FacNew)/FacNew(1);
                end;
                X=reshape(X',newix,newjx);
                G=reshape(G',newig,newjg);
                if length(G_cons(:))>1,
                    G_cons=reshape(G_cons',newig,newjg);
                end;
            end;
            
            %Estimate the new core after each MAIN iteration
            if CoreupdatedInner==0,
                if ~Core_const & Core_cmplex==0,
                    if Core_nonneg==1,
                        g=t3core(reshape(X,DimX),Fac,Factors,0,1);
                        G=reshape(g,Fac(1),prod(Fac(2:N)));
                    elseif Core_nonneg==0,
                        % Convert to new format
                        clear ff,id1 = 0;
                        for i = 1:length(DimX)
                            id2 = sum(DimX(1:i).*Fac(1:i));ff{i} = reshape(Factors(id1+1:id2),DimX(i),Fac(i));id1 = id2;
                        end
                        Fact = ff;
                        G=calcore(reshape(X,DimX),Fact,[],0,MissingExist);
                        G = reshape(G,size(G,1),prod(size(G))/size(G,1));
                    end
                end;
            end;
            CoreupdatedInner=0;
            
            if ~Core_const
                SSG=sum(sum(G.^2));
                if MissingExist,
                    SSG=SSG-SSMis;
                end;
                if abs(SSG-SSGOld)<Options12*SSGOld,
                    Converged1=1;
                end;
                if it1>=Options61,
                    Converged1=1;
                    Converged2=1;
                    itlim1=1;
                end;
                SSGOld=SSG;
            else
                SSF=sum(sum((Factors-OldFactors).^2));
                if SSF<Options12*sum(sum(Factors.^2));
                    Converged1=1;
                end;
                if it1>=Options61,
                    Converged1=1;
                    Converged2=1;
                    itlim1=1;
                end;
                OldFactors=Factors;
                SSG=SSF;
            end;
            
            js=0;
            if Options101>0 & (etime(clock,t0)>Options101),
                save('temp.mat','Factors','G','DimX','Fac');
                t0=clock;
                js=1;
            end;
            %if Options101<0 & mod(it1,abs(Options101))==0,
            keval=it1/Options101;
            if (Options101<0) & (abs( keval - floor(keval) ) <=eps), %Matlab 4.2 comp.
                save('temp.mat','Factors','G','DimX','Fac');
                js=1;
            end;
            %if mod(it1,Options51)==0 | it1==1 | js==1,
            keval=it1/Options51;
            if (abs( keval - floor(keval) ) <=eps) | it1==1  | js==1, %Matlab 4.2 comp.
                % Convert to new format
                clear ff
                id1 = 0;
                for i = 1:length(DimX)
                    if Fac(i)>0
                        id2 = sum(DimX(1:i).*Fac(1:i));
                        ff{i} = reshape(Factors(id1+1:id2),DimX(i),FacNew(i));
                        id1 = id2;
                    end
                end
                Fact = ff;
                Xm = nmodel(Fact,reshape(G,FacNew));
                Xm = reshape(Xm,DimX(1),prod(DimX(2:end)));
                if MissingExist,
                    X(IdxIsNans)=Xm(IdxIsNans);
                    SSMis=sum(sum( Xm(IdxIsNans).^2 ));
                    if abs(SSMis-SSMisOld)<Options11*SSMisOld,
                        Converged2=1;
                    end;
                    SSMisOld=SSMis;
                else
                    Converged2=1;
                end;
                ExplX=100*(1 - sum(sum((X-Xm).^2))/SSX );
                pout=pout+1;
                if pout>pmore,
                    if prlvl>0,
                        fprintf('%s\n',str1);
                        fprintf('%s\n',str2);
                    end;
                    pout=0;
                end;
                if prlvl>0,
                    if MissingExist,
                        fprintf(' %6i       %14.3f     %14.3f         %8.4f',it1,SSG,SSMis,ExplX);
                    else
                        fprintf(' %6i        %14.3f      %8.4f',it1,SSG,ExplX);
                    end;
                    if js,
                        fprintf(' - saved to ''temp.mat'' \n')
                    else
                        fprintf('\n')
                    end;
                end;
            end;
        end; %Inner loop
    end; %Outer loop
    if prlvl>0,
        if itlim1==0,
            fprintf('   Stopped. Convergence criteria reached.\n');
        else
            fprintf('   Stopped. Iteration limits reached in model and expectation loops.\n');
        end;
        if MissingExist,
            fprintf(' %6i       %14.3f     %14.3f         %8.4f',it1,SSG,SSMis,ExplX);
        else
            fprintf(' %6i        %14.3f      %8.4f',it1,SSG,ExplX);
        end;
    end;
    if Options101~=0,
        save('temp.mat','Factors','G','DimX','Fac');
        if prlvl>0,
            fprintf(' - saved to ''temp.mat'' \n')
        end;
    else
        if prlvl>0,
            fprintf('\n')
        end;
    end;
end; %Outer loop

if MissingExist,
    Xf=Xm(IdxIsNans);
end;

Factors=Factors';
format
Xm = reshape(Xm,DimX);
G = reshape(G,FacNew);

% Convert to new format
clear ff
id1 = 0;
for i = 1:length(DimX)
    if Fac(i)
        id2 = sum(DimX(1:i).*Fac(1:i));
        ff{i} = reshape(Factors(id1+1:id2),DimX(i),Fac(i));
        id1 = id2;
    else
        ff{i}=[];
    end
end

Factors = ff;
```
