## Supplementary File S1 for "Robust latent-variable interpretation of *in vivo* regression models by nested resampling": tucktest.m

```
function [fit,label] = tucktest(varargin)

%TUCKTEST makes a scree-plot for Tucker using from [1 1 1] to Fac models
% 
% INPUT
% Same as Tucker
% 
% OUTPUT
% fit      Vector containing the fit of the individual models
% label    Number of components of the corresponding element of fit
%
% USAGE
% [fit,label] = tucktest(X,Fac[,Options[,ConstrF,[ConstrG[,Factors[,G]]]]]);
%
% Alternatively, the output from ncrossdecomp.m may be visualized with
% tucktest using "tucktest(XvalResult);"

% Copyright (C) 1995-2006  Rasmus Bro & Claus Andersson
% Copenhagen University, DK-1958 Frederiksberg, Denmark,
%

% $ Version 2.1 $ Jan 2004 $ Fixed error for 4 and higherway models $ RB $ Not compiled $


if nargin>1 % Then the input is the struct from cross-validation and no fitting is performed
  
  Fac = varargin{2};
  order = length(size(varargin{1}));
  
  if order>10
    error('TUCKTEST only works up to ten-way arrays')
  end
  
  
  PossibleNumber = [1:max(Fac)]'*ones(1,order);
  possibleCombs = unique(nchoosek(PossibleNumber(:),order),'rows');
  for i=1:order
    j = find(possibleCombs(:,i)>Fac(i));
    possibleCombs(j,:)=[];
  end
  p=0;
  
  if length(varargin)>2
    Opt = varargin{3};
    Opt(5)=NaN;
  else
    Opt(5) = NaN;
  end
  varargin{3}=Opt;
  
  for i = 1:size(possibleCombs,1)
    if prod(possibleCombs(i,:))/max(possibleCombs(i,:))>=max(possibleCombs(i,:)),
      p=p+1;
      label(p,:) = possibleCombs(i,:);
      disp([' Fitting model ',num2str(label(p,:))])
      if length(varargin)<4
        [Factors,G,fit(p)]=tucker(varargin{1},label(p,:),varargin{3});
      elseif length(varargin)<5
        [Factors,G,fit(p)]=tucker(varargin{1},label(p,:),varargin{3},varargin{4});
      elseif length(varargin)<6
        [Factors,G,fit(p)]=tucker(varargin{1},label(p,:),varargin{3},varargin{4},varargin{5});
      elseif length(varargin)<7
        [Factors,G,fit(p)]=tucker(varargin{1},label(p,:),varargin{3},varargin{4},varargin{5},varargin{6});
      else
        [Factors,G,fit(p)]=tucker(varargin{1},label(p,:),varargin{3},varargin{4},varargin{5},varargin{6},varargin{7});
      end
      if all(label(p,:)==[1 2*ones(1,order-1)])
        fit(p);
      end
    end  
  end

  
  
  topfit = zeros(length(fit),1);
  uniqlab = unique(sum(label'));
  for i = 1:length(uniqlab);
    j = find(sum(label')==uniqlab(i));
    [a,b] = max(fit(j));
    topfit(j(b(1))) = 1;
  end
  idtopfit = find(topfit);
  

  figure
  plot(sum(label')',fit,'o');hold on
  plot(sum(label(idtopfit,:)')',fit(idtopfit),'bo','linewidth',3,'MarkerFaceColor','r','MarkerSize',9);
  ii = sum(label(idtopfit,:)');
  [a,b]=sort(ii);
  plot(ii(b),fit(idtopfit(b)),'r--');
  hold off
  
  
  grid on;
  for i=1:length(fit(:));
    text(sum(label(i,:))+.1,fit(i),['(' num2str(label(i,:)) ')']);
  end;
  title('SSEx as function of Tucker3 model dimensionality');
  xlabel('Tucker3 model dimensionality (total number of components)');
  ylabel('Explained variation of X');
  shg
  
else % Input is struct from ncrossdecomp
  
  label = varargin{1}.Xval(:,2:end);
  fit   = varargin{1}.Xval(:,1);
  
  
  topfit = zeros(length(fit),1);
  uniqlab = unique(sum(label'));
  for i = 1:length(uniqlab);
    j = find(sum(label')==uniqlab(i));
    [a,b] = max(fit(j));
    topfit(j(b(1))) = 1;
  end
  idtopfit = find(topfit);
  
  
  figure
  plot(sum(label')',fit,'o');hold on
  plot(sum(label(idtopfit,:)')',fit(idtopfit),'bo','linewidth',3,'MarkerFaceColor','r','MarkerSize',9);
  plot(sum(label(idtopfit,:)')',varargin{1}.Fit(idtopfit,1),'go','linewidth',3,'MarkerSize',9);  
  ii = sum(label(idtopfit,:)');
  [a,b]=sort(ii);
  plot(ii(b),fit(idtopfit(b)),'r--');
  hold off
  
  
  grid on;
  for i=1:length(fit(:));
    text(sum(label(i,:))+.1,fit(i),['(' num2str(label(i,:)) ')']);
  end;
  title('Cross-validated SSEx as function of Tucker3 model dimensionality');
  xlabel('Tucker3 model dimensionality (total number of components)');
  ylabel('Explained variation of X (green are fitted versions of blue/red ones)');
  shg
  
end
```
