## Supplementary File S1 for "Robust latent-variable interpretation of *in vivo* regression models by nested resampling": two2n.m

```
function [IndicesN]=Two2N(DimX,Indices2);
%TWO2N Conversion of indices between unfoldings and N-way arrays
%
%
% function [IndicesN]=Two2N(DimX,Indices2);
%
%
% This algorithm requires access to:
% ''
%
% ---------------------------------------------------------
%                Conversion of indices
%         between unfoldings and N-way arrays
% ---------------------------------------------------------
%
% [IndicesN]=Two2N(DimX,Indices2);
%
% DimX     : Dimensions of the N-way array.
% Indices2 : Indices in the unfolded 2-way array.
% indicesN : Indices in the N-way array
%
% This function helps you resolve the correct N-way indices
% to an entry in an unfolded array. If you e.g. want to know
% the N-way indices of the [3 240] element of a 5-way core
% with dimensions [7 6 5 8 7] you would write:
% two2n([7 6 5 8 7],[3 240]) and the answer is
% [3 6 5 8 1].

% Copyright (C) 1995-2006  Rasmus Bro & Claus Andersson
% Copenhagen University, DK-1958 Frederiksberg, Denmark,


% $ Version 0.01 $ Date 11. July 1997 $ Not compiled $

C=size(DimX,2);
a=Indices2(1);
b=Indices2(2);

if a>DimX(1),
   fprintf('two2n.m: The column index cannot be that high!\n');
   return
end;

if b>prod(DimX)/DimX(1),
   fprintf('two2n.m: The row index cannot be that high!\n');
   return
end;

Tb=b;
Par(1)=a;    
for c=C:-1:3,
   factor=prod(DimX(2:c-1));
   Par(c)=floor(Tb/factor)+1;
   if (Tb-(Par(c)-1)*factor)<1,
      Par(c)=Par(c)-1;
   end;
   Tb=Tb-(Par(c)-1)*factor;
end;
Par(2)=Tb;
IndicesN=Par;
```
