## Supplementary File S1 for "Robust latent-variable interpretation of *in vivo* regression models by nested resampling": ulsr.m

```
function [b,All,MaxML]=ulsr(x,NonNeg);

%ULSR 
%
% See also:
% 'unimodal' 'monreg' 'fastnnls'
%
% ------INPUT------
%
% x       is the vector to be approximated
% NonNeg  If NonNeg is one, nonnegativity is imposed
%
%
%
% ------OUTPUT-----
%
% b 	     is the best ULSR vector
% All      is containing in its i'th column the ULSRFIX solution for mode
% 	        location at the i'th element. The ULSR solution given in All
%          is found disregarding the i'th element and hence NOT optimal
% MaxML    is the optimal (leftmost) mode location (i.e. position of maximum)
%
% Reference
% Bro and Sidiropoulos, "Journal of Chemometrics", 1998, 12, 223-247. 
%
%
% [b,All,MaxML]=ulsr(x,NonNeg);
% This file uses MONREG.M


% Copyright (C) 1995-2006  Rasmus Bro & Nikos Sidiropoulos
% Copenhagen University, DK-1958 Frederiksberg, Denmark,
%


% $ Version 1.02 $ Date 28. July 1998 $ Not compiled $
%


x=x(:);
I=length(x);
xmin=min(x);
if xmin<0
  x=x-xmin;
end


% THE SUBSEQUENT 
% CALCULATES BEST BY TWO MONOTONIC REGRESSIONS

% B1(1:i,i) contains the monontonic increasing regr. on x(1:i)
[b1,out,B1]=monreg(x);

% BI is the opposite of B1. Hence BI(i:I,i) holds the monotonic
% decreasing regression on x(i:I)
[bI,out,BI]=monreg(flipud(x));
BI=flipud(fliplr(BI));

% Together B1 and BI can be concatenated to give the solution to
% problem ULSR for any modloc position AS long as we do not pay
% attention to the element of x at this position


All=zeros(I,I+2);
All(1:I,3:I+2)=B1;
All(1:I,1:I)=All(1:I,1:I)+BI;
All=All(:,2:I+1);
Allmin=All;
Allmax=All;
% All(:,i) holds the ULSR solution for modloc = i, disregarding x(i),


iii=find(x>=max(All)');
b=All(:,iii(1));
b(iii(1))=x(iii(1));
Bestfit=sum((b-x).^2);
MaxML=iii(1);
for ii=2:length(iii)
  this=All(:,iii(ii));
  this(iii(ii))=x(iii(ii));
  thisfit=sum((this-x).^2);
  if thisfit<Bestfit
    b=this;
    Bestfit=thisfit;
    MaxML=iii(ii);
  end
end

if xmin<0
  b=b+xmin;
end


% Impose nonnegativity
if NonNeg==1
  if any(b<0)
    id=find(b<0);
    % Note that changing the negative values to zero does not affect the
    % solution with respect to nonnegative parameters and position of the
    % maximum.
    b(id)=zeros(size(id))+0;
  end
end
```
