## Supplementary File S1 for "Robust latent-variable interpretation of *in vivo* regression models by nested resampling": unimodal.m

```
function B=unimodal(X,Y,Bold)

%UNIMODAL unimodal regression
%
% Solves the problem min|Y-XB'| subject to the columns of 
% B are unimodal and nonnegative. The algorithm is iterative
% If an estimate of B (Bold) is given only one iteration is given, hence
% the solution is only improving not least squares
% If Bold is not given the least squares solution is estimated
%
% I/O B=unimodal(X,Y,Bold)
%
% Reference
% Bro and Sidiropoulos, "Journal of Chemometrics", 1998, 12, 223-247. 


% Copyright (C) 1995-2006  Rasmus Bro & Claus Andersson
% Copenhagen University, DK-1958 Frederiksberg, Denmark,


if nargin==3
   B=Bold;
   F=size(B,2);
   for f=1:F
     y=Y-X(:,[1:f-1 f+1:F])*B(:,[1:f-1 f+1:F])';
     beta=pinv(X(:,f))*y;
     B(:,f)=ulsr(beta',1);
   end
else
   F=size(X,2);
   maxit=100;
   B=randn(size(Y,2),F);
   Bold=2*B;
   it=0;
   while norm(Bold-B)/norm(B)>1e-5&it<maxit
     Bold=B;
     it=it+1;
     for f=1:F
       y=Y-X(:,[1:f-1 f+1:F])*B(:,[1:f-1 f+1:F])';
       beta=pinv(X(:,f))*y;
       B(:,f)=ulsr(beta',1);
     end
   end
   if it==maxit
     disp([' UNIMODAL did not converge in ',num2str(maxit),' iterations']);
   end
end
```
