## Supplementary File S1 for "Robust latent-variable interpretation of *in vivo* regression models by nested resampling": unimodalcrossproducts.m

```
function B=unimodalcrossproducts(XtX,XtY,Bold)

%UNIMODALCROSSPRODUCTS
% Solves the problem min|Y-XB'| subject to the columns of 
% B are unimodal and nonnegative. The algorithm is iterative and
% only one iteration is given, hence the solution is only improving 
% the current estimate
%
% I/O B=unimodalcrossproducts(XtX,XtY,Bold)
% Modified from unimodal.m to handle crossproducts in input 1999
% Reference
% Bro and Sidiropoulos, "Journal of Chemometrics", 1998, 12, 223-247. 


% Copyright (C) 1995-2006  Rasmus Bro & Claus Andersson
% Copenhagen University, DK-1958 Frederiksberg, Denmark,
%

B=Bold;
F=size(B,2);
for f=1:F
   xty = XtY(f,:)-XtX(f,[1:f-1 f+1:F])*B(:,[1:f-1 f+1:F])';
   beta=pinv(XtX(f,f))*xty;
   B(:,f)=ulsr(beta',1);
end


function [b,All,MaxML]=ulsr(x,NonNeg);

% ------INPUT------
%
% x          is the vector to be approximated
% NonNeg     If NonNeg is one, nonnegativity is imposed
%
%
%
% ------OUTPUT-----
%
% b 	     is the best ULSR vector
% All 	     is containing in its i'th column the ULSRFIX solution for mode
% 	     location at the i'th element. The ULSR solution given in All
%            is found disregarding the i'th element and hence NOT optimal
% MaxML      is the optimal (leftmost) mode location (i.e. position of maximum)
%
% ___________________________________________________________
%
%
%               Copyright 1997
%
% Nikos Sidiroupolos
% University of Maryland
% Maryland, US
%
%       &
%
% Rasmus Bro
% Royal Veterinary & Agricultural University
% Denmark
%
% 
% ___________________________________________________________


% This file uses MONREG.M

x=x(:);
I=length(x);
xmin=min(x);
if xmin<0
  x=x-xmin;
end


% THE SUBSEQUENT 
% CALCULATES BEST BY TWO MONOTONIC REGRESSIONS

% B1(1:i,i) contains the monontonic increasing regr. on x(1:i)
[b1,out,B1]=monreg(x);

% BI is the opposite of B1. Hence BI(i:I,i) holds the monotonic
% decreasing regression on x(i:I)
[bI,out,BI]=monreg(flipud(x));
BI=flipud(fliplr(BI));

% Together B1 and BI can be concatenated to give the solution to
% problem ULSR for any modloc position AS long as we do not pay
% attention to the element of x at this position


All=zeros(I,I+2);
All(1:I,3:I+2)=B1;
All(1:I,1:I)=All(1:I,1:I)+BI;
All=All(:,2:I+1);
Allmin=All;
Allmax=All;
% All(:,i) holds the ULSR solution for modloc = i, disregarding x(i),


iii=find(x>=max(All)');
b=All(:,iii(1));
b(iii(1))=x(iii(1));
Bestfit=sum((b-x).^2);
MaxML=iii(1);
for ii=2:length(iii)
  this=All(:,iii(ii));
  this(iii(ii))=x(iii(ii));
  thisfit=sum((this-x).^2);
  if thisfit<Bestfit
    b=this;
    Bestfit=thisfit;
    MaxML=iii(ii);
  end
end

if xmin<0
  b=b+xmin;
end


% Impose nonnegativity
if NonNeg==1
  if any(b<0)
    id=find(b<0);
    % Note that changing the negative values to zero does not affect the
    % solution with respect to nonnegative parameters and position of the
    % maximum.
    b(id)=zeros(size(id))+0;
  end
end

function [b,B,AllBs]=monreg(x);

% Monotonic regression according
% to J. B. Kruskal 64
%
% b     = min|x-b| subject to monotonic increase
% B     = b, but condensed
% AllBs = All monotonic regressions, i.e. AllBs(1:i,i) is the 
%         monotonic regression of x(1:i)
%
%
% Copyright 1997
%
% Rasmus Bro
% Royal Veterinary & Agricultural University
% Denmark
%
%


I=length(x);
if size(x,2)==2
   B=x;
else
   B=[x(:) ones(I,1)];
end

   AllBs=zeros(I,I);
   AllBs(1,1)=x(1);
   i=1;
   while i<size(B,1)
      if B(i,1)>B(min(I,i+1),1)
          summ=B(i,2)+B(i+1,2);
          B=[B(1:i-1,:);[(B(i,1)*B(i,2)+B(i+1,1)*B(i+1,2))/(summ) summ];B(i+2:size(B,1),:)];
          OK=1;
          while OK
             if B(i,1)<B(max(1,i-1),1)
                summ=B(i,2)+B(i-1,2);
                B=[B(1:i-2,:);[(B(i,1)*B(i,2)+B(i-1,1)*B(i-1,2))/(summ) summ];B(i+1:size(B,1),:)];
                i=max(1,i-1);
             else
                OK=0;
             end
          end
          bInterim=[];
          for i2=1:i
             bInterim=[bInterim;zeros(B(i2,2),1)+B(i2,1)];
          end
          No=sum(B(1:i,2));
          AllBs(1:No,No)=bInterim;
      else
          i=i+1;
          bInterim=[];
          for i2=1:i
             bInterim=[bInterim;zeros(B(i2,2),1)+B(i2,1)];
          end
          No=sum(B(1:i,2));
          AllBs(1:No,No)=bInterim;
      end
  end

  b=[];
  for i=1:size(B,1)
    b=[b;zeros(B(i,2),1)+B(i,1)];
 end
```
