## Supplementary File S1 for "Robust latent-variable interpretation of *in vivo* regression models by nested resampling": plsPlots_clean.m

```
%Plot PLS data
%Alex Caulk
%Yale University
%January 2019

gfac = 0.5;
fig_count = 0;

%Some helpful plotting parameters
xoff = 0.04; %offsets for plotting text
yoff = 0.04;
fnt_sz_lg = 16;
fnt_sz_sm = 12;
mrk_type = {'o','x','s','+','d'};
mrk_col = jet(length(Ytime_choice));
lin_wid = 1.5;

%Make color map for heat maps
ndiv = 50;
delt = 255/(ndiv-1);
col_map(1:ndiv,1) = [0:(delt):255]'; col_map(1:ndiv,2) = [0:(delt):255]'; col_map(1:ndiv,3) = 255;
col_map(ndiv+1:100,1) = 255; col_map(ndiv+1:100,2) = [255:-(delt):0]'; col_map(ndiv+1:100,3) = [255:-(delt):0]';
col_map = col_map/255;

if ~strcmp(nullPLS,'Yes')
    if ~strcmp(selpar,'Random')
        X_hilite = listdlg('PromptString','Select X Variables to Highlight','SelectionMode','multiple','ListString',vars(varX_choice));
        X_hilite = vars(varX_choice(X_hilite));
        Y_hilite = listdlg('PromptString','Select Y Variables to Highlight','SelectionMode','multiple','ListString',vars(varY_choice));
        Y_hilite = vars(varY_choice(Y_hilite));
    else
        X_hilite = vars(varX_choice);
        Y_hilite = vars(varY_choice);
    end
    col_options = {'red','green','blue','gray','black'};
    plot_col = listdlg('PromptString','Select Display Color','SelectionMode','Single','ListString',col_options);
    if strcmp(col_options(plot_col),'red')
        dot_col = [1 0 0];
        l_col = 'r';
    elseif strcmp(col_options(plot_col),'green')
        dot_col = [0 1 0];
        l_col = 'g';
    elseif strcmp(col_options(plot_col),'blue')
        dot_col = [0 0 1];
        l_col = 'b';
    elseif strcmp(col_options(plot_col),'gray')
        dot_col = [1 1 1]*gfac;
        l_col = 'k';
    elseif strcmp(col_options(plot_col),'black')
        dot_col = [0 0 0];
        l_col = 'k';
    end
else
    X_hilite = {};
    Y_hilite = {};
    dot_col = [0 0 0];
    l_col = 'k';
end

%--------------------------------------------------------------------------
% RMSE
%--------------------------------------------------------------------------

scale = questdlg('Plot correlations of raw or scaled data?','Scale...','Raw','Scaled','Scaled');
clear raw_data pred_data
fig_count = fig_count + 1;
figure(fig_count)
set(gcf,'Units','Normalized','Position',[0.2 0.2 0.6 0.35])

group_cols = [166 206 227;
              31 120 180;
              173 223 138;
              51 160 44]/255; %color palette used in Figs. 3b and 6b

subplot(1,2,2)
hold on

for j = 1:numsamps
    for k = 1:numtimepts
        if strcmp(scale,'Raw')
            raw_data(:,k) = Y(j,:,k);
            pred_data(:,k) = xvalypred(j,:,k);
%             pred_data(:,k) = xvalypredNs_mean(j,:,k);
        else
            raw_data(:,k) = Y_norm(j,:,k);
            pred_data(:,k) = xvalypred_pre(j,:,k);
%             pred_data(:,k) = xvalypredNs_mean(j,:,k);
        end
        
        if length(raw_data(:,k)) <= 4
            plot(raw_data(:,k),pred_data(:,k),'o','MarkerEdgeColor','k','MarkerFaceColor',group_cols(j,:),'MarkerSize',7,'LineWidth',lin_wid)
        else
            plot(raw_data(:,k),pred_data(:,k),'o','MarkerEdgeColor','k','MarkerFaceColor','k','MarkerSize',7,'LineWidth',lin_wid)
        end
    end
end

set(gca,'FontSize',fnt_sz_lg,'FontWeight','bold','LineWidth',2)
axis square
xwin = get(gca,'xlim');
ywin = get(gca,'ylim');
if strcmp(scale,'Raw')
    text(0.05*xwin(2),0.95*ywin(2),strcat('R = ',{' '},num2str(round(R,2))),'FontWeight','bold','Color','red','FontSize',fnt_sz_lg)
    text(0.05*xwin(2),0.90*ywin(2),strcat('\rho = ',{' '},num2str(round(rho,2))),'FontWeight','bold','Color','red','FontSize',fnt_sz_lg)
    xlabel('Measured Value')
    ylabel('Predicted Value')
else
    text(0.05*xwin(2),0.95*ywin(2),strcat('R = ',{' '},num2str(round(Rpre,2))),'FontWeight','bold','Color','red','FontSize',fnt_sz_lg)
    text(0.05*xwin(2),0.90*ywin(2),strcat('\rho = ',{' '},num2str(round(rhopre,2))),'FontWeight','bold','Color','red','FontSize',fnt_sz_lg)
    xlabel('Measured Value - Scaled')
    ylabel('Predicted Value - Scaled')
end
hold off

subplot(1,2,1)
plot(1:fac,Results.RMSEP,'k-o','MarkerFaceColor','white','MarkerEdgeColor','black','LineWidth',2,'MarkerSize',7)
hold on
if ~strcmp(nullPLS,'Yes') && ~strcmp(paired,'None')
    plot(1:fac,RMSEP,'r-o','MarkerFaceColor','white','MarkerEdgeColor','r','LineWidth',2,'MarkerSize',7)
%     plot(1:fac,RMSEP_mean,'b-o','MarkerFaceColor','white','MarkerEdgeColor','b','LineWidth',2,'MarkerSize',7)
    legend('Model of mean','RMSE of mean predictions','Mean of sampled RMSE vals')
    legend boxoff
end
xlabel('# Latent Variables')
xticks([1:fac])
ylabel('RMSE (%)')
box off
set(gca,'FontSize',fnt_sz_lg,'FontWeight','bold','LineWidth',2)
axis square

%--------------------------------------------------------------------------
%Generate heat map of preprocessed data
%--------------------------------------------------------------------------

fig_count = 1;
figure(fig_count)
hold on
count = 0; countX = 0; countY = 0;
Xp = [];
Yp = [];
for j = 1:length(varX_choice)
    Xp = [Xp reshape(X_norm(:,j,:),length(groupChoice),length(Xtime_choice))];
    vcX(((j-1)*length(Xtime_choice)+1):j*length(Xtime_choice)) = strcat(vars(varX_choice(j)),{' '},times(Xtime_choice));
end

Xpmin = min(min(Xp)); Xpmax = max(max(Xp));

for j = 1:length(varY_choice)
    Yp = [Yp reshape(Y_norm(:,j,:),length(groupChoice),length(Ytime_choice))];
    vcY(((j-1)*length(Ytime_choice)+1):j*length(Ytime_choice)) = strcat(vars(varY_choice(j)),{' '},times(Ytime_choice));
end

Ypmin = min(min(Yp)); Ypmax = max(max(Yp));
plmin = min(Xpmin,Ypmin); plmax = max(Xpmax,Ypmax);
plmax = max(abs(plmin),abs(plmax));
plmax = plmax(~isnan(plmax));

for j = 1:length(groupChoice)
    for i = 1:2
        clear data
        count = count+1;
        subplot(length(groupChoice),2,count)
        if i == 1
            data(:,:) = round(X_norm(j,:,:),2);
            tc = times(Xtime_choice);
            vc = vars(varX_choice);
        else
            data(:,:) = round(Y_norm(j,:,:),2);
            tc = times(Ytime_choice);
            vc = vars(varY_choice);
        end
        h = heatmap(vc,tc,data');
        set(h,'colormap',col_map,'ColorLimit',[-plmax plmax])

        if i == 1
            h.YLabel = expGroup(groupChoice(j));
        end

        if j == 1
            if i == 1
                title('Independent Data')
            else
                title('Dependent Data')
            end
        end

    end
end

%--------------------------------------------------------------------------
% X and Y time weights
%--------------------------------------------------------------------------

if length(size(X_norm)) > 2
    fig_count = fig_count + 1;
    figure(fig_count)
    set(gcf,'Position',[100 100 800 800])
    count = 0;
    for i = 1:2:fac*2-1
        count = count+1;
        subplot(fac,2,i)
        hold on
        if ~strcmp(nullPLS,'Yes') && ~strcmp(paired,'None')
            errorbar(time_val(Xtime_choice),Xt_mean(:,count),Xt_std(:,count),'LineWidth',2,'Color',dot_col)
            plot(time_val(Xtime_choice),Xfactors{3}(:,count),'k-o','LineWidth',2,'MarkerFaceColor','white')
            plot(time_val(Xtime_choice),Xt_mean(:,count),strcat(l_col,'-o'),'LineWidth',2,'MarkerFaceColor','white')
        else
            plot(time_val(Xtime_choice),Xfactors{3}(:,count),'k-o','LineWidth',2,'MarkerFaceColor','white')
        end
        hold off
        if i == fac*2-1
            xlabel('Time (days)')
        elseif i == 1
            title('Independent Time Weights')
        end
        ylabel(strcat('LV',num2str(count),{' '},' (w_{j',num2str(count),'})'))
        ylim([-1 1])
        yticks([-1 -0.5 0 0.5 1])
        set(gca,'FontSize',fnt_sz_sm,'FontWeight','bold','LineWidth',2)
    end

    count = 0;
    for i = 2:2:fac*2
        count = count+1;
        subplot(fac,2,i)
        hold on
        if ~strcmp(nullPLS,'Yes') && ~strcmp(paired,'None')
            errorbar(time_val(Ytime_choice),Yt_mean(:,count),Yt_std(:,count),'LineWidth',2,'Color',dot_col)
            plot(time_val(Ytime_choice),Yfactors{3}(:,count),'k-o','LineWidth',2,'MarkerFaceColor','white')
            plot(time_val(Ytime_choice),Yt_mean(:,count),strcat(l_col,'-o'),'LineWidth',2,'MarkerFaceColor','white')
        else
            plot(time_val(Ytime_choice),Yfactors{3}(:,count),'k-o','LineWidth',2,'MarkerFaceColor','white')
        end
        hold off
        if i == fac*2
            xlabel('Time (days)')
        elseif i == 2
            title('Dependent Time Weights')
        end

        ylabel(strcat('LV',num2str(count),{' '},' (q_{l',num2str(count),'})'))
        ylim([-1 1])
        yticks([-1 -0.5 0 0.5 1])
        set(gca,'FontSize',fnt_sz_sm,'FontWeight','bold','LineWidth',2)
    end
end

%--------------------------------------------------------------------------
% Scores and variable weights (2x2 subplot)
%--------------------------------------------------------------------------

for k = 1:fac-1
    for i = (k+1):fac
        fig_count = fig_count+1;
        figure(fig_count)
        set(gcf,'Position',[100 100 800 800])
        %-------X scores---------------------------------------------------
        subplot(2,2,1)
        set(gca,'FontSize',fnt_sz_lg,'FontWeight','bold','LineWidth',2)
        hold on
        if ~strcmp(nullPLS,'Yes') && ~strcmp(paired,'None')
            if strcmp(selpar,'Random')
                errorbar(Xsc_mean(:,k),Xsc_mean(:,i),Xsc_std(:,i),Xsc_std(:,i),Xsc_std(:,k),Xsc_std(:,k),'o','Color',dot_col,'LineWidth',lin_wid)
            end
            plot(Xfactors{1}(:,k),Xfactors{1}(:,i),'ko','MarkerSize',5,'MarkerFaceColor','k')
            plot(Xsc_mean(:,k),Xsc_mean(:,i),strcat(l_col,'o'),'LineWidth',2,'MarkerSize',5,'MarkerFaceColor',l_col)
        else
            plot(Xfactors{1}(:,k),Xfactors{1}(:,i),'o','MarkerSize',5,'MarkerFaceColor',dot_col,'MarkerEdgeColor',dot_col)
        end
%         if k == 3 && i == 4
%             xlim([-6 6])
%             ylim([-6 6])
%         end
        if ~all(sign(Xfactors{1}(:,k))>0) || ~all(sign(Xfactors{1}(:,i))>0)
            plot(xlim,[0 0],'k','LineWidth',2)
            plot([0 0],ylim,'k','LineWidth',2)
            plot(xlim,[0 0],'k','LineWidth',2)
        end
        hold off
        if ~strcmp(nullPLS,'Yes') && ~strcmp(paired,'None')
            text(Xfactors{1}(:,k)*(1+xoff),Xfactors{1}(:,i)*(1+yoff),expGroup(groupChoice),'FontSize',fnt_sz_sm,'Color','k')
            text(Xsc_mean(:,k)*(1+xoff),Xsc_mean(:,i)*(1+yoff),expGroup(groupChoice),'FontSize',fnt_sz_sm,'Color',dot_col)
        else
            text(Xfactors{1}(:,k)*(1+xoff),Xfactors{1}(:,i)*(1+yoff),expGroup(groupChoice),'FontSize',fnt_sz_sm,'Color',dot_col)
        end
        xlabel(strcat('LV',num2str(k),{' '},'(t_{i',num2str(k),'})'))
        ylabel(strcat('LV',num2str(i),{' '},'(t_{i',num2str(i),'})'))
%         if k == 1 && i == 2
%             xticks([0 3 6 9 12 15])
%             yticks([0 2 4 6 8])
%         elseif k == 3 && i == 4
%             xticks([-5 0 5 10])
%         end
        box off

        %-------X variable weights-----------------------------------------
        subplot(2,2,2)
        set(gca,'FontSize',fnt_sz_lg,'FontWeight','bold','LineWidth',2)
        hold on
        if ~strcmp(nullPLS,'Yes') && ~strcmp(paired,'None')
            if strcmp(selpar,'Random')
                errorbar(Xv_mean(:,k),Xv_mean(:,i),Xv_std(:,i),Xv_std(:,i),Xv_std(:,k),Xv_std(:,k),'o','Color',dot_col,'LineWidth',lin_wid)
            end
            plot(Xv_mean(:,k),Xv_mean(:,i),strcat(l_col,'o'),'LineWidth',2,'MarkerSize',5,'MarkerFaceColor',dot_col)
        end
        for j = 1:length(varX_choice)
            if strcmp(nullPLS,'Yes')
                if Xfactors{2}(j,k) >= Xv_mean(k)+Xv_std(k) || Xfactors{2}(j,k) <= Xv_mean(k)-Xv_std(k)
                    dcol(j,:) = [0 0 0];
                elseif Xfactors{2}(j,i) >= Xv_mean(i)+Xv_std(i) || Xfactors{2}(j,i) <= Xv_mean(i)-Xv_std(i)
                    dcol(j,:) = [0 0 0];
                else
                    dcol(j,:) = [1 1 1]*gfac;
                end
            else
                if any(strcmp(X_hilite,vars(varX_choice(j))))
                    dcol(j,:) = dot_col;
                else
                    dcol(j,:) = [1 1 1]*gfac;
                end
            end
        end
        for j = 1:length(varX_choice)
            if ~strcmp(nullPLS,'Yes')
                if strcmp(selpar,'Random')
                    plot(Xfactors{2}(j,k),Xfactors{2}(j,i),'o','MarkerSize',5,'MarkerFaceColor','k','MarkerEdgeColor','k')
                else
                    plot(Xfactors{2}(j,k),Xfactors{2}(j,i),'o','MarkerSize',5,'MarkerFaceColor',dcol(j,:),'MarkerEdgeColor',dcol(j,:))
                    if any(strcmp(X_hilite,vars(varX_choice(j))))
                        text(Xfactors{2}(j,k)+xoff,Xfactors{2}(j,i)+yoff,vars(varX_choice(j)),'FontSize',fnt_sz_sm,'Color','k')
                    end
                end
            else
                plot(Xfactors{2}(j,k),Xfactors{2}(j,i),'o','MarkerSize',5,'MarkerFaceColor',dcol(j,:),'MarkerEdgeColor',dcol(j,:))
                if Xfactors{2}(j,k) >= Xv_mean(k)+Xv_std(k) || Xfactors{2}(j,k) <= Xv_mean(k)-Xv_std(k)
                    text(Xfactors{2}(j,k)+xoff,Xfactors{2}(j,i)+yoff,vars(varX_choice(j)),'FontSize',fnt_sz_sm,'Color',dcol(j,:))
                elseif Xfactors{2}(j,i) >= Xv_mean(i)+Xv_std(i) || Xfactors{2}(j,i) <= Xv_mean(i)-Xv_std(i)
                    text(Xfactors{2}(j,k)+xoff,Xfactors{2}(j,i)+yoff,vars(varX_choice(j)),'FontSize',fnt_sz_sm,'Color',dcol(j,:))
                end
            end
        end
%         if k == 1 && i == 2
%             ylim([-0.4 0.6])
%         elseif k == 3 && i == 4
%             ylim([-0.6 0.6])
%         end
        xwin = xlim;
        ywin = ylim;
        if strcmp(nullPLS,'Yes')
            plot([Xv_mean(k) Xv_mean(k)],ylim,'Color',[1 1 1]*gfac,'LineWidth',lin_wid)
            plot(xlim,[Xv_mean(i) Xv_mean(i)],'Color',[1 1 1]*gfac,'LineWidth',lin_wid)
            plot([Xv_mean(k) Xv_mean(k)]-Xv_std(k),ylim,'LineStyle','--','Color',[1 1 1]*gfac,'LineWidth',lin_wid)
            text([Xv_mean(k)-Xv_std(k)],ywin(2),'-\sigma','Color',[1 1 1]*gfac,'FontSize',fnt_sz_lg,'HorizontalAlignment','center','VerticalAlignment','bottom','FontWeight','bold')
            plot(xlim,[Xv_mean(i) Xv_mean(i)]-Xv_std(i),'LineStyle','--','Color',[1 1 1]*gfac,'LineWidth',lin_wid)
            text(xwin(2),[Xv_mean(i)-Xv_std(i)],'-\sigma','Color',[1 1 1]*gfac,'FontSize',fnt_sz_lg,'HorizontalAlignment','left','VerticalAlignment','middle','FontWeight','bold')
            plot([Xv_mean(k) Xv_mean(k)]+Xv_std(k),ylim,'LineStyle','--','Color',[1 1 1]*gfac,'LineWidth',lin_wid)
            text([Xv_mean(k)+Xv_std(k)],ywin(2),'+\sigma','Color',[1 1 1]*gfac,'FontSize',fnt_sz_lg,'HorizontalAlignment','center','VerticalAlignment','bottom','FontWeight','bold')
            plot(xlim,[Xv_mean(i) Xv_mean(i)]+Xv_std(i),'LineStyle','--','Color',[1 1 1]*gfac,'LineWidth',lin_wid)
            text(xwin(2),[Xv_mean(i)+Xv_std(i)],'+\sigma','Color',[1 1 1]*gfac,'FontSize',fnt_sz_lg,'HorizontalAlignment','left','VerticalAlignment','middle','FontWeight','bold')
        end
        xlabel(strcat('LV',num2str(k),{' '},'(w_{k',num2str(k),'})'))
        ylabel(strcat('LV',num2str(i),{' '},'(w_{k',num2str(i),'})'))
        box off
%         if k == 1 && i == 2
%             xticks([-0.5 -0.25 0 0.25 0.5])
%         elseif k == 3 && i == 4
%             xticks([-0.5 -0.25 0 0.25 0.5])
%         end
        hold off
        
        %-------Y scores---------------------------------------------------
        subplot(2,2,3)
        set(gca,'FontSize',fnt_sz_lg,'FontWeight','bold','LineWidth',2)
        hold on
        if ~strcmp(nullPLS,'Yes') && ~strcmp(paired,'None')
            if strcmp(selpar,'Random')
                errorbar(Ysc_mean(:,k),Ysc_mean(:,i),Ysc_std(:,i),Ysc_std(:,i),Ysc_std(:,k),Ysc_std(:,k),'o','Color',dot_col,'LineWidth',lin_wid)
            end
            plot(Yfactors{1}(:,k),Yfactors{1}(:,i),'ko','MarkerSize',5,'MarkerFaceColor','k')
            plot(Ysc_mean(:,k),Ysc_mean(:,i),strcat(l_col,'o'),'LineWidth',2,'MarkerSize',5,'MarkerFaceColor',dot_col)
        else
            plot(Yfactors{1}(:,k),Yfactors{1}(:,i),'o','MarkerSize',5,'MarkerFaceColor',dot_col,'MarkerEdgeColor',dot_col)
        end
%         if k == 3 && i == 4
%             xlim([-6 6])
%             ylim([-6 6])
%         end
        if ~all(sign(Yfactors{1}(:,k))>0) || ~all(sign(Yfactors{1}(:,i))>0)
            plot(xlim,[0 0],'k','LineWidth',2)
            plot([0 0],ylim,'k','LineWidth',2)
        end
        hold off
        if ~strcmp(nullPLS,'Yes') && ~strcmp(paired,'None')
            text(Yfactors{1}(:,k)*(1+xoff),Yfactors{1}(:,i)*(1+yoff),expGroup(groupChoice),'FontSize',fnt_sz_sm,'Color','k')
            text(Ysc_mean(:,k)*(1+xoff),Ysc_mean(:,i)*(1+yoff),expGroup(groupChoice),'FontSize',fnt_sz_sm,'Color',dot_col)
        else
            text(Yfactors{1}(:,k)*(1+xoff),Yfactors{1}(:,i)*(1+yoff),expGroup(groupChoice),'FontSize',fnt_sz_sm,'Color',dot_col)
        end
        xlabel(strcat('LV',num2str(k),{' '},'(u_{i',num2str(k),'})'))
        ylabel(strcat('LV',num2str(i),{' '},'(u_{i',num2str(i),'})'))
%         if k == 1 && i == 2
%             xticks([0 3 6 9 12 15])
%             yticks([0 2 4 6 8])
%         elseif k == 3 && i == 4
%             xticks([-10 -5 0 5 10])
%         end
        box off

        %-------Y variable weights-----------------------------------------
        subplot(2,2,4)
        set(gca,'FontSize',fnt_sz_lg,'FontWeight','bold','LineWidth',2)
        hold on
        if ~strcmp(nullPLS,'Yes') && ~strcmp(paired,'None')
            if strcmp(selpar,'Random')
                errorbar(Yv_mean(:,k),Yv_mean(:,i),Yv_std(:,i),Yv_std(:,i),Yv_std(:,k),Yv_std(:,k),'o','Color',dot_col,'LineWidth',lin_wid)
            end
            plot(Yv_mean(:,k),Yv_mean(:,i),strcat(l_col,'o'),'LineWidth',2,'MarkerSize',5,'MarkerFaceColor',dot_col)
        end
        
        for j = 1:length(varY_choice)
            if strcmp(nullPLS,'Yes')
                if Yfactors{2}(j,k) >= Yv_mean(k)+Yv_std(k) || Yfactors{2}(j,k) <= Yv_mean(k)-Yv_std(k)
                    dcol(j,:) = [0 0 0];
                elseif Yfactors{2}(j,i) >= Yv_mean(i)+Yv_std(i) || Yfactors{2}(j,i) <= Yv_mean(i)-Yv_std(i)
                    dcol(j,:) = [0 0 0];
                else
                    dcol(j,:) = [1 1 1]*gfac;
                end
            else
                if any(strcmp(Y_hilite,vars(varY_choice(j))))
                    dcol(j,:) = dot_col;
                else
                    dcol(j,:) = [1 1 1]*gfac;
                end
            end

        end
        for j = 1:length(varY_choice)
            if ~strcmp(nullPLS,'Yes')
                if strcmp(selpar,'Random')
                    plot(Yfactors{2}(j,k),Yfactors{2}(j,i),'o','MarkerSize',5,'MarkerFaceColor','k','MarkerEdgeColor','k')
                else
                    plot(Yfactors{2}(j,k),Yfactors{2}(j,i),'o','MarkerSize',5,'MarkerFaceColor',dcol(j,:),'MarkerEdgeColor',dcol(j,:))
                    if any(strcmp(Y_hilite,vars(varY_choice(j))))
                        text(Yfactors{2}(j,k)+xoff,Yfactors{2}(j,i)+yoff,vars(varY_choice(j)),'FontSize',fnt_sz_sm,'Color',dcol(j,:))
                    end
                end
            else
                plot(Yfactors{2}(j,k),Yfactors{2}(j,i),'o','MarkerSize',5,'MarkerFaceColor',dcol(j,:),'MarkerEdgeColor',dcol(j,:))
                if ~strcmp(selpar,'Random')
                    if Yfactors{2}(j,k) >= Yv_mean(k)+Yv_std(k) || Yfactors{2}(j,k) <= Yv_mean(k)-Yv_std(k)
                        text(Yfactors{2}(j,k)+xoff,Yfactors{2}(j,i)+yoff,vars(varY_choice(j)),'FontSize',fnt_sz_sm,'Color',dcol(j,:))
                    elseif Yfactors{2}(j,i) >= Yv_mean(i)+Yv_std(i) || Yfactors{2}(j,i) <= Yv_mean(i)-Yv_std(i)
                        text(Yfactors{2}(j,k)+xoff,Yfactors{2}(j,i)+yoff,vars(varY_choice(j)),'FontSize',fnt_sz_sm,'Color',dcol(j,:))
                    end
                end
            end
        end
%         if k == 1 && i == 2
%             ylim([-0.4 0.6])
%         elseif k == 3 && i == 4
%             ylim([-0.6 0.6])
%         end
        xwin = xlim;
        ywin = ylim;
        if strcmp(nullPLS,'Yes')
            plot([Yv_mean(k) Yv_mean(k)],ylim,'Color',[1 1 1]*gfac,'LineWidth',lin_wid)
            plot(xlim,[Yv_mean(i) Yv_mean(i)],'Color',[1 1 1]*gfac,'LineWidth',lin_wid)
            plot([Yv_mean(k) Yv_mean(k)]-Yv_std(k),ylim,'LineStyle','--','Color',[1 1 1]*gfac,'LineWidth',lin_wid)
            text([Yv_mean(k)-Yv_std(k)],ywin(2),'-\sigma','Color',[1 1 1]*gfac,'FontSize',fnt_sz_lg,'HorizontalAlignment','center','VerticalAlignment','bottom','FontWeight','bold')
            plot(xlim,[Yv_mean(i) Yv_mean(i)]-Yv_std(i),'LineStyle','--','Color',[1 1 1]*gfac,'LineWidth',lin_wid)
            text(xwin(2),[Yv_mean(i)-Yv_std(i)],'-\sigma','Color',[1 1 1]*gfac,'FontSize',fnt_sz_lg,'HorizontalAlignment','left','VerticalAlignment','middle','FontWeight','bold')
            plot([Yv_mean(k) Yv_mean(k)]+Yv_std(k),ylim,'LineStyle','--','Color',[1 1 1]*gfac,'LineWidth',lin_wid)
            text([Yv_mean(k)+Yv_std(k)],ywin(2),'+\sigma','Color',[1 1 1]*gfac,'FontSize',fnt_sz_lg,'HorizontalAlignment','center','VerticalAlignment','bottom','FontWeight','bold')
            plot(xlim,[Yv_mean(i) Yv_mean(i)]+Yv_std(i),'LineStyle','--','Color',[1 1 1]*gfac,'LineWidth',lin_wid)
            text(xwin(2),[Yv_mean(i)+Yv_std(i)],'+\sigma','Color',[1 1 1]*gfac,'FontSize',fnt_sz_lg,'HorizontalAlignment','left','VerticalAlignment','middle','FontWeight','bold')
        end
        xlabel(strcat('LV',num2str(k),{' '},'(q_{m',num2str(k),'})'))
        ylabel(strcat('LV',num2str(i),{' '},'(q_{m',num2str(i),'})'))
        box off
%         if k == 1 && i == 2
%             xticks([-0.5 -0.25 0 0.25 0.5])
% %             xticks([-1 -0.5 0 0.5])
%         elseif k == 3 && i == 4
%             xticks([-0.5 -0.25 0 0.25 0.5])
% %             xticks([-0.5 0 0.5 1])
%             yticks([-0.5 -0.25 0 0.25 0.5])
%         end
        hold off
    end
end

%--------------------------------------------------------------------------
% Plot histograms of sampled scores and weights
%--------------------------------------------------------------------------

pl_col = {'r','b','g','k','c','y'};

if ~strcmp(paired,'None')
    %Plot histogram of X condition scores
    fig_count = fig_count + 1;
    figure(fig_count)
    pl_count = 0;
    for f = 1:fac
        for i = 1:length(groupChoice)
            pl_count = pl_count + 1;
            subplot(fac,length(groupChoice),pl_count)
            histogram(shuff_X_scores(:,i,f))
            pd = fitdist(shuff_X_scores(:,i,f),'Normal');
            xrange = get(gca,'xlim');
            normx = xrange(1):0.01:xrange(2);
            normy = normdist(normx,pd.mu,pd.sigma);
            LL = mean(shuff_X_scores(:,i,f)) - std(shuff_X_scores(:,i,f));
            UL = mean(shuff_X_scores(:,i,f)) + std(shuff_X_scores(:,i,f));
            hold on
%             plot(normx,normy,'r','LineWidth',1.5)
            plot([Xfactors{1}(i,f) Xfactors{1}(i,f)],get(gca,'ylim'),'r','LineWidth',2)
            ymax = get(gca,'ylim');
%             area([LL UL],[ymax(2) ymax(2)],'FaceColor','k','FaceAlpha',0.2)
            hold off
            if i == 1
                ylabel({strcat('LV',num2str(f));'Count'})
            end
            
            if f == fac
                xlabel('X score')
            end
            
            if f == 1
                title(expGroup(groupChoice(i)))
            end
            
            set(gca,'FontWeight','bold')
        end
    end
    
    %Plot histogram of Y condition scores - not used in publication
%     fig_count = fig_count + 1;
%     figure(fig_count)
%     pl_count = 0;
%     for f = 1:fac
%         for i = 1:length(groupChoice)
%             pl_count = pl_count + 1;
%             subplot(fac,length(groupChoice),pl_count)
%             histogram(shuff_Y_scores(:,i,f))
%             pd = fitdist(shuff_Y_scores(:,i,f),'Normal');
%             ci = paramci(pd);
%             LL = ci(1,1);
%             UL = ci(2,1);
%             hold on
%             plot([Yfactors{1}(i,f) Yfactors{1}(i,f)],get(gca,'ylim'),'r','LineWidth',2)
% %             plot([LL LL],get(gca,'ylim'),'Color',[0.8 0.8 0.8],'LineWidth',2)
% %             plot([UL UL],get(gca,'ylim'),'Color',[0.8 0.8 0.8],'LineWidth',2)
%             hold off
%             if i == 1
%                 ylabel({strcat('LV',num2str(f));'Count'})
%             end
%             
%             if f == fac
%                 xlabel('Y score')
%             end
%             
%             if f == 1
%                 title(expGroup(groupChoice(i)))
%             end
%             
%             set(gca,'FontWeight','bold')
%         end
%     end
    
    %Plot histogram of X variable weights - not in publication
%     fig_count = fig_count + 1;
%     figure(fig_count)
%     pl_count = 0;
%     for f = 1:fac
%         for j = 1:length(varX_choice)
%             pl_count = pl_count + 1;
%             subplot(fac,length(varX_choice),pl_count)
%             histogram(shuff_Xvar_weights(:,j,f))
%             pd = fitdist(shuff_Xvar_weights(:,j,f),'Normal');
%             ci = paramci(pd);
%             LL = ci(1,1);
%             UL = ci(2,1);
%             hold on
%             plot([Xfactors{2}(j,f) Xfactors{2}(j,f)],get(gca,'ylim'),'r','LineWidth',2)
% %             plot([LL LL],get(gca,'ylim'),'Color',[0.8 0.8 0.8],'LineWidth',2)
% %             plot([UL UL],get(gca,'ylim'),'Color',[0.8 0.8 0.8],'LineWidth',2)
%             hold off
%             if j == 1
%                 ylabel({strcat('LV',num2str(f));'Count'})
%             end
%             
%             if f == fac
%                 xlabel('X var weight')
%             end
%             
%             if f == 1
%                 title(vars(varX_choice(j)))
%             end
%             
%             set(gca,'FontWeight','bold')
%         end
%     end
    
    %Plot histogram of Y variable weights - not in publication
%     fig_count = fig_count + 1;
%     figure(fig_count)
%     pl_count = 0;
%     for f = 1:fac
%         for j = 1:length(varY_choice)
%             pl_count = pl_count + 1;
%             subplot(fac,length(varY_choice),pl_count)
%             histogram(shuff_Yvar_weights(:,j,f))
%             pd = fitdist(shuff_Yvar_weights(:,j,f),'Normal');
%             ci = paramci(pd);
%             LL = ci(1,1);
%             UL = ci(2,1);
%             hold on
%             plot([Yfactors{2}(j,f) Yfactors{2}(j,f)],get(gca,'ylim'),'r','LineWidth',2)
% %             plot([LL LL],get(gca,'ylim'),'Color',[0.8 0.8 0.8],'LineWidth',2)
% %             plot([UL UL],get(gca,'ylim'),'Color',[0.8 0.8 0.8],'LineWidth',2)
%             hold off
%             if j == 1
%                 ylabel({strcat('LV',num2str(f));'Count'})
%             end
%             
%             if f == fac
%                 xlabel('Y var weight')
%             end
%             
%             if f == 1
%                 title(vars(varY_choice(j)))
%             end
%             
%             set(gca,'FontWeight','bold')
%         end
%     end
    
    %Plot histogram of X time weights - not in publication
%     fig_count = fig_count + 1;
%     figure(fig_count)
%     pl_count = 0;
%     for f = 1:fac
%         for k = 1:length(Xtime_choice)
%             pl_count = pl_count + 1;
%             subplot(fac,length(Xtime_choice),pl_count)
%             histogram(shuff_Xtime_weights(:,k,f))
%             pd = fitdist(shuff_Xtime_weights(:,k,f),'Normal');
%             ci = paramci(pd);
%             LL = ci(1,1);
%             UL = ci(2,1);
%             hold on
%             plot([Xfactors{3}(k,f) Xfactors{3}(k,f)],get(gca,'ylim'),'r','LineWidth',2)
% %             plot([LL LL],get(gca,'ylim'),'Color',[0.8 0.8 0.8],'LineWidth',2)
% %             plot([UL UL],get(gca,'ylim'),'Color',[0.8 0.8 0.8],'LineWidth',2)
%             hold off
%             if k == 1
%                 ylabel({strcat('LV',num2str(f));'Count'})
%             end
%             
%             if f == fac
%                 xlabel('X time val')
%             end
%             
%             if f == 1
%                 title(times(Xtime_choice(k)))
%             end
%             
%             set(gca,'FontWeight','bold')
%         end
%     end

    %Plot histogram of Y time weights - not in publication
%     fig_count = fig_count + 1;
%     figure(fig_count)
%     pl_count = 0;
%     for f = 1:fac
%         for k = 1:length(Ytime_choice)
%             pl_count = pl_count + 1;
%             subplot(fac,length(Ytime_choice),pl_count)
%             histogram(shuff_Ytime_weights(:,k,f))
%             pd = fitdist(shuff_Ytime_weights(:,k,f),'Normal');
%             ci = paramci(pd);
%             LL = ci(1,1);
%             UL = ci(2,1);
%             hold on
%             plot([Yfactors{3}(k,f) Yfactors{3}(k,f)],get(gca,'ylim'),'r','LineWidth',2)
% %             plot([LL LL],get(gca,'ylim'),'Color',[0.8 0.8 0.8],'LineWidth',2)
% %             plot([UL UL],get(gca,'ylim'),'Color',[0.8 0.8 0.8],'LineWidth',2)
%             hold off
%             if k == 1
%                 ylabel({strcat('LV',num2str(f));'Count'})
%             end
%             
%             if f == fac
%                 xlabel('Y time val')
%             end
%             
%             if f == 1
%                 title(times(Ytime_choice(k)))
%             end
%             
%             set(gca,'FontWeight','bold')
%         end
%     end
end

%--------------------------------------------------------------------------
%Plot calibrations and predictions vs. measurements for all Y data -
%commented out because these are not included in publication
%--------------------------------------------------------------------------

% for i = 1:numvars
%     fig_count = fig_count+1;
%     figure(fig_count)
%     set(gcf,'Position',[100 100 1250 420])
%     subplot(1,2,1)
%     hold on
%     for j = 1:numsamps
%         for k = 1:numtimepts
%             raw_data(k) = Y(j,i,k);
%             pred_data(k) = calibrated(j,i,k);
%             eval(strcat('h',num2str(j),num2str(k),'=plot(raw_data(k),pred_data(k),''o'',''MarkerEdgeColor'',mrk_col(k,:),''MarkerSize'',7,''LineWidth'',',num2str(lin_wid),');'))
%         end
%     end
%     title(vars(varY_choice(i)))
%     lims = ylim;
%     xlabel('Measured Value')
%     ylabel('Calibrated Value')
%     for j = 1:numsamps
%         eval(strcat('leg_ent(j) = h',num2str(j),'1;'))
%     end
% %     legend(leg_ent,expGroup(groupChoice),'Location','SouthEast')
% %     legend boxoff
%     hold off
%     
% %     clear leg_ent
%     
%     subplot(1,2,2)
%     hold on
%     for j = 1:numsamps
%         for k = 1:numtimepts
%         raw_data(k) = Y(j,i,k);
%         pred_data(k) = xvalypred(j,i,k);
%         eval(strcat('h',num2str(j),num2str(k),'=plot(raw_data(k),pred_data(k),''o'',''MarkerEdgeColor'',mrk_col(k,:),''MarkerSize'',7,''LineWidth'',',num2str(lin_wid),');'))
%         end
%     end
%     title(vars(varY_choice(i)))
%     xlabel('Measured Value')
%     ylabel('Predicted Value')
%     for k = 1:numtimepts
%         eval(strcat('leg_ent(k) = h1',num2str(k),';'))
%     end
% %     legend(leg_ent,times(Ytime_choice),'Location','SouthEast')
% %     legend boxoff
%     ylim(lims)
%     hold off
% end

%--------------------------------------------------------------------------
%Plot calibrations and measurements vs. time for all X data - commented out
%because these are not included in publication
%--------------------------------------------------------------------------

% pan_count = 0;
% raw_data = [];
% pred_data = [];
% fig_count = fig_count+1;
% figure(fig_count)
% set(gcf,'units','normalized','outerposition',[0 0 1 1]);
% ha = tight_subplot(length(groupChoice),numXvars,[0.04 0.02],[0.1 0.05],0.05);
% for j = 1:numsamps
%     for i = 1:numXvars
%         pan_count = pan_count + 1;
%         axes(ha(pan_count))
% %         subplot(length(groupChoice),numvars,pan_count)
%         hold on
%         raw_data(j,:) = squeeze(X(j,i,:));
%         cal_data(j,:) = squeeze(xcalibrated(j,i,:));
% %         for k = 1:length(Xtime_choice)
% %             pred_data(j,k) = sum(Xsc(j,:).*Xwj(i,:).*Xwk(k,:))+X0(j,i,1);
% %         end
% %         pred_data(j,:) = squeeze(xvalypred(j,i,:));
%         plot(time_val(Xtime_choice),raw_data(j,:),'o','MarkerEdgeColor','b','MarkerSize',7,'LineWidth',lin_wid)
%         plot(time_val(Xtime_choice),cal_data(j,:),'g-','LineWidth',lin_wid)
%         hold off
%         set(ha(pan_count),'XTickLabelMode','auto','YTickLabelMode','auto')
%         if j == numsamps
%             xlabel('Time (days)','FontWeight','bold')
%         end
%         
%         if j == 1
%             title(strcat(vars(varX_choice(i)),' (',var_units(varX_choice(i)),')'))
%         end
%         
%         if i == 1
%             ylabel(expGroup(groupChoice(j)),'FontWeight','bold')
%         end
%     end
%     
%     if j == numsamps
%         legend('Measured','Calibrated')
%         legend boxoff
%     end
% end

%--------------------------------------------------------------------------
%Plot predictions and measurements vs. time for all Y data - commented out
%because these are not included in publication
%--------------------------------------------------------------------------

% clear raw_data pred_data cal_data
% pan_count = 0;
% raw_data = [];
% pred_data = [];
% fig_count = fig_count+1;
% figure(fig_count)
% set(gcf,'units','normalized','outerposition',[0 0 1 1]);
% ha = tight_subplot(length(groupChoice),numvars,[0.04 0.02],[0.1 0.05],0.05);
% for j = 1:numsamps
%     for i = 1:numvars
%         pan_count = pan_count + 1;
%         axes(ha(pan_count))
% %         subplot(length(groupChoice),numvars,pan_count)
%         hold on
%         raw_data(j,:) = squeeze(Y(j,i,:));
%         pred_data(j,:) = squeeze(xvalypred(j,i,:));
%         cal_data(j,:) = squeeze(calibrated(j,i,:));
%         plot(time_val(Ytime_choice),raw_data(j,:),'o','MarkerEdgeColor','b','MarkerSize',7,'LineWidth',lin_wid)
%         plot(time_val(Ytime_choice),pred_data(j,:),'r-','LineWidth',lin_wid)
%         plot(time_val(Ytime_choice),cal_data(j,:),'g-','LineWidth',lin_wid)
%         hold off
%         set(ha(pan_count),'XTickLabelMode','auto','YTickLabelMode','auto')
%         if j == numsamps
%             xlabel('Time (days)','FontWeight','bold')
%         end
%         
%         if j == 1
%             title(strcat(vars(varY_choice(i)),' (',var_units(varY_choice(i)),')'))
%         end
%         
%         if i == 1
%             ylabel(expGroup(groupChoice(j)),'FontWeight','bold')
%         end
%     end
%     
%     if j == numsamps
%         legend('Measured','Predicted','Calibrated')
%         legend boxoff
%     end
% end

%--------------------------------------------------------------------------
%Plot predictions (avg+/-std) and measurements vs. time for Y data
%predicted from randomly sampled data - commented out because these are not
%included in publication
%--------------------------------------------------------------------------

% if strcmp(selpar,'Random')
%     clear raw_data pred_data cal_data
%     pan_count = 0;
%     raw_data = [];
%     pred_data = [];
%     fig_count = fig_count+1;
%     figure(fig_count)
%     set(gcf,'units','normalized','outerposition',[0 0 1 1]);
%     ha = tight_subplot(length(groupChoice),numvars,[0.04 0.02],[0.1 0.05],0.05);
%     for j = 1:numsamps
%         for i = 1:numvars
%             pan_count = pan_count + 1;
%             axes(ha(pan_count))
%     %         subplot(length(groupChoice),numvars,pan_count)
%             hold on
%             raw_data(j,:) = squeeze(Y(j,i,:));
%             pred_mean(j,:) = squeeze(xvalypredNs_mean(j,i,:));
%             pred_std(j,:) = squeeze(xvalypredNs_std(j,i,:));
%             predc_mean(j,:) = squeeze(xvalypredCs_mean(j,i,:));
%             predc_std(j,:) = squeeze(xvalypredCs_std(j,i,:));
% %             errorbar(time_val(Ytime_choice),pred_mean(j,:),pred_std(j,:),'r-','LineWidth',lin_wid)
%             errorbar(time_val(Ytime_choice),predc_mean(j,:),predc_std(j,:),'b-','LineWidth',lin_wid)
%             plot(time_val(Ytime_choice),raw_data(j,:),'o','MarkerEdgeColor','k','MarkerSize',7,'LineWidth',lin_wid)
%             hold off
%             set(ha(pan_count),'XTickLabelMode','auto','YTickLabelMode','auto')
%             if j == numsamps
%                 xlabel('Time (days)','FontWeight','bold')
%             end
% 
%             if j == 1
%                 title(strcat(vars(varY_choice(i)),' (',var_units(varY_choice(i)),')'))
%             end
% 
%             if i == 1
%                 ylabel(expGroup(groupChoice(j)),'FontWeight','bold')
%             end
%         end
% 
%         if j == numsamps
%             legend('Measured','Corrected')
%             legend boxoff
%         end
%     end
% end

%--------------------------------------------------------------------------
% Breakout plots of sampled predictions - commented out because these are
% not included in publication
%--------------------------------------------------------------------------

% if strcmp(selpar,'Random')
%     for i = 1:numvars
%         pan_count = 0;
%         fig_count = fig_count+1;
%         figure(fig_count)
%         set(gcf,'Position',[100 100 800 800])
%         hold on
%         for j = 1:numsamps
%             for k = 1:numtimepts
%                 raw_data(j,k) = Y(j,i,k);
%                 pred_mean(j,k) = xvalypredNs_mean(j,i,k);
%                 pred_std(j,k) = xvalypredNs_std(j,i,k);
%                 predc_mean(j,k) = xvalypredCs_mean(j,i,k);
%                 predc_std(j,k) = xvalypredCs_std(j,i,k);
%             end
%         end
%         
%         for j = 1:numsamps
%             pan_count = pan_count+1;
%             subplot(numsamps,2,pan_count)
%             hold on
%             errorbar(time_val(Ytime_choice),pred_mean(j,:),pred_std(j,:),'r-','LineWidth',lin_wid)
%             plot(time_val(Ytime_choice),raw_data(j,:),'o','MarkerEdgeColor','k','MarkerSize',7,'LineWidth',lin_wid)
%             hold off
%             if pan_count == 1
%                 title(strcat(vars(varY_choice(i)),' (',var_units(varY_choice(i)),')'))
%             end
%             ylabel(expGroup(groupChoice(j)),'FontWeight','bold')
%             if j == numsamps
%                 xlabel('Time (days)')
%             end
%             set(gca,'FontSize',fnt_sz_sm,'FontWeight','bold','LineWidth',2)
%             if pan_count == numsamps*2-1
%                 legend('Predicted','Measured')
%                 legend boxoff
%             end
%             
%             pan_count = pan_count+1;
%             subplot(numsamps,2,pan_count)
%             hold on
%             errorbar(time_val(Ytime_choice),predc_mean(j,:),predc_std(j,:),'b-','LineWidth',lin_wid)
%             plot(time_val(Ytime_choice),raw_data(j,:),'o','MarkerEdgeColor','k','MarkerSize',7,'LineWidth',lin_wid)
%             hold off
%             if pan_count == 2
%                 title(strcat(vars(varY_choice(i)),' (',var_units(varY_choice(i)),')'))
%             end
%             if j == numsamps
%                 xlabel('Time (days)')
%             end
%             set(gca,'FontSize',fnt_sz_sm,'FontWeight','bold','LineWidth',2)
%             if pan_count == numsamps*2
%                 legend('Corrected','Measured')
%                 legend boxoff
%             end
%             
%             ywin = ylim;
%             subplot(numsamps,2,pan_count-1)
%             ylim(ywin);
%         end
%     end
% end
```
