## Supplementary File S1 for "Robust latent-variable interpretation of *in vivo* regression models by nested resampling": runMRBpls_clean.m

```
%Generate PLSR model of data from Bersi et al., 2017
%Alex Caulk
%Yale University
%February 2018

clear
close all
clc

%--------------------------------------------------------------------------
%Data ordered in matrix as MRBa2(i,j,k). Note that data is permuted later
%in code to make variables mode 3 (index k) and time mode 2 (index j).
%
%Rows (i) 1-4: ATA, DTA, SAA, IAA
%
%Columns (j) variables
var_idx{1} = 'properties.loaded.run1(1,1)'; %1: Systolic BP
var_idx{2} = 'properties.unloaded(1,2)'; %2: OD
var_idx{3} = 'properties.unloaded(1,3)'; %3: H
var_idx{4} = 'properties.unloaded(1,1)'; %4: L
var_idx{5} = 'properties.loaded.run1(1,2)'; %5: od systolic
var_idx{7} = 'properties.loaded.run1(1,3)'; %7: h systolic
var_idx{6} = '(var_idx{5} - 2*var_idx{7})/2'; %6: ir systolic
var_idx{8} = 'properties.loaded.run1(1,4)'; %8: estimated in vivo axial stretch
var_idx{9} = 'properties.loaded.run1(1,8)'; %9: systolic circ stress
var_idx{10} = 'properties.loaded.run1(1,7)'; %10: systolic ax stress
var_idx{11} = 'stiffness_SoL.run1(1,1)'; %11: systolic circ stiff
var_idx{12} = 'stiffness_SoL.run1(1,2)'; %12: systolic ax stiff
var_idx{13} = 'var_idx{11}*var_idx{7}'; %13: systolic struct stiff
var_idx{14} = 'properties.loaded.run1(1,6)'; %14: systolic stored energy
var_idx{15} = ''; %15: distensibility
var_idx{16} = 'histo.wall.med.perc'; %16: medial cross-sectional area
var_idx{17} = 'histo.wall.adv.perc'; %17: adventitial cross-sectional area
var_idx{18} = 'histo.eln.med.AF'; %18: medial elastin area fraction
var_idx{19} = 'histo.smc.med.AF'; %19: medial SMC area fraction
var_idx{20} = 'histo.col.med.AF'; %20: medial collagen area fraction
var_idx{21} = 'histo.gag.med.AF'; %21: medial GAG area fraction
var_idx{22} = 'histo.eln.adv.AF'; %22: adventitial elastin area fraction
var_idx{23} = 'histo.col.adv.AF'; %23: adventitial collagen area fraction
var_idx{24} = 'immuno.CD3.med.dens'; %24: medial CD3 cell density
var_idx{25} = 'immuno.CD45.med.dens'; %25: medial CD45 cell density
var_idx{26} = 'immuno.CD68.med.dens'; %26: medial CD68 cell density
var_idx{27} = 'immuno.CD3.adv.dens'; %27: adventitial CD3 cell density
var_idx{28} = 'immuno.CD45.adv.dens'; %28: adventitial CD45 cell density
var_idx{29} = 'immuno.CD68.adv.dens'; %29: adventitial CD68 cell density
var_idx{30} = 'immuno.MMP2.med.AF'; %30: medial MMP2 area fraction
var_idx{31} = 'immuno.MMP12.med.AF'; %31: medial MMP12 area fraction
var_idx{32} = 'immuno.MMP13.med.AF'; %32: medial MMP13 area fraction
var_idx{33} = 'immuno.MMP2.adv.AF'; %33: adventitial MMP2 area fraction
var_idx{34} = 'immuno.MMP12.adv.AF'; %34: adventitial MMP12 area fraction
var_idx{35} = 'immuno.MMP13.adv.AF'; %35: adventitial MMP13 area fraction
%
%Columns (k) 1-6 time points: 0,4,7,14,21,28 days AngII infusion
%--------------------------------------------------------------------------

path = cd;
%Check Mac vs. PC for file loading and directory switching
if contains(path,'\')
    OS = '\';
else
    OS = '/';
end

%--------------------------------------------------------------------------
%Load and label all data
%--------------------------------------------------------------------------

load('MRB_AngII.mat')
MRBa2 = [MRBa2 dMRBa2]; %Includes raw data and differential data (differential data not used in publication)

expGroup = {'ATA','DTA','SAA','IAA'}; %Mode 1 names
times = {'0d','4d','7d','14d','21d','28d'}; %Mode 2 names
time_val = [0 4 7 14 21 28];
vars = {'BP','OD','H','L','od','ir','h','lziv','tqq','tzz','Cqqqq','Czzzz',...
        'Cth','W','Dist','CSAm','CSAa','elnm','SMCm','colm','GAGm',...
        'elna','cola','CD3m','CD45m','CD68m','CD3a','CD45a','CD68a',...
        'MMP2m','MMP12m','MMP13m','MMP2a','MMP12a','MMP13a'}; %Mode 3 names
dvars = {'dBP','dOD','dH','dL','dod','dir','dh','dlziv','dtqq','dtzz','dCqqqq','dCzzzz',...
        'dCth','dW','dDist','dCSAm','dCSAa','deAFm','dSMCAFm','dcAFm','dGAGAFm',...
        'deAFa','dcAFa','dCD3m','dCD45m','dCD68m','dCD3a','dCD45a','dCD68a',...
        'dMMP2m','dMMP12m','dMMP13m','dMMP2a','dMMP12a','dMMP13a'};
vars = [vars dvars];
var_units = {'mmHg','\mum','\mum','mm','\mum','\mum','\mum','-','kPa','kPa','MPa','MPa',...
             'N/m','kPa','MPa^{-1}','mm^2','mm^2','%','%','%','%',...
             '%','%','#/mm^2','#/mm^2','#/mm^2','#/mm^2','#/mm^2','#/mm^2',...
             '%','%','%','%','%','%'}; %Mode 3 units
var_units = [var_units var_units];

%--------------------------------------------------------------------------
%Select mode of evaluation (e.g., randomization, bootstrapping,
%jackknifing) and select data to include in model. Note that default
%settings correlate to those used in publication.
%--------------------------------------------------------------------------

nullPLS = questdlg('Calculate null-PLS model?','Null...','Yes','No','No');
if strcmp(nullPLS,'No')
    paired = questdlg('Paired or unpaired sampling?','Paired...','Paired','Unpaired','None','Unpaired');
    if strcmp(paired,'Paired')
        selpar = questdlg('Selected or random sampling?','Selected...','Selected','Random','Random');
        jk = questdlg('Sampling Method?','Sampling...','Jackknife','Bootstrap','Bootstrap');
        if strcmp(selpar,'Random')
            numsims = inputdlg('Enter number of simulations for sampling','Simulation number',[1 35],{'500'});
            numsims = str2num(numsims{1});
        else
            numsims = 1;
        end
                    
        %Paired filenames for paired analysis
        biosamp(1,:,1) = {'ATA_C31-2_day0','ATA_C62-1_day0'};
        biosamp(1,:,2) = {'ATA_C63-1_day4','ATA_C64-4_day4'};
        biosamp(1,:,3) = {'ATA_C79-2_day14','ATA_C94-2_day14'};
        biosamp(1,:,4) = {'ATA_C57-3_day28','ATA_C71-5_day28'};

        biosamp(2,:,1) = {'DTA_C39-1_day0','DTA_C43-2_day0'};
        biosamp(2,:,2) = {'DTA_C58-1_day4','DTA_C58-2_day4'};
        biosamp(2,:,3) = {'DTA_C70-3_day14','DTA_C79-1_day14'};
        biosamp(2,:,4) = {'DTA_C37-4_day28','DTA_C81-3_day28'};

        biosamp(3,:,1) = {'SAA_C31-1_day0','SAA_C62-1_day0'};
        biosamp(3,:,2) = {'SAA_C58-2_day4','SAA_C71-1_day4'};
        biosamp(3,:,3) = {'SAA_C5-1_day14','SAA_C21-3_day14'};
        biosamp(3,:,4) = {'SAA_C15-1_day28','SAA_C57-1_day28'};

        biosamp(4,:,1) = {'IAA_C24-5_day0','IAA_C31-2_day0'};
        biosamp(4,:,2) = {'IAA_C58-2_day4','IAA_C63-1_day4'};
        biosamp(4,:,3) = {'IAA_C70-3_day14','IAA_C79-2_day14'};
        biosamp(4,:,4) = {'IAA_C37-3_day28','IAA_C81-2_day28'};

        mechsamp(1,:,1) = {'020914-ATA2','042714-ATA'};
        mechsamp(1,:,2) = {'111914-ATA1','112014-ATA1'};
        mechsamp(1,:,3) = {NaN,NaN};
        mechsamp(1,:,4) = {'042015-ATA1','120615-ATA2'};
        mechsamp(1,:,5) = {NaN,NaN};
        mechsamp(1,:,6) = {NaN,'021015-ATA'}; %ind 1 not paired

        mechsamp(2,:,1) = {'022214-PTA','040314-PTA'};
        mechsamp(2,:,2) = {'050114-PTA','050114-PTA2'};
        mechsamp(2,:,3) = {NaN,NaN};
        mechsamp(2,:,4) = {'012615-PTA','042015-PTA2'};
        mechsamp(2,:,5) = {NaN,NaN};
        mechsamp(2,:,6) = {'032214-PTA','051215-PTA'};

        mechsamp(3,:,1) = {'020914-SAA1','042714-SAA'};
        mechsamp(3,:,2) = {'050214-SAA',NaN}; %ind 2 not paired
        mechsamp(3,:,3) = {NaN,NaN};
        mechsamp(3,:,4) = {'061013-SAA','092513-SAA'};
        mechsamp(3,:,5) = {NaN,NaN};
        mechsamp(3,:,6) = {NaN,'053014-SAA1'};

        mechsamp(4,:,1) = {'011214-IAA','020814-IAA2'};
        mechsamp(4,:,2) = {'050114-IAA2','111914-IAA1'};
        mechsamp(4,:,3) = {NaN,NaN};
        mechsamp(4,:,4) = {'012615-IAA','042015-IAA1'};
        mechsamp(4,:,5) = {NaN,NaN};
        mechsamp(4,:,6) = {'032214-IAA','051115-IAA1'};

        bio_samp_size = 2*ones(4,4);
        mech_samp_size = 2*ones(4,6);
    elseif strcmp(paired,'Unpaired')
        numsims = inputdlg('Enter number of simulations for sampling','Simulation number',[1 35],{'500'});
        numsims = str2num(numsims{1});
        selpar = 'Random';
        jk = questdlg('Sampling Method?','Sampling...','Jackknife','Bootstrap','Bootstrap');
        %All filenames for unpaired random sampling
        biosamp(1,:,1) = {'ATA_C31-2_day0','ATA_C62-1_day0'};
        biosamp(1,:,2) = {'ATA_C63-1_day4','ATA_C64-4_day4'};
        biosamp(1,:,3) = {'ATA_C79-2_day14','ATA_C94-2_day14'};
        biosamp(1,:,4) = {'ATA_C57-3_day28','ATA_C71-5_day28'};

        biosamp(2,:,1) = {'DTA_C39-1_day0','DTA_C43-2_day0'};
        biosamp(2,:,2) = {'DTA_C58-1_day4','DTA_C58-2_day4'};
        biosamp(2,:,3) = {'DTA_C70-3_day14','DTA_C79-1_day14'};
        biosamp(2,:,4) = {'DTA_C37-4_day28','DTA_C81-3_day28'};

        biosamp(3,:,1) = {'SAA_C31-1_day0','SAA_C62-1_day0'};
        biosamp(3,:,2) = {'SAA_C58-2_day4','SAA_C71-1_day4'};
        biosamp(3,:,3) = {'SAA_C5-1_day14','SAA_C21-3_day14'};
        biosamp(3,:,4) = {'SAA_C15-1_day28','SAA_C57-1_day28'};

        biosamp(4,:,1) = {'IAA_C24-5_day0','IAA_C31-2_day0'};
        biosamp(4,:,2) = {'IAA_C58-2_day4','IAA_C63-1_day4'};
        biosamp(4,:,3) = {'IAA_C70-3_day14','IAA_C79-2_day14'};
        biosamp(4,:,4) = {'IAA_C37-3_day28','IAA_C81-2_day28'};

        mechsamp(1,:,1) = {'020914-ATA2','040314-ATA','042714-ATA','111414-ATA','120614-ATA2',NaN,NaN};
        mechsamp(1,:,2) = {'111914-ATA1','112014-ATA1','112414-ATA','121615-ATA1','122214-ATA',NaN,NaN};
        mechsamp(1,:,3) = {'030714-ATA','041415-ATA1','050314-ATA','122115-ATA1','122115-ATA2',NaN,NaN};
        mechsamp(1,:,4) = {'012615-ATA','042015-ATA1','050913-ATA','061313-ATA','120615-ATA1','120615-ATA2',NaN};
        mechsamp(1,:,5) = {'031514-ATA','042915-ATA2','051914-ATA2','051914-ATA3','121115-ATA1','121215-ATA1',NaN};
        mechsamp(1,:,6) = {'021015-ATA','032314-ATA','042015-ATA2','051115-ATA2','052713-ATA','053014-ATA1','122115-ATA3'};

        mechsamp(2,:,1) = {'011114-PTA','011314-PTA','020914-PTA1','022214-PTA','040314-PTA',NaN,NaN};
        mechsamp(2,:,2) = {'011614-PTA','041414-PTA','041514-PTA','050114-PTA','050114-PTA2','111914-PTA1','112014-PTA'};
        mechsamp(2,:,3) = {'011814-PTA','011814-PTA2','022714-PTA','122115-PTA1','122115-PTA2',NaN,NaN};
        mechsamp(2,:,4) = {'012615-PTA','030714-PTA2','042015-PTA1','042015-PTA2','120615-PTA',NaN,NaN};
        mechsamp(2,:,5) = {'042915-PTA1','042915-PTA2','051115-PTA1','121115-PTA1','121215-PTA1',NaN,NaN};
        mechsamp(2,:,6) = {'031414-PTA','032214-PTA','020815-PTA','051215-PTA','122115-PTA3',NaN,NaN};

        mechsamp(3,:,1) = {'011314-SAA','020914-SAA1','020914-SAA2','022214-SAA','042714-SAA','111414-SAA',NaN};
        mechsamp(3,:,2) = {'041514-SAA','041514-SAA2','050114-SAA','050214-SAA','111914-SAA1','121615-SAA1',NaN};
        mechsamp(3,:,3) = {'011814-SAA','022714-SAA','050314-SAA','122215-SAA1',NaN,NaN,NaN};
        mechsamp(3,:,4) = {'061013-SAA','092513-SAA','103013-SAA','120615-SAA',NaN,NaN,NaN};
        mechsamp(3,:,5) = {'031514-SAA','051914-SAA1','042915-SAA1','121315-SAA1',NaN,NaN,NaN};
        mechsamp(3,:,6) = {'062413-SAA','053014-SAA1','051115-SAA1','122215-SAA2',NaN,NaN,NaN};

        mechsamp(4,:,1) = {'011214-IAA','011314-IAA','020814-IAA1','020814-IAA2','022314-IAA',NaN,NaN};
        mechsamp(4,:,2) = {'011514-IAA','041414-IAA','041514-IAA','050114-IAA','050114-IAA2','111914-IAA1','111914-IAA2'};
        mechsamp(4,:,3) = {'022614-IAA','030614-IAA','122115-IAA1','122115-IAA2','122115-IAA3',NaN,NaN};
        mechsamp(4,:,4) = {'012615-IAA','042015-IAA1','042015-IAA2','120615-IAA1','120615-IAA2',NaN,NaN};
        mechsamp(4,:,5) = {'031414-IAA','042915-IAA1','042915-IAA2','121115-IAA1','121215-IAA1',NaN,NaN};
        mechsamp(4,:,6) = {'032214-IAA','032214-IAA2','020815-IAA','051115-IAA1','122215-IAA1',NaN,NaN};

        bio_samp_size = 2*ones(4,4);
        mech_samp_size = [5 5 5 6 6 7;
                          5 7 5 5 5 5;
                          6 6 4 4 4 4;
                          5 7 5 5 5 5];
    else
        selpar = '';
        numsims = 1;
        jk = '';
    end
else
    paired = '';
    selpar = '';
    jk = '';
    numsims = inputdlg('Enter number of simulations for null model','Simulation number',[1 35],{'500'});
    numsims = str2num(numsims{1});
end

%Parameters definining normalization (i.e., mean centering and variance
%scaling). 1 = True, 0 = False.
meanctr = [0 0 0]; %no centering (all normalized in preprocessing to time 0)
scalemechdata = [0 0 1]; %mode 3 scaling

%Select groups, variables, and times to include in the data sets. If "def"
%equals 'Yes', then manual selection can be skipped, but default settings
%need to be predetermined in "if" statement.
def = questdlg('Choose default data arrangement settings?','Data arrangement...','Yes','No','Yes');

if strcmp(def,'Yes')
    groupChoice = 1:length(expGroup); %Group indices to include
    varX_choice = [18:21,23:35]; %X variable indices to include
    varY_choice = [2:3,5:12,14:15]; %Y variable indices to include
    Xtime_choice = [1 2 4 6]; %X time indices to include
    if strcmp(nullPLS,'Yes')
        Ytime_choice = 1:6; %Y time indices to include
    else
        if ~strcmp(paired,'Paired')
            Ytime_choice = 1:6; %unpaired random sampling or mean
        else
            Ytime_choice = [1 2 4 6]; %paired random sampling
        end
    end
else
    %--------------------------------------------------------------------------
    %Choose vessel regions to evaluate to set i indices
    %--------------------------------------------------------------------------

    groupChoice = listdlg('PromptString','Select Vessels to Include in Evaluation','SelectionMode','multiple','ListString',expGroup);

    %--------------------------------------------------------------------------
    %Choose data for X matrix to set j indices of X
    %--------------------------------------------------------------------------

    varX_choice = listdlg('PromptString','Select Variables for X Matrix','SelectionMode','multiple','ListString',vars);

    %--------------------------------------------------------------------------
    %Choose data for Y matrix to set j indices of Y
    %--------------------------------------------------------------------------

    vars_leftover = vars;
    ids = 1:length(vars);
    vars_leftover(varX_choice) = []; %Remove choices for X to avoid repeat data in X and Y matrices
    ids(varX_choice) = [];
    varY_choice = listdlg('PromptString','Select Variables for Y Matrix','SelectionMode','multiple','ListString',vars_leftover);
    varY_choice = ids(varY_choice);

    %--------------------------------------------------------------------------
    %Choose X and Y time points for evaluation to set k indices
    %--------------------------------------------------------------------------

    Xtime_choice = listdlg('PromptString','Select Times to Evaluate in X Matrix','SelectionMode','multiple','ListString',times);
    sametime = questdlg('Same X and Y time points?','Time points...','Yes','No','Yes');
    if strcmp(sametime,'Yes')
        Ytime_choice = Xtime_choice;
    else
        Ytime_choice = listdlg('PromptString','Select Times to Evaluate in Y Matrix','SelectionMode','multiple','ListString',times);
    end
end

%--------------------------------------------------------------------------
% Preprocess selected data
%
% Note: The order of modes 2 and 3 matters for normalization! If
% variance scaling modes 2 and 3, the code will first normalize all data
% within mode 2 (e.g., make variance = 1 for all conditions and time points
% within a single variable) and will then normalize all data within mode 3.
% Therefore, some data gets variance scaled twice, and it matters which
% happens first. The data and code are set up (and named) for variables in
% mode 2 and time points in mode 3. If different order is desired, one can
% permute the matrices before and after normalization without propagating
% changes across all scripts.
%--------------------------------------------------------------------------

X = MRBa2(groupChoice,varX_choice,Xtime_choice);
Y = MRBa2(groupChoice,varY_choice,Ytime_choice);

X0 = MRBa2(groupChoice,varX_choice,1);
Y0 = MRBa2(groupChoice,varY_choice,1);

cd(strcat(path,OS,'nway320 exchange'))

%If some preprocessing other than mean centering or variance scaling is
%necessary, execute it here.
for i = 1:length(groupChoice)
    for j = 1:size(X,2)
        Xpre(i,j,:) = X(i,j,:)-X0(i,j,1);
    end
    for j = 1:size(Y,2)
        Ypre(i,j,:) = (Y(i,j,:)-Y0(i,j,1));
    end
end
    
%Permute modes 2 and 3 and normalize X and Y data according to parameters 
%set for centering and scaling
[X_norm,mX,sX]=nprocess(permute(Xpre,[1 3 2]), meanctr, scalemechdata);
[Y_norm,mY,sY]=nprocess(permute(Ypre,[1 3 2]), meanctr, scalemechdata);

%Return data to original configuration to fit coding setup
X_norm = permute(X_norm,[1 3 2]);
Y_norm = permute(Y_norm,[1 3 2]);
mX_temp = mX(2); sX_temp = sX(2); mY_temp = mY(2); sY_temp = sY(2);
mX(2) = mX(3); mX(3) = mX_temp;
sX(2) = sX(3); sX(3) = sX_temp;
mY(2) = mY(3); mY(3) = mY_temp;
sY(2) = sY(3); sY(3) = sY_temp;
cd(path)

%--------------------------------------------------------------------------
%Run PLS regression on preprocessed data
%--------------------------------------------------------------------------

cd(strcat(path,OS,'nway320 exchange'))

numsamps=length(groupChoice);
numtimepts=length(Ytime_choice);
numvars=length(varY_choice);
numXvars=length(varX_choice);

%Run initial PLSR model that identifies latent variable with min RMSE
[Results_Temp] = ncrossreg2(X_norm,Y_norm,10,any(meanctr==1));
RMSE_diff = Results_Temp.RMSEP(2:10) - Results_Temp.RMSEP(1:9);
fac = find(RMSE_diff>=0,1,'first');
if isempty(fac)
    fac = 10; 
end
numcomp = fac;

[Xfactors,Yfactors,Core,B,ypred,ssx,ssy] = npls(X_norm,Y_norm,fac,1); %Genereate PLS model with correct number LVs
[ycalpred,T,ssxP,Xres] = npred(X_norm,fac,Xfactors,Yfactors,Core,B,0); %Predict Y data from X data and PLS model
if ndims(X_norm) > 2
    ycalpred = reshape(ycalpred,numsamps,numvars,numtimepts);  %model calibration performance
end
calibrated = nprocess(ycalpred,meanctr,[0 1 0], ...
    mY,sY,-1); %"Un-center" and "un-scale" calibrated data
[Results, Model] = ncrossreg(X_norm,Y_norm,fac,any(meanctr==1));
xvalypred_pre = shiftdim(Results.ypred(fac,:,:,:),1);
xvalypred = nprocess(shiftdim(Results.ypred(fac,:,:,:),1), ...
    meanctr,[0 1 0],mY,sY,-1);  %model cross-validated predictions
Xsc = Xfactors{1}; %Mode 1 X scores
Xwj = Xfactors{2}; %Mode 2 X weights
Xwk = Xfactors{3}; %Mode 3 X weights
for j = 1:numsamps
    for i = 1:numXvars
        for k = 1:length(Xtime_choice)
            xcal(j,i,k) = sum(Xsc(j,:).*Xwj(i,:).*Xwk(k,:)); %Model X values
        end
    end
end
xcalibrated = nprocess(xcal,meanctr,[0 1 0], ...
    mX,sX,-1);

%If any preprocessing other than mean centering and variance scaling was
%performed, it needs to be accounted for here (e.g., if you subtracted time
%0 values, they need to be added back here).
for i = 1:length(groupChoice)
    for j = 1:size(Y,2)
        xvalypred(i,j,:) = xvalypred(i,j,:)+Y0(i,j,1);
        calibrated(i,j,:) = calibrated(i,j,:)+Y0(i,j,1);
    end
    
    for j = 1:size(X,2)
        xcalibrated(i,j,:) = xcalibrated(i,j,:)+X0(i,j,1);
    end
end

%Calculate correlation coefficients for raw (i.e., measured) and normalized
%data and cross-validated predictions. Note: data can be perfectly 
%correlated (R = 1) and still have poor RMSE, so this is not the only 
%metric for performance!
all_pred_data = reshape(xvalypred,1,prod(size(xvalypred)));
all_pred_pre_data = reshape(xvalypred_pre,1,prod(size(xvalypred_pre)));
all_cal_data = reshape(calibrated,1,prod(size(calibrated)));
all_Y_data = reshape(Y,1,prod(size(Y)));
all_Y_pre_data = reshape(Y_norm,1,prod(size(Y_norm)));

%Linear regression for predicted data
R = corr(all_pred_data',all_Y_data','Type','Pearson');
rho = corr(all_pred_data',all_Y_data','Type','Spearman');

Rpre = corr(all_pred_pre_data',all_Y_pre_data','Type','Pearson');
rhopre = corr(all_pred_pre_data',all_Y_pre_data','Type','Spearman');

cd(path)

%--------------------------------------------------------------------------
%Calculate a null Tri-PLS model
%
%The null-PLS model utilizes randomization of data within modes 2 and 3 for
%a given group in mode 1 in order to determine statistical significance of
%parameters.
%--------------------------------------------------------------------------

if strcmp(nullPLS,'Yes')
    for idx=1:numsims
        %shuffling is only done across signals and time points
        for i = 1:length(groupChoice)
            X_norm_shuff(i,:,:) = shufflematrix(reshape(X_norm(i,:,:),size(X_norm,2),size(X_norm,3)));
            Y_norm_shuff(i,:,:) = shufflematrix(reshape(Y_norm(i,:,:),size(Y_norm,2),size(Y_norm,3)));
        end

        cd(strcat(path,OS,'nway320 exchange'))

        %Normalize data with permuted modes 2 and 3
        [Xnull,mX,sX]=nprocess(permute(X_norm_shuff,[1 3 2]), meanctr, scalemechdata);   %mode 1 centering and mode 2+3 scaling
        [Ynull,mY,sY]=nprocess(permute(Y_norm_shuff,[1 3 2]), meanctr, scalemechdata); %mode 1 centering and mode 2+3 scaling
        
        %Return data to original configuration to fit coding setup
        Xnull = permute(Xnull,[1 3 2]);
        Ynull = permute(Ynull,[1 3 2]);
        mX_temp = mX(2); sX_temp = sX(2); mY_temp = mY(2); sY_temp = sY(2);
        mX(2) = mX(3); mX(3) = mX_temp;
        sX(2) = sX(3); sX(3) = sX_temp;
        mY(2) = mY(3); mY(3) = mY_temp;
        sY(2) = sY(3); sY(3) = sY_temp;

        %Generate model
        [XNfactors,YNfactors,Core,Bnull,ypred,ssxN,ssyN] = npls(Xnull,Ynull,numcomp,NaN);

        cd(path)

        %Collect scores and weights into individual matrices
        shuff_X_scores(idx,:,:) = XNfactors{1};
        shuff_Y_scores(idx,:,:) = YNfactors{1};
        shuff_Xvar_weights(idx,:,:)=XNfactors{2};
        shuff_Xtime_weights(idx,:,:)=XNfactors{3};
        shuff_Yvar_weights(idx,:,:)=YNfactors{2};
        shuff_Ytime_weights(idx,:,:)=YNfactors{3};

        %Check sign conventions between the mean model and the current
        %iteration of the null-PLS model. If signs of groups are opposite,
        %tsign = -1, else tsign = 1;
        for f = 1:fac
            for i = 1:length(groupChoice)
                if Xfactors{1}(i,f)*XNfactors{1}(i,f) < 0
                    tsign(idx,i,f) = -1;
                else
                    tsign(idx,i,f) = 1;
                end
            end

            %If number of flipped signs in a latent variable is greater 
            %than half of the total groups, then assign that latent
            %variable to have signs switched. This ensures uniformity of
            %results when comparing to the mean model.
            if length(find(tsign(idx,:,f)==-1)) > floor(length(groupChoice)/2)
                sign_all(f) = -1;
            else
                sign_all(f) = 1;
            end
        end
        
        %Switch signs and variable weights of latent variables according to
        % "sign_all". Note that two sets of parameters must have signs
        % flipped (in this case mode 1 and mode 2) in order for the model
        % to remain  mathematically identical.
        for f = 1:fac
            shuff_X_scores(idx,:,f) = sign_all(f)*shuff_X_scores(idx,:,f);
            shuff_Y_scores(idx,:,f) = sign_all(f)*shuff_Y_scores(idx,:,f);
            shuff_Xvar_weights(idx,:,f) = sign_all(f)*shuff_Xvar_weights(idx,:,f);
            shuff_Yvar_weights(idx,:,f) = sign_all(f)*shuff_Yvar_weights(idx,:,f);
        end
    end
end

%--------------------------------------------------------------------------
%Randomly sample individual data sets to generate average model by
%jackknifing or bootstrapping (only if "nullPLS" was not selected).
%--------------------------------------------------------------------------

if ~strcmp(nullPLS,'Yes') && ~strcmp(paired,'None')
    
    Xs0 = zeros(length(groupChoice),length(varX_choice),length(Xtime_choice),numsims);
    Ys0 = zeros(length(groupChoice),length(varY_choice),length(Ytime_choice),numsims);
    
    Xs = zeros(length(groupChoice),length(varX_choice),length(Xtime_choice),numsims);
    Ys = zeros(length(groupChoice),length(varY_choice),length(Ytime_choice),numsims);
    
    Xs_norm = zeros(length(groupChoice),length(varX_choice),length(Xtime_choice),numsims);
    Ys_norm = zeros(length(groupChoice),length(varY_choice),length(Ytime_choice),numsims);
    
    bmat = zeros(length(groupChoice),length(Xtime_choice));
    mmat = zeros(length(groupChoice),length(Ytime_choice));

    if strcmp(paired,'Paired') && strcmp(selpar,'Selected')
        bmat = [2 2 2 2;
                2 2 2 2;
                1 1 1 2;
                2 2 1 2]; %paired selected sampling
    end

    %Randomly select independent data and assemble array for model generation
    for ns = 1:numsims
        for i = 1:length(groupChoice)
            for k = 1:length(Xtime_choice)
                if strcmp(paired,'Paired') && strcmp(selpar,'Random')
                    bmat(i,k) = randi(2); %paired random sampling
                    if all(isnan(mechsamp{i,bmat(i,k),Ytime_choice(k)})==1)
                        if bmat(i,k) == 1
                            bmat(i,k) = 2;
                        else
                            bmat(i,k) = 1;
                        end
                    end
                else
                    if strcmp(paired,'Unpaired')
                        bmat(i,k) = randi(bio_samp_size(i,k)); %unpaired sampling
                    end
                end
            end
        end

        for i = 1:length(groupChoice)
            for j = 1:length(varX_choice)
                for k = 1:length(Xtime_choice)
                    load_path = strcat(path,OS,num2str(time_val(Xtime_choice(k))),'d',OS,expGroup(groupChoice(i)),OS);
                    load(char(strcat(load_path,biosamp(groupChoice(i),bmat(i,k),k),'.mat')))
                    temp = var_idx{varX_choice(j)};
                    var_temp = eval(temp);
                    Xs(i,j,k,ns) = var_temp{1};
                end
            end
        end

        %Randomly select dependent data and assemble array for model generation
        for i = 1:length(groupChoice)
            for k = 1:length(Ytime_choice)
                if strcmp(paired,'Paired')
                    mmat(i,k) = bmat(i,k); %paired random or selected sampling
                else
                    mmat(i,k) = randi(mech_samp_size(i,k)); %unpaired random sampling
                end
            end
        end

        for i = 1:length(groupChoice)
            for j = 1:length(varY_choice)
                for k = 1:length(Ytime_choice)
                    if strcmp(jk,'Bootstrap')
                        load_path = strcat(path,OS,num2str(time_val(Ytime_choice(k))),'d',OS,expGroup(groupChoice(i)),OS);
                        load(char(strcat(load_path,mechsamp(groupChoice(i),mmat(i,k),Ytime_choice(k)),'_results_2D_obj6.mat')))
                        if varY_choice(j) == 6
                            odsys = properties.loaded.run1(1,2);
                            hsys = properties.loaded.run1(1,3);
                            irsys = (odsys - 2*hsys)/2;
                            Ys(i,j,k,ns) = irsys;
                        elseif varY_choice(j) == 15
                            odsys = properties.loaded.run1(1,2);
                            hsys = properties.loaded.run1(1,3);
                            oddia = properties.loaded.run1(3,2);
                            hdia = properties.loaded.run1(3,3);
                            idsys = (odsys - 2*hsys);
                            iddia = (oddia - 2*hdia);
                            Psys = properties.loaded.run1(1,1);
                            Pdia = properties.loaded.run1(3,1);
                            Dist = (idsys-iddia)/(iddia*(Psys-Pdia))*7500.6;
                            Ys(i,j,k,ns) = Dist;
                        else
                            temp = var_idx{varY_choice(j)};
                            var_temp = eval(temp);
                            Ys(i,j,k,ns) = var_temp;
                        end
                    elseif strcmp(jk,'Jackknife')
                        clear Ys_temp
                        for z = 1:mech_samp_size(groupChoice(i),Ytime_choice(k))
                            sampname = mechsamp(groupChoice(i),z,Ytime_choice(k));
                            celltest = isnan(sampname{1});
                            if z ~= mmat(i,k) && ~any(celltest)
                                load_path = strcat(path,OS,num2str(time_val(Ytime_choice(k))),'d',OS,expGroup(groupChoice(i)),OS);
                                load(char(strcat(load_path,mechsamp(groupChoice(i),z,Ytime_choice(k)),'_results_2D_obj6.mat')))
                                if varY_choice(j) == 6
                                    odsys = properties.loaded.run1(1,2);
                                    hsys = properties.loaded.run1(1,3);
                                    irsys = (odsys - 2*hsys)/2;
                                    Ys_temp(z) = irsys;
                                elseif varY_choice(j) == 15
                                    odsys = properties.loaded.run1(1,2);
                                    hsys = properties.loaded.run1(1,3);
                                    oddia = properties.loaded.run1(3,2);
                                    hdia = properties.loaded.run1(3,3);
                                    idsys = (odsys - 2*hsys);
                                    iddia = (oddia - 2*hdia);
                                    Psys = properties.loaded.run1(1,1);
                                    Pdia = properties.loaded.run1(3,1);
                                    Dist = (idsys-iddia)/(iddia*(Psys-Pdia))*7500.6;
                                    Ys_temp(z) = Dist;
                                else
                                    temp = var_idx{varY_choice(j)};
                                    var_temp = eval(temp);
                                    Ys_temp(z) = var_temp;
                                end
                            else
                                Ys_temp(z) = NaN;
                            end
                        end
                        Ys(i,j,k,ns) = nanmean(Ys_temp);    
                    end
                end
            end
        end

        cd(strcat(path,OS,'nway320 exchange'))
        Xs0(:,:,1,ns) = Xs(:,:,1,ns);
        Ys0(:,:,1,ns) = Ys(:,:,1,ns);
        
        %Pretreatment of data prior to centering and scaling
        for i = 1:length(groupChoice)
            for j = 1:length(varX_choice)
                Xspre(i,j,:,ns) = Xs(i,j,:,ns)-Xs0(i,j,1,ns);
            end
            for j = 1:length(varY_choice)
                Yspre(i,j,:,ns) = Ys(i,j,:,ns)-Ys0(i,j,1,ns);
            end
        end
        
        %Centering and scaling of randomly sampled data
        [Xs_norm_temp(:,:,:,ns),mXs,sXs]=nprocess(permute(Xspre(:,:,:,ns),[1 3 2]), meanctr, scalemechdata);
        [Ys_norm_temp(:,:,:,ns),mYs,sYs]=nprocess(permute(Yspre(:,:,:,ns),[1 3 2]), meanctr, scalemechdata);

        Xs_norm(:,:,:,ns) = permute(Xs_norm_temp(:,:,:,ns),[1 3 2]);
        Ys_norm(:,:,:,ns) = permute(Ys_norm_temp(:,:,:,ns),[1 3 2]);
        mXs_temp = mXs(2); sXs_temp = sXs(2); mYs_temp = mYs(2); sYs_temp = sYs(2);
        mXs(2) = mXs(3); mXs(3) = mXs_temp;
        sXs(2) = sXs(3); sXs(3) = sXs_temp;
        mYs(2) = mYs(3); mYs(3) = mYs_temp;
        sYs(2) = sYs(3); sYs(3) = sYs_temp;

        %Model generation with number of LVs used in model of mean data
        [XNfactors,YNfactors,CoreN,BN,ypredN,ssxN,ssyN] = npls(Xs_norm(:,:,:,ns),Ys_norm(:,:,:,ns),numcomp,NaN);
        [ycalpredN,TN,ssxN,XresN] = npred(Xs_norm(:,:,:,ns),fac,XNfactors,YNfactors,CoreN,BN,0);
        if ndims(Xs_norm(:,:,:,ns)) > 2
            ycalpredN = reshape(ycalpredN,numsamps,numvars,numtimepts);  %model calibration performance
        end
        calibratedN = nprocess(ycalpredN,meanctr,[0 1 0], ...
            mYs,sYs,-1);
        [ResultsN(ns), ModelN] = ncrossreg2(Xs_norm(:,:,:,ns),Ys_norm(:,:,:,ns),fac,any(meanctr==1));
        xvalypredN_pre = shiftdim(ResultsN(ns).ypred(fac,:,:,:),1);
        xvalypredN = nprocess(shiftdim(ResultsN(ns).ypred(fac,:,:,:),1), ...
            meanctr,[0 1 0],mYs,sYs,-1);  %model cross-validated predictions
        for i = 1:length(groupChoice)
            for j = 1:length(varY_choice)
                xvalypredN(i,j,:) = xvalypredN(i,j,:)+Ys0(i,j,1,ns);
                calibratedN(i,j,:) = calibratedN(i,j,:)+Ys0(i,j,1,ns);
            end
        end

        xvalypredNs(:,:,:,ns) = xvalypredN;
        xvalypredNs_pre(:,:,:,ns) = xvalypredN_pre;
        calibratedNs(:,:,:,ns) = calibratedN;
        
        RMSE_diff = ResultsN(ns).RMSEP(2:fac) - ResultsN(ns).RMSEP(1:fac-1);
        min_ind = find(RMSE_diff>=0,1,'first');
        if isempty(min_ind)
            min_ind = fac;
        end
        
        %run corrected model with appropriate number of LVs based on
        %minimum value of RMSE
        [XCfactors,YCfactors,CoreC,BC,ypredC,ssxC,ssyC] = npls(Xs_norm(:,:,:,ns),Ys_norm(:,:,:,ns),min_ind,NaN);
        [ycalpredC,TC,ssxC,XresC] = npred(Xs_norm(:,:,:,ns),min_ind,XCfactors,YCfactors,CoreC,BC,0);
        if ndims(Xs_norm(:,:,:,ns)) > 2
            ycalpredC = reshape(ycalpredC,numsamps,numvars,numtimepts);  %model calibration performance
        end
        calibratedC = nprocess(ycalpredC,meanctr,[0 1 0], ...
            mYs,sYs,-1);
        [ResultsC(ns), ModelC] = ncrossreg2(Xs_norm(:,:,:,ns),Ys_norm(:,:,:,ns),min_ind,any(meanctr==1));
        xvalypredC_pre = shiftdim(ResultsC(ns).ypred(min_ind,:,:,:),1);
        xvalypredC = nprocess(shiftdim(ResultsC(ns).ypred(min_ind,:,:,:),1), ...
            meanctr,[0 1 0],mYs,sYs,-1);  %model cross-validated predictions
        for i = 1:length(groupChoice)
            for j = 1:length(varY_choice)
                xvalypredC(i,j,:) = xvalypredC(i,j,:)+Ys0(i,j,1,ns);
                calibratedC(i,j,:) = calibratedC(i,j,:)+Ys0(i,j,1,ns);
            end
        end

        xvalypredCs(:,:,:,ns) = xvalypredC;
        xvalypredCs_pre(:,:,:,ns) = xvalypredC_pre;
        calibratedCs(:,:,:,ns) = calibratedC;       

        cd(path)

        shuff_X_scores(ns,:,:) = XNfactors{1};
        shuff_Xvar_weights(ns,:,:)=XNfactors{2};
        shuff_Xtime_weights(ns,:,:)=XNfactors{3};
        shuff_Y_scores(ns,:,:) = YNfactors{1};
        shuff_Yvar_weights(ns,:,:)=YNfactors{2};
        shuff_Ytime_weights(ns,:,:)=YNfactors{3};
        
        if min_ind < fac
            shuff_XC_scores(ns,:,1:min_ind) = XCfactors{1};
            shuff_XCvar_weights(ns,:,1:min_ind)=XCfactors{2};
            shuff_XCtime_weights(ns,:,1:min_ind)=XCfactors{3};
            shuff_YC_scores(ns,:,1:min_ind) = YCfactors{1};
            shuff_YCvar_weights(ns,:,1:min_ind)=YCfactors{2};
            shuff_YCtime_weights(ns,:,1:min_ind)=YCfactors{3};
            
            shuff_XC_scores(ns,:,min_ind+1:fac) = NaN;
            shuff_XCvar_weights(ns,:,min_ind+1:fac)=NaN;
            shuff_XCtime_weights(ns,:,min_ind+1:fac)=NaN;
            shuff_YC_scores(ns,:,min_ind+1:fac) = NaN;
            shuff_YCvar_weights(ns,:,min_ind+1:fac)=NaN;
            shuff_YCtime_weights(ns,:,min_ind+1:fac)=NaN;
        else
            shuff_XC_scores(ns,:,:) = XCfactors{1};
            shuff_XCvar_weights(ns,:,:)=XCfactors{2};
            shuff_XCtime_weights(ns,:,:)=XCfactors{3};
            shuff_YC_scores(ns,:,:) = YCfactors{1};
            shuff_YCvar_weights(ns,:,:)=YCfactors{2};
            shuff_YCtime_weights(ns,:,:)=YCfactors{3};
        end

        %Sign switching following methods for null model
        for f = 1:fac
            for i = 1:length(groupChoice)
                if Xfactors{1}(i,f).*XNfactors{1}(i,f) < 0
                    tsign(ns,i,f) = -1;
                else
                    tsign(ns,i,f) = 1;
                end
            end
        end

        for f = 1:fac
            if length(find(tsign(ns,:,f) == -1)) > floor(length(groupChoice)/2)
                shuff_X_scores(ns,:,f) = -shuff_X_scores(ns,:,f);
                shuff_Y_scores(ns,:,f) = -shuff_Y_scores(ns,:,f);
                shuff_Xvar_weights(ns,:,f) = -shuff_Xvar_weights(ns,:,f);
                shuff_Yvar_weights(ns,:,f) = -shuff_Yvar_weights(ns,:,f);
            end
            
            orig_sign = sign(Xfactors{1}(:,f).*Yfactors{1}(:,f));
            numflip = sign(shuff_X_scores(ns,:,f).*shuff_Y_scores(ns,:,f));
            
            if length(find(orig_sign.*numflip' == -1)) > floor(length(groupChoice)/2)
                shuff_Y_scores(ns,:,f) = -shuff_Y_scores(ns,:,f);
                shuff_Yvar_weights(ns,:,f) = -shuff_Yvar_weights(ns,:,f);
            end
        end

    end
    
    for i = 1:length(groupChoice)
        for j = 1:length(varY_choice)
            for k = 1:length(Ytime_choice)
                xvalypredNs_mean(i,j,k) = mean(xvalypredNs(i,j,k,:));
                xvalypredNs_pre_mean(i,j,k) = mean(xvalypredNs_pre(i,j,k,:));
                xvalypredNs_std(i,j,k) = std(xvalypredNs(i,j,k,:));
                xvalypredCs_mean(i,j,k) = mean(xvalypredCs(i,j,k,:));
                xvalypredCs_pre_mean(i,j,k) = mean(xvalypredCs_pre(i,j,k,:));
                xvalypredCs_std(i,j,k) = std(xvalypredCs(i,j,k,:));
            end
        end
    end
    
    pred_temp = zeros(fac,length(groupChoice),length(varY_choice),length(Ytime_choice),numsims);
    %Calculate RMSE of mean predictions
    for ns = 1:numsims
        pred_temp(:,:,:,:,ns) = ResultsN(ns).ypred;
        RMSE_temp(ns,:) = ResultsN(ns).RMSEP;
    end
    pred_RMSE(:,:,:,:) = mean(pred_temp,5);
    
    Ey = zeros([fac size(Y)]);
    for i = 1:length(groupChoice)
        for f = 1:fac
            Ypredf = squeeze(pred_RMSE(f,:,:));
            if size(Y,2) == 1
              YpredfOut=Ypredf(i);
            else
              YpredfOut=Ypredf(i,:);
            end
            
            if size(Y,2)==1
              Ey(f,i,:)    = squeeze(Y_norm(i,:))'-squeeze(YpredfOut);
            else
              Ey(f,i,:)    = squeeze(Y_norm(i,:))-squeeze(YpredfOut);
            end
        end
    end
    
    for f=1:fac
      id=find(~isnan(Ey(f,:,:,:)));
      PRESS(f) = sum(Ey(f,id).^2);
    end
    RMSEP = sqrt(PRESS/length(groupChoice));
    RMSEP_mean = mean(RMSE_temp,1);
    
end

%--------------------------------------------------------------------------
%Calculate average and standard deviation of scores and weights
%--------------------------------------------------------------------------

if ~strcmp(paired,'None')
    for f = 1:fac
        for i = 1:length(groupChoice)
            Xsc_mean(i,f) = nanmean(shuff_X_scores(:,i,f));
            Xsc_std(i,f) = nanstd(shuff_X_scores(:,i,f));
            if ~strcmp(nullPLS,'Yes')
                XCsc_mean(i,f) = nanmean(shuff_XC_scores(:,i,f));
                XCsc_std(i,f) = nanstd(shuff_XC_scores(:,i,f));
            end
        end

        for i = 1:length(groupChoice)
            Ysc_mean(i,f) = nanmean(shuff_Y_scores(:,i,f));
            Ysc_std(i,f) = nanstd(shuff_Y_scores(:,i,f));
            if ~strcmp(nullPLS,'Yes')
                YCsc_mean(i,f) = nanmean(shuff_YC_scores(:,i,f));
                YCsc_std(i,f) = nanstd(shuff_YC_scores(:,i,f));
            end
        end

        for j = 1:length(varX_choice)
            Xv_mean(j,f) = nanmean(shuff_Xvar_weights(:,j,f));
            Xv_std(j,f) = nanstd(shuff_Xvar_weights(:,j,f));
            if ~strcmp(nullPLS,'Yes')
                XCv_mean(j,f) = nanmean(shuff_XCvar_weights(:,j,f));
                XCv_std(j,f) = nanstd(shuff_XCvar_weights(:,j,f));
            end
        end

        for j = 1:length(varY_choice)
            Yv_mean(j,f) = nanmean(shuff_Yvar_weights(:,j,f));
            Yv_std(j,f) = nanstd(shuff_Yvar_weights(:,j,f));
            if ~strcmp(nullPLS,'Yes')
                YCv_mean(j,f) = nanmean(shuff_YCvar_weights(:,j,f));
                YCv_std(j,f) = nanstd(shuff_YCvar_weights(:,j,f));
            end
        end

        for k = 1:length(Xtime_choice)
            Xt_mean(k,f) = nanmean(shuff_Xtime_weights(:,k,f));
            Xt_std(k,f) = nanstd(shuff_Xtime_weights(:,k,f));
            if ~strcmp(nullPLS,'Yes')
                XCt_mean(k,f) = nanmean(shuff_XCtime_weights(:,k,f));
                XCt_std(k,f) = nanstd(shuff_XCtime_weights(:,k,f));
            end
        end

        for k = 1:length(Ytime_choice)
            Yt_mean(k,f) = nanmean(shuff_Ytime_weights(:,k,f));
            Yt_std(k,f) = nanstd(shuff_Ytime_weights(:,k,f));
            if ~strcmp(nullPLS,'Yes')
                YCt_mean(k,f) = nanmean(shuff_YCtime_weights(:,k,f));
                YCt_std(k,f) = nanstd(shuff_YCtime_weights(:,k,f));
            end
        end

    end

    Xfac{1}=Xsc_mean;Xfac{2}=Xv_mean;Xfac{3}=Xt_mean;
    DimX = size(X_norm);
    cd(strcat(path,OS,'nway320 exchange'))
    Core_mean{3} = calcore(reshape(X_norm,DimX),Xfac,[],0,1);
    cd(path)
    Tm = Xfac{1};
    Um = Ysc_mean;
    B_mean(1:fac,fac) = inv(Tm'*Tm)*Tm'*Um(:,fac);
end

%--------------------------------------------------------------------------
%Model predictions using average results from random sampling
%--------------------------------------------------------------------------

if ~strcmp(nullPLS,'Yes') && ~strcmp(paired,'None')
    all_samp_data = reshape(xvalypredNs_pre_mean,1,prod(size(xvalypredNs_pre_mean)));
    R_samp = corr(all_samp_data',all_Y_pre_data','Type','Pearson');
    rho_samp = corr(all_samp_data',all_Y_pre_data','Type','Spearman');

    all_adj_data = reshape(xvalypredCs_pre_mean,1,prod(size(xvalypredCs_pre_mean)));
    R_adj = corr(all_adj_data',all_Y_pre_data','Type','Pearson');
    rho_adj = corr(all_adj_data',all_Y_pre_data','Type','Spearman');
end

%--------------------------------------------------------------------------
%Plot scores and weights of latent variables
%--------------------------------------------------------------------------
cd(path)

plsPlots_clean
```
