## Supplementary File S1 for "Robust latent-variable interpretation of *in vivo* regression models by nested resampling": shufflematrix.m

```
function sMx = shufflematrix(X)
% sufflematrix randomizes the rows and columns in a matrix

linX=reshape(X,size(X,1)*size(X,2),1);
shuffleindex=randperm(size(X,1)*size(X,2),size(X,1)*size(X,2));
shuflinX=linX(shuffleindex);
sMx=reshape(shuflinX,size(X,1),size(X,2));
```
