## Supplementary File S1 for "Robust latent-variable interpretation of *in vivo* regression models by nested resampling": tight_subplot.m

```
function [ha, pos] = tight_subplot(Nh, Nw, gap, marg_h, marg_w)

% tight_subplot creates "subplot" axes with adjustable gaps and margins
%
% [ha, pos] = tight_subplot(Nh, Nw, gap, marg_h, marg_w)
%
%   in:  Nh      number of axes in hight (vertical direction)
%        Nw      number of axes in width (horizontaldirection)
%        gap     gaps between the axes in normalized units (0...1)
%                   or [gap_h gap_w] for different gaps in height and width 
%        marg_h  margins in height in normalized units (0...1)
%                   or [lower upper] for different lower and upper margins 
%        marg_w  margins in width in normalized units (0...1)
%                   or [left right] for different left and right margins 
%
%  out:  ha     array of handles of the axes objects
%                   starting from upper left corner, going row-wise as in
%                   subplot
%        pos    positions of the axes objects
%
%  Example: ha = tight_subplot(3,2,[.01 .03],[.1 .01],[.01 .01])
%           for ii = 1:6; axes(ha(ii)); plot(randn(10,ii)); end
%           set(ha(1:4),'XTickLabel',''); set(ha,'YTickLabel','')

% Pekka Kumpulainen 21.5.2012   @tut.fi
% Tampere University of Technology / Automation Science and Engineering


if nargin<3; gap = .02; end
if nargin<4 || isempty(marg_h); marg_h = .05; end
if nargin<5; marg_w = .05; end

if numel(gap)==1; 
    gap = [gap gap];
end
if numel(marg_w)==1; 
    marg_w = [marg_w marg_w];
end
if numel(marg_h)==1; 
    marg_h = [marg_h marg_h];
end

axh = (1-sum(marg_h)-(Nh-1)*gap(1))/Nh; 
axw = (1-sum(marg_w)-(Nw-1)*gap(2))/Nw;

py = 1-marg_h(2)-axh; 

% ha = zeros(Nh*Nw,1);
ii = 0;
for ih = 1:Nh
    px = marg_w(1);
    
    for ix = 1:Nw
        ii = ii+1;
        ha(ii) = axes('Units','normalized', ...
            'Position',[px py axw axh], ...
            'XTickLabel','', ...
            'YTickLabel','');
        px = px+axw+gap(2);
    end
    py = py-axh-gap(1);
end
if nargout > 1
    pos = get(ha,'Position');
end
ha = ha(:);
```
